# A Palearctic divide, niche conservatism and host-fungal endophyte interactions shaped the phylogeography of the grass *Brachypodium sylvaticum*

**DOI:** 10.1101/2025.06.24.661274

**Authors:** María de los Ángeles Decena, Miguel Campos Cáceres, Diana Calderón Pardo, Valeriia Shiposha, Marina Olonova, Ernesto Pérez-Collazos, Pilar Catalán

**Affiliations:** Departamento de Ciencias Agrarias y del Medio Natural. Escuela Politécnica Superior de Huesca. Universidad de Zaragoza. C/ Carretera de Cuarte Km 1. E-22071 Huesca. Spain; Grupo de Bioquímica, Biofísica y Biología Computacional (BIFI, UNIZAR), Unidad Asociada al CSIC; Institute of Biology, Tomsk State University, Tomsk 634050, Russia

**Keywords:** *Brachypodium sylvaticum* complex, *Epichloë sylvatica*, grass-endophyte interactions, coevolution of holobionts, ecological adaptation, glacial refugia, microspeciation, phylogeography

## Abstract

*Brachypodium sylvaticum* is a perennial woodland grass selected as model species for perenniality, which is widely distributed across the Palearctic. This plant forms a symbiosis with the endophytic fungus *Epichloë sylvatica*. We integrate whole-genome phylogenomics, plastome analysis, environmental niche modeling (ENM), and coevolutionary analyses to investigate the diversification of *B. sylvaticum* and its fungal symbiont. Using 94 representative individuals spanning Eurasia and North Africa, we recovered two deeply divergent sister lineages (Eastern and Western Palearctic), with cytonuclear discordances suggesting historical plastid capture events in the western group. Admixture analysis revealed four genetic clusters, including signatures of secondary contact and hybridization in the Western lineage. Filtered ITS sequences of *E. sylvatica* recovered from holobiont genome skimming reads enabled phylogenetic reconstruction, revealing two fungal clades that broadly mirror their host’s evolutionary history in the West. PACo and ParaFit analyses supported partial co-divergence between hosts and endophytes. ENM projections identified climatically stable glacial refugia for both *B. sylvaticum* main lineages during the Last Glacial Maximum and asymmetric postglacial expansion, with moderate niche shifts in the West and stronger turnover in the East. Evidence of niche overlap and similarity indicated niche conservatism among clades, suggesting that geographic isolation, rather than adaptive divergence, was the primary driver of lineage splitting. IBD and IBE patterns significantly influenced divergences in the Western, but not the Eastern, group, highlighting contrasting demographic and ecological dynamics. Our results provide the first evidence of coevolutionary and ecological structuring in *B. sylvaticum*–*E. sylvatica* holobionts across their Western native range.

## 1. Introduction

*Brachypodium sylvaticum*, a perennial Palearctic grass, is one of the most widely distributed species of the small cold seasonal genus *Brachypodium*, which includes about 23 world species (Catalan et al., 2016, 2023; Scholthof et al., 2018). *Brachypodium sylvaticum* is native to Europe, Asia, and North Africa (Catalán et al., 2023) and was introduced in western North America, becoming an invasive species (Rosenthal et al., 2008; Roy et al., 2011). This grass has gained attention in plant research as a model species for perennial grasses due to its adaptability and wide distribution across various habitats (Carroll & Somerville, 2009; Dohleman & Long, 2009; Glover et al., 2010). Its significance as a model plant for perenniality-to-annuality transitions has been underscored by extensive genomic (reference genomes Ain1 and Sin1) and biological resources (Gordon et al., 2015; Steinwand et al., 2013; Lei et al., 2024; Phytozome v13, https://phytozome-next.jgi.doe.gov/ ).

*Brachypodium sylvaticum* is characterized by its short and slender rhizomes, nodding panicles, densely hairy habit, and long-awned lemmas (Schippmann, 1991). The species is self-compatible and thrives in mesic and humid forest habitats (Khan & Stace, 1999; Steinwand et al., 2013). Alongside typical *B. sylvaticum*, four microtaxa (*B. breviglume, B. kurilense, B. spryginii, B. miserum*) have been identified as close relatives, sharing certain morphological characteristics and distribution areas with *B. sylvaticum sensu stricto* (*s.s.*) (Catalán et al., 2023; Tzvelev & Probatova, 2019). The entire *B. sylvaticum* complex is constituted by diploid species, with a consistent chromosome number of *2n* = 2x = 18 (x = 9) (Catalán et al., 2016; Díaz-Pérez et al., 2018). Another closely related taxon is the perennial Mediterranean *B. glaucovirens*, also diploid but with a different chromosome number of *2n* = 2x = 16 (x = 8). *B. glaucovirens* exhibits overall features similar to those of *B. sylvaticum* (short rhizome, large awns) and *B. pinnatum* (bright green leaves, broad leaf ribs, erect panicle) (Schippmann, 1991). Previous studies suggested that *B. glaucovirens* may have a hybrid origin (Scholz, 2007), and plastome-based phylogenies confirmed that *B. sylvaticum s.s.* served as the maternal progenitor for this taxon (Catalán et al., 2023). Despite its wide distribution and significance as a model organism, the *B. sylvaticum* complex remains an active area of research (Catalán et al., 2023; Keng, 1982; Tzvelev, 2015). Some of these micro-taxa exhibit large disjunct geographic distributions in their native Palearctic area, suggesting genetic isolation and evolutionary divergence, warranting further phylogenetic and genomic investigation.

Phylogenetic studies based on nuclear and plastid loci consistently positioned the *B. sylvaticum* complex within the recently evolved core-perennial clade of *Brachypodium* (Catalán et al., 2016; Catalán & Olmstead, 2000; Díaz-Pérez et al., 2018). Molecular dating suggested that the *B. sylvaticum s.s* lineage originated in the Mid-Late Pleistocene (Díaz-Pérez et al., 2018), coinciding with major climatic oscillations that likely influenced its evolutionary trajectory. Comparative chromosome barcoding has further detailed the karyotypic profiles of *B. sylvaticum* and its close congeners, providing insights into their intermediate karyotype evolutionary history and chromosome dynamics (Lusinska et al., 2019; Sancho et al., 2022).

*Brachypodium sylvaticum* maintains a ubiquitous and highly stable symbiosis with its *Epichloë sylvatica* fungal endophyte (Leuchtmann & Schardl, 2022). This association is predominantly characterized by vertical transmission of the endophyte through the host seeds, ensuring the systemic colonization of new generations of host plants without visible symptoms (Meijer & Leuchtmann, 1999). However, *E. sylvatica* also retains the capacity for sexual reproduction via stromata formation and horizontal transmission, which can inhibit host flowering, though such events are rare and typically confined to specific microhabitats (Brem & Leuchtmann, 2001). This dual transmission strategy underscores the evolutionary flexibility of *E. sylvatica*, oscillating between predominantly mutualistic and, under certain conditions, antagonistic interactions with its host grass.

The ecological advantages of this mutualistic symbiosis are of great importance for the holobiont. Experimental feeding trials have demonstrated that *E. sylvatica* infection enhances herbivore resistance in *B. sylvaticum* (Brem & Leuchtmann, 2001). While the precise chemical basis of deterrence to herbivores remains unclear, *Epichloë* endophytes in related grasses are known to influence host physiology through metabolic alterations or secondary metabolite (alkaloids) production (Vikuk et al., 2019; Gundel et al., 2017; Saikkonen et al., 2016). Moreover, *Epichloë* endophytes in other grass species confer additional adaptive advantages, such as increased drought tolerance and pathogen resistance, raising the possibility that *B. sylvaticum* benefits similarly (Vázquez de Aldana et al., 2015). Given the nearly ubiquitous presence of *E. sylvatica* in natural populations of *B. sylvaticum*, its role as a key ecological and evolutionary factor in the phylogeography and adaptive strategies of its host species should not be overlooked.

Environmental niche modeling (ENM) has been extensively used to predict the potential distribution ranges of species and to evaluate their ecological niches under different evolutionary and climatic scenarios (Purcell & Stigall, 2021; Vásquez-Cruz et al., 2025; Žerdoner Čalasan et al., 2021; Zhang et al., 2024). These models have provided valuable insights into processes such as ecological speciation, hybridization niches, species isolation, , and the impacts of climate change on biodiversity (Arenas-Castro & Sillero, 2021; López-Alvarez et al., 2015). The Pleistocene climatic cycles, characterized by alternating glacial and interglacial periods, profoundly influenced the demographic history and distribution patterns of Palearctic species (Friesen et al., 2016; Hewitt, 2004; Zhang et al., 2024).

Despite the considerable genomic and evolutionary knowledge gained from *B. sylvaticum*, no phylogeographic and environmental niche modeling analyses have been performed for this species complex, nor have grass-endophyte holobionts been considered as an evolutionary unit. In this study, we sampled *B. sylvaticum* accessions across its native Eurasian and North African range, retrieved genetic data from its ubiquitous *E. sylvatica* endophytes, and integrated phylogenomic, population genetic, coevolutionary, and environmental niche modeling approaches to address key evolutionary questions. We reconstructed the environmental niches of the *B. sylvaticum* across three geoclimatic scenarios spanning the Last Glacial Maximum (LGM, ∼20 kya) to the present. These models aimed to identify potential glacial refugia, evaluate oceanic *vs*. continental occurrence patterns, and infer postglacial range shifts that may have shaped the current distribution of the *B. sylvaticum* complex species. Our objectives were to: (i) characterize the genomic diversity and phylogenetic relationship among *B. sylvaticum* complex populations using nuclear genome and plastome data, (ii) assess the coevolutionary dynamics between *B. sylvaticum* and its endophyte *E. sylvatica*, (iii) reconstruct historical and contemporary niche distribution patterns to identify glacial refugia and postglacial expansion routes, and (iv) synthesize these findings to elucidate the interplay between niche dynamics and grass-fungal interactions in shaping the plant species’ phylogeography.

## 2. Materials and methods

The study included 94 representative samples of the *B. sylvaticum* complex, strategically collected to cover extensive genetic, geographic, and ecological diversity (Figure 1, Appendix 1). Genome skimming sequencing was performed from total DNA, and whole plastomes were assembled and annotated using Novoplasty and GeSeq, respectively. Reads were mapped to reference genomes of *B. sylvaticum* and its fungal endophyte *E. sylvatica*, filtered, and SNPs were called using a custom bioinformatics pipeline involving Minimap2, Samtools, and BCFtools. For phylogenomic analysis, maximum likelihood trees were built using nuclear SNPs and plastome sequences with IQ-TREE2, and divergence dates were estimated using TreePL with specified calibrations and biogeographic patterns analyzed with BioGeoBEARS. Population genomic structure was assessed using ADMIXTURE, and potential cytonuclear discordance was analyzed using generalized Robinson–Foulds distance. To study plant-endophyte coevolution, PACo and ParaFit analyses were applied. Genomic diversity metrics (Ho, He, Fis) were computed, and isolation by distance (IBD) and environment (IBE) patterns were evaluated using distance based redundancy analyses (dbRDA). Environmental niche modeling (ENM) was conducted using Maxent and niche comparisons were performed with statistical tests to assess niche conservatism and divergence among *B. sylvaticum* complex groups (see expanded Material and Methods in Appendix 2).

**Figure 1.**
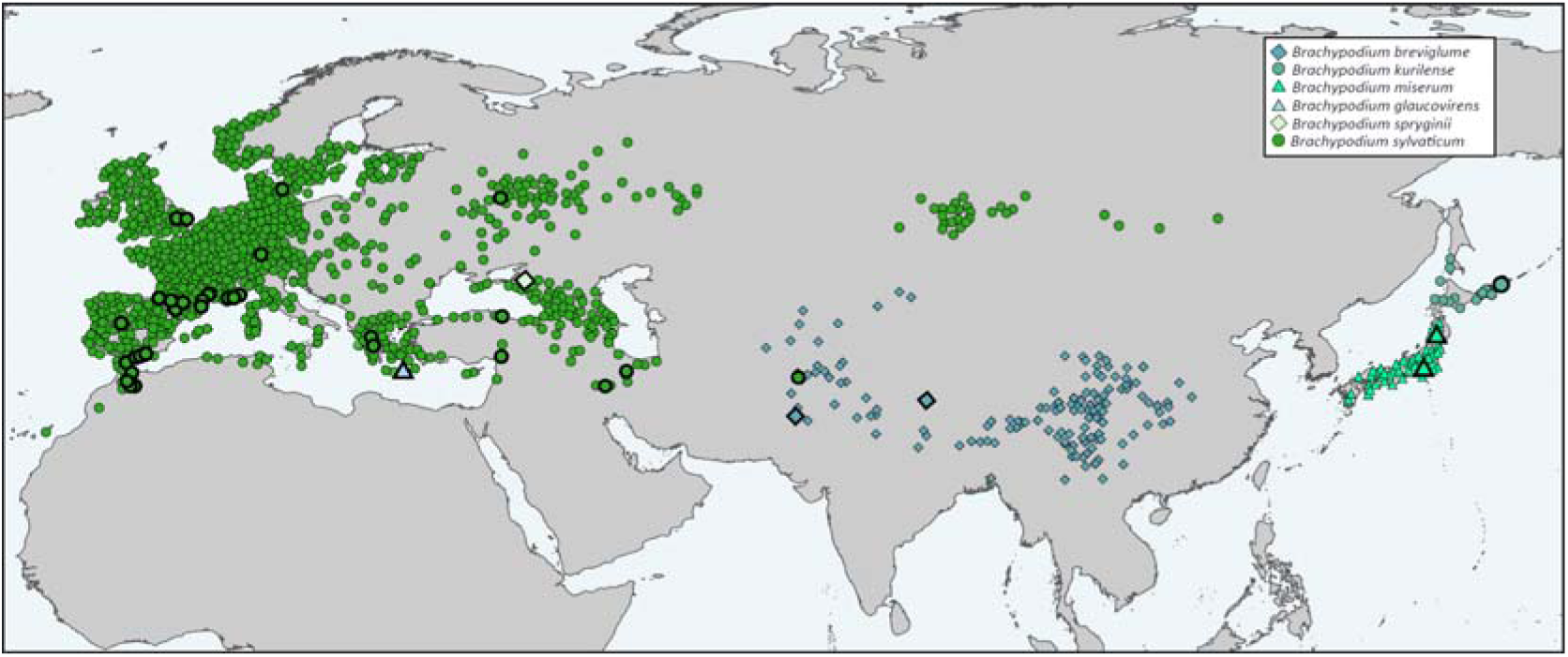
Geographic distribution map of the studied *B. sylvaticum* complex samples in the Palearctic. The map shows georeferenced locations obtained from sampled populations and from GBIF) (see Table S1, Appendix 1). Color and shape codes corresponding to the different taxa (Western group: *B. sylvaticum s.s.*, *B. spryginii, B. glaucovirens*; Eastern group: *B. breviglume, B. miserum, B. kurilense*) are indicated in the chart. Bold symbols correspond to individuals included in the genomic analyses, while non-bold symbols represent additional occurrences retrieved from GBIF.

## 3. Results

### 3.1 Nuclear and plastome datasets, phylogenomics, and bio/phylogeography of the Brachypodium sylvaticum complex

The total number of raw Illumina reads across all samples ranged from 10,904,196 to 48,966,188 (Table S2). After filtering and mapping the reads to the *B. sylvaticum* Ain-1 nuclear reference genome, the number of mapped reads varied between 10,246,338 and 45,697,368. The total number of genes recovered with at least 90% of their sequence information ranged between 8,020 for *B. kurilense* to 24,287 for *B. sylvaticum* Bsyl36H (Table S2). The initial whole-genome data matrix comprised 8,372,248 positions; after discarding gaps and applying a linkage disequilibrium (LD) filtering, we obtained a final dataset of 160,124 nuclear SNPs, uniformly distributed across the nuclear chromosomes of the *B. sylvaticum* complex samples, that were used in downstream analyses. Syntenic SNPs from other diploid Brachypodium species, were retrieved from their respective genomic resources (Table S1). The strongly supported *B. sylvaticum* complex nuclear ML phylogenomic tree based on the nuclear SNP dataset revealed two sister lineages: Eastern and Western Palearctic (Figure 2). In the Eastern clade, the B. breviglume lineages were basal and paraphyletic, and the more recently evolved *B. kurilense* lineage was sister to the B. miserum clade. Within the Western clade, *B. glaucovirens* was recovered as the earliest diverging lineage, followed by two well-supported sister subclades. Subclade A included *B. spryginii* and *B. sylvaticum s.s.* lineages distributed across western, central, and eastern Europe as well as Afghanistan, while subclade B comprised *B. sylvaticum s.s.* lineages from Greece, Denmark, British Islands, Spain and France (Pyrenees and Alps), Morocco, Turkey, and Iran. Most individuals from the same population or region clustered together, and the topology retrieved some geographical structure, although some individuals from the same population nested in separate clades (Figure 2).

**Figure 2.**
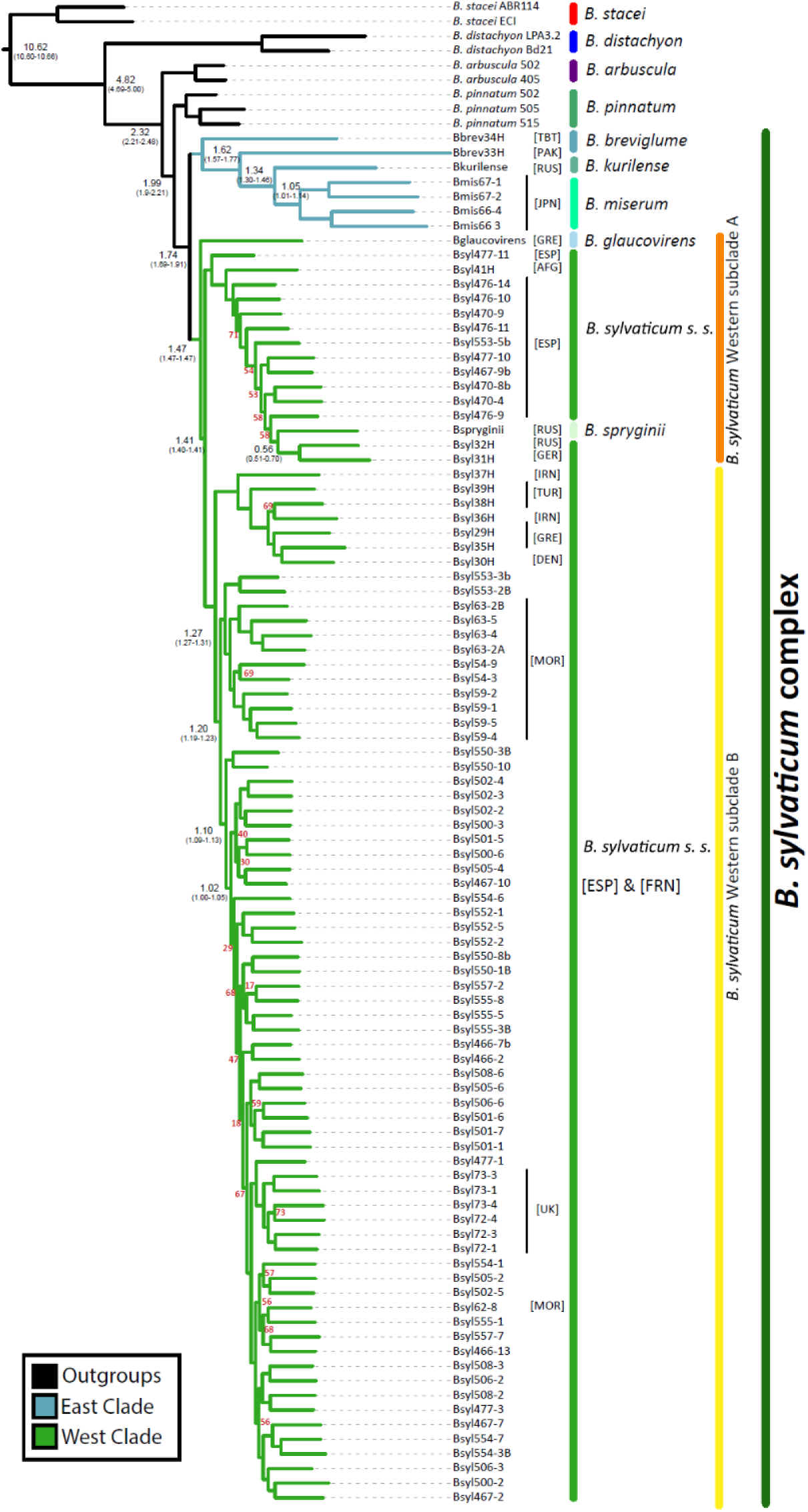
Nuclear Maximum Likelihood phylogenetic tree of the *Brachypodium sylvaticum* complex. Blue branches correspond to the Eastern Palearctic lineages (*B. breviglume, B. miserum, B. kurilense*), green branches to the Western Palearctic lineages (*B. sylvaticum s.s., B. spryginii*, *B. glaucovirens*), and black branches to outgroups. All branches showed bootstrap support values ≥80% except those indicated in red. Node ages estimated with TreePL are indicated in black, along with their 95% confidence intervals. [AFG] Afghanistan; [DEN] Denmark; [ESP] Spain; [GRE] Greece; [IRN] Iran; [JPN] Japan; [MOR] Morocco; [PAK] Pakistan; [RUS] Russia; [TBT] Tibet; [TUR] Turkey; [UK] United Kingdom (British Islands).

The BioGeoBEARS ancestral range reconstructions inferred from the TreePL dated phylogeny, supported DIVA+J (Dispersal-Vicariance Analysis with jump dispersals) as optimal model for the Eastern and Western clades (Figure 3). Divergence dating indicated that *the B. sylvaticum* complex lineage split from its stem ancestor in the early Pleistocene (Gelasian, ∼1.99 Ma), and the crown node of the complex split into the sister Eastern and Western Palearctic clades in the Calabrian (∼1.74 Ma) (Figure 2). The estimated ages of the main Eastern clade lineages were slightly older than those of the main Western clade lineages. Within the Eastern clade, inferred biogeographic patterns indicated that the crown ancestor and later ancestor of the Eastern group (*B. breviglume*) was distributed in the Himalayas (A, 1.74–1.62 Ma). This was followed by dispersal from the Himalayas to the Asian Pacific coast (CD), and then by vicariance between Japan (C) and the Kuril Islands (D) resulting in *B. miserum* and *B. kurilense* in the middle Calabrian (∼1.05 Ma; Figures 2, 3A). Within the Western clade, the most likely inferred ancestral area was a composite range including ABDE areas (Pyrenees-Alps, Middle East, eastern Mediterranean, and Morocco-southern Spain; Figure 3B). From this wide ancestral range, an estimated peripheral isolation event separated the *B. glaucovirens* lineage in the eastern Mediterranean (D) (∼1.47 Ma; Figures 2, 3B). Subsequently, a vicariance event at ∼1.41 Ma likely split the subclade A, distributed in the Pyrenees-Alps region, from its sister subclade B, distributed in the Middle East - Eastern Mediterranean. Subclade A comprised lineages predominantly associated with Pyrenean populations, which likely experienced three independent dispersal events: i) a first long-distance dispersal (LDD) to the Middle East (B) ∼1.08 Ma; ii) a second LDD to Morocco-southern Spain (E) ∼0.80 Ma; and iii) a third LDD to the eastern Mediterranean (C) ∼0.70 Ma, which led to the divergence of *B. spryginii* and related eastern Mediterranean lineages at ∼0.56 Ma (Figures 2, 3). Subclade B showed a more robust geographic structure, with an inferred early vicariance event separating a Middle Eastern lineage (B) from its sister Morocco-Southern Spain lineage (E) ∼1.27 Ma. In the first group, three independent dispersals were estimated to have occurred between nearby geographic areas (Middle East to Eastern Mediterranean ∼0.90 Ma, Eastern Mediterranean to Middle East ∼0.60 Ma, and Eastern Mediterranean to Western Russia and Central Europe ∼0.50 Ma). Within the second group, two independent colonizations of the Pyrenees-Alps from Morocco-southern Spain were estimated to have occurred ∼1.10 Ma and ∼1.00 Ma, respectively, and more recent independent dispersals from the Pyrenees-Alps to the British Isles ∼0.70 Ma and back to Morocco-southern Spain (∼0.70 Ma, ∼0.70 Ma and ∼0.60 Ma) (Figure 3B).

**Figure 3.**
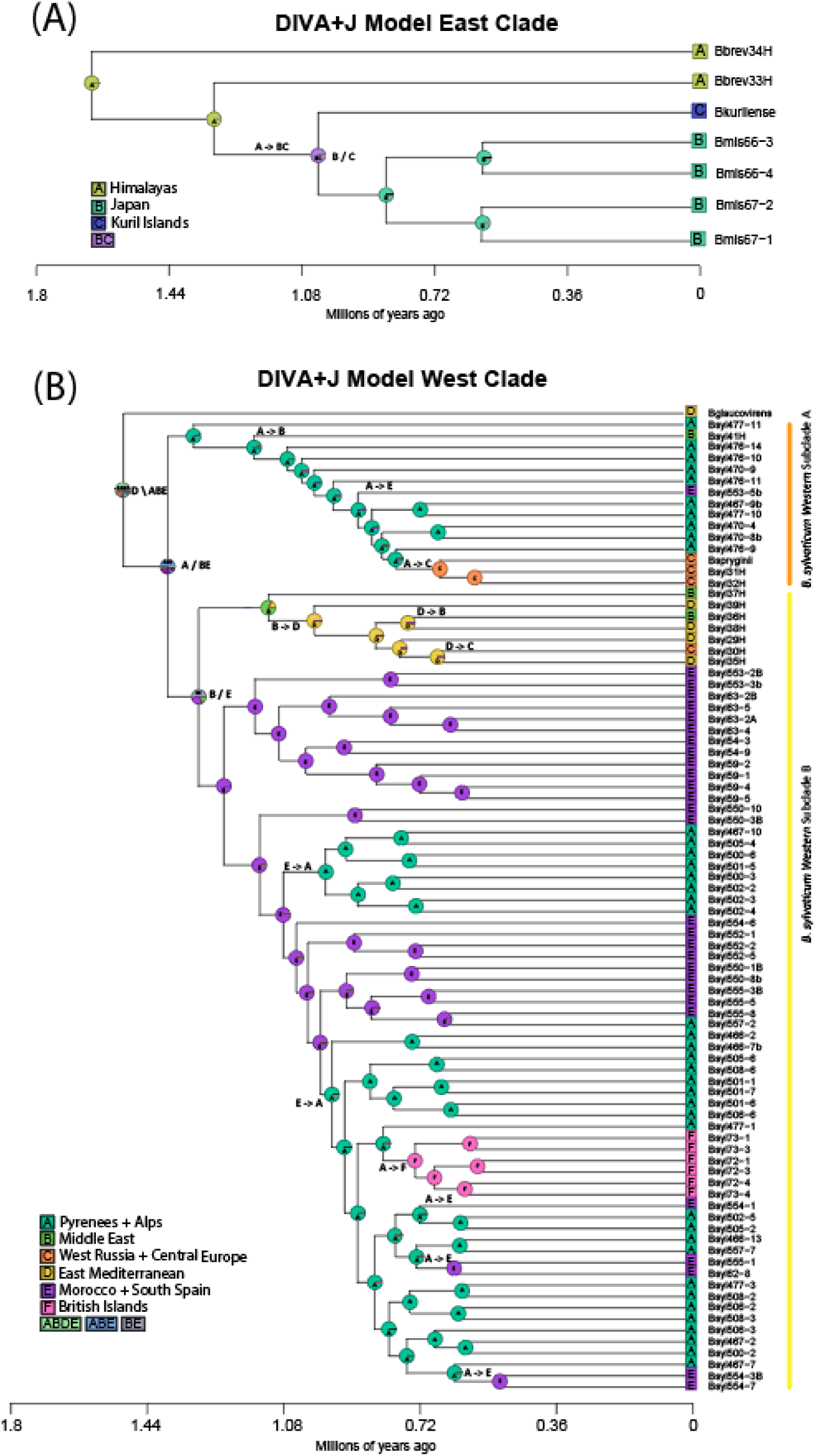
Ancestral ranges and biogeographical events of the *Brachypodium sylvaticum* complex Eastern and Western Palearctic lineages inferred from BioGeoBEARS under DIVA+J models. (A) Optimal model selected for the Eastern Palearctic group; (B) optimal model selected for the Western Palearctic group. Pie charts indicate the relative probability for alternative ancestral ranges. The inferred biogeographic events are indicated at the nodes and branches of the tree (X/Y vicariance; X\Y peripheral isolation; X->Y dispersal).

The *B. sylvaticum* complex whole-plastome alignment included 137,356 positions. The total length of the assembled plastomes ranged from 136,073 to 136,571 bp (Table S2). The plastome sequences of the *B. sylvaticum* complex were highly conserved in terms of synteny and gene number, containing 133 genes (76 protein coding genes, 20 non-redundant tRNAs, four rRNAs in both inverted repeats, four pseudogenes, and two hypothetical open reading frames) (Figure S1). The plastomes of *B. sylvaticum* complex studied exhibited a ∼1,143 bp insertion between the psaI and rbcL genes in the LSC region, corresponding to a rpl23 pseudogene. Additionally, they also showed a ∼301 bp insertion that aligns with a duplicated rps19 gene located between the psbA and trnH genes in the IRb repeat (Figure S1).

The plastome phylogenetic tree of the B. sylvaticum complex (Figure S2) showed a relatively similar topology to that of the nuclear tree (Figure 2) for the major lineages, recovering the basal split of the Eastern and Western Palearctic lineages. However, the *B. pinnatum* lineage was resolved here as sister to the Western Palearctic lineage of *B. sylvaticum*, indicating that the *B. sylvaticum* complex is not monophyletic in the plastome-based reconstruction (Figure S2). The Eastern Palearctic clade displayed the same resolution for its lineages as the nuclear tree, whereas successive divergences within lineages of the Western Palearctic clade differed from those of the nuclear tree (Figures S2, 4). The most ancestral splits of the plastome tree displayed complete support; however, the most recent splits showed poor support (<50% BS) due to the similarity in plastome sequences between these closely related lineages (Figure S2). A cytonuclear discordance analysis performed between the nuclear and plastomic phylogenies using the generalized Robinson-Foulds distance (Smith, 2020) indicated substantial incongruence between both trees (gRF = 0.7286; Figure 4).

**Figure 4.**
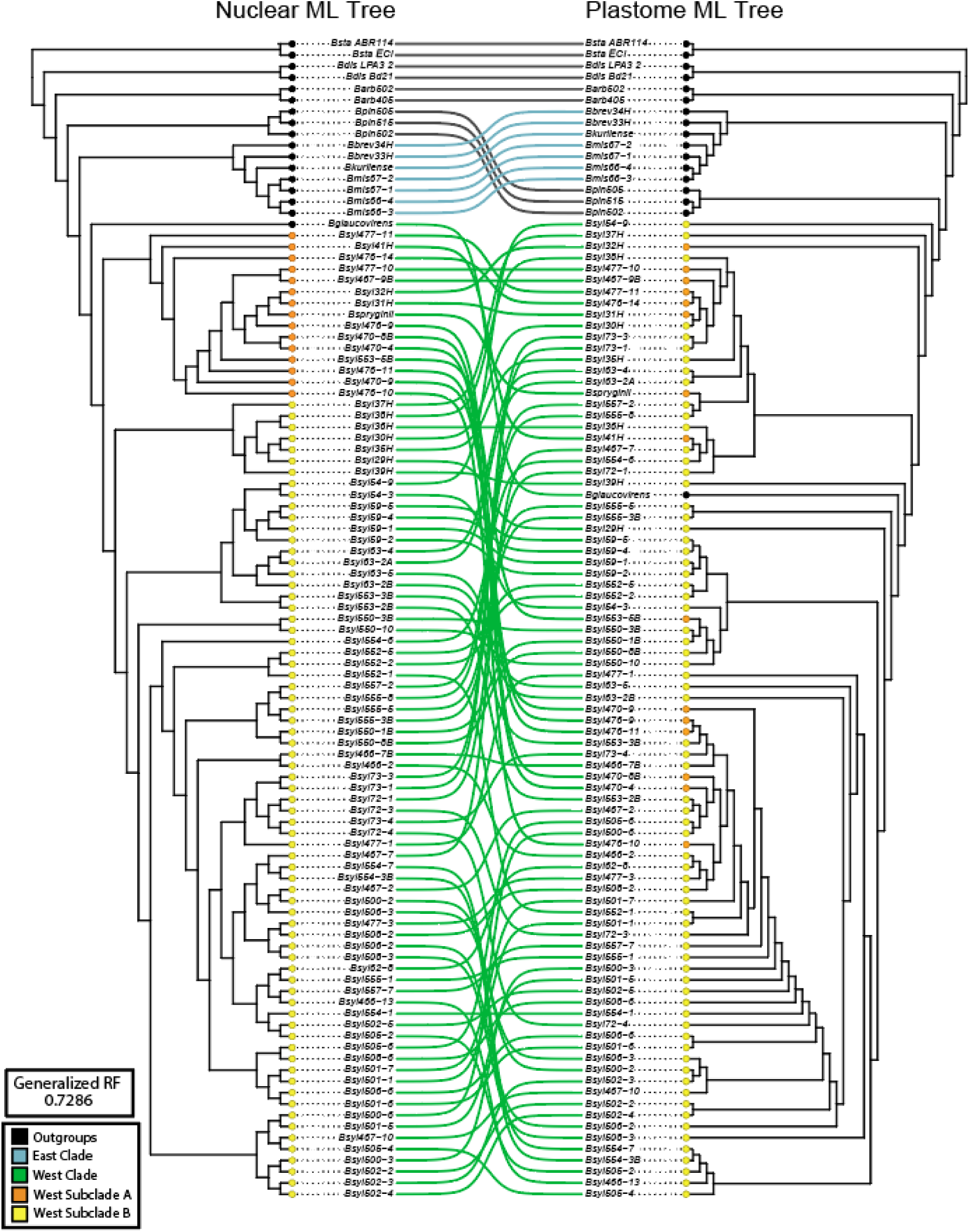
Cophylogenetic comparison of the *Brachypodium sylvaticum* complex nuclear and plastome phylogenetic trees. Blue branches represent the Eastern Palearctic group, green branches the Western Palearctic group, and black branches the outgroups. Colored circles represent subclade A (Orange) and subclade B (Yellow) of the Western group (see Figure 2). Topological incongruence between nuclear and plastid trees was quantified using the generalized Robinson–Foulds distance.

### 3.2 Detection of fungal endophytes and phylogeny of the Epichloë sylvatica symbionts

Evidence of symbiosis with *Epichloë sylvatica* was found in 83 of the 94 samples of the *B. sylvaticum* complex studied (Table S2). The total number of *E. sylvatica* reads ranged from 15,224 to 564,201, with the percentage coverage of the 35S rDNA cistron ranging from 63.82% to 89.44% in these samples. All Blast searches indicated that the assembled cistron sequences aligned with the *E. sylvatica* reference genome (Table S2). Among the 11 excluded individuals, six belonged to the Eastern Palearctic clade, specifically *B. breviglume* (Bbrev33H, Bbrev34H) and *B. miserum* (Bmis66-3, Bmis66-4, Bmis67-1, Bmis67-2), and eight samples belonged to the Western Palearctic clade (*B. glaucovirens*, Bsyl554-3B, Bsyl555-8, Bsyl59-2, Bsyl72-1). However, the failure to detect *E. sylvatica* in these samples could be due to the quality of their DNA or a lack of reliable detection, so they were discarded from the analysis.

The ITS phylogenetic tree of *E. sylvatica* revealed a well-to-moderately supported topology composed of two major clades (Figure S3). The first clade (Clade I) included endophytes found on host plants of *B. kurilense*, the only Eastern Palearctic species where the symbiont was detected, and lineages from Subclade B of *B. sylvaticum* from the Western Palearctic (plus three individuals from Subclade A). The second clade (Clade II) consisted of endophytes found on *B. sylvaticum* from a wide geographic distribution, corresponding to lineages from Subclades A and B in the Western Palearctic (Figure S3). *E. sylvatica* lineages with very low support values (<20% BS) were collapsed into polytomies; these corresponded to samples with very similar or identical ITS sequences (Figure S3).

Potential coevolutionary relationships between *B. sylvaticum* host plants and their *E. sylvatica* endophytes were assessed using the ParaFit and PACo methods. The global ParaFit test, based on the *B. sylvaticum* nuclear consensus phylogenies and *E. sylvatica* ITS tree over 1000 replicates, yielded a significant result, indicating overall phylogenetic congruence between hosts and symbionts (Table S3). However, only 11 individual host–endophyte interactions out of 83 were significant (Figure S4; Table S3), suggesting that congruence at the global level does not extend evenly to all pairwise associations. Interestingly, the majority of significant interactions corresponded to the Western Subclade A of *B. sylvaticum*, which included *B. spryginii* and *B. sylvaticum* s. s. individuals distributed across western, central, and eastern Europe, as well as Afghanistan (Figures 2, S4). In contrast, PACo analysis using 1000 bootstrap phylogenies also indicated statistically significant congruence, although the relatively high residual suggests a moderate level of phylogenetic fit (Table S4). Notably, PACo identified a higher number of consistent individual congruent assemblages, including 50 individual holobionts of *B. sylvaticum* and *E. sylvatica* from the Western Palearctic, 17 of 37 in subclade A and 33 of 46 in subclade B (Figures 5, S5).

**Figure 5.**
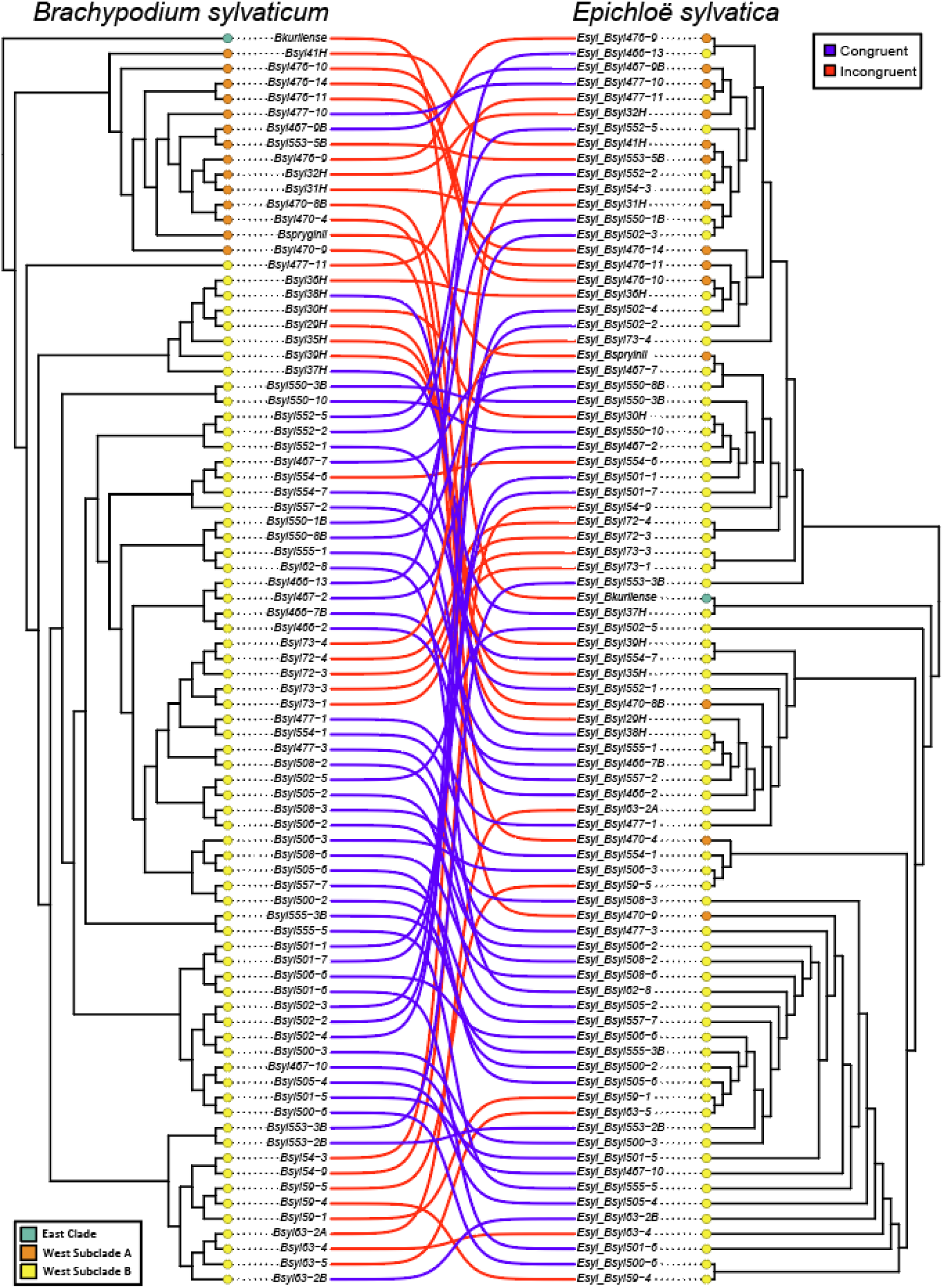
Co-evolutionary analysis of *Brachypodium sylvaticum* and *Epichloë sylvatica* interactions based on Procrustean Approach to Cophylogenetics (PACo) analysis. The figure displays the cophylogeny between the *B. sylvaticum* nuclear phylogenetic tree and the *E. sylvatica* ITS tree. Blue lines indicate congruent branches and red lines mark incongruent associations. Colored circles represent in blue the *B. sylvaticum* complex Eastern Palearctic clade (*B. kurilense*), in orange the *B. sylvaticum* Western Palearctic Subclade A, and in yellow the *B. sylvaticum* Western Palearctic Subclade B.

### 3.3 Population genomics and genomic diversity of Brachypodium sylvaticum complex taxa and groups, and impact of IBD and IBE

The optimal number of population groups (K), determined using the Admixture and StructureSelector method, was K=4 (Figures 6, S6). The first group corresponded to the Eastern Palearctic group (*B. breviglume*, *B. miserum*, *B. kurilense*) (Figure 6, blue), with one sample (Bbrev34H, Pakistan) showing some admixture with group 2. The second group included *B. spryginii* and a subset of *B. sylvaticum* individuals belonging to the Western Palearctic A subclade (Figures 2, 6, orange); these individuals were predominantly from Pyrenean populations, with one individual showing admixture with group 4 (Bsyl477-11, Spain). The third cluster included *B. glaucovirens* and *B. sylvaticum* samples from Morocco, southern Spain, the Middle East, and the eastern Mediterranean, corresponding to the Western Palearctic B subclade (Figures 2, 6 (magenta); it contained samples showing admixture mainly with clusters 1 (Bsyl37H, Iran) and 2 (Bsyl39H, Turkey). The fourth cluster consisted of B. sylvaticum individuals of the Western Palearctic B subclade from the Pyrenees, the Alps, southern Spain, and the British Isles (Figures 2, 6 (green), with one individual (Bsyl550-10, Spain) showing some admixture with cluster 2.

**Figure 6.**
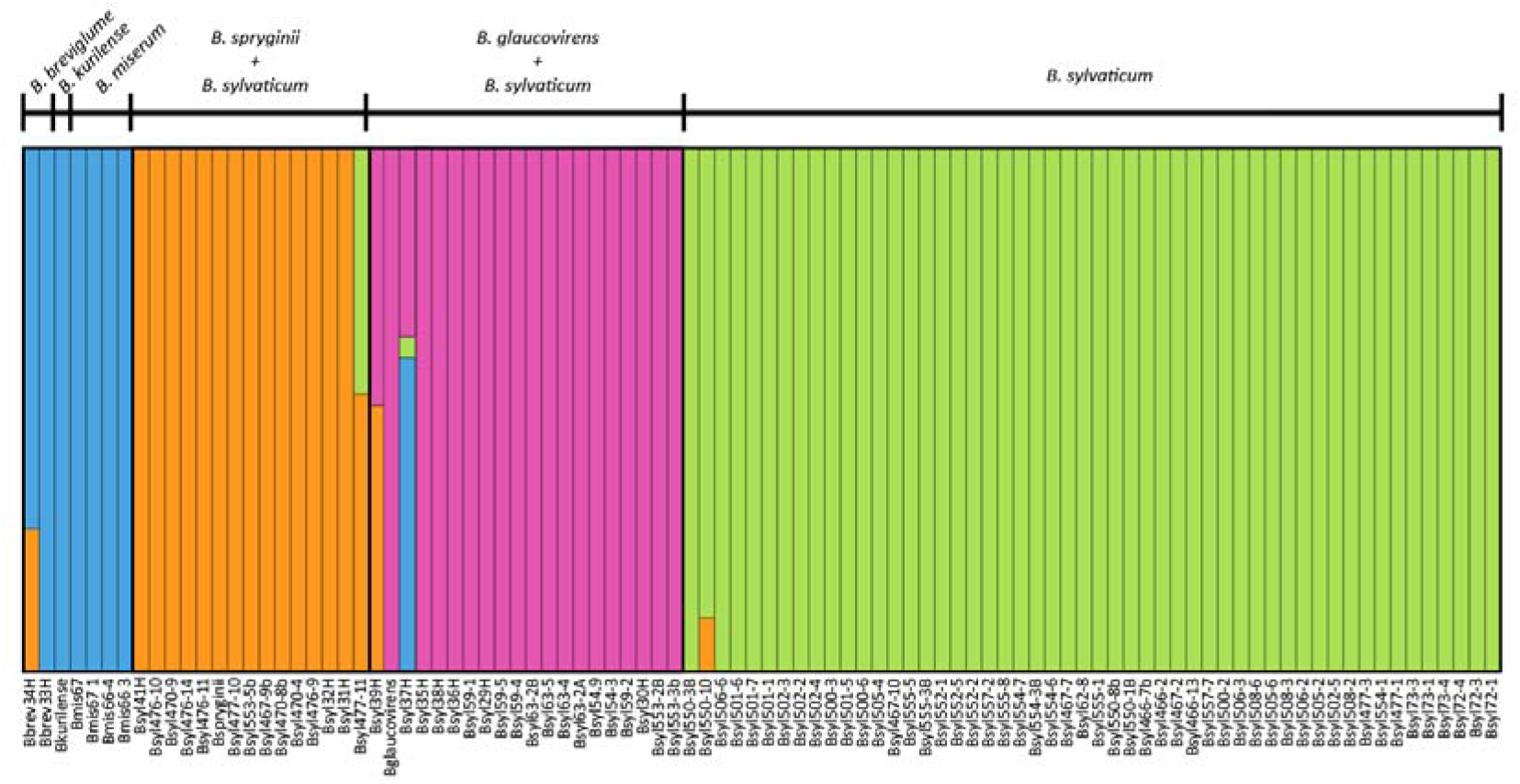
Genetic structure of the studied *Brachypodium sylvaticum* complex samples based on SNPs matrix, showing an optimal number of four population clusters (K = 4). Cluster 1 (blue) encompassed samples of the Eastern Palearctic clade. Cluster 2 (orange) included individuals of *B. spryginii* and *B. sylvaticum* from the Pyrenees (Western Palearctic Subclade A). Cluster 3 (magenta) grouped *B. glaucovirens* and *B. sylvaticum* from Morocco, southern Spain, the Middle East, and the Eastern Mediterranean (Western Palearctic Subclade B). Cluster 4 (green) comprised exclusively *B. sylvaticum* individuals from the Pyrenees, Alps, southern Spain, and the British Islands (Western Palearctic Subclade B).

All taxa within the *Brachypodium sylvaticum* complex studied exhibited low overall levels of genomic diversity, consistent with the predominantly selfing nature of the group (Catalán et al., 2016; Khan and Stace, 1999). Observed heterozygosity values ranged from 0.067 in B. miserum to 0.114 in *B. glaucovirens*, with correspondingly high inbreeding coefficients and selfing rates (Fis = 0.113–0.553; s = 0.203–0.712) (Table 1A). All taxa except *B. glaucovirens* showed low heterozygosity and high Fis values, consistent with this autogamous reproductive system. The comparatively higher heterozygosity and lower inbreeding in *B. glaucovirens* could reflect its putative hybrid origin (Catalán et al., 2023), although the small sample size limits a solid inference. In the ADMIXTURE group (Table 1B), similar trends were observed. The Eastern Palearctic group showed the lowest observed heterozygosity (Ho = 0.071) and one of the highest Fis values (0.534), translating to a selfing rate of 0.696. The groups combining B. sylvaticum with either B. glaucovirens (Cluster 3) or B. spryginii (Cluster 2) exhibited intermediate diversity and inbreeding values (Fis = 0.428–0.438; s = 0.599–0.609), whereas the group composed exclusively of *B. sylvaticum* samples (Cluster 4) showed the highest heterozygosity (Ho = 0.090) and the lowest selfing estimate (s = 0.492), although still within the range expected for a selfing species. Overall, several Fis values deviated significantly from Hardy–Weinberg expectations (Table 1), further supporting the strong effect of selfing in shaping genetic diversity patterns in this complex.

**Table 1.**
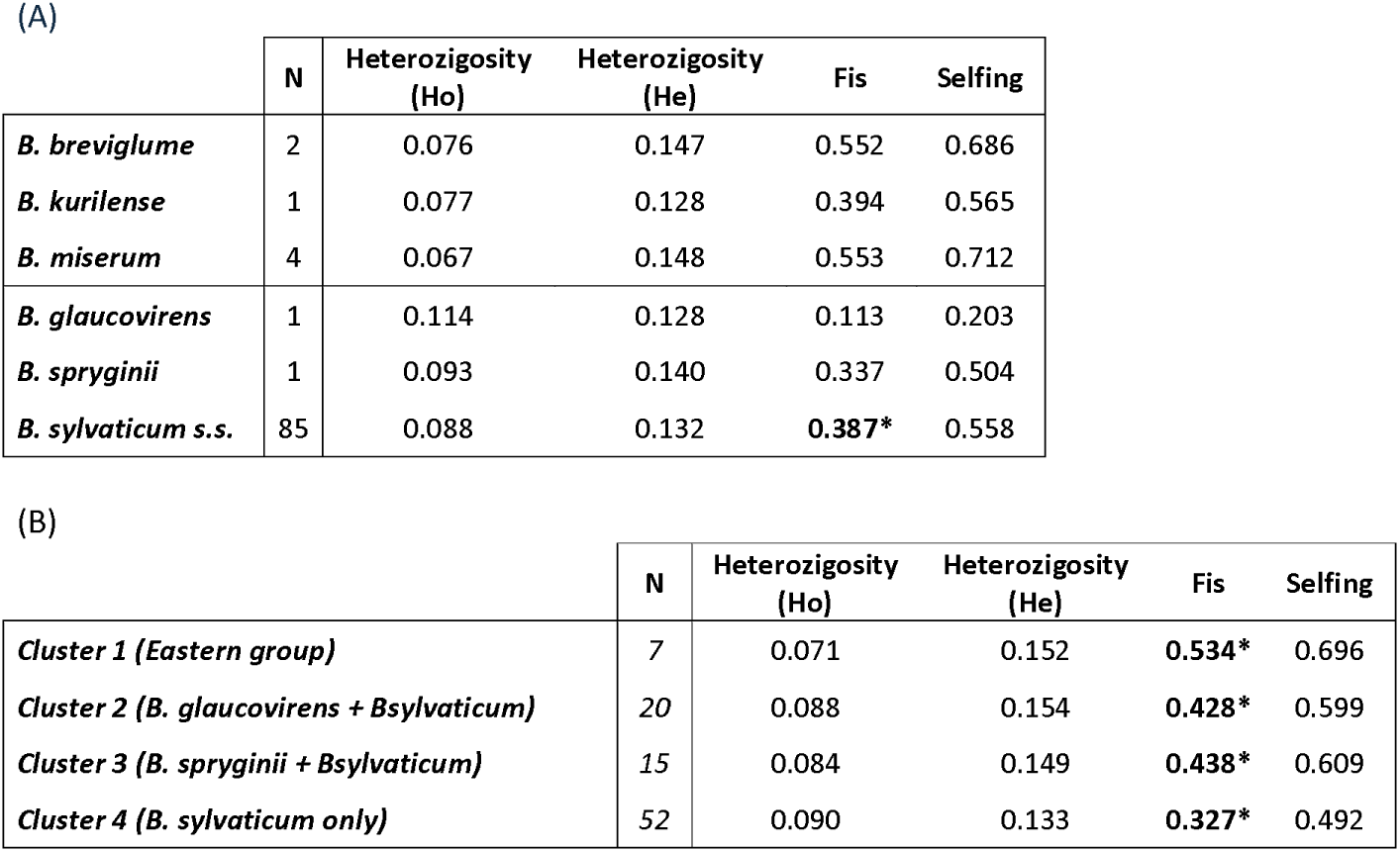
Genetic diversity parameters of the *Brachypodium sylvaticum* complex taxa (A) and the four population genomic clusters obtained by ADMIXTURE analysis (B) (see text and Figure 6). Heterozygosity, inbreeding coefficient (Fis), and selfing rate (s) values were estimated using nuclear genomic data from a 160,124-SNP data matrix. Bold values with an asterisk indicate significant Fis values.

To disentangle the relative contributions of geographic distance (IBD) and environmental factors (IBE) to the genomic structure of *B. sylvaticum* complex populations, separate analyses were conducted for the Eastern Palearctic and Western Palearctic groups (Table 2). In the Eastern group (Table 2A), neither geographic nor environmental distances explained a significant portion of the genetic variation, whether assessed independently or conditionally. These results suggest that the genomic structure within this clade may reflect other factors, such as demographic history, genetic drift, or recent divergence. In contrast, in the Western group (Table 2B), although overall explained variances were low, the effects of IBD and IBE remained significant (p < 0.05), each accounting for approximately 13% of the genetic variation independently. After controlling for geographic distance, environmental factors still explained 10.79% of the variation, indicating that niche-related differentiation may also contribute to genomic structure in this region.

**Table 2.**
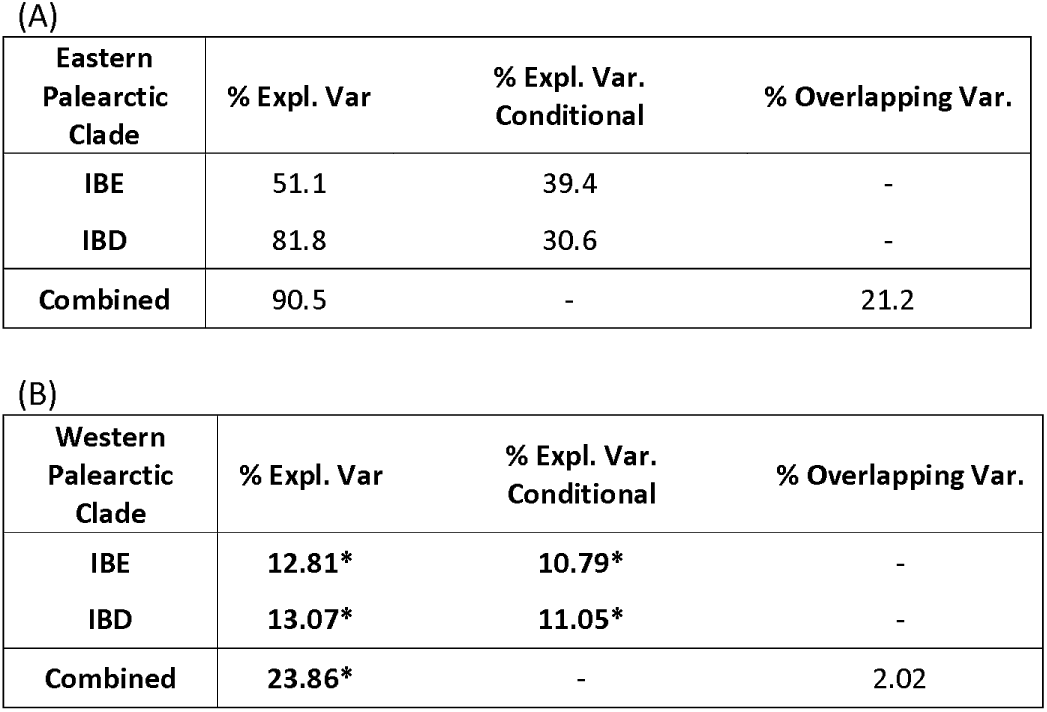
Summary of results from the dbRDA analysis assessing the effects of geographic (IBD) and environmental (IBE) distances on genomic differentiation within samples of the *Brachypodium sylvaticum* complex. Genetic, geographic, and environmental distance matrices were subjected to Principal Coordinate Analysis (PCoA), and dbRDA models were fitted using the resulting axes as predictors. Separate analyses were performed for the Eastern Palearctic clade (A) and the Western Palearctic clade (B). p-values were obtained from permutation tests (n = 10,000). Marginal variance represents the percentage of genetic variation explained by each predictor when tested independently. Conditional variance corresponds to the variation explained by a given predictor after accounting for the effect of the other predictor (partial dbRDA). Shared variance represents the proportion of variance jointly explained by geography and environment in the full model. Values in bold with an asterisk indicate significant p-values.

### 3.4 Bioclimatic analysis and environmental niche shifts of the Brachypodium sylvaticum complex taxa and its Eastern and Western Palearctic groups from the LGM to the present

Taxa and populations of the *B. sylvaticum* complex inhabit regions characterized by relatively mild thermal conditions and marked seasonal variability (Table 3, Appendix 2). Across the entire distribution of the *B. sylvaticum* group (Figure 1), the mean annual temperature averaged 9.43 °C (±4.58), with a marked temperature seasonality of 686.46 °C (±217.06). Thermal extremes ranged from a maximum temperature of 24.03 °C (±5.74) during the warmest month to a minimum of −3.39 °C (±6.48) in the coldest month. The annual temperature range was 27.42 °C (±7.39), with a mean diurnal variation of 8.88 °C (±2.50). In terms of humidity, the *B. sylvaticum* complex is associated with an annual precipitation of 851.56mm (±415.91), with substantial contrasts between the wettest (116.98±66.41mm) and driest (34.74±24.60mm) months.

**Table 3.**
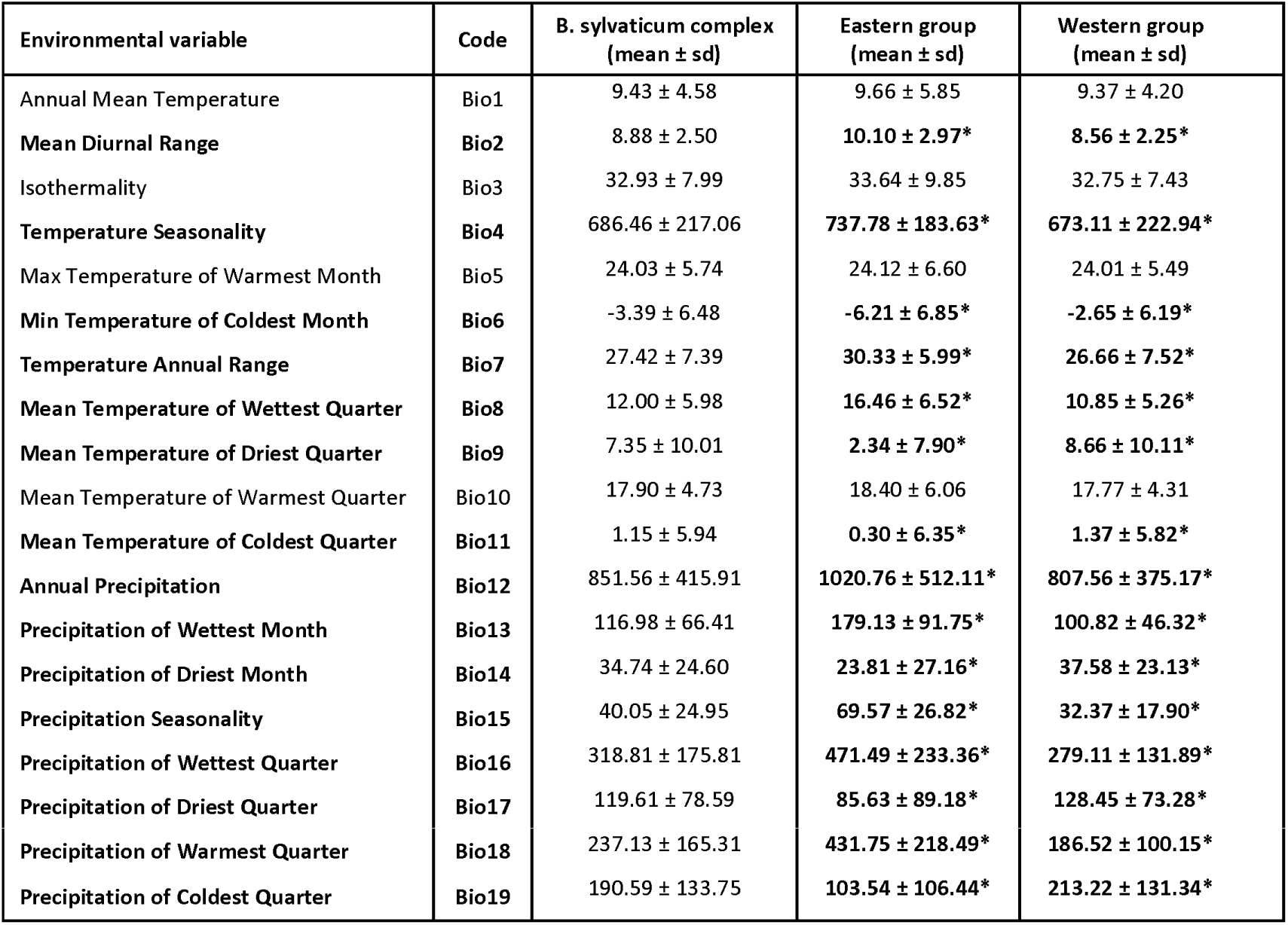
Climatic characteristics of the *Brachypodium sylvaticum* complex and its Eastern and Western Palearctic groups. Mean and standard deviation values for 19 bioclimatic variables. Values highlighted in bold represent bioclimatic variables that showed significant differences (p < 0.05).

When the environmental data were divided into the Eastern Palearctic (including *B. breviglume*, *B. miserum, B. kurilense*) and Western Palearctic (including *B. sylvaticum*, *B. glaucovirens*, *B. spryginii*) groups, significant differences were detected for 16 of the 19 bioclimatic variables (Kruskall-Wallis, p < 0.05; Table 3). While both groups showed comparable mean annual temperatures (Bio1), isothermality (Bio3), and maximum temperatures in the warmest month (Bio5), the Eastern group was characterized by significantly more extreme climatic conditions. This included a greater diurnal temperature range (Bio2), greater temperature seasonality (Bio4), lower minimum temperatures during the coldest month (Bio6), and wider annual temperature ranges (Bio7). Furthermore, the Eastern group experienced significantly higher annual precipitation (Bio12) and greater seasonality of precipitation (Bio15), as well as marked differences in precipitation between the wettest and driest months (Bio13-14) and quarters (Bio16-19).

Principal component analysis (PCA) revealed that the first three axes explained 30.7%, 28.44%, and 17.66% of the total variance, respectively. In the 3D PCA plots (Figures S7A and S7B), *B. sylvaticum* populations largely overlapped with those of the other microtaxa, indicating that overall niche differentiation within the *B. sylvaticum* complex is relatively subtle, although distinct clusters were observed for *B. miserum*, *B. glaucovirens*, and *B. kurilense*. Ecological niche modeling for the *B. sylvaticum* complex yielded highly accurate predictions, indicating excellent model performance (AUC = 0.941, SD = 0.001; Figure 7A). The predicted current species distribution represented a fragmented, ocean-driven climatic niche, with greatest suitability along the northeast Atlantic margins as well as along northwest Pacific coasts, including Japan, the Kuril Islands, and an interior fringe of the Himalayas.

**Figure 7.**
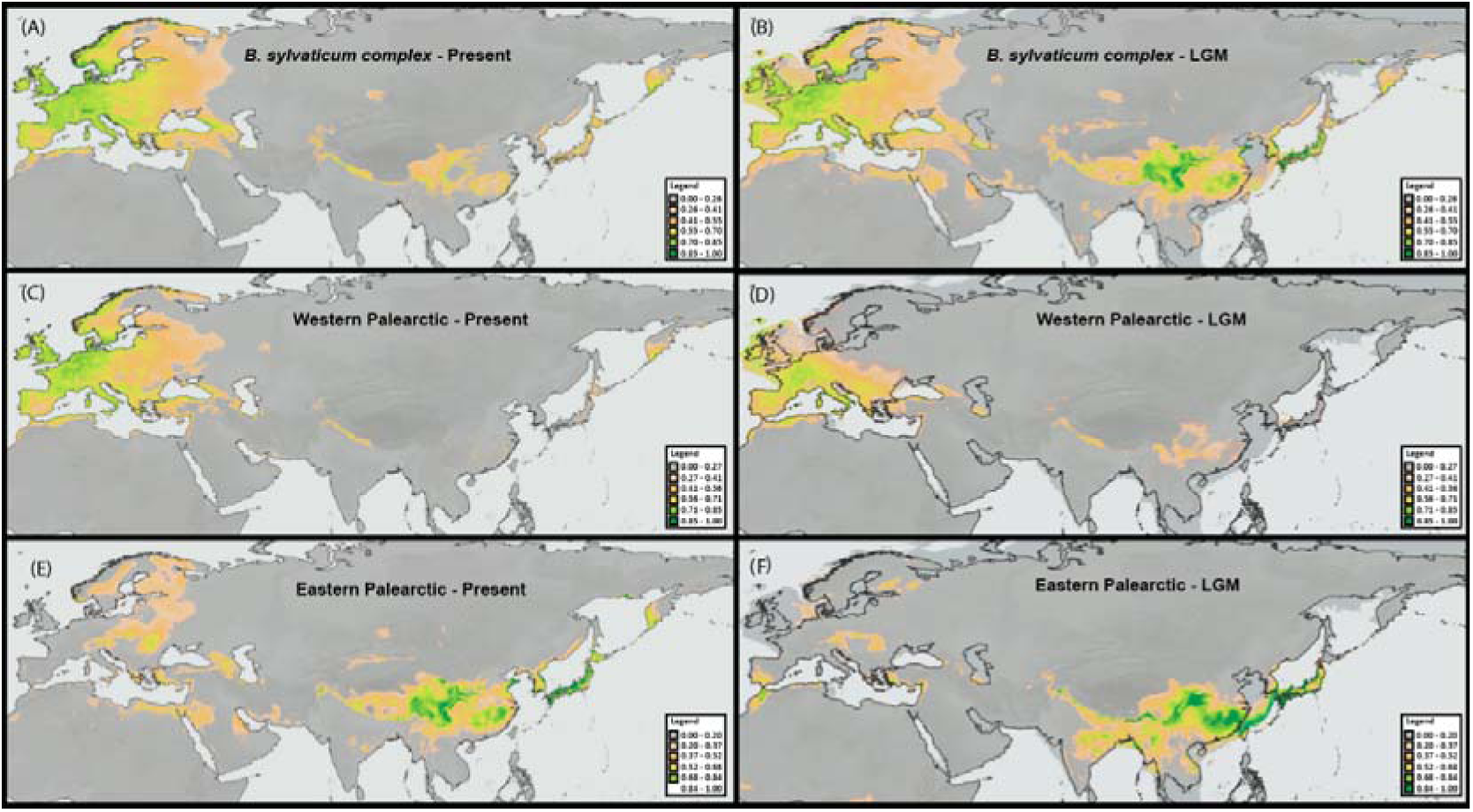
Environmental niche modeling (ENM) for the *Brachypodium sylvaticum* complex and its two main groups under current and Last Glacial Maximum (LGM, ∼20 kya) climatic conditions. Current (A) and LGM (B) ENM projections for the *B. sylvaticum* complex. (B). Current (C) and LGM (D) ENM projections for the Western Palearctic group. Current (E) and LGM (F) projections for the Eastern Palearctic group.

Paleoclimatic projections for the Last Glacial Maximum (LGM) indicate the persistence of glacial refugia in coastal, island, and lowland regions of western Europe (Atlantic margin), eastern Asia (Pacific margin), and in mountain ranges of western and southern Asia (Figure 7B). These results support a longitudinal shift in distribution, with species likely shifting from glacial refugia in eastern and western coastal areas to more continental interiors as postglacial climates became progressively milder and wetter (Figure 7). Separate models for the Eastern and Western Palearctic groups of the *B. sylvaticum* complex further highlighted their distinct ecological preferences and possible historical distributions (Figures 7C–7F). The Western Palearctic group displayed a primarily European distribution, with high suitability extending eastward to the Caspian region and scattered areas of moderate suitability in central Siberia, the Himalayas, and the Kuril Islands (Figure 7C). In contrast, the Eastern Palearctic group exhibited a niche centered on China and the Himalayan region, with high probabilities of occurrence extending to Japan and the Pacific coast of East Asia, including the Kuril Islands, and lower suitability inferred for parts of Eastern Europe (Figure 7E). LGM reconstructions suggest that both groups persisted in distinct glacial refugia: the western group along the Atlantic coast of Europe (Figure 7D) and the eastern group in the Himalayas and East Asia (Figure 7F). The present-day distributions of both groups (Figures 7C-7E) reflected partial postglacial expansions from these refugia.

Niche overlap between the Western and Eastern Palearctic groups was moderate according to Schoener’s D (0.473) and Hellinger’s I (0.702), with a non-significant result in the equivalence test (p = 1), indicating that the two niches were statistically indistinguishable when group identity was randomized. In contrast, the similarity test yielded a significant result (p = 0.027), suggesting that the observed overlap was greater than expected by chance given the environmental context (Table 4A). According to Warren et al. (2021), this pattern supports niche conservatism, implying that despite their deep genomic divergence and geographic separation, the two clades retain broadly similar climatic preferences. Standardized niche breadth values (Levins’ B2) remained similarly low across time periods within each clade, with the western group showing a slight increase from 0.109 under present-day conditions to 0.117 during the LGM, and the eastern group increasing from 0.107 to 0.111 (Table 4B). Temporal niche overlap between present-day and LGM projections, as measured by Schoener’s D, was moderate in both cases, with values of 0.625 for the western group and 0.487 for the eastern group, indicating partial shifts in climatic suitability over time.

**Table 4.**
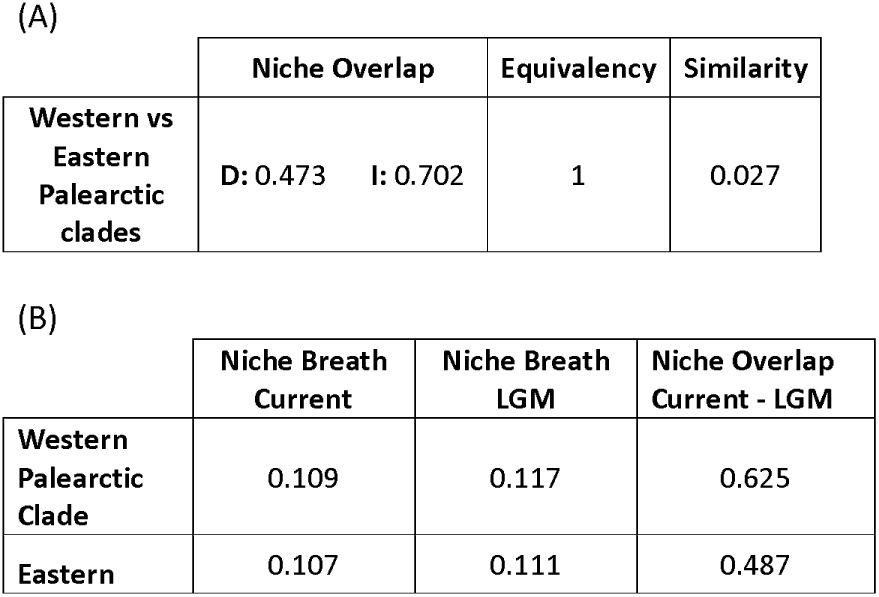
Summary of niche overlap and identity tests between the Western and Eastern Palearctic groups of *Brachypodium sylvaticum* (A) and between Last Glacial Maximum (LGM) and present-day climate data (B). Niche overlap was quantified using Schoener’s D and Hellinger’s I indices. For the equivalence test, p-values less than 0.05 indicate distinct niches, whereas for the similarity test, p-values less than 0.05 represent similar niches; all p-values were based on 1000 permutations. Subtable B presents niche breadth and temporal niche overlap for each group. Niche breadth was calculated for the current and Last Glacial Maximum (LGM) periods using Levins’ B2 indices (standardized). The last column shows the Schoener D value representing the niche overlap between present-day and LGM projections within each clade.

## 4. Discussion

### 4.1 Evolutionary diversification of the main Brachypodium sylvaticum complex lineages and microtaxa

Our phylogenomic analyses, integrating genome-wide SNP and plastome data from 94 accessions, resolved the *B. sylvaticum* complex into two deeply divergent sister lineages, the Eastern Palearctic clade (Himalayas to the Pacific) and the Western Palearctic clade (Europe, North Africa, and Siberia). These clades diverged approximately 1.75 Ma ago (95% HPD: 1.2– 2.1 Ma; Figure 2), coinciding with Middle Pleistocene climate shifts that fragmented widely distributed ancestral angiosperm populations into refugia (Díaz-Pérez et al., 2018; Hewitt, 2004). This split partially corroborates previous findings (Catalán et al., 2023), but with unprecedented resolution, revealing reciprocal monophyly and distinct evolutionary trajectories for the Eastern and Western Palearctic lineages (Figures 2, 3, 4, S2). Importantly, however, the plastome-based phylogeny reconstructed the *B. sylvaticum* complex as paraphyletic, with *B. pinnatum* resolved as sister to the Western Palearctic lineage (Figs. 4, S2), highlighting the proximity of the core perennial lineages *B. sylvaticum* and *B. pinnatum* and their high intercrossability (Khan & Stace 1999), resulting in their shared matrilineal plastomes (Catalan et al. 2016).

Our broader sampling has also refined the geographic and taxonomic boundaries of the studied taxa within each lineage. The East Palearctic clade, comprising *B. breviglume* (Himalayas), *B. miserum* (Japan), and *B. kurilense* (Kuril Islands), exhibits a robust geographic structure with no evidence of hybridization (Figures 2, 4, 6). The divergence dates (∼1.35–1.06 Ma; Figure 2) predate the Last Glacial Maximum and could correspond to earlier Pleistocene climatic and geological events. The emergence of the Great Kuril Range during the Early Pleistocene (Bulgakov, 1996) and subsequent glacial cycles could have promoted geographic isolation and speciation. Biogeographic reconstruction (DIVA+J model; Figure 3A) supports an ancestral distribution in the Himalayas for *B. breviglume*, followed by eastward dispersal and allopatric speciation. The paraphyly of *B. breviglume* and the sister relationship of *B. miserum* and *B. kurilense* (Figures 2, 3) suggest a stepwise colonization from the Himalayas to the Pacific coast, with subsequent isolation likely driven by Pleistocene sea-level changes that fragmented the temperate biota across East Asia (Qiu et al., 2011).

In contrast, the Western Palearctic clade (*B. sylvaticum*, *B. glaucovirens*, *B. spryginii*) revealed phylogeographic patterns driven by Pleistocene climatic oscillations. The SNP nuclear tree separated two well-supported subclades, Subclade A, which includes Pyrenean lineages and Caucasian *B. spryginii*, and Subclade B, a more heterogeneous assemblage including Mediterranean, North African, and British Islands populations. These two subclades showed different patterns of diversification and geographic expansion, with Subclade B exhibiting higher levels of admixture and secondary contact (Figure 6), suggesting contrasting demographic histories within the Western clade. Cytonuclear discordance (RF=0.7286; Figure 4) highlighted repeated chloroplast capture events, particularly in populations of *B. spryginii* (Caucasus) and Mediterranean *B. sylvaticum*, suggesting historical connections between glacial refugia in Anatolia and the Caucasus (Figure S2). The continuity of temperate deciduous forests in these regions during Quaternary climate oscillations likely facilitated gene flow or historical dispersal (Novák et al., 2023). However, phylogeographic patterns in the western A and B subclades reflected the survival of populations in warmer circum-Mediterranean refugia in the Early Pleistocene (>1 Ma) and recent colonization of cooler high-latitude areas in the Late Pleistocene (<1 Ma) (Figure 3), consistent with similar survival and dispersal patterns in other angiosperms (Hewitt, 2004; Nieto Feliner, 2014). Genomic admixture analysis (K=4, Figure 6) further reveals genomic admixture in Pyrenean and Balkan populations, reflecting secondary contact zones in these southern European refugia, where divergent lineages were found during the Holocene warming (Hewitt, 2004). Similarly, the topological incongruence observed between nuclear and plastome trees (Figure 4) reflected the high level of introgression between geographically distant populations of *B. sylvaticum*, facilitated by the recurrence of LDD in the Western Palearctic regions during the Pleistocene (Fig. 3).

In this context, *B. glaucovirens*, a putative hybrid between *B. sylvaticum* and *B. pinnatum* (Catalán et al., 2023; Scholz, 2007), stands out for its high heterozygosity (Ho = 0.114 vs. 0.067– 0.093 in other taxa; Table 1), supporting its hybrid origin. This is further corroborated by experimental and natural hybridization studies reporting intermediate individuals between *B. glaucovirens* and *B. sylvaticum* with 2n = 17 chromosomes and reduced pollen fertility, indicating spontaneous hybrid formation (Khan and Stace, 1999). Despite displaying partial sterility, these hybrids were able to backcross, suggesting the potential for introgression into sympatric populations. Notably, *B. glaucovirens* displays lower self-compatibility than *B. sylvaticum*, implying a tendency towards greater allogamy, which could further promote genetic exchange where their ranges overlap (Khan and Stace, 1999). The predominantly autogamous mating system of *B. sylvaticum* explains its low nuclear diversity (Ho = 0.067– 0.090; Table 1), in contrast to allogamous relatives, such as *B. rupestre*, which displays higher heterozygosity (Ho ∼0.13) and low inbreeding coefficients, even in clonal populations, due to mixed sexual and asexual reproduction strategies (Durán et al., 2025). Despite this, selfing facilitated rapid postglacial colonization events of *B. sylvaticum*, such as LDDs from western Europe to Iceland and Siberia (Figures 6, 7, 8) and invasive success in North America (Rosenthal et al., 2008), mirroring the “colonizer syndrome” of the selfing annuals *B. distachyon* and *B. hybridum* (Catalán et al. 2016), and closely resembling the rapid spread of the highly selfing annual *B. stacei* across the Mediterranean through repeated LDDs and bottleneck events (Campos et al., 2024).

### 4.2 Coevolution and co-speciation of Brachypodium sylvaticum complex and its fungal endophyte Epichloë sylvatica

Our analysis of fungal ITS reads derived from total holobiont DNA successfully recovered *Epichloë sylvatica* sequences from nearly all *B. sylvaticum* complex accessions (>95% infection rate), enabling robust phylogenetic reconstruction despite low fungal coverage in plant-targeted sequencing. This represents one of the first studies to recover high-quality *Epichloë* genetic data directly from genome skimming datasets of the host plant (holobiont). The nearly complete recovery of the ITS region with high sequence quality across geographically diverse samples highlights the utility of this approach for symbiont characterization in non-targeted genomic workflows. Strikingly, *E. sylvatica* was detected in the Eastern Palearctic taxon *B. kurilense*, marking its first confirmed occurrence outside the Western Palearctic clade. It suggests *E. sylvatica* may occur more widely across the *B. sylvaticum* complex, though its absence in other Eastern taxa (e.g., *B. breviglume*, *B. miserum*) could reflect sampling gaps or ecological exclusion. Notably, *E. sylvatica* has not been detected in related Western species like *B. pinnatum* and *B. phoenicoides*, which host distinct endophytes (e.g., *E. typhina;* Leuchtmann & Schardl, 2022), underscoring its unique specificity to the *B. sylvaticum* lineage. The ecological preferences of the *B. sylvaticum* complex taxa to forested humid habitats across the Palearctic (Figure 7, Table 3) probably fostered the infection of their presumably humid-adapted crown grass ancestor by the endophyte’s ancestor since the early Pleistocene (Figures 3, 4, S3, S4) resulting in the co-speciation of both organisms after 1.7 Ma of shared symbiosis. The phylogeny revealed two well-supported *E. sylvatica* clades, I and II (Figure S3), demonstrating genetic structuring within this ubiquitous endophyte. Interestingly, the co-evolutionary analyses detected significant global concordance from both Parafit and PACo approaches (Figures 5, S4) while PACo further revealed a relatively high number of host-endophyte individual interactions (Figure 4), thus supporting a general common evolutionary framework of *B. sylvaticum* population and their respective endophytes. This pattern suggests a long-term association with the co-evolution reinforced by vertical transmission and the host’s selfing mating system.

While ITS data could not distinguish mating (sexual) and non-mating (asexual) strains of *E. sylvatica* (Leuchtmann & Schardl, 2022), we observed geographic structuring within fungal clades. For instance, identical *E. sylvatica* haplotypes occurred in distant populations (e.g., Pyrenees *vs*. Caucasus), aligning with prior reports of genetic homogeneity in this endophyte. Phylogenetic congruence analyses revealed complex dynamics; thus, PACo detected significant global host–fungus concordance (p < 0.001; Figure 4), reflecting fidelity imposed by vertical transmission in the highly selfing host (s = 0.492–0.712; Table 1). In contrast, ParaFit identified only localized associations (12% significant links; Figure S4). While both PACo and ParaFit are designed to assess global phylogenetic congruence in host–symbiont systems, PACo has been shown to exhibit greater statistical power and robustness to noise, particularly under scenarios of partial congruence or moderate topological uncertainty (Balbuena et al., 2013; Hutchinson et al., 2017). This algorithmic background likely explains the broader host–endophyte signal recovered by PACo, without necessarily implying differences in their sensitivity to horizontal transmission. The near ubiquitous presence of *E. sylvatica* in *B. sylvaticum* holobionts suggests mutualistic benefits, potentially enhancing host ecological success.

### 4.3 Past climate niche conservatism, glacial refugia, and postglacial spread

Our integrated phylogenomic and environmental niche modeling (ENM) analyses revealed distinct Quaternary responses of *Brachypodium sylvaticum* Eastern and Western Palearctic clades, which diverged ∼1.74 Ma (Figure 2). While both clades share adaptations to humid, oceanic habitats, their evolutionary trajectories and niche dynamics reflect independent histories shaped by contrasting glacial-interglacial pressures. The Western Palearctic lineage likely persisted in western Europe and Mediterranean LGM refugia (Pyrennees + Alps, Morocco + South Spain, Middle East, SW Europe) ), (Figures 3, 7D), in agreement with general refugial patterns identified for temperate taxa across the Mediterranean and Eurasia (Médail & Diadema, 2009; Svenning et al., 2008; Taberlet et al., 1998). Post-LGM expansion of the Western clade into northern Europe and western Asia—particularly into Scandinavia, the British Islands, and the Caucasus—is supported by phylogenomic and genetic patterns of genomic admixture, cytonuclear discordance, and shallow node divergence times (Figures 2, 4, 6), and by the biogeographic models (Figure 3), which may oversimplify the complexity of recent postglacial migration. This expansion likely reflects a latitudinal migration from southern refugia to the North under increasingly oceanic postglacial climates, a trend observed in other mesic-adapted taxa (Leipold et al., 2017). In contrast, the Eastern Palearctic clade (Himalayas to Asian Pacific coast) exhibited spatial niche persistence since the LGM (Figures 7E-7F), maintaining its distribution within climatically stable mountainous and coastal regions. This pattern reflects long-term range continuity, rather than climatic niche stability, and is consistent with its long-term isolation, low genetic diversity (Table 1), and ancestral origin in East Asia –Himalayas inferred from dating phylogenies (Figure 2; Catalán et al., 2023) and ancestral area reconstruction (Figure 3). Oceanic islands (e.g., Japan, Kurils) and the Himalayan corridor likely acted as climatically stable refugia, buffering the Eastern Palearctic lineage against Quaternary climatic upheavals. Similar long-term persistence has been reported in other parts of temperate Asia, including periglacial lowlands, as supported by ancient DNA analyses and plant regeneration studies from ancient permafrost (Thomsen & Willerslev, 2015; Yashina et al., 2012).

Niche overlap between clades was moderate (Schoener’s D = 0.473; Hellinger’s I = 0.702), with statistically significant similarity (p-value = 0.027) and equivalent niches (p-value = 1.00), indicating that the niches occupied by the Eastern and Western Palearctic lineages are more similar than expected by chance. This pattern is best interpreted as evidence of niche conservatism, suggesting that divergence between clades was likely due to geographic and historical isolation, rather than adaptive ecological speciation. Despite profound genomic divergence and long-term separation, both clades converged on similar mesic climatic niches, likely constrained by ancestral ecological preferences.

Temporal comparisons of niches between the LGM and the present revealed asymmetric niche shifts, the Western Palearctic clade maintained moderate stability (present-day LGM Schoener’s D = 0.625), whereas the Eastern Palearctic clade experienced greater climatic niche turnover (present-day LGM Schoener’s D = 0.487), likely due to increased climatic fragmentation and heterogeneity in high-altitude refugia in East Asia. This is consistent with phylogeographic evidence from the region showing that Quaternary climatic oscillations promoted population isolation and profound lineage divergence in the Himalayas, eastern China, and Japan due to mountain barriers and habitat fragmentation (Qiu et al., 2011). Both lineages showed a slight expansion of niche breadth (Levins B2: western clade 0.109 1 0.117; eastern clade 0.107 1 0.111), suggesting gradual postglacial diversification rather than abrupt ecological changes.

## Data availability

All the data matrices assembled and related phylogenomic and biogeographic results in this study have been deposited to the public database Github (https://github.com/Bioflora/BsylvaticumPhylogeography). The raw sequencing data have been submitted to the European Nucleotide Archive (ENA) under the project numbers PRJEB64465 and PRJEB89851.

## Aknowledgements

We thank L.A. Inda Aramendía and J. Ascaso Martorell for their assistance with field sampling and collection of *Brachypodium sylvaticum* specimens, M.F. Moreno-Aguilar for helpful support with the implementation of custom scripts, and the herbaria of B (Berlin), M (München), LD (Lund), VLA (Vladivostok), RIKEN (Japan) for providing access to voucher material.

## Author contributions

PC, MD, MC and EP designed the study. PC and EP collected samples. MD, MC, EP, DC and PC conducted and supervised the analyses. PC, MD, MC, DC and EP wrote the original draft, and all authors reviewed it.

---

**Figure S1** Comparison of plastome maps constructed from whole-plastome consensus sequences of *B. sylvaticum s. s.* (outermost first), *B. miserum* (second), *B. kurilense* (third) and *B. breviglume* (inner). The small inner black circle represents the four major plastome regions (Long Single Copy (LSC), Small Single Copy (SSC), Inverted Repeat A (IRA), and Inverted Repeat B (IRB). Maps highlight a ∼1K bp insertion in the LSC region containing a rpl23 pseudogene and a ∼300 bp insertion containing a copy of rps19 in the IRB, which are present in all four *B. sylvaticum* complex taxa plastomes.

**Figure S2** Maximum likelihood phylogenomic tree of the *Brachypodium sylvaticum* complex samples based on whole-plastome sequences; values on branches show bootstrap support (BS). The blue clade represents the Eastern Palearctic group, while the green clade corresponds to the Western Palearctic group.

**Figure S3** Maximum likelihood phylogenetic tree of *Epichloë sylvatica* based on the complete sequence of the rDNA ITS region (ITS1, 5.8S, ITS2); values on branches correspond to bootstrap support (BS), branches with BS values below 20 are collapsed into polytomies. Red branches correspond accessions of the *Epichloë* spp. reference genomes used, and brown branches to the samples analyzed in this study.

**Figure S4** Cophylogenetic analysis of the *Brachypodium sylvaticum* complex nuclear ML tree and the *Epichloë sylvatica* ITS ML tree based on host-endophyte topological conflicts ParaFit results. Green lines indicate congruent branches, whereas lilac lines mark incongruent associations. Colored circles represent *B. sylvaticum* complex Eastern Palearctic Clade (blue), Western Palearctic Subclade A (orange), and Western Palearctic Subclade B (yellow) (see Figure 2).

**Figure S5** Bar plot illustrating normalized squared residuals from PACo analyses between phylogenetic trees of *Brachypodium sylvaticum* (host, nuclear SNPs data) and *Epichloë sylvatica* endophyte, ITS data). Each bar represents a specific host-endophyte association, showing the median of normalized squared residuals with 95% confidence intervals (blue lines). The black horizontal line indicates the incongruence threshold calculated with the Q3+Q4 quartile; bars exceeding this threshold represent incongruent associations (red), while bars below denote congruent associations (blue).

**Figure S6** Graphical representation of Bayesian genetic structure Puechmaille’s estimators for determining the optimal number of genetic groups (K) within *Brachypodium sylvaticum* complex samples. The graphs display the values of MedMed, MedMean, MaxMed, and MaxMean estimators calculated over 10 replicates for each K (ranging from 1 to 10) based on the admixture analysis of the studied samples. The red lines indicate the optimal number of genetic clusters (K) estimated from each parameter.

**Figure S7** Climatic principal component analysis (PCA) of the *Brachypodium sylvaticum* complex samples, retrieved from their respective Palearctic geographical areas (see Figure 1), based on 19 worldclim bioclimatic variables. The plots show different views of the projections (A, B). Different colors and shapes represent the *B. sylvaticum* complex taxa (see Figure 1).

**Table S1** List of the 94 *Brachypodium sylvaticum* complex samples (85 *B. sylvaticum s. s., 2 B. breviglume, 1 B. glaucovirens, 1 B. kurilense, 4 B. miserum, 1 B. spryginii*) included in the phylogeographic study of the complex. Information on taxon name, population code, sampling size (N), geographic location, georeferenced coordinates, and ENA accession code is provided in the table, and that on samples from seed germplasms [RIKEN, Japanese Center for Sustainable Resource Science] or herbarium specimens [B (Berlin), M (München), LD (Lund), VLA (Vladivostok)] under Location. The remaining samples were collected in the field. All studied *B. sylvaticum* complex samples are diploids. Taxonomic identities are based in the taxonomic works of Keng (1982), Probatova (1985), Schippmann (1991), Catalán et al. (2016), Tzvelev (2015), Tzvelev & Probatova (2019). Samples corresponding to the outgroups used for this study (*B. stacei, B. distachyon, B. arbuscula*, *B. pinnatum*) are listed at the bottom of the table. Newly studied accessions are indicated in bold.

**Table S2** Summary statistics of the genomic data for the *Brachypodium sylvaticum* complex samples under study and their *Epichloë* endophytes. Information is provided on taxon name, population code, total number of Illumina raw reads generated from genome skimming analysis (RawReads), number of *Brachypodium* reads that successfully mapped to the *B. sylvaticum* reference genome Ain1 (BsylReads), number of *B. sylvaticum* protein-coding genes with at least 90% of sequence positions covered at a minimum depth of 10 (Genes90), assembled plastome length (Plastome), number of *Epichloë* reads mapped against *Epichloë* reference genomes (EpichloëReads), percentage of fungal endophyte rDNA 35S cistron recovered (Cistron%), and BLAST result (Blast) indicating whether the recovered fungal cistron belongs to *Epichloë sylvatica* (Yes) or to other *Epichloë* endophyte (No).

**Table S3** Results of co-evolutionary ParaFit analysis for individual *Brachypodium sylvaticum* complex *– E. sylvatica* associations detected over 1000 replicates. The F1 statistic quantifies the degree of phylogenetic congruence between each *Brachypodium* host plant and its associated *Epichloë* endophyte. In bold, the p-values with p < 0.05 supporting significant co-evolutionary relationships. Global ParaFit value and p-value are also indicated.

**Table S4** Results of Procrustean Approach to Cophylogenetics (PACo) analysis for individual *Brachypodium sylvaticum* complex *– Epichloë sylvatica* associations. The sum of squared normalized residual values represents the deviation of each host–endophyte link from the expected coevolutionary pattern, with higher residuals indicating greater phylogenetic incongruence. Associations are classified as Congruent (median residual ≤ 0.012) or Incongruent (median residual > 0.012), using 0.012 as the threshold defined by the combined Q3+Q4 quartiles of the bootstrap distributions.

**Table S5**: Genetic diversity parameters at the Individual level for each *Brachypodium sylvaticum* complex sample calculated using plink2. For each accession, observed heterozygosity (Ho), expected heterozygosity (He) and the method-of-moments F coefficient are shown in the table.

**Appendix 1** List of geographic coordinates and values of 19 bioclimatic variables for each of the 1,263 *Brachypodium sylvaticum* complex samples used in this study, extracted from the WorldClim database (2.5 arc-minute resolution). The dataset includes 169 *B. breviglume*, 4 *B. glaucovirens*, 29 *B. kurilense*, 49 *B. miserum*, 2 *B. spryginii*, and 1,009 *B. sylvaticum* samples. Samples shown in bold correspond to individuals for which genomic data were available; the remaining records were retrieved from GBIF based on taxonomic identity and geolocation. *Appendix 2* Expanded Materials and Methods

