## Supplementary figures and images for "A Palearctic divide, niche conservatism and host-fungal endophyte interactions shaped the phylogeography of the grass *Brachypodium sylvaticum*"

### Figure S1

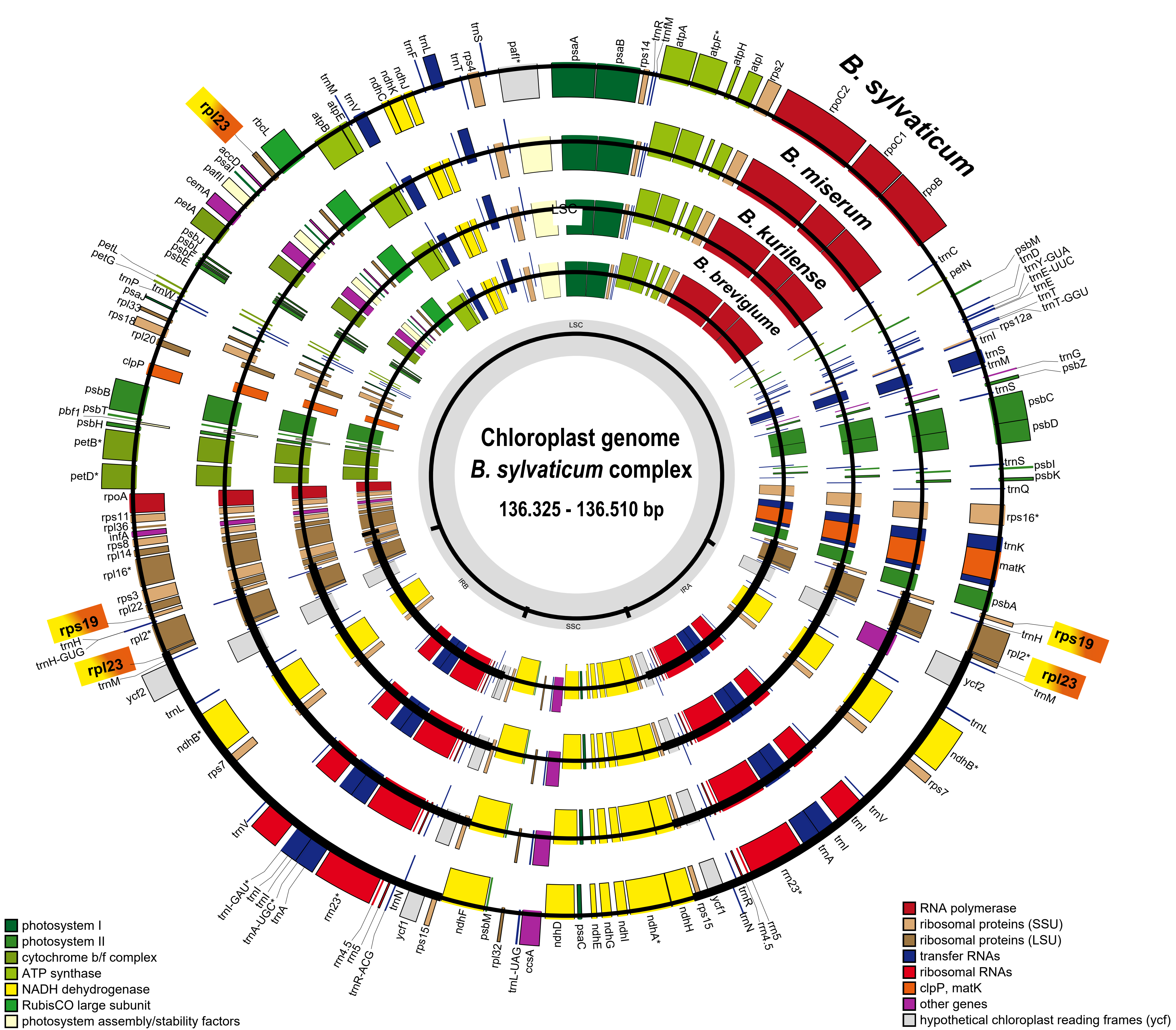

### Figure S2

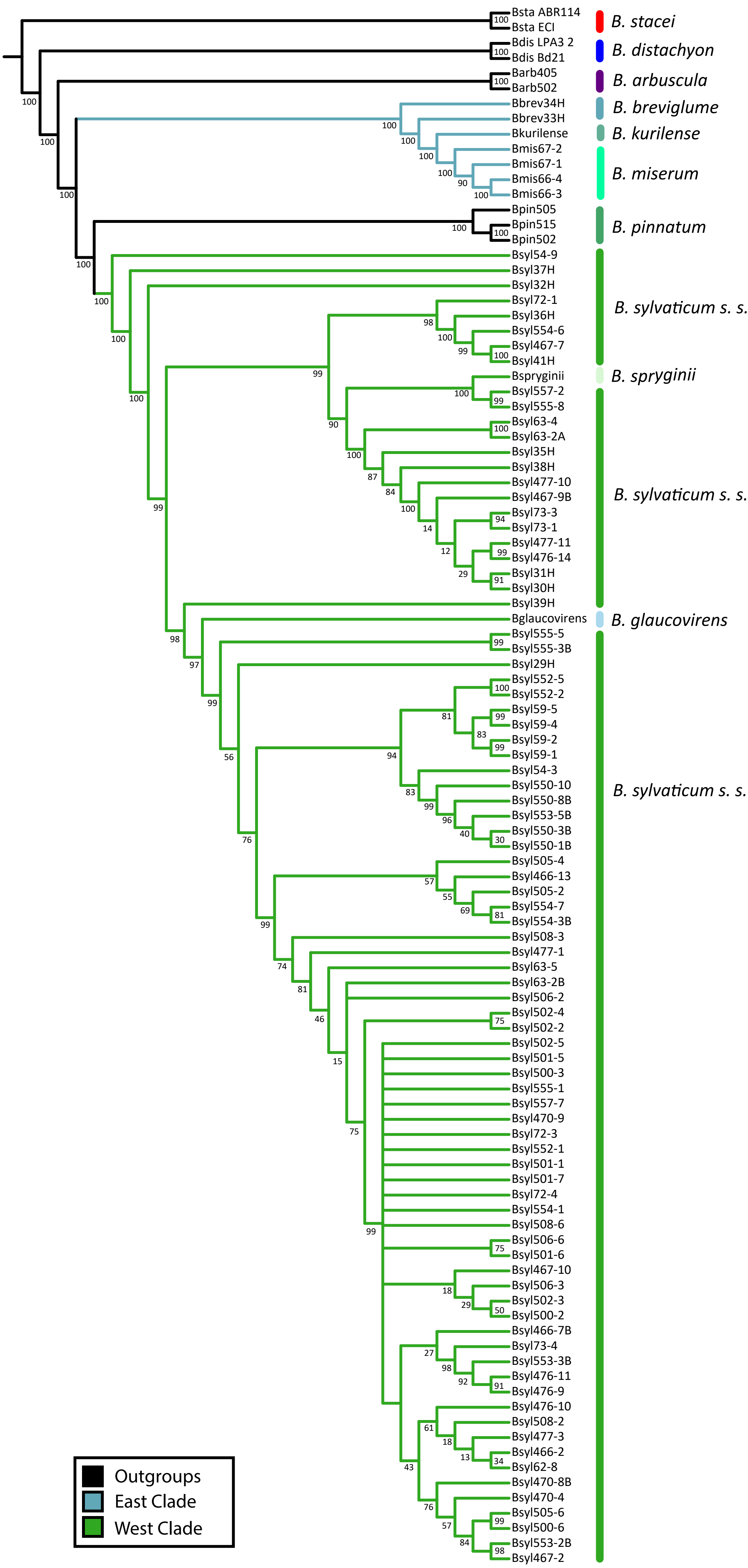

### Figure S3

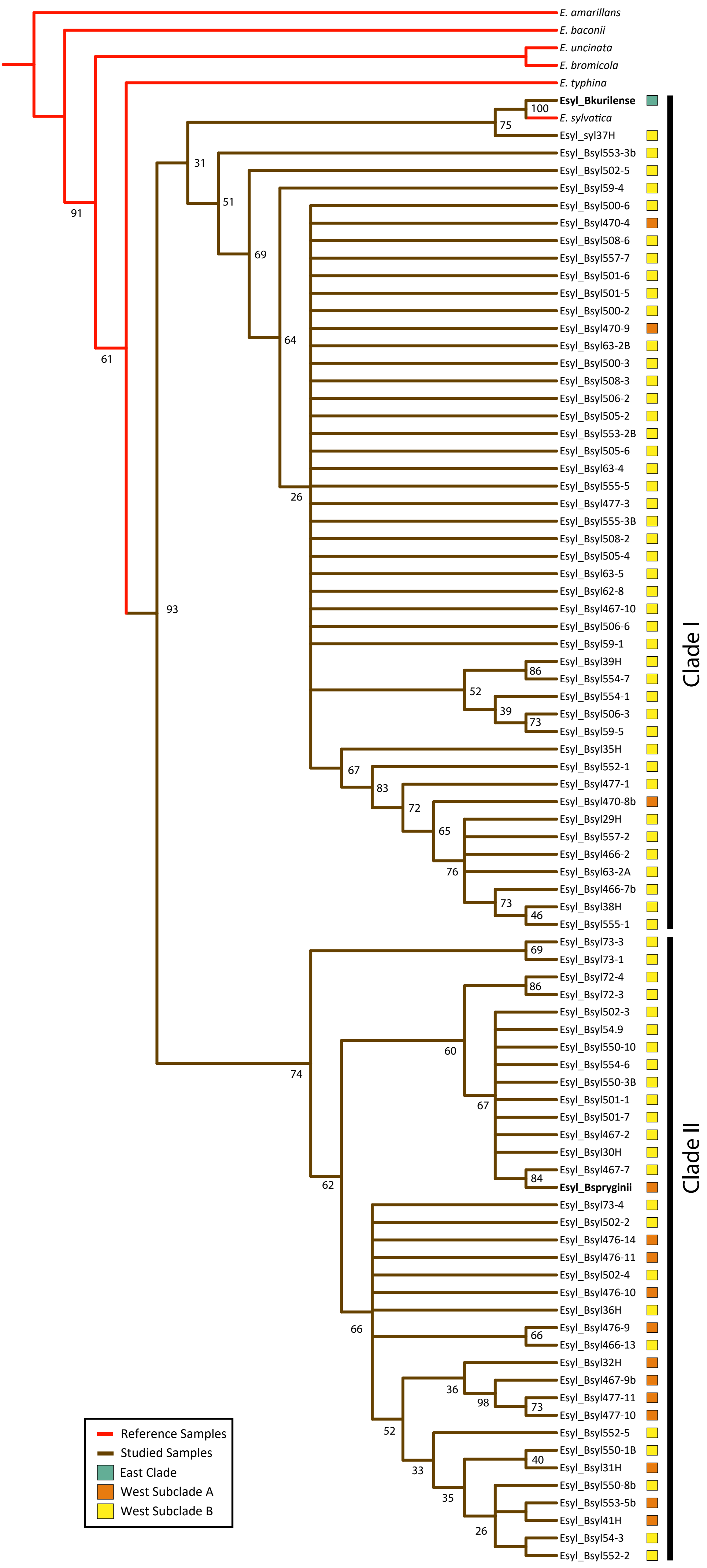

### Figure S4

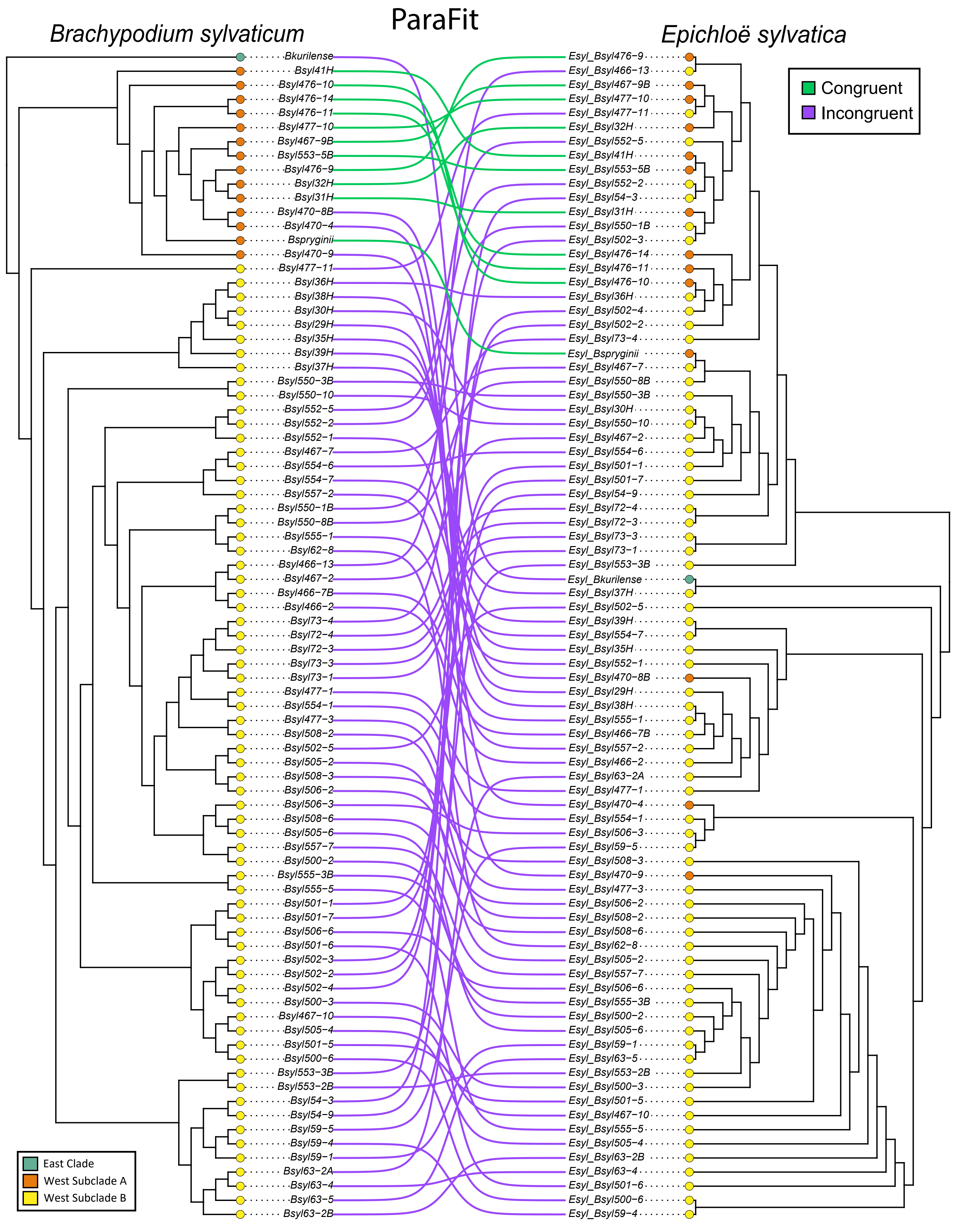

### Figure S5

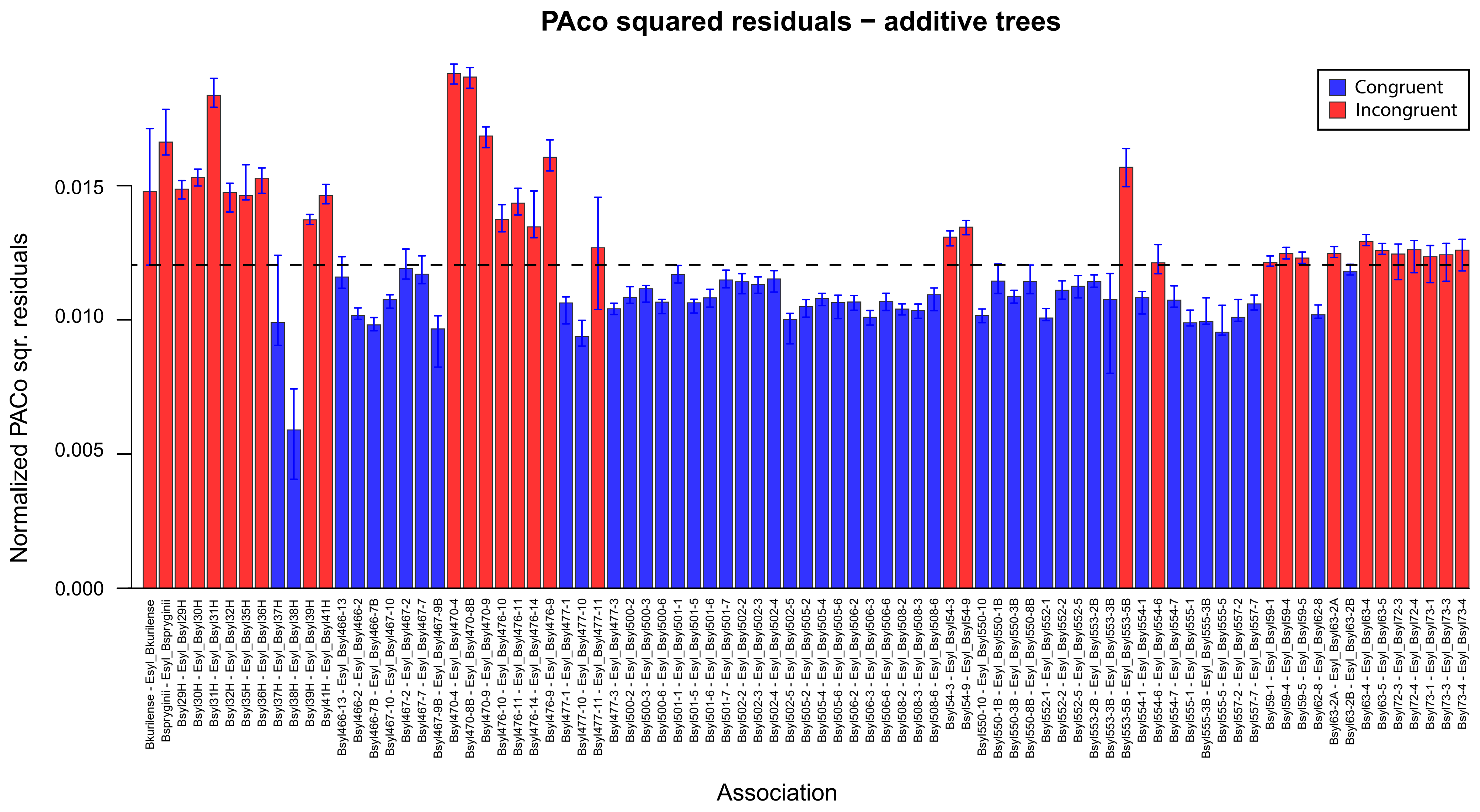

### Figure S6

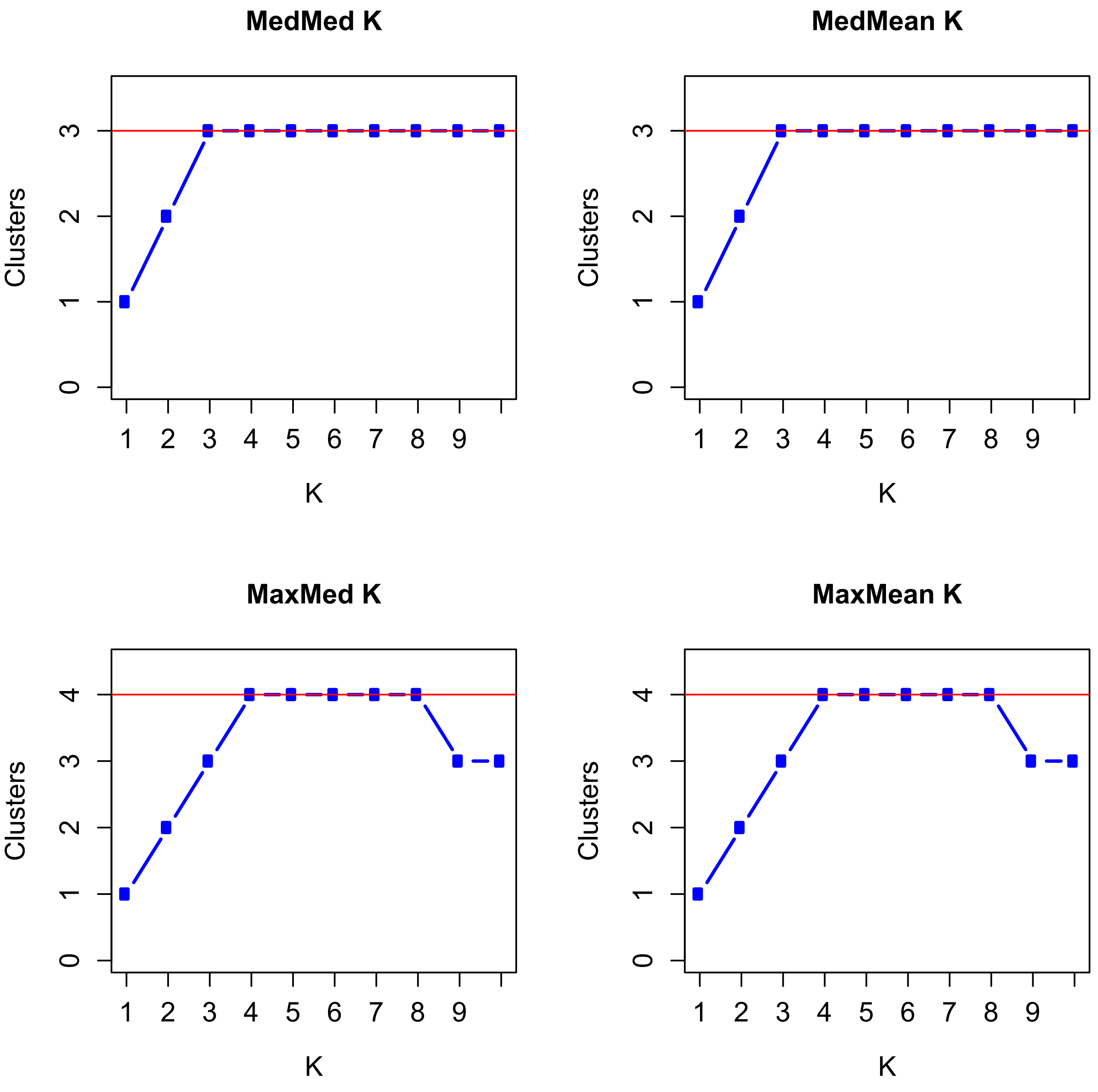

### Figure S7

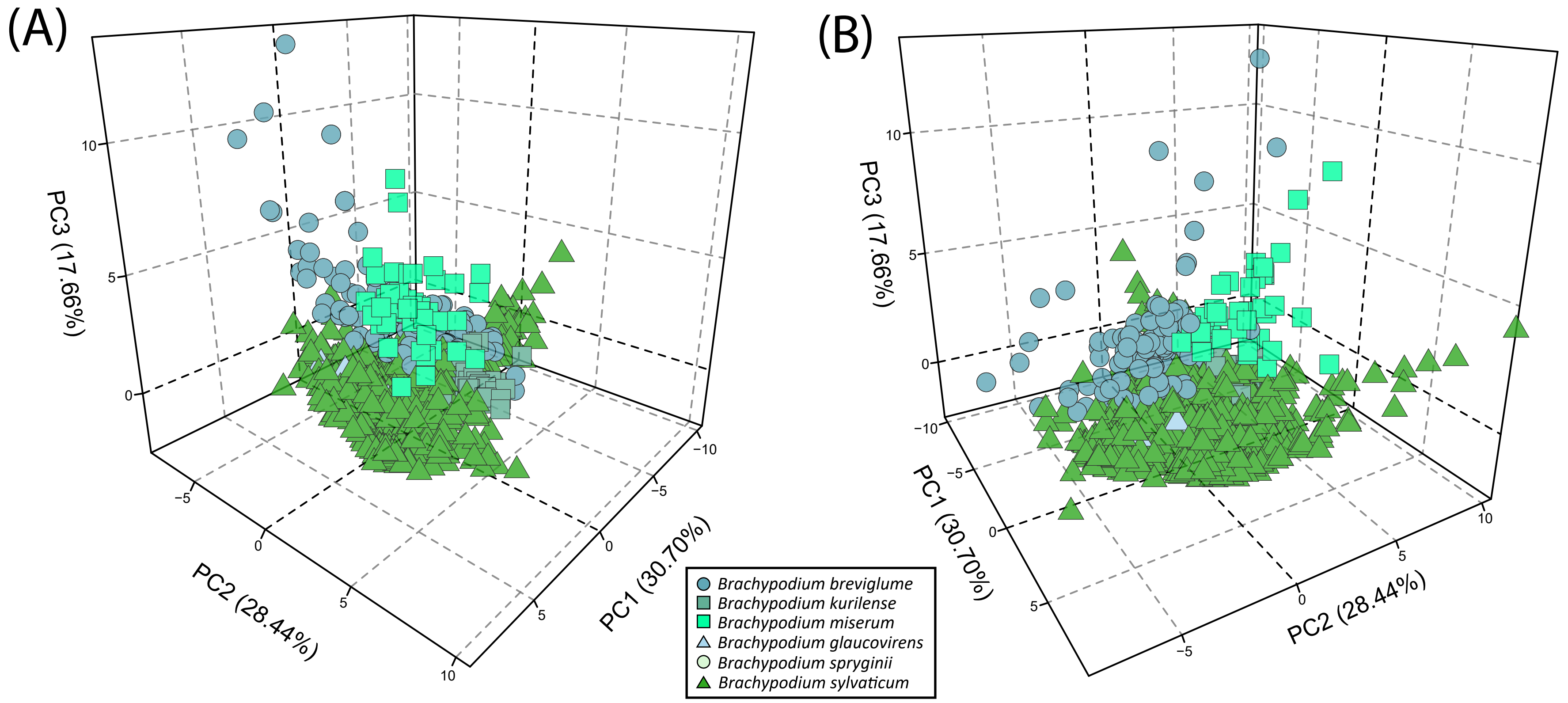
