## Appendix 1 for "A Palearctic divide, niche conservatism and host-fungal endophyte interactions shaped the phylogeography of the grass *Brachypodium sylvaticum*"

**Appendix 1.** List of geographic coordinates and values of 19 bioclimatic variables for each of the 1,263 *Brachypodium sylvaticum* complex samples used in this study, extracted from the WorldClim database (2.5 arc-minute resolution). The dataset includes 169 *B. breviglume*, 4 *B. glaucovirens*, 29 *B. kurilense*, 49 *B. miserum*, 2 *B. spryginii*, and 1,009 *B. sylvaticum* samples. Samples shown in bold correspond to individuals for which genomic data were available; the remaining records were retrieved from GBIF based on taxonomic identity and geolocation.

| **Taxa** | **Ind** | **Latitude** | **Longitude** | **Bio1** | **Bio2** | **Bio3** | **Bio4** | **Bio5** | **Bio6** | **Bio7** | **Bio8** | **Bio9** | **Bio10** | **Bio11** | **Bio12** | **Bio13** | **Bio14** | **Bio15** | **Bio16** | **Bio17** | **Bio18** | **Bio19** |
| --- | --- | --- | --- | --- | --- | --- | --- | --- | --- | --- | --- | --- | --- | --- | --- | --- | --- | --- | --- | --- | --- | --- |
| ***B. breviglume*** | **Bbrev33H** | **30.36** | **69.00** | **18.44** | **13.79** | **38.65** | **818.22** | **35.38** | **-0.29** | **35.67** | **27.67** | **19.08** | **27.67** | **7.62** | **249** | **51** | **5** | **68.28** | **107** | **17** | **107** | **49** |
| ***B. breviglume*** | **Bbrev34H** | **31.80** | **83.47** | **-7.56** | **13.22** | **38.22** | **800.67** | **9.01** | **-25.59** | **34.60** | **2.10** | **-8.56** | **2.37** | **-17.09** | **101** | **28** | **2** | **95.25** | **64** | **8** | **60** | **15** |
| *B. breviglume* | Bbrev_1 | 29.19 | 90.70 | 0.75 | 14.42 | 44.75 | 634.01 | 14.91 | -17.32 | 32.23 | 8.22 | -6.64 | 8.22 | -7.35 | 307 | 87 | 1 | 120.24 | 220 | 5 | 220 | 5 |
| *B. breviglume* | Bbrev_2 | 29.76 | 70.74 | 25.33 | 14.39 | 39.44 | 809.78 | 41.94 | 5.46 | 36.48 | 32.41 | 20.30 | 33.98 | 14.46 | 197 | 59 | 1 | 108.57 | 129 | 10 | 80 | 27 |
| *B. breviglume* | Bbrev_3 | 31.40 | 109.60 | 15.56 | 8.38 | 27.32 | 809.05 | 31.01 | 0.32 | 30.69 | 24.67 | 5.24 | 25.27 | 5.24 | 1186 | 207 | 17 | 65.47 | 520 | 60 | 515 | 60 |
| *B. breviglume* | Bbrev_4 | 25.00 | 102.70 | 15.20 | 10.17 | 46.04 | 451.40 | 24.07 | 1.99 | 22.08 | 19.96 | 9.04 | 19.96 | 9.04 | 922 | 185 | 13 | 83.00 | 511 | 44 | 511 | 44 |
| *B. breviglume* | Bbrev_5 | 31.40 | 109.60 | 15.56 | 8.38 | 27.32 | 809.05 | 31.01 | 0.32 | 30.69 | 24.67 | 5.24 | 25.27 | 5.24 | 1186 | 207 | 17 | 65.47 | 520 | 60 | 515 | 60 |
| *B. breviglume* | Bbrev_6 | 29.60 | 94.40 | 8.75 | 11.64 | 43.20 | 572.18 | 20.85 | -6.09 | 26.94 | 15.41 | 1.32 | 15.41 | 1.32 | 669 | 133 | 2 | 91.71 | 368 | 9 | 368 | 9 |
| *B. breviglume* | Bbrev_7 | 27.40 | 101.50 | 11.86 | 12.65 | 49.54 | 474.75 | 22.54 | -3.00 | 25.54 | 17.13 | 5.60 | 17.13 | 5.60 | 832 | 178 | 5 | 89.38 | 467 | 22 | 467 | 22 |
| *B. breviglume* | Bbrev_8 | 31.40 | 109.60 | 15.56 | 8.38 | 27.32 | 809.05 | 31.01 | 0.32 | 30.69 | 24.67 | 5.24 | 25.27 | 5.24 | 1186 | 207 | 17 | 65.47 | 520 | 60 | 515 | 60 |
| *B. breviglume* | Bbrev_9 | 25.00 | 102.70 | 15.20 | 10.17 | 46.04 | 451.40 | 24.07 | 1.99 | 22.08 | 19.96 | 9.04 | 19.96 | 9.04 | 922 | 185 | 13 | 83.00 | 511 | 44 | 511 | 44 |
| *B. breviglume* | Bbrev_10 | 30.40 | 102.80 | 12.66 | 9.20 | 33.17 | 671.09 | 25.70 | -2.03 | 27.74 | 20.46 | 3.79 | 20.46 | 3.79 | 1023 | 201 | 12 | 84.16 | 551 | 40 | 551 | 40 |
| *B. breviglume* | Bbrev_11 | 25.00 | 102.70 | 15.20 | 10.17 | 46.04 | 451.40 | 24.07 | 1.99 | 22.08 | 19.96 | 9.04 | 19.96 | 9.04 | 922 | 185 | 13 | 83.00 | 511 | 44 | 511 | 44 |
| *B. breviglume* | Bbrev_12 | 31.40 | 109.60 | 15.56 | 8.38 | 27.32 | 809.05 | 31.01 | 0.32 | 30.69 | 24.67 | 5.24 | 25.27 | 5.24 | 1186 | 207 | 17 | 65.47 | 520 | 60 | 515 | 60 |
| *B. breviglume* | Bbrev_13 | 28.00 | 102.80 | 11.37 | 11.02 | 42.77 | 538.02 | 22.55 | -3.22 | 25.77 | 17.54 | 4.23 | 17.54 | 4.23 | 913 | 188 | 8 | 90.83 | 516 | 29 | 516 | 29 |
| *B. breviglume* | Bbrev_14 | 31.40 | 109.60 | 15.56 | 8.38 | 27.32 | 809.05 | 31.01 | 0.32 | 30.69 | 24.67 | 5.24 | 25.27 | 5.24 | 1186 | 207 | 17 | 65.47 | 520 | 60 | 515 | 60 |
| *B. breviglume* | Bbrev_15 | 23.30 | 103.50 | 15.33 | 9.19 | 46.00 | 426.07 | 23.64 | 3.65 | 19.98 | 19.78 | 9.45 | 19.78 | 9.45 | 1351 | 274 | 12 | 85.47 | 759 | 54 | 759 | 54 |
| *B. breviglume* | Bbrev_16 | 31.50 | 98.30 | -1.47 | 12.46 | 39.55 | 695.41 | 12.73 | -18.76 | 31.49 | 6.78 | -9.26 | 6.78 | -10.20 | 617 | 133 | 5 | 92.75 | 356 | 19 | 356 | 19 |
| *B. breviglume* | Bbrev_17 | 31.50 | 102.10 | 10.82 | 13.88 | 44.30 | 625.08 | 25.06 | -6.28 | 31.33 | 16.80 | 2.65 | 18.00 | 2.65 | 623 | 127 | 2 | 86.58 | 312 | 11 | 308 | 11 |
| *B. breviglume* | Bbrev_18 | 31.80 | 110.70 | 12.17 | 9.05 | 29.18 | 811.60 | 27.22 | -3.81 | 31.03 | 21.18 | 1.88 | 22.01 | 1.88 | 1119 | 194 | 16 | 65.03 | 496 | 57 | 489 | 57 |
| *B. breviglume* | Bbrev_19 | 31.50 | 102.10 | 10.82 | 13.88 | 44.30 | 625.08 | 25.06 | -6.28 | 31.33 | 16.80 | 2.65 | 18.00 | 2.65 | 623 | 127 | 2 | 86.58 | 312 | 11 | 308 | 11 |
| *B. breviglume* | Bbrev_20 | 29.30 | 108.50 | 13.50 | 7.27 | 25.89 | 755.76 | 27.64 | -0.42 | 28.07 | 20.64 | 3.89 | 22.56 | 3.89 | 1388 | 215 | 24 | 59.77 | 599 | 84 | 556 | 84 |
| *B. breviglume* | Bbrev_21 | 34.70 | 101.60 | 0.30 | 15.38 | 39.96 | 797.26 | 16.34 | -22.14 | 38.48 | 9.47 | -10.19 | 9.47 | -10.19 | 595 | 123 | 2 | 91.85 | 329 | 10 | 329 | 10 |
| *B. breviglume* | Bbrev_22 | 28.30 | 103.10 | 11.36 | 9.80 | 37.83 | 581.61 | 23.06 | -2.85 | 25.92 | 18.05 | 3.62 | 18.05 | 3.62 | 894 | 175 | 9 | 88.19 | 499 | 28 | 499 | 28 |
| *B. breviglume* | Bbrev_23 | 27.70 | 102.80 | 10.50 | 11.03 | 44.27 | 520.13 | 21.16 | -3.77 | 24.93 | 16.42 | 3.63 | 16.42 | 3.63 | 908 | 191 | 7 | 92.46 | 517 | 24 | 517 | 24 |
| *B. breviglume* | Bbrev_24 | 24.70 | 101.60 | 16.26 | 11.24 | 47.66 | 444.09 | 25.76 | 2.18 | 23.58 | 20.87 | 10.20 | 20.87 | 10.20 | 889 | 177 | 13 | 83.52 | 491 | 43 | 491 | 43 |
| *B. breviglume* | Bbrev_25 | 31.50 | 102.10 | 10.82 | 13.88 | 44.30 | 625.08 | 25.06 | -6.28 | 31.33 | 16.80 | 2.65 | 18.00 | 2.65 | 623 | 127 | 2 | 86.58 | 312 | 11 | 308 | 11 |
| *B. breviglume* | Bbrev_26 | 31.80 | 110.70 | 12.17 | 9.05 | 29.18 | 811.60 | 27.22 | -3.81 | 31.03 | 21.18 | 1.88 | 22.01 | 1.88 | 1119 | 194 | 16 | 65.03 | 496 | 57 | 489 | 57 |
| *B. breviglume* | Bbrev_27 | 32.90 | 101.70 | 3.33 | 15.44 | 43.10 | 711.70 | 18.77 | -17.04 | 35.82 | 11.45 | -5.18 | 11.45 | -6.03 | 725 | 132 | 4 | 85.74 | 369 | 17 | 369 | 17 |
| *B. breviglume* | Bbrev_28 | 31.20 | 69.60 | 16.35 | 12.77 | 36.95 | 822.09 | 32.52 | -2.05 | 34.57 | 25.67 | 17.29 | 25.67 | 5.57 | 326 | 69 | 5 | 70.06 | 140 | 24 | 140 | 69 |
| *B. breviglume* | Bbrev_29 | 44.10 | 80.90 | 8.60 | 13.84 | 28.51 | 1259.93 | 31.85 | -16.68 | 48.54 | 17.17 | -5.60 | 22.84 | -8.20 | 254 | 32 | 14 | 27.86 | 87 | 46 | 73 | 47 |
| *B. breviglume* | Bbrev_30 | 27.90 | 101.30 | 11.23 | 11.83 | 47.21 | 502.11 | 21.84 | -3.22 | 25.06 | 16.94 | 4.67 | 16.94 | 4.67 | 815 | 174 | 4 | 90.59 | 459 | 19 | 459 | 19 |
| *B. breviglume* | Bbrev_31 | 31.40 | 103.20 | 4.93 | 11.14 | 37.52 | 645.90 | 18.58 | -11.12 | 29.70 | 11.12 | -3.49 | 12.51 | -3.49 | 738 | 134 | 4 | 80.34 | 355 | 17 | 348 | 17 |
| *B. breviglume* | Bbrev_32 | 29.20 | 94.20 | 8.02 | 11.30 | 41.89 | 583.05 | 20.10 | -6.87 | 26.97 | 14.76 | 0.45 | 14.76 | 0.45 | 673 | 138 | 2 | 92.44 | 378 | 10 | 378 | 10 |
| *B. breviglume* | Bbrev_33 | 31.20 | 96.60 | 2.90 | 13.29 | 41.89 | 680.70 | 17.25 | -14.49 | 31.74 | 10.91 | -4.87 | 10.91 | -5.76 | 569 | 125 | 4 | 95.26 | 335 | 15 | 335 | 15 |
| *B. breviglume* | Bbrev_34 | 30.90 | 98.30 | 3.86 | 15.07 | 46.49 | 645.41 | 18.17 | -14.24 | 32.41 | 11.41 | -4.37 | 11.41 | -4.37 | 556 | 120 | 3 | 93.72 | 323 | 13 | 323 | 13 |
| *B. breviglume* | Bbrev_35 | 30.40 | 102.80 | 12.66 | 9.20 | 33.17 | 671.09 | 25.70 | -2.03 | 27.74 | 20.46 | 3.79 | 20.46 | 3.79 | 1023 | 201 | 12 | 84.16 | 551 | 40 | 551 | 40 |
| *B. breviglume* | Bbrev_36 | 25.40 | 103.00 | 13.93 | 10.89 | 47.34 | 444.64 | 23.41 | 0.42 | 23.00 | 18.58 | 7.91 | 18.58 | 7.91 | 911 | 182 | 12 | 84.34 | 505 | 39 | 505 | 39 |
| *B. breviglume* | Bbrev_37 | 29.60 | 103.50 | 17.75 | 7.39 | 27.90 | 698.10 | 30.55 | 4.06 | 26.49 | 25.24 | 8.61 | 25.88 | 8.61 | 1577 | 397 | 18 | 97.46 | 942 | 60 | 937 | 60 |
| *B. breviglume* | Bbrev_38 | 30.90 | 101.90 | 11.78 | 12.05 | 40.60 | 635.89 | 25.29 | -4.38 | 29.67 | 19.03 | 3.40 | 19.03 | 3.40 | 650 | 134 | 2 | 90.75 | 347 | 9 | 347 | 9 |
| *B. breviglume* | Bbrev_39 | 25.40 | 103.00 | 13.93 | 10.89 | 47.34 | 444.64 | 23.41 | 0.42 | 23.00 | 18.58 | 7.91 | 18.58 | 7.91 | 911 | 182 | 12 | 84.34 | 505 | 39 | 505 | 39 |
| *B. breviglume* | Bbrev_40 | 26.10 | 101.70 | 18.19 | 12.26 | 48.03 | 466.67 | 28.92 | 3.40 | 25.52 | 23.04 | 11.97 | 23.05 | 11.97 | 737 | 158 | 7 | 91.36 | 426 | 23 | 339 | 23 |
| *B. breviglume* | Bbrev_41 | 31.80 | 98.60 | 3.53 | 14.20 | 43.69 | 674.01 | 18.22 | -14.30 | 32.51 | 11.43 | -4.33 | 11.43 | -5.07 | 569 | 118 | 4 | 94.14 | 327 | 14 | 327 | 14 |
| *B. breviglume* | Bbrev_42 | 32.50 | 107.90 | 12.55 | 8.29 | 27.95 | 787.95 | 27.42 | -2.24 | 29.67 | 21.26 | 2.62 | 22.20 | 2.62 | 988 | 174 | 10 | 69.85 | 464 | 38 | 423 | 38 |
| *B. breviglume* | Bbrev_43 | 31.40 | 100.70 | 6.26 | 15.06 | 45.37 | 654.10 | 20.65 | -12.55 | 33.20 | 13.69 | -2.30 | 13.69 | -2.30 | 624 | 131 | 2 | 96.03 | 348 | 9 | 348 | 9 |
| *B. breviglume* | Bbrev_44 | 32.30 | 100.30 | 1.12 | 15.98 | 44.10 | 720.26 | 16.39 | -19.85 | 36.24 | 9.45 | -7.28 | 9.45 | -8.26 | 677 | 134 | 3 | 90.47 | 363 | 14 | 363 | 14 |
| *B. breviglume* | Bbrev_45 | 35.80 | 108.00 | 9.31 | 10.73 | 28.86 | 966.98 | 27.04 | -10.15 | 37.19 | 19.41 | -3.08 | 20.86 | -3.08 | 544 | 106 | 4 | 83.51 | 306 | 14 | 267 | 14 |
| *B. breviglume* | Bbrev_46 | 27.20 | 99.30 | 12.61 | 10.82 | 43.63 | 531.39 | 23.18 | -1.62 | 24.81 | 18.27 | 6.69 | 18.79 | 5.78 | 949 | 153 | 13 | 61.03 | 400 | 57 | 397 | 95 |
| *B. breviglume* | Bbrev_47 | 30.10 | 102.00 | 7.30 | 10.99 | 39.14 | 610.32 | 19.92 | -8.15 | 28.07 | 14.43 | -0.64 | 14.43 | -0.64 | 784 | 164 | 6 | 90.74 | 424 | 19 | 424 | 19 |
| *B. breviglume* | Bbrev_48 | 31.80 | 98.60 | 3.53 | 14.20 | 43.69 | 674.01 | 18.22 | -14.30 | 32.51 | 11.43 | -4.33 | 11.43 | -5.07 | 569 | 118 | 4 | 94.14 | 327 | 14 | 327 | 14 |
| *B. breviglume* | Bbrev_49 | 31.40 | 109.60 | 15.56 | 8.38 | 27.32 | 809.05 | 31.01 | 0.32 | 30.69 | 24.67 | 5.24 | 25.27 | 5.24 | 1186 | 207 | 17 | 65.47 | 520 | 60 | 515 | 60 |
| *B. breviglume* | Bbrev_50 | 31.20 | 98.80 | 5.39 | 13.20 | 43.06 | 638.81 | 19.29 | -11.35 | 30.64 | 12.84 | -2.03 | 12.84 | -2.80 | 575 | 122 | 3 | 95.20 | 334 | 13 | 334 | 13 |
| *B. breviglume* | Bbrev_51 | 27.20 | 99.30 | 12.61 | 10.82 | 43.63 | 531.39 | 23.18 | -1.62 | 24.81 | 18.27 | 6.69 | 18.79 | 5.78 | 949 | 153 | 13 | 61.03 | 400 | 57 | 397 | 95 |
| *B. breviglume* | Bbrev_52 | 28.20 | 106.90 | 13.28 | 7.42 | 27.10 | 728.14 | 27.43 | 0.04 | 27.40 | 20.27 | 3.90 | 21.94 | 3.90 | 1090 | 179 | 17 | 66.32 | 498 | 55 | 478 | 55 |
| *B. breviglume* | Bbrev_53 | 30.10 | 102.00 | 7.30 | 10.99 | 39.14 | 610.32 | 19.92 | -8.15 | 28.07 | 14.43 | -0.64 | 14.43 | -0.64 | 784 | 164 | 6 | 90.74 | 424 | 19 | 424 | 19 |
| *B. breviglume* | Bbrev_54 | 29.60 | 103.50 | 17.75 | 7.39 | 27.90 | 698.10 | 30.55 | 4.06 | 26.49 | 25.24 | 8.61 | 25.88 | 8.61 | 1577 | 397 | 18 | 97.46 | 942 | 60 | 937 | 60 |
| *B. breviglume* | Bbrev_55 | 32.10 | 101.10 | -0.96 | 13.54 | 40.67 | 698.70 | 13.52 | -19.78 | 33.30 | 7.16 | -8.92 | 7.16 | -9.92 | 730 | 143 | 3 | 89.20 | 390 | 16 | 390 | 16 |
| *B. breviglume* | Bbrev_56 | 32.10 | 103.00 | 3.07 | 12.25 | 40.19 | 630.01 | 16.76 | -13.73 | 30.49 | 9.03 | -5.09 | 10.47 | -5.09 | 742 | 132 | 4 | 79.43 | 351 | 20 | 349 | 20 |
| *B. breviglume* | Bbrev_57 | 33.90 | 106.50 | 11.09 | 8.66 | 28.09 | 818.64 | 26.32 | -4.52 | 30.84 | 19.93 | 0.68 | 21.00 | 0.68 | 707 | 131 | 4 | 78.86 | 370 | 16 | 325 | 16 |
| *B. breviglume* | Bbrev_58 | 28.00 | 98.50 | 2.90 | 8.74 | 35.91 | 589.34 | 13.83 | -10.52 | 24.35 | 9.71 | -2.91 | 9.71 | -4.50 | 802 | 146 | 13 | 63.63 | 374 | 53 | 374 | 61 |
| *B. breviglume* | Bbrev_59 | 28.50 | 98.90 | 4.43 | 10.72 | 41.59 | 581.52 | 15.92 | -9.85 | 25.77 | 11.23 | -1.61 | 11.23 | -2.86 | 638 | 124 | 8 | 68.44 | 304 | 35 | 304 | 37 |
| *B. breviglume* | Bbrev_60 | 36.30 | 74.60 | 8.73 | 12.23 | 33.63 | 888.19 | 27.73 | -8.62 | 36.35 | 7.96 | 3.98 | 19.61 | -2.42 | 163 | 34 | 2 | 60.24 | 76 | 15 | 43 | 23 |
| *B. breviglume* | Bbrev_61 | 35.90 | 74.30 | 10.56 | 10.89 | 30.31 | 920.46 | 29.36 | -6.57 | 35.94 | 9.90 | 5.52 | 21.89 | -1.01 | 240 | 43 | 4 | 54.63 | 109 | 25 | 60 | 40 |
| *B. breviglume* | Bbrev_62 | 34.30 | 70.00 | 17.87 | 13.74 | 37.48 | 873.28 | 35.87 | -0.80 | 36.67 | 12.26 | 18.72 | 28.35 | 6.82 | 464 | 98 | 12 | 76.70 | 251 | 37 | 77 | 110 |
| *B. breviglume* | Bbrev_63 | 36.90 | 101.30 | 0.17 | 12.84 | 34.73 | 846.27 | 16.68 | -20.28 | 36.97 | 10.01 | -9.59 | 10.01 | -10.81 | 485 | 107 | 1 | 98.90 | 289 | 5 | 289 | 5 |
| *B. breviglume* | Bbrev_64 | 35.90 | 71.80 | 14.39 | 13.28 | 36.25 | 898.74 | 34.26 | -2.38 | 36.64 | 8.26 | 25.62 | 25.62 | 3.29 | 383 | 81 | 9 | 75.27 | 200 | 29 | 29 | 113 |
| *B. breviglume* | Bbrev_65 | 29.70 | 94.10 | 2.29 | 11.01 | 39.24 | 627.80 | 15.16 | -12.90 | 28.06 | 9.71 | -4.47 | 9.71 | -5.67 | 580 | 123 | 3 | 91.99 | 330 | 14 | 330 | 14 |
| *B. breviglume* | Bbrev_66 | 28.10 | 83.90 | 19.69 | 11.31 | 46.52 | 497.43 | 29.88 | 5.58 | 24.30 | 24.62 | 13.55 | 24.62 | 12.71 | 2477 | 624 | 6 | 110.67 | 1628 | 39 | 1628 | 50 |
| *B. breviglume* | Bbrev_67 | 29.70 | 91.10 | 6.54 | 14.35 | 44.60 | 637.56 | 20.76 | -11.42 | 32.18 | 13.91 | -1.81 | 13.91 | -1.81 | 412 | 115 | 1 | 126.81 | 299 | 3 | 299 | 3 |
| *B. breviglume* | Bbrev_68 | 32.00 | 101.10 | -0.40 | 13.83 | 41.60 | 686.63 | 13.98 | -19.27 | 33.25 | 7.58 | -9.20 | 7.58 | -9.20 | 721 | 144 | 3 | 90.10 | 387 | 14 | 387 | 14 |
| *B. breviglume* | Bbrev_69 | 32.30 | 77.20 | 9.38 | 9.60 | 34.53 | 660.75 | 22.25 | -5.55 | 27.80 | 15.66 | 6.55 | 16.38 | 0.46 | 1023 | 135 | 20 | 43.05 | 351 | 102 | 305 | 240 |
| *B. breviglume* | Bbrev_70 | 35.10 | 69.60 | 7.53 | 10.91 | 32.49 | 830.36 | 24.30 | -9.29 | 33.59 | 1.75 | 16.65 | 17.64 | -3.03 | 776 | 178 | 6 | 98.04 | 479 | 22 | 24 | 266 |
| *B. breviglume* | Bbrev_71 | 29.60 | 94.90 | 6.57 | 11.00 | 41.14 | 587.11 | 18.62 | -8.13 | 26.75 | 13.34 | -1.01 | 13.34 | -1.01 | 700 | 140 | 3 | 89.19 | 383 | 13 | 383 | 13 |
| *B. breviglume* | Bbrev_72 | 29.00 | 93.90 | 2.03 | 11.35 | 41.16 | 606.71 | 13.92 | -13.64 | 27.57 | 9.15 | -5.66 | 9.15 | -5.66 | 519 | 115 | 2 | 94.37 | 304 | 11 | 304 | 11 |
| *B. breviglume* | Bbrev_73 | 34.60 | 73.10 | 15.73 | 12.07 | 38.28 | 718.61 | 31.36 | -0.16 | 31.52 | 22.90 | 12.43 | 24.09 | 6.26 | 1168 | 207 | 25 | 58.55 | 486 | 113 | 479 | 226 |
| *B. breviglume* | Bbrev_74 | 28.90 | 93.20 | -4.64 | 11.15 | 39.78 | 627.03 | 7.14 | -20.90 | 28.04 | 2.82 | -11.18 | 2.82 | -12.38 | 431 | 97 | 4 | 92.11 | 253 | 16 | 253 | 16 |
| *B. breviglume* | Bbrev_75 | 34.50 | 69.40 | 12.18 | 13.78 | 34.01 | 977.18 | 32.22 | -8.29 | 40.51 | 6.47 | 18.94 | 23.88 | -0.53 | 403 | 91 | 6 | 88.34 | 239 | 24 | 28 | 143 |
| *B. breviglume* | Bbrev_76 | 35.30 | 72.60 | 12.74 | 11.18 | 34.80 | 815.10 | 29.16 | -2.96 | 32.12 | 11.85 | 8.97 | 22.57 | 2.36 | 670 | 92 | 16 | 44.20 | 248 | 79 | 192 | 142 |
| *B. breviglume* | Bbrev_77 | 25.80 | 100.10 | 14.51 | 10.83 | 47.75 | 450.89 | 23.83 | 1.16 | 22.67 | 19.46 | 9.07 | 19.46 | 8.50 | 848 | 142 | 14 | 63.83 | 394 | 61 | 394 | 61 |
| *B. breviglume* | Bbrev_78 | 34.90 | 70.70 | 9.94 | 10.42 | 33.13 | 802.43 | 25.89 | -5.55 | 31.44 | 4.03 | 11.22 | 19.71 | -0.10 | 792 | 168 | 20 | 77.78 | 428 | 72 | 92 | 218 |
| *B. breviglume* | Bbrev_79 | 28.30 | 99.80 | 2.48 | 10.85 | 40.99 | 588.69 | 13.79 | -12.69 | 26.48 | 9.31 | -3.75 | 9.31 | -4.97 | 665 | 136 | 6 | 76.60 | 342 | 29 | 342 | 30 |
| *B. breviglume* | Bbrev_80 | 35.70 | 106.20 | 3.44 | 9.72 | 29.56 | 853.64 | 19.12 | -13.76 | 32.88 | 12.60 | -7.39 | 13.67 | -7.39 | 531 | 105 | 3 | 83.51 | 290 | 13 | 270 | 13 |
| *B. breviglume* | Bbrev_81 | 30.40 | 78.30 | 14.56 | 9.48 | 38.64 | 530.98 | 25.88 | 1.35 | 24.53 | 18.86 | 11.46 | 20.10 | 7.32 | 2172 | 651 | 14 | 126.95 | 1587 | 124 | 862 | 182 |
| *B. breviglume* | Bbrev_82 | 34.10 | 75.00 | 6.77 | 10.88 | 33.51 | 822.31 | 22.03 | -10.44 | 32.47 | 0.54 | 3.56 | 16.35 | -3.66 | 1067 | 156 | 31 | 45.98 | 426 | 139 | 220 | 293 |
| *B. breviglume* | Bbrev_83 | 31.00 | 78.40 | 6.99 | 8.96 | 36.01 | 589.20 | 18.18 | -6.71 | 24.88 | 12.75 | 4.32 | 13.30 | -0.90 | 1446 | 278 | 26 | 66.75 | 722 | 152 | 617 | 297 |
| *B. breviglume* | Bbrev_84 | 27.00 | 100.20 | 10.20 | 11.36 | 46.49 | 488.23 | 20.61 | -3.82 | 24.43 | 15.83 | 4.53 | 15.83 | 3.91 | 791 | 141 | 10 | 65.92 | 370 | 44 | 370 | 53 |
| *B. breviglume* | Bbrev_85 | 29.80 | 96.30 | -3.82 | 10.48 | 36.59 | 676.61 | 9.05 | -19.59 | 28.64 | 4.08 | -10.92 | 4.08 | -12.30 | 651 | 136 | 7 | 84.53 | 355 | 26 | 355 | 26 |
| *B. breviglume* | Bbrev_86 | 30.90 | 101.90 | 11.78 | 12.05 | 40.60 | 635.89 | 25.29 | -4.38 | 29.67 | 19.03 | 3.40 | 19.03 | 3.40 | 650 | 134 | 2 | 90.75 | 347 | 9 | 347 | 9 |
| *B. breviglume* | Bbrev_87 | 28.70 | 97.50 | 3.08 | 10.68 | 41.19 | 575.52 | 14.77 | -11.16 | 25.92 | 9.74 | -2.72 | 9.74 | -4.14 | 815 | 161 | 9 | 76.77 | 425 | 36 | 425 | 40 |
| *B. breviglume* | Bbrev_88 | 33.30 | 105.60 | 11.41 | 8.58 | 29.12 | 764.16 | 25.84 | -3.61 | 29.46 | 19.61 | 1.62 | 20.63 | 1.62 | 697 | 137 | 4 | 82.09 | 368 | 15 | 345 | 15 |
| *B. breviglume* | Bbrev_89 | 30.10 | 102.00 | 7.30 | 10.99 | 39.14 | 610.32 | 19.92 | -8.15 | 28.07 | 14.43 | -0.64 | 14.43 | -0.64 | 784 | 164 | 6 | 90.74 | 424 | 19 | 424 | 19 |
| *B. breviglume* | Bbrev_90 | 32.60 | 103.60 | 4.89 | 12.03 | 38.39 | 676.55 | 19.31 | -12.02 | 31.34 | 11.35 | -3.92 | 12.84 | -3.92 | 714 | 115 | 3 | 76.20 | 329 | 17 | 317 | 17 |
| *B. breviglume* | Bbrev_91 | 27.40 | 101.50 | 11.86 | 12.65 | 49.54 | 474.75 | 22.54 | -3.00 | 25.54 | 17.13 | 5.60 | 17.13 | 5.60 | 832 | 178 | 5 | 89.38 | 467 | 22 | 467 | 22 |
| *B. breviglume* | Bbrev_92 | 29.90 | 93.20 | 4.34 | 12.17 | 40.42 | 653.64 | 17.75 | -12.35 | 30.10 | 11.87 | -3.16 | 11.87 | -4.18 | 489 | 115 | 2 | 103.37 | 304 | 8 | 304 | 8 |
| *B. breviglume* | Bbrev_93 | 31.80 | 98.60 | 3.53 | 14.20 | 43.69 | 674.01 | 18.22 | -14.30 | 32.51 | 11.43 | -4.33 | 11.43 | -5.07 | 569 | 118 | 4 | 94.14 | 327 | 14 | 327 | 14 |
| *B. breviglume* | Bbrev_94 | 30.90 | 102.50 | -0.56 | 12.40 | 41.35 | 611.98 | 12.40 | -17.59 | 29.99 | 6.66 | -8.31 | 6.66 | -8.31 | 783 | 153 | 4 | 87.16 | 406 | 16 | 406 | 16 |
| *B. breviglume* | Bbrev_95 | 29.70 | 111.00 | 15.96 | 7.49 | 24.94 | 819.47 | 30.85 | 0.82 | 30.04 | 23.79 | 5.59 | 25.82 | 5.59 | 1296 | 198 | 28 | 55.68 | 552 | 101 | 535 | 101 |
| *B. breviglume* | Bbrev_96 | 31.40 | 100.70 | 6.26 | 15.06 | 45.37 | 654.10 | 20.65 | -12.55 | 33.20 | 13.69 | -2.30 | 13.69 | -2.30 | 624 | 131 | 2 | 96.03 | 348 | 9 | 348 | 9 |
| *B. breviglume* | Bbrev_97 | 31.60 | 100.00 | 6.68 | 15.06 | 45.53 | 662.33 | 21.17 | -11.92 | 33.09 | 14.20 | -1.97 | 14.20 | -1.97 | 630 | 130 | 4 | 92.89 | 345 | 14 | 345 | 14 |
| *B. breviglume* | Bbrev_98 | 31.40 | 109.60 | 15.56 | 8.38 | 27.32 | 809.05 | 31.01 | 0.32 | 30.69 | 24.67 | 5.24 | 25.27 | 5.24 | 1186 | 207 | 17 | 65.47 | 520 | 60 | 515 | 60 |
| *B. breviglume* | Bbrev_99 | 27.90 | 101.30 | 11.23 | 11.83 | 47.21 | 502.11 | 21.84 | -3.22 | 25.06 | 16.94 | 4.67 | 16.94 | 4.67 | 815 | 174 | 4 | 90.59 | 459 | 19 | 459 | 19 |
| *B. breviglume* | Bbrev_100 | 24.90 | 105.80 | 16.98 | 8.02 | 34.58 | 575.36 | 27.51 | 4.32 | 23.19 | 23.32 | 9.17 | 23.32 | 9.17 | 1293 | 250 | 12 | 84.75 | 723 | 57 | 723 | 57 |
| *B. breviglume* | Bbrev_101 | 35.80 | 108.00 | 9.31 | 10.73 | 28.86 | 966.98 | 27.04 | -10.15 | 37.19 | 19.41 | -3.08 | 20.86 | -3.08 | 544 | 106 | 4 | 83.51 | 306 | 14 | 267 | 14 |
| *B. breviglume* | Bbrev_102 | 27.90 | 102.20 | 17.04 | 11.04 | 45.66 | 497.68 | 27.28 | 3.10 | 24.19 | 22.49 | 10.36 | 22.49 | 10.36 | 943 | 225 | 4 | 101.08 | 575 | 16 | 575 | 16 |
| *B. breviglume* | Bbrev_103 | 31.80 | 108.60 | 12.37 | 8.16 | 27.45 | 787.55 | 27.36 | -2.38 | 29.73 | 21.15 | 2.39 | 21.93 | 2.39 | 1140 | 189 | 14 | 65.43 | 500 | 52 | 475 | 52 |
| *B. breviglume* | Bbrev_104 | 35.80 | 104.10 | 5.68 | 10.99 | 29.91 | 924.18 | 23.00 | -13.75 | 36.75 | 15.40 | -6.35 | 16.53 | -6.35 | 408 | 88 | 1 | 88.36 | 228 | 7 | 218 | 7 |
| *B. breviglume* | Bbrev_105 | 25.70 | 101.30 | 16.46 | 11.97 | 48.21 | 460.86 | 26.81 | 1.97 | 24.84 | 21.28 | 10.27 | 21.28 | 10.27 | 751 | 148 | 10 | 82.08 | 406 | 34 | 406 | 34 |
| *B. breviglume* | Bbrev_106 | 31.30 | 110.80 | 13.77 | 8.76 | 28.24 | 820.80 | 28.89 | -2.15 | 31.04 | 23.67 | 3.36 | 23.67 | 3.36 | 1150 | 202 | 17 | 64.80 | 511 | 62 | 511 | 62 |
| *B. breviglume* | Bbrev_107 | 34.70 | 104.90 | 6.41 | 9.70 | 28.60 | 855.60 | 22.86 | -11.06 | 33.92 | 15.51 | -4.67 | 16.64 | -4.67 | 590 | 108 | 3 | 81.32 | 303 | 13 | 290 | 13 |
| *B. breviglume* | Bbrev_108 | 33.30 | 106.20 | 13.65 | 8.11 | 27.49 | 791.82 | 28.32 | -1.18 | 29.50 | 22.18 | 3.59 | 23.27 | 3.59 | 774 | 158 | 5 | 83.10 | 420 | 18 | 382 | 18 |
| *B. breviglume* | Bbrev_109 | 31.20 | 98.80 | 5.39 | 13.20 | 43.06 | 638.81 | 19.29 | -11.35 | 30.64 | 12.84 | -2.03 | 12.84 | -2.80 | 575 | 122 | 3 | 95.20 | 334 | 13 | 334 | 13 |
| *B. breviglume* | Bbrev_110 | 30.10 | 102.80 | 16.19 | 8.12 | 30.42 | 679.18 | 28.86 | 2.17 | 26.69 | 23.40 | 7.22 | 24.06 | 7.22 | 1412 | 344 | 19 | 94.03 | 835 | 62 | 816 | 62 |
| *B. breviglume* | Bbrev_111 | 30.90 | 100.30 | 4.64 | 13.67 | 43.88 | 643.73 | 18.11 | -13.03 | 31.14 | 11.97 | -3.73 | 11.97 | -3.73 | 610 | 129 | 2 | 99.86 | 360 | 8 | 360 | 8 |
| *B. breviglume* | Bbrev_112 | 33.30 | 104.20 | 6.67 | 10.46 | 33.81 | 718.53 | 21.07 | -9.86 | 30.93 | 14.42 | -2.72 | 15.19 | -2.72 | 668 | 116 | 3 | 78.97 | 325 | 14 | 315 | 14 |
| *B. breviglume* | Bbrev_113 | 43.50 | 82.20 | 8.30 | 12.83 | 26.89 | 1266.30 | 30.15 | -17.57 | 47.72 | 20.79 | -8.95 | 22.11 | -8.95 | 202 | 31 | 8 | 42.69 | 83 | 26 | 76 | 26 |
| *B. breviglume* | Bbrev_114 | 33.80 | 106.10 | 12.87 | 9.30 | 30.05 | 802.15 | 28.06 | -2.89 | 30.96 | 21.47 | 2.68 | 22.59 | 2.68 | 688 | 131 | 4 | 80.79 | 365 | 15 | 329 | 15 |
| *B. breviglume* | Bbrev_115 | 25.00 | 102.70 | 15.20 | 10.17 | 46.04 | 451.40 | 24.07 | 1.99 | 22.08 | 19.96 | 9.04 | 19.96 | 9.04 | 922 | 185 | 13 | 83.00 | 511 | 44 | 511 | 44 |
| *B. breviglume* | Bbrev_116 | 31.20 | 97.20 | 6.32 | 14.69 | 44.88 | 672.73 | 21.04 | -11.69 | 32.73 | 14.20 | -2.27 | 14.20 | -2.27 | 509 | 111 | 3 | 96.13 | 302 | 12 | 302 | 12 |
| *B. breviglume* | Bbrev_117 | 32.40 | 113.40 | 15.23 | 9.27 | 27.80 | 903.91 | 30.92 | -2.42 | 33.34 | 26.06 | 3.64 | 26.06 | 3.64 | 975 | 186 | 15 | 63.13 | 437 | 66 | 437 | 66 |
| *B. breviglume* | Bbrev_118 | 34.00 | 112.00 | 12.43 | 10.28 | 30.46 | 871.01 | 28.36 | -5.39 | 33.76 | 21.64 | 1.18 | 22.94 | 1.18 | 741 | 151 | 8 | 73.49 | 374 | 30 | 346 | 30 |
| *B. breviglume* | Bbrev_119 | 36.50 | 101.60 | 3.87 | 13.96 | 36.74 | 855.98 | 20.34 | -17.66 | 38.00 | 13.61 | -6.34 | 13.61 | -7.38 | 436 | 98 | 1 | 97.06 | 254 | 5 | 254 | 5 |
| *B. breviglume* | Bbrev_120 | 27.20 | 105.70 | 12.82 | 7.84 | 30.17 | 670.52 | 25.55 | -0.44 | 25.99 | 19.30 | 4.00 | 20.64 | 4.00 | 1074 | 200 | 16 | 72.87 | 518 | 51 | 513 | 51 |
| *B. breviglume* | Bbrev_121 | 29.40 | 110.20 | 15.68 | 7.73 | 25.95 | 806.18 | 30.57 | 0.80 | 29.78 | 23.32 | 5.49 | 25.34 | 5.49 | 1425 | 241 | 28 | 59.85 | 640 | 101 | 596 | 101 |
| *B. breviglume* | Bbrev_122 | 27.50 | 88.90 | 6.92 | 11.87 | 47.36 | 507.38 | 17.54 | -7.54 | 25.07 | 12.79 | 1.51 | 12.79 | 0.38 | 741 | 127 | 5 | 67.82 | 346 | 27 | 346 | 45 |
| *B. breviglume* | Bbrev_123 | 37.00 | 103.10 | 3.28 | 11.78 | 31.59 | 919.90 | 20.98 | -16.31 | 37.29 | 14.24 | -8.54 | 14.24 | -8.54 | 349 | 78 | 1 | 96.65 | 207 | 5 | 207 | 5 |
| *B. breviglume* | Bbrev_124 | 28.70 | 83.60 | 10.56 | 10.51 | 46.97 | 452.78 | 20.70 | -1.69 | 22.38 | 14.78 | 5.04 | 15.32 | 4.43 | 1063 | 232 | 11 | 78.65 | 561 | 66 | 557 | 95 |
| *B. breviglume* | Bbrev_125 | 36.50 | 103.90 | 7.92 | 13.73 | 32.84 | 1011.35 | 27.64 | -14.17 | 41.81 | 18.34 | -5.27 | 19.74 | -5.27 | 293 | 69 | 1 | 92.17 | 170 | 6 | 166 | 6 |
| *B. breviglume* | Bbrev_126 | 34.00 | 74.40 | 3.44 | 10.10 | 31.16 | 855.82 | 18.73 | -13.70 | 32.43 | 13.01 | 0.32 | 13.30 | -7.49 | 1114 | 148 | 28 | 42.65 | 365 | 131 | 351 | 247 |
| *B. breviglume* | Bbrev_127 | 27.40 | 89.90 | 17.26 | 10.63 | 47.46 | 445.57 | 27.03 | 4.64 | 22.39 | 22.13 | 12.16 | 22.13 | 11.20 | 1343 | 267 | 10 | 85.25 | 741 | 46 | 741 | 66 |
| *B. breviglume* | Bbrev_128 | 28.30 | 103.10 | 11.36 | 9.80 | 37.83 | 581.61 | 23.06 | -2.85 | 25.92 | 18.05 | 3.62 | 18.05 | 3.62 | 894 | 175 | 9 | 88.19 | 499 | 28 | 499 | 28 |
| *B. breviglume* | Bbrev_129 | 29.00 | 101.50 | 8.78 | 13.11 | 47.61 | 537.30 | 20.28 | -7.26 | 27.54 | 14.93 | 1.77 | 14.93 | 1.77 | 876 | 187 | 2 | 98.34 | 503 | 9 | 503 | 9 |
| *B. breviglume* | Bbrev_130 | 32.80 | 102.50 | 2.83 | 15.13 | 42.73 | 680.89 | 18.19 | -17.21 | 35.40 | 10.80 | -6.00 | 10.80 | -6.00 | 707 | 124 | 3 | 83.42 | 348 | 16 | 348 | 16 |
| *B. breviglume* | Bbrev_131 | 31.70 | 103.90 | 9.10 | 10.14 | 34.77 | 685.44 | 22.88 | -6.29 | 29.17 | 16.51 | 0.15 | 17.19 | 0.15 | 798 | 144 | 5 | 79.00 | 391 | 23 | 388 | 23 |
| *B. breviglume* | Bbrev_132 | 33.00 | 98.10 | -0.67 | 14.75 | 40.98 | 769.86 | 15.16 | -20.82 | 35.98 | 8.26 | -9.70 | 8.26 | -10.54 | 540 | 117 | 4 | 94.19 | 313 | 15 | 313 | 16 |
| *B. breviglume* | Bbrev_133 | 32.30 | 100.30 | 1.12 | 15.98 | 44.10 | 720.26 | 16.39 | -19.85 | 36.24 | 9.45 | -7.28 | 9.45 | -8.26 | 677 | 134 | 3 | 90.47 | 363 | 14 | 363 | 14 |
| *B. breviglume* | Bbrev_134 | 27.70 | 103.30 | 14.44 | 9.60 | 37.36 | 588.11 | 26.14 | 0.44 | 25.70 | 21.18 | 6.62 | 21.18 | 6.62 | 892 | 175 | 11 | 86.94 | 499 | 34 | 499 | 34 |
| *B. breviglume* | Bbrev_135 | 26.70 | 102.30 | 14.78 | 11.33 | 46.80 | 472.09 | 24.96 | 0.74 | 24.22 | 19.88 | 8.49 | 19.88 | 8.49 | 933 | 207 | 7 | 96.83 | 559 | 24 | 559 | 24 |
| *B. breviglume* | Bbrev_136 | 31.00 | 101.10 | 7.63 | 15.13 | 45.71 | 643.20 | 21.91 | -11.18 | 33.09 | 14.92 | -0.16 | 14.92 | -0.81 | 608 | 131 | 2 | 97.95 | 347 | 8 | 347 | 8 |
| *B. breviglume* | Bbrev_137 | 31.20 | 103.10 | 3.41 | 11.00 | 38.16 | 614.99 | 16.43 | -12.39 | 28.82 | 9.26 | -4.54 | 10.62 | -4.54 | 774 | 144 | 5 | 80.07 | 373 | 21 | 369 | 21 |
| *B. breviglume* | Bbrev_138 | 30.80 | 103.50 | 15.49 | 8.19 | 29.82 | 699.34 | 28.53 | 1.06 | 27.47 | 23.00 | 6.40 | 23.71 | 6.40 | 1099 | 246 | 9 | 92.73 | 646 | 38 | 622 | 38 |
| *B. breviglume* | Bbrev_139 | 27.40 | 99.30 | 6.84 | 9.77 | 39.80 | 559.05 | 17.55 | -7.00 | 24.54 | 13.40 | 0.93 | 13.40 | -0.18 | 811 | 141 | 12 | 61.15 | 360 | 52 | 360 | 67 |
| *B. breviglume* | Bbrev_140 | 37.90 | 66.10 | 16.54 | 13.74 | 34.85 | 976.66 | 38.64 | -0.79 | 39.43 | 10.81 | 26.90 | 28.91 | 4.71 | 227 | 54 | 0 | 90.44 | 124 | 1 | 3 | 96 |
| *B. breviglume* | Bbrev_141 | 32.80 | 68.90 | 7.87 | 12.98 | 33.82 | 931.86 | 26.10 | -12.28 | 38.38 | 1.97 | 14.24 | 18.76 | -4.43 | 318 | 57 | 1 | 75.00 | 157 | 19 | 29 | 141 |
| *B. breviglume* | Bbrev_142 | 35.20 | 71.60 | 13.71 | 11.54 | 34.00 | 863.97 | 31.49 | -2.44 | 33.93 | 7.50 | 14.98 | 24.35 | 2.95 | 669 | 131 | 22 | 64.64 | 326 | 76 | 102 | 170 |
| *B. breviglume* | Bbrev_143 | 40.50 | 69.80 | 12.51 | 12.50 | 32.90 | 928.32 | 33.33 | -4.66 | 37.99 | 12.46 | 22.56 | 23.96 | 0.92 | 377 | 62 | 5 | 57.10 | 165 | 23 | 31 | 108 |
| *B. breviglume* | Bbrev_144 | 31.60 | 73.70 | 23.60 | 13.39 | 37.67 | 788.53 | 40.31 | 4.78 | 35.53 | 30.45 | 18.94 | 32.00 | 13.07 | 414 | 117 | 4 | 107.19 | 267 | 19 | 157 | 51 |
| *B. breviglume* | Bbrev_145 | 30.10 | 76.20 | 23.70 | 13.33 | 39.37 | 728.63 | 39.64 | 5.78 | 33.86 | 29.24 | 19.21 | 31.61 | 13.79 | 565 | 165 | 3 | 125.88 | 432 | 29 | 203 | 44 |
| *B. breviglume* | Bbrev_146 | 30.30 | 77.70 | 22.34 | 11.76 | 38.46 | 629.66 | 36.91 | 6.34 | 30.58 | 26.70 | 25.35 | 29.13 | 13.82 | 1787 | 600 | 7 | 138.66 | 1381 | 71 | 691 | 124 |
| *B. breviglume* | Bbrev_147 | 28.30 | 78.30 | 24.79 | 13.16 | 39.70 | 671.32 | 40.40 | 7.24 | 33.16 | 29.28 | 27.97 | 32.10 | 15.64 | 889 | 277 | 3 | 137.82 | 720 | 26 | 304 | 39 |
| *B. breviglume* | Bbrev_148 | 26.80 | 83.00 | 24.85 | 12.58 | 41.29 | 586.94 | 38.66 | 8.20 | 30.46 | 28.66 | 17.49 | 30.65 | 16.50 | 1146 | 348 | 3 | 133.01 | 866 | 28 | 161 | 35 |
| *B. breviglume* | Bbrev_149 | 27.00 | 87.60 | 19.16 | 9.20 | 43.96 | 444.50 | 27.19 | 6.27 | 20.92 | 23.56 | 12.78 | 23.56 | 12.78 | 1562 | 366 | 7 | 97.19 | 944 | 39 | 944 | 39 |
| *B. breviglume* | Bbrev_150 | 23.40 | 91.10 | 25.33 | 9.28 | 45.14 | 357.68 | 32.38 | 11.82 | 20.56 | 27.98 | 19.91 | 28.08 | 19.91 | 2329 | 474 | 5 | 90.87 | 1332 | 32 | 1204 | 32 |
| *B. breviglume* | Bbrev_151 | 37.00 | 99.30 | -3.50 | 12.46 | 34.72 | 847.92 | 12.78 | -23.11 | 35.88 | 6.52 | -13.13 | 6.52 | -14.33 | 348 | 82 | 0 | 105.12 | 227 | 3 | 227 | 4 |
| *B. breviglume* | Bbrev_152 | 35.90 | 102.90 | 7.99 | 12.66 | 32.05 | 949.04 | 26.25 | -13.26 | 39.51 | 17.81 | -4.45 | 19.05 | -4.45 | 427 | 90 | 0 | 91.48 | 241 | 4 | 232 | 4 |
| *B. breviglume* | Bbrev_153 | 34.50 | 106.10 | 10.15 | 9.54 | 29.30 | 847.61 | 26.11 | -6.44 | 32.55 | 19.17 | -0.70 | 20.33 | -0.70 | 628 | 114 | 4 | 79.45 | 328 | 15 | 296 | 15 |
| *B. breviglume* | Bbrev_154 | 30.70 | 109.80 | 13.46 | 7.71 | 26.41 | 783.70 | 28.06 | -1.15 | 29.21 | 20.90 | 3.50 | 22.88 | 3.50 | 1377 | 223 | 25 | 61.61 | 592 | 84 | 588 | 84 |
| *B. breviglume* | Bbrev_155 | 38.70 | 68.80 | 11.97 | 11.33 | 31.68 | 894.79 | 31.23 | -4.54 | 35.77 | 11.35 | 22.05 | 23.14 | 0.85 | 583 | 114 | 2 | 79.30 | 300 | 14 | 23 | 182 |
| *B. breviglume* | Bbrev_156 | 38.60 | 71.90 | 3.49 | 11.75 | 30.55 | 981.33 | 23.68 | -14.80 | 38.48 | 2.47 | 14.79 | 14.79 | -8.85 | 338 | 50 | 5 | 51.91 | 143 | 25 | 25 | 99 |
| *B. breviglume* | Bbrev_157 | 40.60 | 74.10 | -1.08 | 13.32 | 32.33 | 1042.18 | 19.55 | -21.66 | 41.20 | 8.45 | -14.41 | 10.60 | -14.41 | 397 | 68 | 14 | 51.09 | 173 | 48 | 96 | 48 |
| *B. breviglume* | Bbrev_158 | 42.00 | 70.20 | 4.42 | 11.73 | 31.65 | 929.31 | 24.46 | -12.61 | 37.06 | 3.19 | 15.17 | 15.50 | -6.92 | 759 | 113 | 14 | 49.21 | 285 | 60 | 78 | 232 |
| *B. breviglume* | Bbrev_159 | 42.50 | 78.50 | 4.75 | 11.71 | 31.94 | 928.94 | 23.23 | -13.43 | 36.66 | 13.69 | -7.13 | 15.58 | -7.13 | 519 | 72 | 17 | 46.27 | 207 | 58 | 204 | 58 |
| *B. breviglume* | Bbrev_160 | 34.00 | 107.20 | 7.73 | 8.07 | 26.93 | 804.83 | 22.46 | -7.50 | 29.96 | 16.50 | -2.46 | 17.51 | -2.46 | 741 | 132 | 5 | 75.90 | 377 | 20 | 330 | 20 |
| *B. breviglume* | Bbrev_161 | 35.10 | 102.90 | 2.93 | 13.55 | 37.00 | 799.56 | 19.04 | -17.59 | 36.63 | 11.51 | -7.58 | 12.22 | -7.58 | 556 | 110 | 1 | 89.96 | 306 | 6 | 297 | 6 |
| *B. breviglume* | Bbrev_162 | 26.50 | 100.00 | 8.50 | 11.49 | 47.65 | 480.31 | 18.34 | -5.78 | 24.12 | 13.97 | 2.94 | 13.97 | 2.25 | 830 | 146 | 10 | 65.78 | 389 | 48 | 389 | 55 |
| *B. breviglume* | Bbrev_163 | 25.80 | 100.60 | 18.70 | 12.23 | 49.36 | 456.58 | 29.03 | 4.26 | 24.77 | 23.48 | 12.53 | 23.48 | 12.53 | 787 | 141 | 11 | 71.93 | 397 | 46 | 397 | 46 |
| *B. breviglume* | Bbrev_164 | 23.20 | 99.30 | 18.33 | 12.08 | 51.85 | 382.68 | 27.62 | 4.33 | 23.29 | 22.08 | 14.35 | 22.08 | 12.96 | 1432 | 296 | 10 | 87.59 | 792 | 38 | 792 | 44 |
| *B. breviglume* | Bbrev_165 | 34.70 | 103.40 | 3.44 | 13.27 | 36.98 | 785.45 | 19.12 | -16.78 | 35.89 | 11.84 | -6.87 | 12.57 | -6.87 | 588 | 111 | 1 | 86.47 | 314 | 8 | 302 | 8 |
| *B. breviglume* | Bbrev_166 | 27.50 | 90.70 | 9.05 | 10.09 | 42.18 | 508.79 | 19.35 | -4.57 | 23.92 | 14.92 | 3.37 | 14.92 | 2.43 | 1028 | 227 | 5 | 87.96 | 568 | 22 | 568 | 25 |
| *B. breviglume* | Bbrev_167 | 27.30 | 91.50 | 19.90 | 10.29 | 46.29 | 465.15 | 29.31 | 7.08 | 22.23 | 24.84 | 14.47 | 24.84 | 13.49 | 1067 | 235 | 4 | 83.90 | 584 | 28 | 584 | 40 |
| ***B. glaucovirens*** | **Bglaucovirens** | **35.26** | **24.83** | **12.94** | **7.40** | **31.76** | **605.04** | **26.01** | **2.72** | **23.30** | **5.93** | **20.55** | **20.55** | **5.93** | **1034** | **211** | **3** | **83.09** | **544** | **23** | **23** | **544** |
| *B. glaucovirens* | Bgla_1 | 35.26 | 24.83 | 12.94 | 7.40 | 31.76 | 605.04 | 26.01 | 2.72 | 23.30 | 5.93 | 20.54 | 20.54 | 5.93 | 1034 | 211 | 3 | 83.09 | 544 | 23 | 23 | 544 |
| *B. glaucovirens* | Bgla_2 | 37.93 | 22.22 | 8.85 | 9.28 | 35.15 | 650.08 | 24.10 | -2.31 | 26.41 | 2.94 | 17.04 | 17.04 | 1.35 | 931 | 136 | 27 | 52.60 | 379 | 85 | 85 | 371 |
| *B. glaucovirens* | Bgla_3 | 38.46 | 20.65 | 16.51 | 7.95 | 35.12 | 547.52 | 29.33 | 6.70 | 22.63 | 11.82 | 23.31 | 23.40 | 10.42 | 875 | 159 | 8 | 72.30 | 415 | 34 | 63 | 305 |
| ***B. kurilense*** | **Bkurilense** | **45.06** | **147.80** | **2.78** | **5.08** | **18.93** | **776.26** | **16.68** | **-10.13** | **26.81** | **11.15** | **-6.81** | **12.34** | **-6.81** | **1163** | **164** | **54** | **35.28** | **434** | **190** | **364** | **190** |
| *B. kurilense* | Bkur_1 | 45.22 | 147.87 | 3.53 | 5.26 | 19.94 | 758.73 | 17.20 | -9.20 | 26.40 | 7.87 | -5.84 | 12.85 | -5.84 | 1105 | 155 | 52 | 34.95 | 412 | 180 | 347 | 180 |
| *B. kurilense* | Bkur_2 | 45.00 | 147.50 | 3.30 | 5.08 | 19.21 | 757.50 | 17.00 | -9.42 | 26.42 | 11.68 | -6.09 | 12.66 | -6.09 | 1114 | 158 | 51 | 34.20 | 412 | 181 | 357 | 181 |
| *B. kurilense* | Bkur_3 | 43.78 | 146.69 | 4.92 | 5.06 | 19.64 | 748.70 | 18.52 | -7.25 | 25.78 | 13.08 | -4.35 | 14.24 | -4.35 | 1148 | 166 | 47 | 34.78 | 422 | 177 | 378 | 177 |
| *B. kurilense* | Bkur_4 | 44.18 | 145.87 | 3.56 | 5.35 | 18.89 | 809.51 | 17.62 | -10.72 | 28.33 | 12.23 | -5.93 | 13.59 | -6.49 | 1090 | 161 | 47 | 34.51 | 405 | 178 | 374 | 179 |
| *B. kurilense* | Bkur_5 | 43.78 | 145.47 | 5.92 | 5.91 | 20.40 | 830.83 | 20.20 | -8.77 | 28.97 | 14.64 | -4.14 | 16.04 | -4.40 | 1036 | 161 | 44 | 36.73 | 392 | 165 | 368 | 171 |
| *B. kurilense* | Bkur_6 | 44.30 | 146.15 | 4.14 | 5.49 | 19.63 | 803.46 | 18.38 | -9.62 | 27.99 | 12.79 | -5.74 | 14.09 | -5.74 | 1066 | 157 | 45 | 34.85 | 398 | 174 | 357 | 174 |
| *B. kurilense* | Bkur_7 | 44.99 | 147.79 | 2.96 | 5.04 | 19.29 | 752.50 | 16.62 | -9.49 | 26.11 | 11.17 | -6.34 | 12.29 | -6.34 | 1174 | 166 | 53 | 35.61 | 439 | 188 | 369 | 188 |
| *B. kurilense* | Bkur_8 | 43.93 | 145.65 | 5.36 | 5.33 | 19.22 | 792.64 | 19.22 | -8.54 | 27.76 | 13.98 | -3.98 | 15.18 | -4.45 | 1035 | 158 | 43 | 36.42 | 391 | 164 | 363 | 168 |
| *B. kurilense* | Bkur_9 | 44.39 | 146.44 | 4.59 | 5.00 | 18.76 | 767.35 | 18.20 | -8.45 | 26.65 | 13.12 | -4.94 | 14.14 | -4.94 | 1059 | 154 | 44 | 34.55 | 392 | 168 | 352 | 168 |
| *B. kurilense* | Bkur_10 | 45.40 | 147.93 | 1.39 | 4.76 | 18.73 | 729.69 | 14.78 | -10.66 | 25.44 | 9.51 | -7.63 | 10.56 | -7.63 | 1183 | 163 | 56 | 33.84 | 436 | 193 | 376 | 193 |
| *B. kurilense* | Bkur_11 | 44.17 | 145.95 | 4.19 | 5.48 | 19.28 | 809.39 | 18.43 | -10.01 | 28.44 | 12.93 | -5.81 | 14.21 | -5.81 | 1064 | 158 | 45 | 35.04 | 398 | 173 | 362 | 173 |
| *B. kurilense* | Bkur_12 | 43.80 | 146.74 | 5.01 | 5.04 | 19.59 | 747.30 | 18.60 | -7.15 | 25.75 | 13.16 | -4.24 | 14.32 | -4.24 | 1130 | 164 | 46 | 35.32 | 418 | 174 | 372 | 174 |
| *B. kurilense* | Bkur_13 | 45.08 | 147.88 | 1.74 | 4.94 | 18.28 | 784.68 | 15.86 | -11.15 | 27.00 | 10.19 | -7.93 | 11.45 | -7.93 | 1213 | 170 | 56 | 34.99 | 451 | 196 | 382 | 196 |
| *B. kurilense* | Bkur_14 | 46.81 | 142.06 | 2.86 | 6.11 | 18.86 | 976.75 | 19.49 | -12.92 | 32.41 | 12.02 | -4.84 | 14.82 | -9.09 | 844 | 107 | 38 | 33.44 | 304 | 127 | 302 | 165 |
| *B. kurilense* | Bkur_15 | 43.85 | 145.52 | 5.06 | 5.36 | 18.76 | 826.87 | 19.34 | -9.22 | 28.55 | 13.76 | -4.75 | 15.22 | -5.20 | 1061 | 163 | 44 | 36.64 | 401 | 166 | 377 | 172 |
| *B. kurilense* | Bkur_16 | 44.39 | 146.23 | 2.66 | 4.99 | 17.61 | 833.64 | 17.16 | -11.16 | 28.32 | 11.39 | -7.62 | 12.95 | -7.62 | 1091 | 157 | 47 | 33.74 | 403 | 179 | 367 | 179 |
| *B. kurilense* | Bkur_17 | 47.80 | 142.10 | 2.74 | 6.78 | 19.60 | 1005.37 | 19.66 | -14.90 | 34.56 | 12.25 | -4.85 | 15.01 | -9.85 | 821 | 97 | 45 | 28.58 | 285 | 138 | 274 | 175 |
| *B. kurilense* | Bkur_18 | 45.20 | 147.90 | 3.17 | 5.17 | 19.40 | 771.98 | 17.04 | -9.60 | 26.64 | 7.52 | -6.34 | 12.66 | -6.34 | 1138 | 159 | 54 | 34.48 | 422 | 188 | 355 | 188 |
| *B. kurilense* | Bkur_19 | 43.40 | 142.60 | 0.92 | 8.45 | 22.71 | 1045.38 | 19.35 | -17.86 | 37.21 | 9.92 | -11.62 | 13.52 | -11.74 | 1327 | 174 | 59 | 33.28 | 484 | 213 | 474 | 219 |
| *B. kurilense* | Bkur_20 | 43.00 | 142.00 | 5.91 | 8.56 | 24.27 | 972.18 | 23.34 | -11.93 | 35.28 | 17.68 | -5.62 | 17.68 | -5.99 | 1156 | 158 | 55 | 34.14 | 426 | 185 | 426 | 207 |
| *B. kurilense* | Bkur_21 | 43.20 | 141.30 | 8.19 | 7.98 | 23.53 | 957.68 | 25.52 | -8.39 | 33.91 | 16.79 | 11.37 | 19.87 | -3.48 | 1216 | 142 | 63 | 29.08 | 399 | 196 | 372 | 328 |
| *B. kurilense* | Bkur_22 | 42.30 | 143.20 | 5.48 | 8.27 | 23.28 | 975.41 | 22.08 | -13.44 | 35.52 | 14.40 | -6.91 | 17.06 | -6.91 | 1124 | 169 | 41 | 38.31 | 415 | 151 | 414 | 151 |
| *B. kurilense* | Bkur_23 | 43.20 | 145.00 | 6.72 | 6.41 | 21.06 | 878.15 | 21.51 | -8.94 | 30.45 | 15.49 | -4.27 | 17.29 | -4.27 | 1070 | 165 | 39 | 39.14 | 404 | 145 | 388 | 145 |
| *B. kurilense* | Bkur_24 | 45.30 | 141.00 | 5.91 | 5.19 | 17.69 | 865.75 | 21.12 | -8.23 | 29.34 | 14.84 | -1.35 | 16.91 | -4.40 | 1109 | 128 | 56 | 29.48 | 373 | 178 | 346 | 266 |
| *B. kurilense* | Bkur_25 | 43.30 | 140.40 | 7.18 | 6.71 | 21.48 | 906.18 | 23.74 | -7.51 | 31.25 | -1.16 | 9.67 | 18.50 | -3.67 | 1268 | 155 | 60 | 29.77 | 413 | 195 | 367 | 394 |
| *B. kurilense* | Bkur_26 | 44.40 | 146.10 | 1.65 | 5.31 | 18.26 | 849.60 | 16.43 | -12.68 | 29.10 | 10.49 | -8.46 | 12.15 | -8.81 | 1186 | 171 | 51 | 34.33 | 441 | 190 | 398 | 194 |
| *B. kurilense* | Bkur_27 | 43.80 | 146.70 | 4.77 | 5.02 | 19.41 | 749.00 | 18.36 | -7.53 | 25.89 | 12.92 | -4.52 | 14.11 | -4.52 | 1141 | 165 | 46 | 35.10 | 420 | 175 | 376 | 175 |
| *B. kurilense* | Bkur_28 | 44.00 | 145.70 | 4.62 | 5.36 | 18.96 | 813.09 | 18.73 | -9.55 | 28.28 | 13.28 | -5.03 | 14.66 | -5.41 | 1058 | 160 | 44 | 36.01 | 398 | 169 | 369 | 171 |
| ***B. miserum*** | **Bmis66-3** | **35.40** | **139.05** | **13.23** | **8.91** | **29.53** | **755.69** | **28.12** | **-2.04** | **30.16** | **20.08** | **4.12** | **22.58** | **4.12** | **1590** | **220** | **45** | **48.07** | **606** | **155** | **603** | **155** |
| ***B. miserum*** | **Bmis66-4** | **35.40** | **139.05** | **13.23** | **8.91** | **29.53** | **755.69** | **28.12** | **-2.04** | **30.16** | **20.08** | **4.12** | **22.58** | **4.12** | **1590** | **220** | **45** | **48.07** | **606** | **155** | **603** | **155** |
| ***B. miserum*** | **Bmis67-1** | **38.32** | **140.60** | **9.83** | **7.50** | **24.37** | **869.37** | **26.30** | **-4.50** | **30.80** | **20.54** | **-0.41** | **20.54** | **-0.47** | **1311** | **175** | **68** | **37.17** | **509** | **211** | **509** | **216** |
| ***B. miserum*** | **Bmis67-2** | **38.32** | **140.60** | **9.83** | **7.50** | **24.37** | **869.37** | **26.30** | **-4.50** | **30.80** | **20.54** | **-0.41** | **20.54** | **-0.47** | **1311** | **175** | **68** | **37.17** | **509** | **211** | **509** | **216** |
| *B. miserum* | Bmis_1 | 38.26 | 140.84 | 12.35 | 7.93 | 25.93 | 834.39 | 28.18 | -2.40 | 30.58 | 22.68 | 2.36 | 22.68 | 2.36 | 1225 | 192 | 39 | 49.48 | 515 | 133 | 515 | 133 |
| *B. miserum* | Bmis_2 | 36.90 | 139.30 | 5.19 | 8.75 | 26.02 | 917.64 | 22.22 | -11.40 | 33.62 | 16.40 | -5.79 | 16.40 | -5.99 | 1966 | 334 | 64 | 54.60 | 869 | 212 | 869 | 247 |
| *B. miserum* | Bmis_4 | 35.20 | 134.10 | 11.55 | 8.58 | 27.27 | 854.97 | 28.04 | -3.42 | 31.46 | 22.11 | 3.82 | 22.11 | 1.31 | 1722 | 248 | 72 | 42.31 | 631 | 248 | 631 | 248 |
| *B. miserum* | Bmis_6 | 39.30 | 140.60 | 9.83 | 8.01 | 23.65 | 961.52 | 27.92 | -5.95 | 33.87 | 21.71 | 7.15 | 21.71 | -1.59 | 1588 | 178 | 93 | 20.62 | 487 | 293 | 487 | 416 |
| *B. miserum* | Bmis_7 | 40.20 | 140.90 | 7.58 | 7.82 | 23.52 | 938.72 | 25.09 | -8.16 | 33.25 | 19.10 | -0.10 | 19.10 | -3.62 | 1455 | 174 | 86 | 26.50 | 516 | 278 | 516 | 300 |
| *B. miserum* | Bmis_8 | 40.10 | 140.20 | 10.58 | 6.90 | 22.30 | 888.49 | 27.59 | -3.34 | 30.93 | 21.74 | 3.23 | 21.74 | 0.22 | 1513 | 171 | 89 | 22.28 | 488 | 284 | 488 | 355 |
| *B. miserum* | Bmis_9 | 35.20 | 135.80 | 10.97 | 8.21 | 26.50 | 851.46 | 27.27 | -3.70 | 30.97 | 21.13 | 3.34 | 21.50 | 0.77 | 1906 | 252 | 85 | 38.62 | 691 | 283 | 677 | 283 |
| *B. miserum* | Bmis_10 | 32.10 | 130.80 | 12.93 | 7.77 | 26.70 | 782.16 | 27.01 | -2.10 | 29.12 | 21.98 | 5.39 | 22.39 | 3.13 | 2840 | 515 | 72 | 64.33 | 1345 | 255 | 1089 | 280 |
| *B. miserum* | Bmis_11 | 34.50 | 136.80 | 15.25 | 7.29 | 26.01 | 763.79 | 29.49 | 1.45 | 28.04 | 22.31 | 6.05 | 24.75 | 6.05 | 2122 | 315 | 61 | 46.13 | 784 | 215 | 774 | 215 |
| *B. miserum* | Bmis_12 | 35.80 | 140.70 | 14.57 | 7.86 | 26.77 | 758.29 | 28.54 | -0.84 | 29.38 | 21.80 | 5.11 | 23.87 | 5.11 | 1458 | 205 | 54 | 38.75 | 520 | 183 | 467 | 183 |
| *B. miserum* | Bmis_14 | 36.20 | 137.50 | 5.35 | 9.42 | 27.23 | 929.29 | 22.80 | -11.79 | 34.58 | 16.53 | -6.07 | 16.53 | -6.07 | 1896 | 264 | 68 | 48.41 | 772 | 213 | 772 | 213 |
| *B. miserum* | Bmis_16 | 35.50 | 140.20 | 15.05 | 7.87 | 27.06 | 755.88 | 29.31 | 0.23 | 29.08 | 22.07 | 5.83 | 24.38 | 5.83 | 1482 | 197 | 54 | 37.91 | 522 | 188 | 448 | 188 |
| *B. miserum* | Bmis_17 | 36.50 | 139.20 | 11.67 | 9.83 | 30.42 | 826.93 | 28.09 | -4.23 | 32.32 | 21.86 | 4.24 | 21.86 | 1.84 | 1449 | 210 | 50 | 48.97 | 603 | 180 | 603 | 183 |
| *B. miserum* | Bmis_18 | 36.30 | 138.70 | 10.64 | 9.79 | 29.88 | 847.56 | 27.47 | -5.30 | 32.77 | 21.07 | 0.54 | 21.07 | 0.54 | 1207 | 178 | 31 | 54.21 | 505 | 121 | 505 | 121 |
| *B. miserum* | Bmis_19 | 36.20 | 140.10 | 13.40 | 8.59 | 27.77 | 793.63 | 28.48 | -2.44 | 30.92 | 20.65 | 3.70 | 23.14 | 3.70 | 1336 | 187 | 34 | 46.56 | 499 | 128 | 474 | 128 |
| *B. miserum* | Bmis_20 | 36.50 | 136.90 | 11.35 | 7.58 | 23.90 | 892.36 | 28.15 | -3.56 | 31.71 | 22.26 | 4.00 | 22.26 | 0.60 | 2075 | 254 | 124 | 24.20 | 679 | 407 | 679 | 439 |
| *B. miserum* | Bmis_21 | 35.40 | 137.50 | 9.59 | 9.45 | 29.03 | 845.55 | 25.93 | -6.62 | 32.56 | 19.91 | -0.75 | 19.91 | -0.75 | 2078 | 296 | 60 | 51.69 | 843 | 198 | 843 | 198 |
| *B. miserum* | Bmis_22 | 33.80 | 135.70 | 13.49 | 7.32 | 25.89 | 772.54 | 27.46 | -0.79 | 28.25 | 22.46 | 3.99 | 22.96 | 3.99 | 2166 | 323 | 56 | 48.57 | 803 | 204 | 773 | 204 |
| *B. miserum* | Bmis_23 | 37.70 | 139.50 | 10.97 | 7.82 | 23.88 | 921.48 | 28.49 | -4.25 | 32.74 | 22.27 | 8.47 | 22.27 | 0.00 | 1660 | 179 | 95 | 20.65 | 485 | 300 | 485 | 392 |
| *B. miserum* | Bmis_24 | 34.70 | 134.50 | 14.95 | 7.21 | 24.59 | 790.17 | 30.12 | 0.80 | 29.32 | 20.98 | 5.75 | 24.92 | 5.46 | 1326 | 211 | 31 | 53.71 | 498 | 122 | 467 | 179 |
| *B. miserum* | Bmis_25 | 34.90 | 135.30 | 13.78 | 8.58 | 27.62 | 838.22 | 30.06 | -0.99 | 31.06 | 20.72 | 3.76 | 24.18 | 3.76 | 1444 | 216 | 45 | 45.39 | 540 | 167 | 485 | 167 |
| *B. miserum* | Bmis_26 | 35.60 | 134.40 | 13.77 | 8.28 | 27.22 | 814.25 | 29.99 | -0.41 | 30.41 | 23.95 | 11.34 | 23.95 | 4.16 | 1879 | 233 | 117 | 21.57 | 545 | 369 | 545 | 472 |
| *B. miserum* | Bmis_27 | 34.20 | 134.80 | 16.31 | 6.08 | 22.62 | 737.12 | 30.18 | 3.30 | 26.88 | 25.62 | 7.76 | 25.62 | 7.38 | 1525 | 252 | 41 | 51.46 | 541 | 152 | 541 | 204 |
| *B. miserum* | Bmis_28 | 35.10 | 134.80 | 11.89 | 8.43 | 27.09 | 849.29 | 28.22 | -2.90 | 31.12 | 19.00 | 1.76 | 22.39 | 1.76 | 1679 | 225 | 73 | 37.77 | 582 | 253 | 580 | 253 |
| *B. miserum* | Bmis_29 | 40.70 | 140.90 | 5.76 | 7.33 | 22.68 | 922.44 | 22.78 | -9.54 | 32.32 | 17.14 | -5.07 | 17.14 | -5.24 | 1451 | 183 | 84 | 27.66 | 519 | 270 | 519 | 285 |
| *B. miserum* | Bmis_30 | 37.70 | 140.10 | 7.42 | 8.31 | 25.26 | 909.03 | 24.61 | -8.31 | 32.92 | 18.53 | -3.33 | 18.53 | -3.58 | 1561 | 224 | 74 | 41.22 | 624 | 226 | 624 | 238 |
| *B. miserum* | Bmis_32 | 37.10 | 140.20 | 11.08 | 9.16 | 28.61 | 845.48 | 27.60 | -4.42 | 32.02 | 21.46 | 0.97 | 21.46 | 0.97 | 1300 | 192 | 37 | 52.69 | 542 | 125 | 542 | 125 |
| *B. miserum* | Bmis_33 | 35.60 | 139.30 | 14.36 | 9.14 | 29.11 | 789.40 | 29.83 | -1.56 | 31.39 | 21.42 | 4.73 | 24.06 | 4.73 | 1511 | 207 | 45 | 48.39 | 578 | 147 | 560 | 147 |
| *B. miserum* | Bmis_34 | 36.80 | 139.70 | 10.54 | 8.59 | 27.40 | 836.22 | 26.86 | -4.51 | 31.36 | 20.88 | 0.65 | 20.88 | 0.65 | 1496 | 213 | 54 | 47.68 | 610 | 178 | 610 | 178 |
| *B. miserum* | Bmis_35 | 32.80 | 131.10 | 12.60 | 8.07 | 27.30 | 802.37 | 27.18 | -2.37 | 29.55 | 21.94 | 4.94 | 22.32 | 2.60 | 2478 | 468 | 63 | 65.33 | 1192 | 221 | 989 | 241 |
| *B. miserum* | Bmis_36 | 34.80 | 133.30 | 12.84 | 8.23 | 26.56 | 853.46 | 28.98 | -2.02 | 30.99 | 23.15 | 4.99 | 23.42 | 2.59 | 1584 | 249 | 55 | 51.06 | 636 | 184 | 597 | 189 |
| *B. miserum* | Bmis_37 | 37.40 | 140.70 | 9.43 | 8.28 | 26.67 | 839.14 | 25.27 | -5.77 | 31.04 | 19.75 | -0.64 | 19.75 | -0.65 | 1332 | 198 | 34 | 52.31 | 548 | 121 | 548 | 158 |
| *B. miserum* | Bmis_38 | 35.20 | 139.60 | 15.59 | 7.86 | 27.56 | 730.44 | 30.02 | 1.50 | 28.52 | 22.40 | 6.81 | 24.70 | 6.81 | 1794 | 226 | 62 | 36.52 | 594 | 220 | 542 | 220 |
| *B. miserum* | Bmis_39 | 34.70 | 131.90 | 14.68 | 7.92 | 27.36 | 785.17 | 29.89 | 0.96 | 28.93 | 21.11 | 5.39 | 24.54 | 5.39 | 1804 | 272 | 81 | 41.69 | 687 | 270 | 619 | 270 |
| *B. miserum* | Bmis_40 | 35.20 | 138.50 | 13.83 | 8.58 | 28.62 | 765.27 | 28.51 | -1.48 | 29.99 | 22.70 | 4.45 | 23.24 | 4.45 | 1760 | 238 | 49 | 44.86 | 652 | 180 | 637 | 180 |
| *B. miserum* | Bmis_41 | 35.80 | 138.80 | 8.17 | 9.83 | 30.53 | 813.86 | 24.28 | -7.91 | 32.18 | 18.16 | -1.61 | 18.16 | -1.62 | 1629 | 266 | 30 | 61.06 | 718 | 114 | 718 | 173 |
| *B. miserum* | Bmis_42 | 33.80 | 133.10 | 12.33 | 6.91 | 23.81 | 815.53 | 26.74 | -2.27 | 29.01 | 19.14 | 2.25 | 22.26 | 2.25 | 2081 | 331 | 59 | 54.28 | 869 | 205 | 754 | 205 |
| *B. miserum* | Bmis_43 | 36.70 | 138.00 | 8.92 | 9.60 | 27.56 | 928.74 | 26.66 | -8.18 | 34.84 | 20.09 | -2.42 | 20.09 | -2.42 | 1319 | 183 | 60 | 39.77 | 495 | 196 | 495 | 196 |
| *B. miserum* | Bmis_44 | 35.40 | 136.40 | 12.50 | 8.14 | 26.35 | 848.22 | 28.82 | -2.06 | 30.88 | 22.60 | 4.87 | 23.02 | 2.34 | 2028 | 267 | 102 | 36.07 | 715 | 324 | 701 | 327 |
| *B. miserum* | Bmis_45 | 36.10 | 136.10 | 12.80 | 7.34 | 24.38 | 835.53 | 28.93 | -1.16 | 30.10 | 5.54 | 10.36 | 23.22 | 2.90 | 2261 | 252 | 144 | 21.90 | 681 | 441 | 599 | 574 |
| *B. miserum* | Bmis_46 | 35.90 | 138.20 | 6.76 | 9.80 | 29.41 | 860.26 | 23.44 | -9.86 | 33.31 | 17.19 | -3.69 | 17.19 | -3.69 | 1693 | 251 | 36 | 57.67 | 730 | 134 | 730 | 134 |
| *B. miserum* | Bmis_47 | 35.40 | 133.50 | 10.85 | 8.11 | 26.19 | 862.99 | 27.04 | -3.94 | 30.98 | 21.16 | 3.14 | 21.45 | 0.44 | 1811 | 261 | 94 | 36.99 | 653 | 301 | 653 | 308 |
| *B. miserum* | Bmis_51 | 37.00 | 138.60 | 10.61 | 8.83 | 25.24 | 960.59 | 28.55 | -6.44 | 35.00 | -0.88 | 7.98 | 22.23 | -0.94 | 1804 | 202 | 97 | 23.40 | 539 | 309 | 521 | 463 |
| *B. miserum* | Bmis_53 | 33.80 | 133.80 | 11.87 | 7.03 | 24.12 | 815.30 | 26.23 | -2.91 | 29.14 | 18.65 | 1.76 | 21.79 | 1.76 | 2126 | 333 | 56 | 53.22 | 863 | 203 | 767 | 203 |
| *B. miserum* | Bmis_54 | 33.20 | 133.20 | 15.42 | 7.44 | 26.61 | 750.27 | 28.88 | 0.92 | 27.96 | 21.47 | 6.09 | 24.62 | 6.09 | 2309 | 347 | 58 | 55.22 | 960 | 228 | 763 | 228 |
| ***B. spryginii*** | **Bspryginii** | **45.08** | **38.91** | **11.72** | **10.00** | **30.53** | **865.46** | **29.25** | **-3.49** | **32.74** | **2.74** | **17.54** | **22.40** | **1.12** | **727** | **80** | **46** | **17.58** | **218** | **155** | **185** | **198** |
| *B. spryginii* | Bspry_1 | 45.11 | 38.82 | 11.57 | 10.34 | 31.16 | 873.46 | 29.49 | -3.70 | 33.19 | 2.53 | 5.97 | 22.34 | 0.89 | 717 | 80 | 46 | 17.46 | 216 | 153 | 183 | 197 |
| ***B. sylvaticum*** | **Bsyl29H** | **38.99** | **21.69** | **13.31** | **10.51** | **36.15** | **683.81** | **29.60** | **0.53** | **29.07** | **6.70** | **21.86** | **21.86** | **5.20** | **800** | **118** | **21** | **52.47** | **321** | **68** | **68** | **297** |
| ***B. sylvaticum*** | **Bsyl30H** | **55.49** | **11.78** | **8.55** | **4.76** | **21.49** | **665.56** | **20.77** | **-1.38** | **22.15** | **16.20** | **3.15** | **16.99** | **0.74** | **615** | **65** | **33** | **20.29** | **181** | **110** | **176** | **142** |
| ***B. sylvaticum*** | **Bsyl31H** | **48.22** | **9.41** | **6.89** | **8.30** | **32.19** | **666.36** | **20.96** | **-4.83** | **25.79** | **15.10** | **-0.16** | **15.10** | **-1.21** | **913** | **114** | **56** | **25.35** | **310** | **172** | **310** | **179** |
| ***B. sylvaticum*** | **Bsyl32H** | **54.54** | **36.12** | **5.20** | **8.33** | **24.49** | **973.04** | **23.10** | **-10.91** | **34.01** | **16.98** | **-5.57** | **16.98** | **-6.84** | **618** | **82** | **31** | **31.57** | **224** | **100** | **224** | **115** |
| ***B. sylvaticum*** | **Bsyl35H** | **38.02** | **21.94** | **11.66** | **10.59** | **37.96** | **650.31** | **27.68** | **-0.22** | **27.90** | **5.63** | **19.90** | **19.90** | **4.17** | **841** | **131** | **20** | **58.88** | **367** | **64** | **64** | **345** |
| ***B. sylvaticum*** | **Bsyl36H** | **33.52** | **47.77** | **17.36** | **16.01** | **39.94** | **894.71** | **39.79** | **-0.30** | **40.09** | **7.56** | **27.97** | **28.52** | **6.55** | **446** | **85** | **0** | **86.18** | **222** | **1** | **3** | **211** |
| ***B. sylvaticum*** | **Bsyl37H** | **35.18** | **50.13** | **9.55** | **13.09** | **32.93** | **969.82** | **30.80** | **-8.97** | **39.77** | **2.96** | **20.79** | **21.37** | **-2.69** | **235** | **47** | **0** | **77.97** | **116** | **3** | **8** | **85** |
| ***B. sylvaticum*** | **Bsyl38H** | **36.83** | **36.20** | **19.66** | **9.15** | **34.87** | **663.28** | **32.87** | **6.64** | **26.23** | **11.45** | **27.29** | **27.70** | **11.45** | **833** | **120** | **12** | **54.63** | **330** | **54** | **61** | **330** |
| ***B. sylvaticum*** | **Bsyl39H** | **41.28** | **36.29** | **13.34** | **7.50** | **31.33** | **623.26** | **26.13** | **2.19** | **23.94** | **11.03** | **21.21** | **21.21** | **6.21** | **675** | **82** | **31** | **27.62** | **233** | **113** | **113** | **164** |
| ***B. sylvaticum*** | **Bsyl41H** | **34.48** | **69.30** | **12.56** | **14.00** | **34.34** | **977.86** | **32.84** | **-7.94** | **40.78** | **6.96** | **19.41** | **24.27** | **-0.18** | **365** | **82** | **5** | **89.75** | **219** | **21** | **23** | **132** |
| ***B. sylvaticum*** | **Bsyl466-13** | **42.25** | **-0.24** | **12.61** | **10.45** | **36.41** | **653.50** | **29.37** | **0.68** | **28.69** | **13.38** | **21.02** | **21.02** | **5.15** | **628** | **71** | **26** | **23.84** | **186** | **126** | **126** | **149** |
| ***B. sylvaticum*** | **Bsyl466-2** | **42.25** | **-0.24** | **12.61** | **10.45** | **36.41** | **653.50** | **29.37** | **0.68** | **28.69** | **13.38** | **21.02** | **21.02** | **5.15** | **628** | **71** | **26** | **23.84** | **186** | **126** | **126** | **149** |
| ***B. sylvaticum*** | **Bsyl466-7b** | **42.25** | **-0.24** | **12.61** | **10.45** | **36.41** | **653.50** | **29.37** | **0.68** | **28.69** | **13.38** | **21.02** | **21.02** | **5.15** | **628** | **71** | **26** | **23.84** | **186** | **126** | **126** | **149** |
| ***B. sylvaticum*** | **Bsyl467-10** | **42.06** | **-0.12** | **14.61** | **11.02** | **36.09** | **698.11** | **32.27** | **1.73** | **30.54** | **15.44** | **7.78** | **23.51** | **6.44** | **519** | **61** | **20** | **26.62** | **159** | **102** | **105** | **113** |
| ***B. sylvaticum*** | **Bsyl467-2** | **42.06** | **-0.12** | **14.61** | **11.02** | **36.09** | **698.11** | **32.27** | **1.73** | **30.54** | **15.44** | **7.78** | **23.51** | **6.44** | **519** | **61** | **20** | **26.62** | **159** | **102** | **105** | **113** |
| ***B. sylvaticum*** | **Bsyl467-7** | **42.06** | **-0.12** | **14.61** | **11.02** | **36.09** | **698.11** | **32.27** | **1.73** | **30.54** | **15.44** | **7.78** | **23.51** | **6.44** | **519** | **61** | **20** | **26.62** | **159** | **102** | **105** | **113** |
| ***B. sylvaticum*** | **Bsyl467-9b** | **42.06** | **-0.12** | **14.61** | **11.02** | **36.09** | **698.11** | **32.27** | **1.73** | **30.54** | **15.44** | **7.78** | **23.51** | **6.44** | **519** | **61** | **20** | **26.62** | **159** | **102** | **105** | **113** |
| ***B. sylvaticum*** | **Bsyl470-10** | **43.32** | **-1.97** | **14.32** | **7.70** | **40.28** | **432.00** | **24.39** | **5.28** | **19.11** | **12.46** | **19.54** | **19.87** | **9.40** | **1271** | **148** | **66** | **22.65** | **399** | **229** | **248** | **347** |
| ***B. sylvaticum*** | **Bsyl470-4** | **43.32** | **-1.97** | **14.32** | **7.70** | **40.28** | **432.00** | **24.39** | **5.28** | **19.11** | **12.46** | **19.54** | **19.87** | **9.40** | **1271** | **148** | **66** | **22.65** | **399** | **229** | **248** | **347** |
| ***B. sylvaticum*** | **Bsyl470-8b** | **43.32** | **-1.97** | **14.32** | **7.70** | **40.28** | **432.00** | **24.39** | **5.28** | **19.11** | **12.46** | **19.54** | **19.87** | **9.40** | **1271** | **148** | **66** | **22.65** | **399** | **229** | **248** | **347** |
| ***B. sylvaticum*** | **Bsyl470-9** | **43.32** | **-1.97** | **14.32** | **7.70** | **40.28** | **432.00** | **24.39** | **5.28** | **19.11** | **12.46** | **19.54** | **19.87** | **9.40** | **1271** | **148** | **66** | **22.65** | **399** | **229** | **248** | **347** |
| ***B. sylvaticum*** | **Bsyl476-10** | **43.03** | **-0.62** | **11.84** | **8.97** | **39.88** | **495.10** | **24.74** | **2.23** | **22.50** | **7.04** | **18.37** | **18.37** | **6.42** | **1004** | **107** | **56** | **19.27** | **301** | **186** | **186** | **281** |
| ***B. sylvaticum*** | **Bsyl476-11** | **43.03** | **-0.62** | **11.84** | **8.97** | **39.88** | **495.10** | **24.74** | **2.23** | **22.50** | **7.04** | **18.37** | **18.37** | **6.42** | **1004** | **107** | **56** | **19.27** | **301** | **186** | **186** | **281** |
| ***B. sylvaticum*** | **Bsyl476-14** | **43.03** | **-0.62** | **11.84** | **8.97** | **39.88** | **495.10** | **24.74** | **2.23** | **22.50** | **7.04** | **18.37** | **18.37** | **6.42** | **1004** | **107** | **56** | **19.27** | **301** | **186** | **186** | **281** |
| ***B. sylvaticum*** | **Bsyl476-9** | **43.03** | **-0.62** | **11.84** | **8.97** | **39.88** | **495.10** | **24.74** | **2.23** | **22.50** | **7.04** | **18.37** | **18.37** | **6.42** | **1004** | **107** | **56** | **19.27** | **301** | **186** | **186** | **281** |
| ***B. sylvaticum*** | **Bsyl477-10** | **42.83** | **0.72** | **8.40** | **9.70** | **37.36** | **585.01** | **23.15** | **-2.80** | **25.95** | **5.66** | **15.97** | **15.97** | **1.77** | **1159** | **118** | **69** | **14.11** | **325** | **248** | **248** | **283** |
| ***B. sylvaticum*** | **Bsyl477-11** | **42.83** | **0.72** | **8.40** | **9.70** | **37.36** | **585.01** | **23.15** | **-2.80** | **25.95** | **5.66** | **15.97** | **15.97** | **1.77** | **1159** | **118** | **69** | **14.11** | **325** | **248** | **248** | **283** |
| ***B. sylvaticum*** | **Bsyl477-1** | **42.83** | **0.72** | **8.40** | **9.70** | **37.36** | **585.01** | **23.15** | **-2.80** | **25.95** | **5.66** | **15.97** | **15.97** | **1.77** | **1159** | **118** | **69** | **14.11** | **325** | **248** | **248** | **283** |
| ***B. sylvaticum*** | **Bsyl477-3** | **42.83** | **0.72** | **8.40** | **9.70** | **37.36** | **585.01** | **23.15** | **-2.80** | **25.95** | **5.66** | **15.97** | **15.97** | **1.77** | **1159** | **118** | **69** | **14.11** | **325** | **248** | **248** | **283** |
| ***B. sylvaticum*** | **Bsyl500-2** | **43.34** | **5.74** | **12.53** | **8.95** | **36.12** | **580.11** | **26.59** | **1.80** | **24.78** | **9.57** | **19.98** | **19.98** | **5.98** | **751** | **108** | **15** | **37.31** | **262** | **97** | **97** | **214** |
| ***B. sylvaticum*** | **Bsyl500-3** | **43.34** | **5.74** | **12.53** | **8.95** | **36.12** | **580.11** | **26.59** | **1.80** | **24.78** | **9.57** | **19.98** | **19.98** | **5.98** | **751** | **108** | **15** | **37.31** | **262** | **97** | **97** | **214** |
| ***B. sylvaticum*** | **Bsyl500-6** | **43.34** | **5.74** | **12.53** | **8.95** | **36.12** | **580.11** | **26.59** | **1.80** | **24.78** | **9.57** | **19.98** | **19.98** | **5.98** | **751** | **108** | **15** | **37.31** | **262** | **97** | **97** | **214** |
| ***B. sylvaticum*** | **Bsyl501-1** | **43.66** | **7.04** | **14.24** | **7.87** | **33.90** | **559.31** | **26.70** | **3.48** | **23.22** | **11.42** | **21.34** | **21.39** | **7.90** | **847** | **133** | **17** | **43.48** | **325** | **97** | **129** | **248** |
| ***B. sylvaticum*** | **Bsyl501-5** | **43.66** | **7.04** | **14.24** | **7.87** | **33.90** | **559.31** | **26.70** | **3.48** | **23.22** | **11.42** | **21.34** | **21.39** | **7.90** | **847** | **133** | **17** | **43.48** | **325** | **97** | **129** | **248** |
| ***B. sylvaticum*** | **Bsyl501-6** | **43.66** | **7.04** | **14.24** | **7.87** | **33.90** | **559.31** | **26.70** | **3.48** | **23.22** | **11.42** | **21.34** | **21.39** | **7.90** | **847** | **133** | **17** | **43.48** | **325** | **97** | **129** | **248** |
| ***B. sylvaticum*** | **Bsyl501-7** | **43.66** | **7.04** | **14.24** | **7.87** | **33.90** | **559.31** | **26.70** | **3.48** | **23.22** | **11.42** | **21.34** | **21.39** | **7.90** | **847** | **133** | **17** | **43.48** | **325** | **97** | **129** | **248** |
| ***B. sylvaticum*** | **Bsyl502-2** | **43.37** | **6.37** | **14.48** | **9.23** | **37.07** | **569.86** | **28.28** | **3.40** | **24.88** | **11.59** | **21.77** | **21.77** | **7.99** | **753** | **106** | **12** | **39.60** | **275** | **91** | **91** | **230** |
| ***B. sylvaticum*** | **Bsyl502-3** | **43.37** | **6.37** | **14.48** | **9.23** | **37.07** | **569.86** | **28.28** | **3.40** | **24.88** | **11.59** | **21.77** | **21.77** | **7.99** | **753** | **106** | **12** | **39.60** | **275** | **91** | **91** | **230** |
| ***B. sylvaticum*** | **Bsyl502-4** | **43.37** | **6.37** | **14.48** | **9.23** | **37.07** | **569.86** | **28.28** | **3.40** | **24.88** | **11.59** | **21.77** | **21.77** | **7.99** | **753** | **106** | **12** | **39.60** | **275** | **91** | **91** | **230** |
| ***B. sylvaticum*** | **Bsyl502-5** | **43.37** | **6.37** | **14.48** | **9.23** | **37.07** | **569.86** | **28.28** | **3.40** | **24.88** | **11.59** | **21.77** | **21.77** | **7.99** | **753** | **106** | **12** | **39.60** | **275** | **91** | **91** | **230** |
| ***B. sylvaticum*** | **Bsyl505-2** | **43.80** | **3.64** | **13.23** | **8.36** | **34.05** | **592.09** | **27.00** | **2.46** | **24.54** | **9.93** | **20.85** | **20.85** | **6.48** | **688** | **99** | **24** | **31.68** | **227** | **109** | **109** | **191** |
| ***B. sylvaticum*** | **Bsyl505-4** | **43.80** | **3.64** | **13.23** | **8.36** | **34.05** | **592.09** | **27.00** | **2.46** | **24.54** | **9.93** | **20.85** | **20.85** | **6.48** | **688** | **99** | **24** | **31.68** | **227** | **109** | **109** | **191** |
| ***B. sylvaticum*** | **Bsyl505-6** | **43.80** | **3.64** | **13.23** | **8.36** | **34.05** | **592.09** | **27.00** | **2.46** | **24.54** | **9.93** | **20.85** | **20.85** | **6.48** | **688** | **99** | **24** | **31.68** | **227** | **109** | **109** | **191** |
| ***B. sylvaticum*** | **Bsyl506-2** | **43.71** | **3.54** | **13.59** | **8.23** | **33.85** | **588.44** | **27.21** | **2.90** | **24.31** | **10.34** | **21.16** | **21.16** | **6.89** | **662** | **95** | **22** | **32.08** | **220** | **103** | **103** | **187** |
| ***B. sylvaticum*** | **Bsyl506-3** | **43.71** | **3.54** | **13.59** | **8.23** | **33.85** | **588.44** | **27.21** | **2.90** | **24.31** | **10.34** | **21.16** | **21.16** | **6.89** | **662** | **95** | **22** | **32.08** | **220** | **103** | **103** | **187** |
| ***B. sylvaticum*** | **Bsyl506-6** | **43.71** | **3.54** | **13.59** | **8.23** | **33.85** | **588.44** | **27.21** | **2.90** | **24.31** | **10.34** | **21.16** | **21.16** | **6.89** | **662** | **95** | **22** | **32.08** | **220** | **103** | **103** | **187** |
| ***B. sylvaticum*** | **Bsyl508-2** | **43.04** | **2.86** | **13.92** | **7.99** | **33.91** | **566.42** | **27.03** | **3.46** | **23.57** | **10.93** | **21.20** | **21.20** | **7.49** | **615** | **80** | **17** | **31.24** | **203** | **91** | **91** | **182** |
| ***B. sylvaticum*** | **Bsyl508-3** | **43.04** | **2.86** | **13.92** | **7.99** | **33.91** | **566.42** | **27.03** | **3.46** | **23.57** | **10.93** | **21.20** | **21.20** | **7.49** | **615** | **80** | **17** | **31.24** | **203** | **91** | **91** | **182** |
| ***B. sylvaticum*** | **Bsyl508-6** | **43.04** | **2.86** | **13.92** | **7.99** | **33.91** | **566.42** | **27.03** | **3.46** | **23.57** | **10.93** | **21.20** | **21.20** | **7.49** | **615** | **80** | **17** | **31.24** | **203** | **91** | **91** | **182** |
| ***B. sylvaticum*** | **Bsyl54-3** | **35.02** | **-5.01** | **12.94** | **11.66** | **39.18** | **628.66** | **30.16** | **0.40** | **29.76** | **6.43** | **21.07** | **21.07** | **5.68** | **960** | **166** | **1** | **78.65** | **479** | **16** | **16** | **403** |
| ***B. sylvaticum*** | **Bsyl54-9** | **35.02** | **-5.01** | **12.94** | **11.66** | **39.18** | **628.66** | **30.16** | **0.40** | **29.76** | **6.43** | **21.07** | **21.07** | **5.68** | **960** | **166** | **1** | **78.65** | **479** | **16** | **16** | **403** |
| ***B. sylvaticum*** | **Bsyl550-10** | **40.50** | **-6.18** | **9.29** | **7.75** | **33.48** | **567.48** | **23.00** | **-0.16** | **23.16** | **4.26** | **16.96** | **16.96** | **3.27** | **1289** | **194** | **18** | **54.37** | **524** | **90** | **90** | **519** |
| ***B. sylvaticum*** | **Bsyl550-1B** | **40.50** | **-6.18** | **9.29** | **7.75** | **33.48** | **567.48** | **23.00** | **-0.16** | **23.16** | **4.26** | **16.96** | **16.96** | **3.27** | **1289** | **194** | **18** | **54.37** | **524** | **90** | **90** | **519** |
| ***B. sylvaticum*** | **Bsyl550-3B** | **40.50** | **-6.18** | **9.29** | **7.75** | **33.48** | **567.48** | **23.00** | **-0.16** | **23.16** | **4.26** | **16.96** | **16.96** | **3.27** | **1289** | **194** | **18** | **54.37** | **524** | **90** | **90** | **519** |
| ***B. sylvaticum*** | **Bsyl550-8b** | **40.50** | **-6.18** | **9.29** | **7.75** | **33.48** | **567.48** | **23.00** | **-0.16** | **23.16** | **4.26** | **16.96** | **16.96** | **3.27** | **1289** | **194** | **18** | **54.37** | **524** | **90** | **90** | **519** |
| ***B. sylvaticum*** | **Bsyl552-1** | **36.15** | **-5.63** | **17.25** | **7.94** | **38.11** | **454.38** | **28.76** | **7.92** | **20.84** | **13.14** | **22.80** | **23.23** | **12.06** | **891** | **146** | **0** | **77.34** | **419** | **16** | **23** | **409** |
| ***B. sylvaticum*** | **Bsyl552-2** | **36.15** | **-5.63** | **17.25** | **7.94** | **38.11** | **454.38** | **28.76** | **7.92** | **20.84** | **13.14** | **22.80** | **23.23** | **12.06** | **891** | **146** | **0** | **77.34** | **419** | **16** | **23** | **409** |
| ***B. sylvaticum*** | **Bsyl552-5** | **36.15** | **-5.63** | **17.25** | **7.94** | **38.11** | **454.38** | **28.76** | **7.92** | **20.84** | **13.14** | **22.80** | **23.23** | **12.06** | **891** | **146** | **0** | **77.34** | **419** | **16** | **23** | **409** |
| ***B. sylvaticum*** | **Bsyl553-2B** | **36.72** | **-4.61** | **17.64** | **10.60** | **43.24** | **508.68** | **31.16** | **6.65** | **24.51** | **12.81** | **24.02** | **24.34** | **11.94** | **560** | **102** | **1** | **73.97** | **285** | **18** | **23** | **252** |
| ***B. sylvaticum*** | **Bsyl553-3b** | **36.72** | **-4.61** | **17.64** | **10.60** | **43.24** | **508.68** | **31.16** | **6.65** | **24.51** | **12.81** | **24.02** | **24.34** | **11.94** | **560** | **102** | **1** | **73.97** | **285** | **18** | **23** | **252** |
| ***B. sylvaticum*** | **Bsyl553-5b** | **36.72** | **-4.61** | **17.64** | **10.60** | **43.24** | **508.68** | **31.16** | **6.65** | **24.51** | **12.81** | **24.02** | **24.34** | **11.94** | **560** | **102** | **1** | **73.97** | **285** | **18** | **23** | **252** |
| ***B. sylvaticum*** | **Bsyl554-1** | **36.89** | **-4.06** | **12.14** | **11.92** | **39.89** | **631.63** | **29.82** | **-0.07** | **29.89** | **6.17** | **20.43** | **20.44** | **5.20** | **678** | **93** | **5** | **57.96** | **273** | **35** | **37** | **267** |
| ***B. sylvaticum*** | **Bsyl554-3B** | **36.89** | **-4.06** | **12.14** | **11.92** | **39.89** | **631.63** | **29.82** | **-0.07** | **29.89** | **6.17** | **20.43** | **20.44** | **5.20** | **678** | **93** | **5** | **57.96** | **273** | **35** | **37** | **267** |
| ***B. sylvaticum*** | **Bsyl554-6** | **36.89** | **-4.06** | **12.14** | **11.92** | **39.89** | **631.63** | **29.82** | **-0.07** | **29.89** | **6.17** | **20.43** | **20.44** | **5.20** | **678** | **93** | **5** | **57.96** | **273** | **35** | **37** | **267** |
| ***B. sylvaticum*** | **Bsyl554-7** | **36.89** | **-4.06** | **12.14** | **11.92** | **39.89** | **631.63** | **29.82** | **-0.07** | **29.89** | **6.17** | **20.43** | **20.44** | **5.20** | **678** | **93** | **5** | **57.96** | **273** | **35** | **37** | **267** |
| ***B. sylvaticum*** | **Bsyl555-1** | **37.13** | **-3.45** | **11.53** | **12.62** | **39.10** | **686.72** | **30.60** | **-1.67** | **32.27** | **4.84** | **20.42** | **20.53** | **3.82** | **552** | **74** | **6** | **52.31** | **212** | **36** | **37** | **206** |
| ***B. sylvaticum*** | **Bsyl555-3B** | **37.13** | **-3.45** | **11.53** | **12.62** | **39.10** | **686.72** | **30.60** | **-1.67** | **32.27** | **4.84** | **20.42** | **20.53** | **3.82** | **552** | **74** | **6** | **52.31** | **212** | **36** | **37** | **206** |
| ***B. sylvaticum*** | **Bsyl555-5** | **37.13** | **-3.45** | **11.53** | **12.62** | **39.10** | **686.72** | **30.60** | **-1.67** | **32.27** | **4.84** | **20.42** | **20.53** | **3.82** | **552** | **74** | **6** | **52.31** | **212** | **36** | **37** | **206** |
| ***B. sylvaticum*** | **Bsyl555-8** | **37.13** | **-3.45** | **11.53** | **12.62** | **39.10** | **686.72** | **30.60** | **-1.67** | **32.27** | **4.84** | **20.42** | **20.53** | **3.82** | **552** | **74** | **6** | **52.31** | **212** | **36** | **37** | **206** |
| ***B. sylvaticum*** | **Bsyl557-2** | **42.39** | **2.74** | **11.53** | **8.19** | **34.90** | **553.21** | **24.41** | **0.95** | **23.46** | **12.38** | **18.65** | **18.68** | **5.41** | **804** | **93** | **37** | **21.13** | **240** | **159** | **170** | **194** |
| ***B. sylvaticum*** | **Bsyl557-3** | **42.39** | **2.74** | **11.53** | **8.19** | **34.90** | **553.21** | **24.41** | **0.95** | **23.46** | **12.38** | **18.65** | **18.68** | **5.41** | **804** | **93** | **37** | **21.13** | **240** | **159** | **170** | **194** |
| ***B. sylvaticum*** | **Bsyl557-7** | **42.39** | **2.74** | **11.53** | **8.19** | **34.90** | **553.21** | **24.41** | **0.95** | **23.46** | **12.38** | **18.65** | **18.68** | **5.41** | **804** | **93** | **37** | **21.13** | **240** | **159** | **170** | **194** |
| ***B. sylvaticum*** | **Bsyl557-8** | **42.39** | **2.74** | **11.53** | **8.19** | **34.90** | **553.21** | **24.41** | **0.95** | **23.46** | **12.38** | **18.65** | **18.68** | **5.41** | **804** | **93** | **37** | **21.13** | **240** | **159** | **170** | **194** |
| ***B. sylvaticum*** | **Bsyl59-1** | **33.53** | **-4.69** | **12.44** | **12.69** | **40.06** | **670.99** | **30.34** | **-1.34** | **31.67** | **11.09** | **20.97** | **20.98** | **4.56** | **239** | **34** | **6** | **39.87** | **91** | **34** | **35** | **49** |
| ***B. sylvaticum*** | **Bsyl59-2** | **33.53** | **-4.69** | **12.44** | **12.69** | **40.06** | **670.99** | **30.34** | **-1.34** | **31.67** | **11.09** | **20.97** | **20.98** | **4.56** | **239** | **34** | **6** | **39.87** | **91** | **34** | **35** | **49** |
| ***B. sylvaticum*** | **Bsyl59-4** | **33.53** | **-4.69** | **12.44** | **12.69** | **40.06** | **670.99** | **30.34** | **-1.34** | **31.67** | **11.09** | **20.97** | **20.98** | **4.56** | **239** | **34** | **6** | **39.87** | **91** | **34** | **35** | **49** |
| ***B. sylvaticum*** | **Bsyl59-5** | **33.53** | **-4.69** | **12.44** | **12.69** | **40.06** | **670.99** | **30.34** | **-1.34** | **31.67** | **11.09** | **20.97** | **20.98** | **4.56** | **239** | **34** | **6** | **39.87** | **91** | **34** | **35** | **49** |
| ***B. sylvaticum*** | **Bsyl62-8** | **33.52** | **-5.17** | **12.16** | **12.09** | **39.61** | **644.73** | **29.52** | **-1.00** | **30.52** | **7.76** | **20.34** | **20.58** | **4.72** | **384** | **50** | **5** | **45.90** | **140** | **33** | **33** | **114** |
| ***B. sylvaticum*** | **Bsyl63-2A** | **34.06** | **-5.53** | **16.65** | **12.02** | **42.00** | **572.36** | **32.88** | **4.27** | **28.62** | **10.10** | **23.74** | **24.28** | **10.10** | **593** | **87** | **1** | **66.32** | **244** | **16** | **17** | **244** |
| ***B. sylvaticum*** | **Bsyl63-2B** | **34.06** | **-5.53** | **16.65** | **12.02** | **42.00** | **572.36** | **32.88** | **4.27** | **28.62** | **10.10** | **23.74** | **24.28** | **10.10** | **593** | **87** | **1** | **66.32** | **244** | **16** | **17** | **244** |
| ***B. sylvaticum*** | **Bsyl63-4** | **34.06** | **-5.53** | **16.65** | **12.02** | **42.00** | **572.36** | **32.88** | **4.27** | **28.62** | **10.10** | **23.74** | **24.28** | **10.10** | **593** | **87** | **1** | **66.32** | **244** | **16** | **17** | **244** |
| ***B. sylvaticum*** | **Bsyl63-5** | **34.06** | **-5.53** | **16.65** | **12.02** | **42.00** | **572.36** | **32.88** | **4.27** | **28.62** | **10.10** | **23.74** | **24.28** | **10.10** | **593** | **87** | **1** | **66.32** | **244** | **16** | **17** | **244** |
| ***B. sylvaticum*** | **Bsyl72-1** | **52.24** | **0.02** | **9.78** | **7.99** | **37.81** | **487.30** | **21.92** | **0.79** | **21.14** | **10.50** | **5.99** | **15.94** | **4.21** | **569** | **54** | **35** | **11.67** | **158** | **123** | **145** | **134** |
| ***B. sylvaticum*** | **Bsyl72-3** | **52.24** | **0.02** | **9.78** | **7.99** | **37.81** | **487.30** | **21.92** | **0.79** | **21.14** | **10.50** | **5.99** | **15.94** | **4.21** | **569** | **54** | **35** | **11.67** | **158** | **123** | **145** | **134** |
| ***B. sylvaticum*** | **Bsyl72-4** | **52.24** | **0.02** | **9.78** | **7.99** | **37.81** | **487.30** | **21.92** | **0.79** | **21.14** | **10.50** | **5.99** | **15.94** | **4.21** | **569** | **54** | **35** | **11.67** | **158** | **123** | **145** | **134** |
| ***B. sylvaticum*** | **Bsyl73-1** | **52.19** | **1.13** | **9.55** | **7.78** | **37.19** | **483.89** | **21.71** | **0.80** | **20.91** | **7.23** | **5.70** | **15.70** | **4.07** | **598** | **62** | **38** | **15.07** | **176** | **122** | **155** | **147** |
| ***B. sylvaticum*** | **Bsyl73-3** | **52.19** | **1.13** | **9.55** | **7.78** | **37.19** | **483.89** | **21.71** | **0.80** | **20.91** | **7.23** | **5.70** | **15.70** | **4.07** | **598** | **62** | **38** | **15.07** | **176** | **122** | **155** | **147** |
| ***B. sylvaticum*** | **Bsyl73-4** | **52.19** | **1.13** | **9.55** | **7.78** | **37.19** | **483.89** | **21.71** | **0.80** | **20.91** | **7.23** | **5.70** | **15.70** | **4.07** | **598** | **62** | **38** | **15.07** | **176** | **122** | **155** | **147** |
| ***B. sylvaticum*** | **Bsyl73-3** | **52.19** | **1.13** | **9.55** | **7.78** | **37.19** | **483.89** | **21.71** | **0.80** | **20.91** | **7.23** | **5.70** | **15.70** | **4.07** | **598** | **62** | **38** | **15.07** | **176** | **122** | **155** | **147** |
| ***B. sylvaticum*** | **Bsyl73-4** | **52.19** | **1.13** | **9.55** | **7.78** | **37.19** | **483.89** | **21.71** | **0.80** | **20.91** | **7.23** | **5.70** | **15.70** | **4.07** | **598** | **62** | **38** | **15.07** | **176** | **122** | **155** | **147** |
| *B. sylvaticum* | Bsyl_1 | 48.20 | 10.90 | 7.94 | 8.61 | 32.66 | 678.44 | 22.34 | -4.02 | 26.36 | 16.34 | 0.89 | 16.34 | -0.30 | 918 | 113 | 49 | 29.87 | 328 | 154 | 328 | 162 |
| *B. sylvaticum* | Bsyl_2 | 48.00 | 9.30 | 7.51 | 8.43 | 32.01 | 679.82 | 22.17 | -4.15 | 26.32 | 15.95 | 0.41 | 15.95 | -0.73 | 867 | 109 | 50 | 27.45 | 302 | 153 | 302 | 165 |
| *B. sylvaticum* | Bsyl_3 | 48.40 | 8.60 | 7.92 | 8.81 | 34.30 | 646.21 | 22.36 | -3.31 | 25.67 | 14.34 | 12.62 | 16.01 | 0.21 | 1008 | 99 | 66 | 12.54 | 285 | 221 | 265 | 265 |
| *B. sylvaticum* | Bsyl_4 | 54.00 | 11.70 | 8.49 | 6.69 | 28.57 | 635.04 | 21.62 | -1.78 | 23.40 | 16.43 | 3.87 | 16.43 | 0.94 | 627 | 70 | 35 | 18.14 | 194 | 122 | 194 | 138 |
| *B. sylvaticum* | Bsyl_5 | 51.60 | 13.80 | 8.84 | 8.23 | 31.68 | 681.03 | 23.26 | -2.74 | 25.99 | 17.25 | 4.13 | 17.25 | 0.52 | 617 | 81 | 34 | 25.47 | 208 | 119 | 208 | 128 |
| *B. sylvaticum* | Bsyl_6 | 47.80 | 8.50 | 7.64 | 9.30 | 34.31 | 670.53 | 22.70 | -4.39 | 27.09 | 14.40 | 3.33 | 16.00 | -0.46 | 930 | 101 | 59 | 15.28 | 271 | 200 | 271 | 236 |
| *B. sylvaticum* | Bsyl_7 | 54.30 | 10.10 | 8.56 | 7.02 | 31.02 | 596.29 | 21.38 | -1.24 | 22.63 | 5.58 | 3.94 | 16.10 | 1.57 | 776 | 78 | 43 | 18.00 | 222 | 148 | 214 | 187 |
| *B. sylvaticum* | Bsyl_8 | 51.40 | 6.40 | 10.23 | 7.69 | 33.84 | 576.45 | 23.00 | 0.28 | 22.72 | 16.03 | 6.14 | 17.43 | 3.36 | 738 | 74 | 45 | 13.04 | 205 | 155 | 202 | 178 |
| *B. sylvaticum* | Bsyl_9 | 51.00 | 14.00 | 8.87 | 7.76 | 29.56 | 697.55 | 23.35 | -2.91 | 26.26 | 17.46 | 4.26 | 17.46 | 0.37 | 716 | 88 | 38 | 26.50 | 250 | 132 | 250 | 145 |
| *B. sylvaticum* | Bsyl_10 | 50.60 | 9.50 | 8.32 | 8.55 | 33.42 | 642.57 | 22.66 | -2.92 | 25.58 | 4.49 | 3.97 | 16.41 | 0.62 | 756 | 75 | 53 | 10.58 | 205 | 166 | 198 | 191 |
| *B. sylvaticum* | Bsyl_11 | 48.90 | 12.00 | 8.40 | 8.30 | 30.57 | 721.17 | 23.24 | -3.91 | 27.14 | 17.24 | 4.01 | 17.24 | -0.47 | 692 | 84 | 40 | 25.49 | 239 | 129 | 239 | 138 |
| *B. sylvaticum* | Bsyl_12 | 54.50 | 9.30 | 8.36 | 6.90 | 30.91 | 588.01 | 20.94 | -1.38 | 22.33 | 9.07 | 3.78 | 15.73 | 1.43 | 827 | 91 | 43 | 23.49 | 262 | 143 | 228 | 184 |
| *B. sylvaticum* | Bsyl_13 | 51.20 | 9.80 | 7.81 | 7.73 | 31.87 | 626.49 | 21.22 | -3.05 | 24.26 | 14.13 | 3.42 | 15.60 | 0.24 | 843 | 82 | 56 | 11.29 | 234 | 182 | 230 | 203 |
| *B. sylvaticum* | Bsyl_14 | 51.10 | 8.70 | 7.44 | 8.54 | 34.00 | 619.54 | 21.46 | -3.66 | 25.12 | 1.20 | 3.06 | 15.25 | 0.05 | 921 | 98 | 61 | 13.71 | 275 | 204 | 223 | 256 |
| *B. sylvaticum* | Bsyl_15 | 48.30 | 7.90 | 9.81 | 7.65 | 31.21 | 641.48 | 23.42 | -1.10 | 24.51 | 16.18 | 3.23 | 17.82 | 2.09 | 1040 | 112 | 69 | 17.21 | 326 | 208 | 302 | 227 |
| *B. sylvaticum* | Bsyl_16 | 52.60 | 11.60 | 9.27 | 8.38 | 32.66 | 657.94 | 23.85 | -1.82 | 25.67 | 17.45 | 4.91 | 17.45 | 1.36 | 548 | 61 | 34 | 18.91 | 174 | 109 | 174 | 120 |
| *B. sylvaticum* | Bsyl_17 | 53.70 | 10.50 | 8.43 | 7.70 | 32.20 | 613.02 | 22.01 | -1.90 | 23.91 | 16.16 | 3.90 | 16.16 | 1.17 | 759 | 75 | 46 | 15.62 | 217 | 154 | 217 | 186 |
| *B. sylvaticum* | Bsyl_18 | 51.70 | 10.80 | 6.64 | 7.68 | 31.73 | 631.42 | 20.10 | -4.09 | 24.19 | 0.37 | 9.60 | 14.55 | -0.88 | 1002 | 109 | 68 | 14.70 | 297 | 225 | 250 | 272 |
| *B. sylvaticum* | Bsyl_19 | 52.90 | 12.40 | 9.17 | 8.09 | 31.07 | 684.65 | 23.74 | -2.31 | 26.06 | 17.66 | 4.63 | 17.66 | 0.90 | 553 | 72 | 26 | 25.33 | 185 | 106 | 185 | 129 |
| *B. sylvaticum* | Bsyl_20 | 52.80 | 7.30 | 9.13 | 7.66 | 34.24 | 568.15 | 21.73 | -0.65 | 22.38 | 16.23 | 4.87 | 16.23 | 2.39 | 789 | 80 | 47 | 16.30 | 222 | 158 | 222 | 191 |
| *B. sylvaticum* | Bsyl_21 | 52.40 | 13.40 | 9.23 | 8.13 | 31.03 | 694.93 | 23.86 | -2.34 | 26.20 | 17.94 | 4.61 | 17.94 | 0.88 | 559 | 63 | 35 | 19.70 | 176 | 114 | 176 | 128 |
| *B. sylvaticum* | Bsyl_22 | 53.40 | 9.70 | 8.84 | 7.76 | 32.41 | 615.15 | 22.35 | -1.60 | 23.95 | 16.54 | 4.44 | 16.54 | 1.50 | 743 | 72 | 43 | 15.22 | 209 | 147 | 209 | 178 |
| *B. sylvaticum* | Bsyl_23 | 48.80 | 9.50 | 9.19 | 9.16 | 34.96 | 650.56 | 23.72 | -2.48 | 26.20 | 15.81 | 2.47 | 17.32 | 1.31 | 930 | 102 | 63 | 15.65 | 287 | 195 | 273 | 208 |
| *B. sylvaticum* | Bsyl_24 | 50.20 | 12.00 | 6.20 | 7.52 | 29.95 | 662.06 | 20.20 | -4.90 | 25.11 | 14.45 | 1.76 | 14.45 | -1.74 | 858 | 93 | 56 | 17.06 | 261 | 177 | 261 | 216 |
| *B. sylvaticum* | Bsyl_25 | 53.90 | 9.60 | 8.61 | 7.74 | 33.02 | 602.71 | 21.86 | -1.59 | 23.44 | 5.40 | 4.19 | 16.15 | 1.40 | 840 | 82 | 47 | 18.43 | 241 | 158 | 236 | 198 |
| *B. sylvaticum* | Bsyl_26 | 52.20 | 9.10 | 9.65 | 7.66 | 32.35 | 613.85 | 22.88 | -0.79 | 23.68 | 17.29 | 5.34 | 17.29 | 2.26 | 771 | 75 | 50 | 12.89 | 217 | 162 | 217 | 193 |
| *B. sylvaticum* | Bsyl_27 | 50.60 | 11.40 | 8.36 | 8.51 | 32.75 | 670.85 | 22.94 | -3.03 | 25.98 | 15.22 | 1.30 | 16.74 | 0.30 | 704 | 81 | 41 | 19.90 | 223 | 139 | 221 | 146 |
| *B. sylvaticum* | Bsyl_28 | 51.70 | 7.30 | 9.76 | 8.12 | 34.25 | 582.36 | 23.09 | -0.61 | 23.70 | 3.91 | 5.60 | 17.04 | 2.75 | 834 | 81 | 53 | 13.96 | 233 | 174 | 231 | 208 |
| *B. sylvaticum* | Bsyl_29 | 50.90 | 7.20 | 9.68 | 8.18 | 34.58 | 586.43 | 22.82 | -0.84 | 23.66 | 15.60 | 5.58 | 17.02 | 2.64 | 869 | 91 | 54 | 15.09 | 256 | 181 | 249 | 203 |
| *B. sylvaticum* | Bsyl_30 | 50.70 | 13.40 | 6.68 | 7.44 | 29.26 | 694.98 | 20.62 | -4.81 | 25.43 | 15.15 | 7.08 | 15.15 | -1.83 | 953 | 119 | 57 | 22.32 | 311 | 203 | 311 | 214 |
| *B. sylvaticum* | Bsyl_31 | 53.10 | 10.70 | 8.60 | 8.13 | 32.85 | 632.50 | 22.52 | -2.23 | 24.75 | 16.52 | 4.04 | 16.52 | 1.04 | 670 | 69 | 42 | 14.79 | 199 | 140 | 199 | 163 |
| *B. sylvaticum* | Bsyl_32 | 52.50 | 8.50 | 9.30 | 7.99 | 33.43 | 602.53 | 22.98 | -0.93 | 23.90 | 16.87 | 4.97 | 16.87 | 2.14 | 698 | 76 | 42 | 15.15 | 199 | 143 | 199 | 166 |
| *B. sylvaticum* | Bsyl_33 | 49.10 | 12.80 | 6.85 | 8.35 | 31.40 | 694.18 | 21.38 | -5.21 | 26.58 | 15.47 | 2.30 | 15.47 | -1.56 | 920 | 110 | 55 | 21.73 | 301 | 177 | 301 | 212 |
| *B. sylvaticum* | Bsyl_34 | 50.60 | 8.30 | 8.60 | 8.12 | 32.68 | 633.43 | 22.52 | -2.34 | 24.86 | 2.01 | 4.41 | 16.56 | 0.97 | 774 | 82 | 52 | 12.48 | 221 | 171 | 190 | 206 |
| *B. sylvaticum* | Bsyl_35 | 52.50 | 0.70 | 9.67 | 7.43 | 37.41 | 469.25 | 21.00 | 1.14 | 19.86 | 7.36 | 5.94 | 15.66 | 4.33 | 642 | 67 | 39 | 13.98 | 185 | 137 | 163 | 154 |
| *B. sylvaticum* | Bsyl_36 | 52.50 | 1.60 | 9.72 | 6.77 | 34.59 | 471.53 | 20.66 | 1.10 | 19.56 | 7.73 | 5.82 | 15.73 | 4.34 | 575 | 63 | 35 | 17.09 | 174 | 117 | 148 | 141 |
| *B. sylvaticum* | Bsyl_37 | 50.10 | 36.30 | 7.14 | 7.71 | 22.56 | 1002.69 | 24.86 | -9.32 | 34.18 | 19.16 | 0.88 | 19.16 | -5.29 | 594 | 75 | 34 | 22.09 | 191 | 111 | 191 | 128 |
| *B. sylvaticum* | Bsyl_38 | 52.30 | 35.40 | 5.95 | 7.83 | 23.66 | 977.16 | 23.31 | -9.80 | 33.11 | 17.70 | -4.86 | 17.70 | -6.10 | 647 | 85 | 34 | 28.11 | 226 | 113 | 226 | 121 |
| *B. sylvaticum* | Bsyl_39 | 54.90 | -2.10 | 7.91 | 7.25 | 38.74 | 426.24 | 18.67 | -0.05 | 18.72 | 3.90 | 9.17 | 13.42 | 3.11 | 918 | 95 | 60 | 16.47 | 277 | 184 | 207 | 256 |
| *B. sylvaticum* | Bsyl_40 | 43.80 | 6.10 | 11.96 | 11.05 | 39.39 | 617.83 | 27.71 | -0.35 | 28.06 | 8.53 | 19.86 | 19.86 | 4.88 | 778 | 101 | 30 | 25.99 | 251 | 138 | 138 | 197 |
| *B. sylvaticum* | Bsyl_41 | 44.00 | 7.60 | 9.90 | 6.52 | 29.24 | 587.84 | 22.35 | 0.04 | 22.30 | 10.56 | 17.48 | 17.48 | 3.37 | 906 | 121 | 29 | 31.05 | 299 | 134 | 134 | 231 |
| *B. sylvaticum* | Bsyl_42 | 43.60 | 7.00 | 14.43 | 7.75 | 33.72 | 556.83 | 26.78 | 3.80 | 22.98 | 11.64 | 21.50 | 21.55 | 8.14 | 847 | 133 | 16 | 44.79 | 330 | 94 | 127 | 254 |
| *B. sylvaticum* | Bsyl_43 | 44.30 | 5.80 | 11.09 | 11.08 | 37.70 | 660.28 | 27.66 | -1.74 | 29.40 | 11.53 | 19.50 | 19.50 | 3.35 | 848 | 96 | 42 | 20.21 | 266 | 178 | 178 | 195 |
| *B. sylvaticum* | Bsyl_44 | 43.30 | 5.70 | 11.95 | 8.78 | 35.77 | 579.89 | 26.01 | 1.45 | 24.56 | 9.04 | 19.43 | 19.43 | 5.51 | 817 | 116 | 18 | 36.74 | 286 | 106 | 106 | 234 |
| *B. sylvaticum* | Bsyl_45 | 44.90 | 5.80 | 9.83 | 10.68 | 36.64 | 665.67 | 26.16 | -2.98 | 29.14 | 10.29 | 18.21 | 18.21 | 1.89 | 1014 | 118 | 55 | 18.77 | 314 | 220 | 220 | 234 |
| *B. sylvaticum* | Bsyl_46 | 44.60 | 5.20 | 9.73 | 9.87 | 35.87 | 639.47 | 25.46 | -2.07 | 27.53 | 10.25 | 17.91 | 17.91 | 2.36 | 1082 | 125 | 53 | 22.59 | 347 | 210 | 210 | 245 |
| *B. sylvaticum* | Bsyl_47 | 45.60 | 6.90 | 4.48 | 8.15 | 32.58 | 623.34 | 18.42 | -6.60 | 25.02 | -1.42 | 12.30 | 12.30 | -2.60 | 1415 | 145 | 87 | 16.04 | 419 | 278 | 278 | 406 |
| *B. sylvaticum* | Bsyl_48 | 46.20 | 5.20 | 11.17 | 8.69 | 32.90 | 659.27 | 25.90 | -0.52 | 26.42 | 11.39 | 4.54 | 19.43 | 3.19 | 968 | 99 | 63 | 15.39 | 287 | 204 | 227 | 218 |
| *B. sylvaticum* | Bsyl_49 | 58.00 | -4.10 | 7.83 | 6.82 | 39.82 | 396.54 | 17.24 | 0.10 | 17.14 | 5.79 | 9.11 | 12.85 | 3.22 | 1058 | 115 | 61 | 21.74 | 328 | 193 | 229 | 286 |
| *B. sylvaticum* | Bsyl_50 | 57.50 | -6.40 | 8.31 | 4.91 | 33.67 | 365.23 | 16.23 | 1.65 | 14.58 | 6.77 | 9.16 | 13.03 | 4.38 | 1796 | 211 | 85 | 32.17 | 624 | 277 | 337 | 517 |
| *B. sylvaticum* | Bsyl_51 | 54.90 | -4.00 | 8.40 | 6.76 | 37.63 | 427.98 | 18.61 | 0.65 | 17.96 | 4.02 | 10.08 | 13.88 | 3.38 | 1167 | 131 | 66 | 26.66 | 388 | 205 | 224 | 350 |
| *B. sylvaticum* | Bsyl_52 | 56.80 | -2.60 | 7.88 | 8.03 | 39.39 | 473.01 | 19.16 | -1.23 | 20.38 | 11.48 | 4.72 | 13.88 | 2.25 | 676 | 70 | 42 | 15.71 | 196 | 137 | 166 | 171 |
| *B. sylvaticum* | Bsyl_53 | 55.70 | -5.20 | 9.30 | 5.59 | 36.63 | 375.09 | 17.88 | 2.62 | 15.26 | 5.82 | 10.61 | 14.06 | 5.07 | 1629 | 192 | 81 | 29.21 | 551 | 258 | 317 | 498 |
| *B. sylvaticum* | Bsyl_54 | 54.70 | -4.90 | 8.62 | 6.46 | 38.25 | 391.74 | 17.99 | 1.09 | 16.90 | 6.85 | 9.82 | 13.60 | 4.18 | 1053 | 117 | 55 | 25.41 | 342 | 182 | 219 | 303 |
| *B. sylvaticum* | Bsyl_55 | 49.50 | 11.90 | 8.09 | 8.62 | 31.60 | 724.79 | 23.28 | -4.01 | 27.29 | 17.02 | 3.59 | 17.02 | -0.73 | 779 | 86 | 48 | 18.39 | 243 | 154 | 243 | 185 |
| *B. sylvaticum* | Bsyl_56 | 47.80 | 12.60 | 8.08 | 9.13 | 33.32 | 695.85 | 22.60 | -4.79 | 27.39 | 16.57 | 0.83 | 16.57 | -0.49 | 1324 | 165 | 77 | 28.03 | 477 | 256 | 477 | 257 |
| *B. sylvaticum* | Bsyl_57 | 49.00 | 11.10 | 7.54 | 8.22 | 31.74 | 677.11 | 21.90 | -4.00 | 25.90 | 14.45 | 0.47 | 15.97 | -0.63 | 834 | 94 | 54 | 19.04 | 264 | 169 | 264 | 185 |
| *B. sylvaticum* | Bsyl_58 | 47.70 | 11.90 | 6.69 | 7.70 | 31.51 | 640.89 | 19.77 | -4.67 | 24.44 | 14.55 | -1.02 | 14.55 | -1.02 | 1335 | 178 | 71 | 33.45 | 505 | 231 | 505 | 231 |
| *B. sylvaticum* | Bsyl_59 | 48.60 | 13.60 | 7.88 | 8.52 | 31.05 | 719.38 | 22.59 | -4.84 | 27.42 | 16.62 | 3.47 | 16.62 | -1.04 | 960 | 109 | 62 | 19.21 | 306 | 193 | 306 | 229 |
| *B. sylvaticum* | Bsyl_60 | 50.10 | 9.10 | 9.38 | 8.10 | 32.11 | 656.75 | 23.53 | -1.68 | 25.21 | 5.28 | 5.13 | 17.62 | 1.47 | 814 | 84 | 53 | 13.31 | 226 | 176 | 217 | 209 |
| *B. sylvaticum* | Bsyl_61 | 50.10 | 11.20 | 7.92 | 8.44 | 32.45 | 676.89 | 22.41 | -3.60 | 26.01 | 16.33 | 3.46 | 16.33 | -0.26 | 782 | 82 | 51 | 15.89 | 225 | 162 | 225 | 197 |
| *B. sylvaticum* | Bsyl_62 | 48.30 | 11.70 | 8.39 | 9.12 | 33.37 | 693.06 | 23.26 | -4.08 | 27.34 | 16.89 | 1.15 | 16.89 | -0.12 | 835 | 112 | 43 | 33.56 | 310 | 135 | 310 | 139 |
| *B. sylvaticum* | Bsyl_63 | 57.20 | 17.00 | 7.47 | 7.09 | 28.41 | 655.35 | 21.51 | -3.44 | 24.94 | 15.52 | 1.82 | 16.04 | 0.18 | 477 | 52 | 27 | 22.93 | 149 | 85 | 141 | 103 |
| *B. sylvaticum* | Bsyl_64 | 56.60 | 16.60 | 7.28 | 7.09 | 28.47 | 658.63 | 21.35 | -3.56 | 24.92 | 15.14 | 1.81 | 15.82 | -0.22 | 514 | 54 | 30 | 19.84 | 155 | 97 | 147 | 120 |
| *B. sylvaticum* | Bsyl_65 | 53.40 | -1.20 | 9.10 | 7.96 | 38.71 | 463.55 | 20.98 | 0.40 | 20.57 | 4.77 | 13.34 | 15.05 | 3.84 | 697 | 72 | 47 | 12.57 | 202 | 158 | 165 | 190 |
| *B. sylvaticum* | Bsyl_66 | 47.20 | 13.10 | 5.80 | 10.26 | 36.49 | 677.74 | 20.41 | -7.70 | 28.11 | 14.00 | -1.23 | 14.00 | -2.61 | 1176 | 168 | 52 | 39.96 | 472 | 179 | 472 | 180 |
| *B. sylvaticum* | Bsyl_67 | 48.00 | 13.20 | 7.77 | 8.63 | 32.31 | 690.20 | 21.92 | -4.79 | 26.70 | 16.20 | 0.57 | 16.20 | -0.71 | 1219 | 159 | 68 | 30.56 | 451 | 228 | 451 | 231 |
| *B. sylvaticum* | Bsyl_68 | 47.80 | 13.80 | 6.83 | 8.06 | 31.79 | 663.63 | 20.30 | -5.05 | 25.35 | 14.99 | -0.15 | 14.99 | -1.17 | 1375 | 181 | 77 | 27.67 | 490 | 269 | 490 | 270 |
| *B. sylvaticum* | Bsyl_69 | 47.30 | 12.30 | 3.62 | 7.56 | 30.72 | 638.07 | 16.74 | -7.88 | 24.62 | 11.45 | -3.97 | 11.45 | -3.97 | 1258 | 171 | 63 | 34.60 | 478 | 205 | 478 | 205 |
| *B. sylvaticum* | Bsyl_70 | 47.10 | 13.90 | 2.49 | 8.32 | 34.15 | 611.77 | 15.27 | -9.09 | 24.36 | 10.05 | -4.14 | 10.05 | -4.70 | 1597 | 192 | 77 | 27.01 | 542 | 267 | 542 | 269 |
| *B. sylvaticum* | Bsyl_71 | 61.10 | 6.90 | 6.44 | 5.63 | 26.48 | 583.30 | 18.53 | -2.73 | 21.27 | 3.02 | 9.34 | 14.02 | -0.23 | 997 | 121 | 41 | 36.51 | 348 | 132 | 179 | 317 |
| *B. sylvaticum* | Bsyl_72 | 59.30 | 9.50 | 6.40 | 6.21 | 24.17 | 743.55 | 20.79 | -4.90 | 25.69 | 6.24 | 1.43 | 15.89 | -2.37 | 921 | 114 | 45 | 28.86 | 321 | 155 | 249 | 179 |
| *B. sylvaticum* | Bsyl_73 | 57.20 | -5.70 | 8.31 | 6.01 | 39.85 | 352.10 | 16.57 | 1.50 | 15.07 | 5.08 | 9.33 | 12.77 | 4.41 | 2149 | 256 | 96 | 34.78 | 755 | 316 | 370 | 658 |
| *B. sylvaticum* | Bsyl_74 | 55.80 | -3.30 | 6.83 | 6.62 | 36.93 | 423.90 | 17.10 | -0.81 | 17.92 | 2.86 | 8.25 | 12.31 | 2.07 | 1033 | 108 | 63 | 19.58 | 315 | 197 | 223 | 292 |
| *B. sylvaticum* | Bsyl_75 | 53.20 | 56.30 | 2.94 | 10.50 | 24.37 | 1244.05 | 24.83 | -18.26 | 43.09 | 17.58 | -4.66 | 17.58 | -12.61 | 521 | 63 | 26 | 26.97 | 172 | 84 | 172 | 112 |
| *B. sylvaticum* | Bsyl_76 | 52.80 | 40.60 | 5.03 | 8.71 | 22.35 | 1130.37 | 24.75 | -14.22 | 38.98 | 18.39 | -2.33 | 18.39 | -8.89 | 555 | 70 | 29 | 26.27 | 180 | 89 | 180 | 115 |
| *B. sylvaticum* | Bsyl_77 | 54.60 | 48.40 | 3.98 | 9.05 | 22.58 | 1168.63 | 24.63 | -15.44 | 40.07 | 17.77 | -9.39 | 17.77 | -10.73 | 456 | 63 | 21 | 36.62 | 174 | 72 | 174 | 77 |
| *B. sylvaticum* | Bsyl_78 | 50.50 | 44.90 | 5.91 | 9.56 | 23.79 | 1178.48 | 27.11 | -13.07 | 40.18 | 20.09 | -1.75 | 20.09 | -8.49 | 417 | 48 | 21 | 22.73 | 132 | 72 | 132 | 98 |
| *B. sylvaticum* | Bsyl_79 | 43.00 | 41.50 | 5.77 | 9.47 | 33.96 | 660.81 | 20.63 | -7.25 | 27.88 | 3.31 | -2.05 | 13.71 | -2.15 | 1661 | 182 | 96 | 19.87 | 523 | 318 | 411 | 379 |
| *B. sylvaticum* | Bsyl_80 | 54.00 | 37.20 | 5.16 | 8.12 | 23.14 | 1001.49 | 23.50 | -11.59 | 35.09 | 17.18 | -6.17 | 17.18 | -7.29 | 595 | 82 | 31 | 30.94 | 216 | 100 | 216 | 114 |
| *B. sylvaticum* | Bsyl_81 | 48.20 | 16.30 | 10.09 | 8.41 | 29.86 | 747.40 | 25.45 | -2.71 | 28.16 | 17.73 | 0.93 | 19.34 | 0.93 | 582 | 71 | 30 | 29.32 | 205 | 98 | 201 | 98 |
| *B. sylvaticum* | Bsyl_82 | 48.70 | 21.60 | 8.78 | 9.30 | 30.63 | 796.92 | 24.67 | -5.69 | 30.36 | 18.27 | 0.38 | 18.27 | -1.37 | 661 | 88 | 30 | 36.41 | 247 | 94 | 247 | 103 |
| *B. sylvaticum* | Bsyl_83 | 40.00 | 4.00 | 16.33 | 7.58 | 34.75 | 531.25 | 28.48 | 6.66 | 21.81 | 14.50 | 22.95 | 23.33 | 10.63 | 607 | 89 | 5 | 49.47 | 240 | 46 | 87 | 173 |
| *B. sylvaticum* | Bsyl_84 | 39.60 | 47.00 | 11.92 | 9.45 | 29.53 | 818.27 | 28.98 | -3.01 | 32.00 | 15.65 | 2.08 | 22.04 | 2.08 | 517 | 83 | 22 | 43.74 | 205 | 78 | 108 | 78 |
| *B. sylvaticum* | Bsyl_85 | 49.60 | 12.60 | 7.09 | 8.24 | 31.66 | 686.43 | 21.46 | -4.56 | 26.02 | 15.58 | 2.61 | 15.58 | -1.27 | 720 | 91 | 44 | 24.37 | 245 | 136 | 245 | 158 |
| *B. sylvaticum* | Bsyl_86 | 44.10 | -0.60 | 12.45 | 10.97 | 43.71 | 531.53 | 25.81 | 0.72 | 25.09 | 6.75 | 18.99 | 18.99 | 6.08 | 934 | 114 | 49 | 20.93 | 297 | 185 | 185 | 287 |
| *B. sylvaticum* | Bsyl_87 | 52.00 | 6.20 | 9.69 | 7.96 | 35.01 | 563.15 | 22.54 | -0.20 | 22.74 | 4.04 | 5.59 | 16.74 | 2.99 | 783 | 77 | 46 | 14.42 | 218 | 162 | 204 | 196 |
| *B. sylvaticum* | Bsyl_89 | 51.40 | -2.50 | 10.12 | 7.28 | 37.53 | 448.63 | 21.16 | 1.76 | 19.40 | 6.00 | 14.23 | 15.87 | 5.06 | 815 | 86 | 49 | 18.50 | 250 | 168 | 179 | 229 |
| *B. sylvaticum* | Bsyl_90 | 55.00 | 12.50 | 8.18 | 4.56 | 22.00 | 606.55 | 19.68 | -1.05 | 20.73 | 15.58 | 2.76 | 15.78 | 1.26 | 606 | 60 | 33 | 18.32 | 180 | 113 | 176 | 123 |
| *B. sylvaticum* | Bsyl_91 | 55.80 | 11.40 | 8.26 | 5.04 | 23.23 | 641.86 | 20.35 | -1.34 | 21.69 | 15.79 | 2.97 | 16.26 | 0.74 | 573 | 61 | 29 | 21.92 | 176 | 98 | 171 | 125 |
| *B. sylvaticum* | Bsyl_92 | 54.80 | 11.60 | 8.78 | 4.51 | 21.93 | 627.20 | 20.54 | -0.01 | 20.55 | 16.37 | 3.73 | 16.73 | 1.62 | 587 | 65 | 33 | 20.08 | 175 | 108 | 173 | 131 |
| *B. sylvaticum* | Bsyl_93 | 54.90 | 10.70 | 8.63 | 5.34 | 25.21 | 602.99 | 20.47 | -0.70 | 21.17 | 9.74 | 3.56 | 16.19 | 1.68 | 572 | 63 | 32 | 20.94 | 173 | 101 | 163 | 125 |
| *B. sylvaticum* | Bsyl_94 | 54.50 | 56.60 | 3.10 | 10.55 | 24.18 | 1233.22 | 25.05 | -18.57 | 43.62 | 17.62 | -4.16 | 17.62 | -12.49 | 571 | 69 | 24 | 30.33 | 192 | 86 | 192 | 118 |
| *B. sylvaticum* | Bsyl_95 | 52.80 | 33.50 | 5.75 | 8.28 | 25.14 | 948.12 | 23.00 | -9.96 | 32.96 | 17.13 | -4.77 | 17.13 | -5.98 | 634 | 85 | 32 | 30.90 | 230 | 107 | 230 | 117 |
| *B. sylvaticum* | Bsyl_96 | 53.20 | 34.60 | 5.42 | 7.33 | 23.01 | 937.19 | 21.95 | -9.91 | 31.86 | 16.70 | -4.90 | 16.70 | -6.14 | 644 | 87 | 31 | 31.57 | 234 | 105 | 234 | 117 |
| *B. sylvaticum* | Bsyl_97 | 52.60 | 17.20 | 8.24 | 8.22 | 29.97 | 735.75 | 23.34 | -4.09 | 27.43 | 17.29 | 0.19 | 17.29 | -0.82 | 518 | 75 | 23 | 33.77 | 193 | 85 | 193 | 93 |
| *B. sylvaticum* | Bsyl_98 | 45.20 | 9.10 | 12.33 | 9.27 | 30.75 | 781.32 | 28.11 | -2.02 | 30.14 | 12.76 | 2.46 | 21.76 | 2.46 | 770 | 105 | 39 | 26.12 | 249 | 156 | 163 | 156 |
| *B. sylvaticum* | Bsyl_99 | 46.80 | 12.80 | 7.25 | 10.41 | 32.12 | 808.49 | 23.78 | -8.63 | 32.41 | 16.81 | -3.08 | 16.81 | -3.08 | 947 | 129 | 36 | 37.09 | 347 | 128 | 347 | 128 |
| *B. sylvaticum* | Bsyl_100 | 51.00 | 5.90 | 9.99 | 7.89 | 34.29 | 572.26 | 22.93 | -0.09 | 23.02 | 15.63 | 5.89 | 17.15 | 3.16 | 768 | 80 | 49 | 12.70 | 217 | 166 | 212 | 186 |
| *B. sylvaticum* | Bsyl_102 | 58.20 | -5.30 | 8.30 | 5.33 | 37.79 | 343.20 | 15.84 | 1.73 | 14.11 | 6.63 | 9.31 | 12.57 | 4.25 | 1584 | 191 | 72 | 31.63 | 545 | 243 | 371 | 459 |
| *B. sylvaticum* | Bsyl_103 | 55.90 | -2.30 | 7.45 | 7.78 | 40.60 | 432.32 | 18.43 | -0.74 | 19.17 | 5.12 | 8.95 | 12.98 | 2.44 | 808 | 84 | 47 | 17.40 | 240 | 159 | 196 | 212 |
| *B. sylvaticum* | Bsyl_104 | 44.20 | 0.70 | 12.73 | 10.19 | 40.15 | 570.13 | 26.88 | 1.50 | 25.38 | 6.64 | 19.86 | 19.86 | 6.02 | 912 | 89 | 56 | 14.10 | 255 | 184 | 184 | 246 |
| *B. sylvaticum* | Bsyl_105 | 44.50 | 0.20 | 12.93 | 10.18 | 41.48 | 547.74 | 26.34 | 1.80 | 24.54 | 7.10 | 19.75 | 19.75 | 6.47 | 943 | 99 | 53 | 18.18 | 287 | 179 | 179 | 271 |
| *B. sylvaticum* | Bsyl_106 | 62.40 | 6.50 | 5.84 | 5.05 | 27.42 | 510.92 | 16.04 | -2.38 | 18.42 | 3.29 | 7.84 | 12.40 | -0.01 | 1872 | 229 | 76 | 34.11 | 637 | 269 | 330 | 591 |
| *B. sylvaticum* | Bsyl_107 | 39.40 | 2.80 | 16.42 | 10.66 | 41.81 | 549.48 | 30.54 | 5.05 | 25.49 | 14.38 | 23.32 | 23.59 | 10.47 | 480 | 77 | 6 | 49.59 | 193 | 39 | 74 | 144 |
| *B. sylvaticum* | Bsyl_108 | 39.80 | 3.10 | 16.54 | 9.42 | 39.88 | 523.70 | 29.49 | 5.86 | 23.63 | 14.64 | 23.06 | 23.45 | 10.91 | 707 | 103 | 14 | 45.69 | 270 | 66 | 115 | 209 |
| *B. sylvaticum* | Bsyl_109 | 52.30 | -2.10 | 9.20 | 7.54 | 38.79 | 445.69 | 20.24 | 0.80 | 19.44 | 5.03 | 7.96 | 14.82 | 4.08 | 864 | 88 | 58 | 15.59 | 258 | 184 | 207 | 233 |
| *B. sylvaticum* | Bsyl_110 | 44.70 | 34.40 | 12.31 | 7.92 | 28.10 | 759.70 | 28.03 | -0.16 | 28.19 | 5.28 | 10.48 | 22.06 | 3.66 | 549 | 68 | 36 | 18.81 | 168 | 117 | 139 | 163 |
| *B. sylvaticum* | Bsyl_111 | 55.10 | 58.40 | 1.59 | 9.08 | 22.94 | 1133.87 | 21.89 | -17.70 | 39.59 | 15.20 | -10.99 | 15.20 | -12.46 | 594 | 93 | 24 | 42.90 | 237 | 80 | 237 | 90 |
| *B. sylvaticum* | Bsyl_112 | 45.40 | 40.80 | 10.74 | 9.97 | 28.83 | 925.04 | 28.99 | -5.60 | 34.59 | 19.94 | 0.42 | 21.86 | -0.88 | 614 | 77 | 34 | 22.80 | 201 | 115 | 191 | 132 |
| *B. sylvaticum* | Bsyl_113 | 55.50 | 31.70 | 4.96 | 7.85 | 24.63 | 906.51 | 21.70 | -10.17 | 31.88 | 15.98 | -0.79 | 15.98 | -6.26 | 695 | 91 | 35 | 31.17 | 247 | 113 | 247 | 132 |
| *B. sylvaticum* | Bsyl_114 | 55.90 | 37.50 | 5.04 | 7.64 | 22.77 | 975.87 | 22.88 | -10.66 | 33.54 | 17.00 | -0.90 | 17.00 | -6.98 | 668 | 87 | 29 | 32.96 | 243 | 99 | 243 | 129 |
| *B. sylvaticum* | Bsyl_115 | 52.70 | 41.50 | 5.52 | 8.53 | 23.35 | 1068.68 | 24.70 | -11.83 | 36.53 | 18.30 | -1.21 | 18.30 | -7.62 | 535 | 70 | 28 | 27.28 | 172 | 88 | 172 | 113 |
| *B. sylvaticum* | Bsyl_116 | 56.00 | 45.90 | 3.72 | 7.85 | 20.49 | 1121.61 | 23.20 | -15.10 | 38.30 | 17.05 | -8.89 | 17.05 | -10.40 | 572 | 74 | 27 | 29.22 | 199 | 93 | 199 | 110 |
| *B. sylvaticum* | Bsyl_117 | 49.40 | 10.20 | 8.41 | 8.53 | 32.71 | 675.12 | 23.01 | -3.06 | 26.07 | 15.23 | 1.35 | 16.81 | 0.24 | 689 | 74 | 48 | 14.01 | 206 | 150 | 200 | 159 |
| *B. sylvaticum* | Bsyl_118 | 50.40 | -4.10 | 10.67 | 6.30 | 37.34 | 395.52 | 20.03 | 3.16 | 16.87 | 7.19 | 14.15 | 15.69 | 6.27 | 999 | 123 | 54 | 28.65 | 348 | 175 | 182 | 326 |
| *B. sylvaticum* | Bsyl_119 | 51.70 | 0.60 | 9.84 | 7.65 | 37.01 | 483.35 | 21.74 | 1.08 | 20.66 | 10.58 | 6.03 | 15.99 | 4.36 | 566 | 57 | 34 | 15.46 | 164 | 115 | 142 | 137 |
| *B. sylvaticum* | Bsyl_120 | 52.10 | -4.50 | 9.52 | 5.45 | 34.92 | 376.26 | 18.12 | 2.51 | 15.62 | 8.14 | 10.23 | 14.33 | 5.32 | 1181 | 133 | 68 | 23.65 | 386 | 217 | 274 | 337 |
| *B. sylvaticum* | Bsyl_121 | 53.20 | 0.20 | 9.50 | 6.81 | 35.20 | 468.59 | 20.46 | 1.10 | 19.36 | 5.07 | 5.80 | 15.42 | 4.13 | 610 | 60 | 39 | 13.05 | 176 | 130 | 152 | 156 |
| *B. sylvaticum* | Bsyl_122 | 51.20 | -3.50 | 10.04 | 6.39 | 36.30 | 417.11 | 19.96 | 2.37 | 17.59 | 6.41 | 11.06 | 15.38 | 5.42 | 1070 | 124 | 62 | 24.65 | 354 | 189 | 221 | 312 |
| *B. sylvaticum* | Bsyl_123 | 51.20 | 1.40 | 10.28 | 6.24 | 32.30 | 474.69 | 21.22 | 1.90 | 19.31 | 11.34 | 6.39 | 16.35 | 4.88 | 570 | 67 | 35 | 23.07 | 190 | 113 | 140 | 141 |
| *B. sylvaticum* | Bsyl_124 | 54.40 | -1.80 | 8.16 | 6.93 | 37.62 | 428.69 | 18.78 | 0.36 | 18.42 | 4.14 | 9.39 | 13.70 | 3.36 | 1008 | 103 | 67 | 16.42 | 305 | 207 | 231 | 275 |
| *B. sylvaticum* | Bsyl_125 | 54.20 | -3.10 | 8.76 | 6.42 | 36.68 | 431.41 | 18.39 | 0.88 | 17.51 | 6.53 | 10.27 | 14.22 | 3.74 | 1389 | 158 | 76 | 26.61 | 466 | 239 | 281 | 402 |
| *B. sylvaticum* | Bsyl_126 | 51.00 | -4.50 | 9.94 | 5.94 | 36.82 | 380.10 | 18.77 | 2.62 | 16.14 | 6.81 | 10.57 | 14.81 | 5.74 | 1076 | 128 | 59 | 27.42 | 370 | 188 | 238 | 289 |
| *B. sylvaticum* | Bsyl_127 | 50.10 | -5.40 | 10.84 | 5.44 | 37.46 | 338.58 | 19.04 | 4.51 | 14.53 | 8.12 | 13.52 | 15.28 | 7.16 | 1057 | 128 | 52 | 30.39 | 372 | 173 | 211 | 315 |
| *B. sylvaticum* | Bsyl_128 | 50.60 | -1.20 | 10.01 | 6.76 | 36.64 | 437.52 | 20.41 | 1.94 | 18.46 | 6.07 | 13.88 | 15.54 | 5.10 | 894 | 106 | 45 | 30.36 | 313 | 151 | 158 | 278 |
| *B. sylvaticum* | Bsyl_129 | 51.70 | -0.80 | 9.58 | 7.52 | 36.29 | 482.90 | 21.59 | 0.86 | 20.72 | 7.08 | 5.83 | 15.81 | 4.12 | 742 | 76 | 45 | 15.34 | 220 | 163 | 164 | 199 |
| *B. sylvaticum* | Bsyl_130 | 55.20 | -6.40 | 9.12 | 5.90 | 38.47 | 364.64 | 17.61 | 2.26 | 15.35 | 5.77 | 10.08 | 13.72 | 5.01 | 1206 | 136 | 62 | 26.24 | 394 | 200 | 280 | 362 |
| *B. sylvaticum* | Bsyl_131 | 58.40 | 25.60 | 5.34 | 8.08 | 25.07 | 854.90 | 22.95 | -9.29 | 32.24 | 14.73 | -1.12 | 15.89 | -4.99 | 648 | 77 | 32 | 30.86 | 218 | 98 | 205 | 130 |
| *B. sylvaticum* | Bsyl_132 | 58.90 | 26.80 | 4.96 | 8.16 | 25.33 | 853.74 | 23.02 | -9.20 | 32.22 | 14.44 | -4.96 | 15.72 | -5.17 | 615 | 81 | 27 | 35.47 | 225 | 90 | 221 | 106 |
| *B. sylvaticum* | Bsyl_133 | 57.60 | 26.60 | 5.58 | 7.97 | 26.40 | 828.77 | 21.86 | -8.32 | 30.18 | 15.90 | -0.28 | 15.90 | -4.51 | 651 | 80 | 30 | 31.11 | 226 | 100 | 226 | 118 |
| *B. sylvaticum* | Bsyl_134 | 43.80 | -1.40 | 13.76 | 8.57 | 41.75 | 446.24 | 24.57 | 4.05 | 20.52 | 9.33 | 17.51 | 19.42 | 8.60 | 1207 | 146 | 54 | 27.68 | 409 | 198 | 253 | 370 |
| *B. sylvaticum* | Bsyl_135 | 55.50 | 45.30 | 3.79 | 8.47 | 21.19 | 1157.97 | 23.76 | -16.20 | 39.96 | 17.32 | -9.61 | 17.32 | -10.98 | 544 | 75 | 25 | 31.59 | 192 | 86 | 192 | 101 |
| *B. sylvaticum* | Bsyl_136 | 41.90 | 44.80 | 10.65 | 10.14 | 32.40 | 794.58 | 27.02 | -4.28 | 31.30 | 14.31 | 0.80 | 20.15 | 0.80 | 659 | 100 | 25 | 42.70 | 264 | 89 | 214 | 89 |
| *B. sylvaticum* | Bsyl_137 | 48.70 | 40.40 | 7.83 | 9.21 | 25.55 | 1033.78 | 27.38 | -8.66 | 36.04 | 18.60 | -3.73 | 20.36 | -4.87 | 483 | 58 | 26 | 24.16 | 158 | 94 | 150 | 114 |
| *B. sylvaticum* | Bsyl_138 | 45.30 | 4.90 | 11.98 | 9.38 | 34.70 | 649.11 | 27.11 | 0.08 | 27.03 | 12.28 | 4.26 | 20.17 | 4.26 | 846 | 100 | 51 | 23.09 | 268 | 162 | 193 | 162 |
| *B. sylvaticum* | Bsyl_139 | 45.70 | 5.50 | 11.49 | 9.73 | 34.80 | 673.34 | 26.90 | -1.04 | 27.95 | 11.68 | 4.67 | 19.91 | 3.29 | 1035 | 112 | 70 | 17.42 | 316 | 220 | 237 | 222 |
| *B. sylvaticum* | Bsyl_140 | 54.90 | 38.30 | 4.96 | 8.19 | 23.19 | 1016.28 | 23.41 | -11.88 | 35.29 | 17.29 | -6.56 | 17.29 | -7.54 | 589 | 83 | 28 | 34.02 | 216 | 91 | 216 | 105 |
| *B. sylvaticum* | Bsyl_141 | 33.30 | -5.40 | 14.37 | 12.72 | 41.23 | 631.86 | 31.78 | 0.93 | 30.85 | 10.06 | 22.36 | 22.62 | 7.05 | 451 | 57 | 4 | 52.62 | 168 | 28 | 28 | 154 |
| *B. sylvaticum* | Bsyl_142 | 47.50 | 7.80 | 8.90 | 8.83 | 34.08 | 643.70 | 23.32 | -2.58 | 25.90 | 15.34 | 4.77 | 16.96 | 1.16 | 1225 | 122 | 83 | 11.15 | 346 | 270 | 338 | 301 |
| *B. sylvaticum* | Bsyl_143 | 46.10 | 7.20 | 5.38 | 8.37 | 33.92 | 616.16 | 19.10 | -5.58 | 24.68 | -0.62 | 13.00 | 13.11 | -1.67 | 1265 | 131 | 83 | 15.02 | 374 | 265 | 286 | 373 |
| *B. sylvaticum* | Bsyl_144 | 46.40 | 8.50 | 4.15 | 7.87 | 33.22 | 592.01 | 17.28 | -6.40 | 23.68 | 11.67 | -2.52 | 11.67 | -2.52 | 1356 | 143 | 72 | 22.83 | 417 | 230 | 417 | 230 |
| *B. sylvaticum* | Bsyl_145 | 47.10 | 8.60 | 6.87 | 7.47 | 32.11 | 601.70 | 19.63 | -3.64 | 23.27 | 14.40 | 0.55 | 14.40 | -0.18 | 1559 | 181 | 93 | 23.28 | 530 | 298 | 530 | 308 |
| *B. sylvaticum* | Bsyl_146 | 46.00 | 9.00 | 9.06 | 8.55 | 33.12 | 638.18 | 23.10 | -2.71 | 25.80 | 13.82 | 1.58 | 17.19 | 1.58 | 1330 | 149 | 58 | 29.64 | 419 | 197 | 399 | 197 |
| *B. sylvaticum* | Bsyl_147 | 46.20 | 7.90 | 3.35 | 7.93 | 32.71 | 605.43 | 16.86 | -7.38 | 24.24 | 5.17 | -3.44 | 11.03 | -3.44 | 856 | 86 | 61 | 11.75 | 235 | 187 | 218 | 187 |
| *B. sylvaticum* | Bsyl_148 | 47.50 | 9.40 | 8.98 | 7.76 | 30.45 | 671.60 | 23.00 | -2.49 | 25.50 | 17.31 | 2.05 | 17.31 | 0.82 | 1152 | 143 | 59 | 30.07 | 413 | 192 | 413 | 199 |
| *B. sylvaticum* | Bsyl_149 | 47.30 | 7.20 | 7.34 | 7.92 | 32.58 | 620.89 | 20.84 | -3.47 | 24.31 | 13.25 | 0.85 | 15.06 | -0.08 | 1174 | 119 | 79 | 11.70 | 335 | 255 | 322 | 281 |
| *B. sylvaticum* | Bsyl_150 | 46.70 | 7.20 | 5.11 | 7.65 | 32.56 | 596.67 | 18.13 | -5.38 | 23.50 | -1.68 | 3.52 | 12.59 | -1.68 | 1388 | 139 | 97 | 11.45 | 388 | 307 | 356 | 388 |
| *B. sylvaticum* | Bsyl_151 | 57.30 | 15.50 | 5.98 | 8.75 | 31.50 | 695.37 | 21.77 | -6.01 | 27.78 | 13.81 | 0.64 | 14.95 | -2.10 | 617 | 67 | 36 | 20.44 | 192 | 114 | 187 | 135 |
| *B. sylvaticum* | Bsyl_152 | 59.00 | 16.30 | 6.07 | 8.56 | 29.85 | 733.21 | 22.19 | -6.49 | 28.68 | 14.22 | -1.90 | 15.54 | -2.51 | 592 | 71 | 30 | 27.82 | 201 | 102 | 190 | 121 |
| *B. sylvaticum* | Bsyl_153 | 52.50 | 6.00 | 9.68 | 7.92 | 35.47 | 556.94 | 22.38 | 0.06 | 22.32 | 10.14 | 5.83 | 16.60 | 2.93 | 808 | 78 | 45 | 16.26 | 228 | 159 | 212 | 195 |
| *B. sylvaticum* | Bsyl_154 | 52.50 | 4.60 | 9.88 | 6.32 | 31.37 | 510.24 | 20.94 | 0.80 | 20.15 | 10.96 | 11.52 | 16.13 | 3.90 | 854 | 102 | 46 | 26.88 | 295 | 159 | 194 | 204 |
| *B. sylvaticum* | Bsyl_155 | 49.70 | 6.40 | 9.38 | 8.03 | 33.15 | 621.38 | 23.20 | -1.02 | 24.22 | 5.62 | 5.27 | 17.18 | 1.94 | 794 | 80 | 53 | 11.37 | 224 | 171 | 200 | 203 |
| *B. sylvaticum* | Bsyl_156 | 45.40 | 3.20 | 9.58 | 9.82 | 38.10 | 581.44 | 24.20 | -1.59 | 25.78 | 14.96 | 3.67 | 16.96 | 2.84 | 669 | 82 | 36 | 26.54 | 206 | 114 | 193 | 123 |
| *B. sylvaticum* | Bsyl_157 | 47.90 | -4.20 | 12.03 | 6.01 | 35.62 | 401.83 | 21.58 | 4.70 | 16.88 | 8.38 | 17.05 | 17.13 | 7.48 | 1028 | 133 | 45 | 34.11 | 382 | 156 | 182 | 347 |
| *B. sylvaticum* | Bsyl_158 | 43.20 | 0.50 | 11.85 | 10.10 | 41.06 | 538.09 | 25.17 | 0.58 | 24.59 | 13.32 | 18.69 | 18.69 | 5.72 | 1123 | 118 | 69 | 13.83 | 322 | 227 | 227 | 284 |
| *B. sylvaticum* | Bsyl_159 | 48.00 | 5.40 | 9.28 | 7.44 | 31.04 | 628.03 | 22.70 | -1.29 | 23.99 | 2.60 | 5.14 | 17.11 | 1.78 | 859 | 83 | 55 | 11.80 | 240 | 187 | 209 | 234 |
| *B. sylvaticum* | Bsyl_160 | 47.50 | -0.30 | 11.83 | 9.19 | 39.49 | 535.11 | 25.32 | 2.05 | 23.27 | 6.32 | 18.60 | 18.60 | 5.58 | 704 | 74 | 41 | 19.78 | 219 | 130 | 130 | 208 |
| *B. sylvaticum* | Bsyl_161 | 47.80 | -1.50 | 11.36 | 8.31 | 38.52 | 500.00 | 23.84 | 2.28 | 21.56 | 8.71 | 17.65 | 17.65 | 5.56 | 761 | 90 | 42 | 21.28 | 240 | 143 | 143 | 215 |
| *B. sylvaticum* | Bsyl_162 | 49.20 | 2.10 | 10.49 | 7.95 | 35.15 | 545.37 | 23.55 | 0.94 | 22.60 | 7.48 | 6.68 | 17.31 | 4.00 | 650 | 64 | 45 | 10.93 | 180 | 144 | 154 | 166 |
| *B. sylvaticum* | Bsyl_163 | 46.10 | 6.00 | 9.29 | 9.25 | 34.16 | 657.12 | 24.60 | -2.49 | 27.09 | 9.58 | 8.63 | 17.54 | 1.40 | 1074 | 105 | 73 | 13.17 | 311 | 234 | 252 | 277 |
| *B. sylvaticum* | Bsyl_164 | 46.70 | -1.00 | 11.95 | 8.94 | 39.13 | 531.51 | 25.08 | 2.22 | 22.86 | 6.48 | 18.59 | 18.59 | 5.66 | 768 | 90 | 39 | 26.44 | 254 | 126 | 126 | 234 |
| *B. sylvaticum* | Bsyl_165 | 48.30 | -1.50 | 11.21 | 7.99 | 38.50 | 483.98 | 23.07 | 2.32 | 20.75 | 8.74 | 17.27 | 17.27 | 5.58 | 725 | 82 | 47 | 19.52 | 229 | 141 | 141 | 198 |
| *B. sylvaticum* | Bsyl_166 | 48.30 | 2.80 | 10.74 | 8.76 | 35.91 | 578.50 | 24.78 | 0.38 | 24.39 | 16.25 | 6.75 | 18.02 | 3.84 | 631 | 59 | 46 | 8.88 | 170 | 141 | 157 | 154 |
| *B. sylvaticum* | Bsyl_167 | 44.90 | 1.80 | 12.54 | 10.23 | 39.74 | 577.80 | 27.04 | 1.28 | 25.75 | 14.87 | 6.72 | 19.81 | 5.68 | 761 | 79 | 50 | 12.89 | 213 | 173 | 175 | 184 |
| *B. sylvaticum* | Bsyl_168 | 50.30 | 1.70 | 10.84 | 6.38 | 32.22 | 499.33 | 21.79 | 2.00 | 19.80 | 8.35 | 7.09 | 16.95 | 4.86 | 673 | 81 | 44 | 21.34 | 218 | 135 | 149 | 171 |
| *B. sylvaticum* | Bsyl_169 | 47.40 | 3.70 | 10.16 | 8.84 | 35.12 | 606.38 | 24.50 | -0.67 | 25.16 | 6.77 | 6.01 | 17.80 | 2.87 | 788 | 75 | 54 | 10.82 | 211 | 171 | 198 | 199 |
| *B. sylvaticum* | Bsyl_170 | 47.10 | -1.40 | 11.75 | 8.71 | 39.47 | 510.06 | 24.39 | 2.32 | 22.07 | 6.54 | 18.16 | 18.16 | 5.74 | 777 | 87 | 42 | 25.72 | 256 | 128 | 128 | 244 |
| *B. sylvaticum* | Bsyl_171 | 45.30 | 1.40 | 12.08 | 9.19 | 38.47 | 552.19 | 25.50 | 1.62 | 23.88 | 8.89 | 19.04 | 19.04 | 5.64 | 948 | 92 | 58 | 13.61 | 268 | 196 | 196 | 250 |
| *B. sylvaticum* | Bsyl_172 | 48.90 | 1.40 | 10.28 | 8.05 | 35.73 | 534.48 | 23.31 | 0.80 | 22.52 | 7.39 | 9.34 | 16.99 | 3.97 | 623 | 70 | 42 | 13.90 | 184 | 139 | 150 | 164 |
| *B. sylvaticum* | Bsyl_173 | 49.60 | 2.90 | 10.04 | 7.97 | 35.47 | 542.33 | 23.00 | 0.52 | 22.48 | 15.16 | 6.20 | 16.82 | 3.61 | 651 | 63 | 46 | 10.10 | 176 | 142 | 167 | 160 |
| *B. sylvaticum* | Bsyl_174 | 43.80 | 5.40 | 11.61 | 10.01 | 36.41 | 633.04 | 27.23 | -0.27 | 27.50 | 12.20 | 19.75 | 19.75 | 4.34 | 849 | 114 | 29 | 29.26 | 283 | 139 | 139 | 215 |
| *B. sylvaticum* | Bsyl_175 | 49.00 | -1.40 | 11.15 | 7.00 | 37.05 | 447.46 | 21.70 | 2.80 | 18.90 | 9.16 | 16.68 | 16.73 | 5.92 | 797 | 94 | 44 | 25.21 | 266 | 138 | 156 | 234 |
| *B. sylvaticum* | Bsyl_176 | 45.90 | 4.70 | 11.12 | 8.66 | 33.34 | 641.15 | 25.76 | -0.23 | 25.99 | 11.33 | 4.71 | 19.24 | 3.52 | 823 | 86 | 51 | 18.16 | 241 | 159 | 216 | 164 |
| *B. sylvaticum* | Bsyl_177 | 46.40 | 2.30 | 10.43 | 8.94 | 36.98 | 574.21 | 24.32 | 0.14 | 24.17 | 14.97 | 6.47 | 17.69 | 3.68 | 741 | 75 | 45 | 17.11 | 214 | 145 | 208 | 160 |
| *B. sylvaticum* | Bsyl_178 | 48.50 | 4.70 | 10.67 | 9.13 | 36.69 | 595.96 | 24.91 | 0.01 | 24.90 | 4.43 | 6.65 | 18.21 | 3.60 | 764 | 73 | 52 | 9.54 | 208 | 165 | 193 | 201 |
| *B. sylvaticum* | Bsyl_179 | 42.90 | -0.10 | 7.90 | 9.44 | 36.26 | 598.53 | 23.04 | -3.00 | 26.04 | 5.04 | 15.65 | 15.65 | 1.11 | 1076 | 110 | 59 | 17.70 | 314 | 207 | 207 | 291 |
| *B. sylvaticum* | Bsyl_180 | 46.50 | 4.60 | 10.53 | 8.59 | 33.48 | 635.82 | 25.15 | -0.51 | 25.66 | 10.76 | 6.47 | 18.56 | 2.95 | 831 | 83 | 51 | 14.40 | 233 | 170 | 217 | 189 |
| *B. sylvaticum* | Bsyl_181 | 46.10 | 1.00 | 11.35 | 8.59 | 37.26 | 548.40 | 24.45 | 1.40 | 23.06 | 8.25 | 18.27 | 18.27 | 4.99 | 888 | 87 | 57 | 13.81 | 254 | 181 | 181 | 244 |
| *B. sylvaticum* | Bsyl_182 | 50.70 | 2.70 | 10.21 | 7.43 | 34.49 | 516.10 | 22.38 | 0.85 | 21.53 | 11.00 | 6.36 | 16.55 | 4.05 | 696 | 73 | 42 | 15.50 | 208 | 141 | 172 | 162 |
| *B. sylvaticum* | Bsyl_183 | 42.60 | 2.10 | 6.38 | 8.98 | 35.27 | 599.60 | 20.94 | -4.51 | 25.45 | 3.60 | 14.15 | 14.15 | -0.30 | 1123 | 119 | 61 | 17.77 | 335 | 226 | 226 | 284 |
| *B. sylvaticum* | Bsyl_184 | 46.80 | -1.90 | 12.52 | 7.88 | 37.79 | 488.35 | 24.39 | 3.53 | 20.86 | 7.67 | 18.61 | 18.61 | 6.77 | 770 | 89 | 38 | 28.94 | 262 | 119 | 119 | 246 |
| *B. sylvaticum* | Bsyl_185 | 43.00 | 2.90 | 13.39 | 7.85 | 33.38 | 567.24 | 26.50 | 2.99 | 23.51 | 10.43 | 20.69 | 20.69 | 6.99 | 657 | 84 | 19 | 30.27 | 216 | 99 | 99 | 192 |
| *B. sylvaticum* | Bsyl_186 | 47.30 | 0.40 | 11.46 | 8.77 | 38.49 | 537.51 | 24.66 | 1.86 | 22.79 | 5.89 | 18.24 | 18.24 | 5.16 | 666 | 69 | 39 | 16.63 | 198 | 131 | 131 | 192 |
| *B. sylvaticum* | Bsyl_187 | 48.10 | -2.20 | 11.52 | 6.91 | 36.51 | 450.40 | 22.39 | 3.48 | 18.92 | 7.20 | 17.11 | 17.21 | 6.33 | 821 | 96 | 46 | 24.04 | 271 | 144 | 160 | 245 |
| *B. sylvaticum* | Bsyl_188 | 43.60 | 1.30 | 12.84 | 9.73 | 38.43 | 577.32 | 27.01 | 1.70 | 25.31 | 14.68 | 20.06 | 20.15 | 6.21 | 704 | 80 | 46 | 15.28 | 216 | 152 | 164 | 166 |
| *B. sylvaticum* | Bsyl_189 | 43.40 | 3.40 | 15.09 | 8.97 | 35.79 | 592.68 | 29.00 | 3.95 | 25.05 | 11.74 | 22.72 | 22.72 | 8.30 | 584 | 83 | 17 | 33.02 | 195 | 86 | 86 | 168 |
| *B. sylvaticum* | Bsyl_190 | 44.10 | 1.70 | 12.22 | 10.19 | 38.82 | 589.66 | 26.95 | 0.71 | 26.24 | 14.41 | 19.38 | 19.68 | 5.29 | 770 | 83 | 48 | 13.49 | 219 | 167 | 169 | 196 |
| *B. sylvaticum* | Bsyl_191 | 39.50 | -0.40 | 17.59 | 9.44 | 40.46 | 521.75 | 29.90 | 6.56 | 23.34 | 15.10 | 24.11 | 24.49 | 11.79 | 446 | 75 | 10 | 45.26 | 176 | 52 | 78 | 118 |
| *B. sylvaticum* | Bsyl_192 | 48.00 | 0.30 | 11.19 | 8.57 | 37.99 | 532.40 | 24.26 | 1.70 | 22.56 | 5.71 | 17.90 | 17.90 | 4.94 | 671 | 68 | 41 | 15.36 | 199 | 134 | 134 | 193 |
| *B. sylvaticum* | Bsyl_193 | 38.50 | 25.90 | 16.52 | 8.09 | 33.50 | 623.62 | 29.91 | 5.76 | 24.15 | 9.63 | 24.33 | 24.36 | 9.45 | 526 | 107 | 3 | 83.79 | 285 | 14 | 21 | 240 |
| *B. sylvaticum* | Bsyl_194 | 40.10 | 22.50 | 14.16 | 9.96 | 33.09 | 761.85 | 30.76 | 0.67 | 30.09 | 9.99 | 23.04 | 23.74 | 5.02 | 444 | 56 | 22 | 27.53 | 148 | 73 | 75 | 113 |
| *B. sylvaticum* | Bsyl_195 | 48.40 | 17.50 | 9.65 | 9.60 | 31.83 | 774.51 | 26.14 | -4.02 | 30.16 | 19.06 | 1.52 | 19.06 | 0.00 | 648 | 81 | 32 | 32.32 | 236 | 102 | 236 | 113 |
| *B. sylvaticum* | Bsyl_196 | 37.50 | 44.70 | 3.88 | 9.69 | 26.93 | 943.07 | 23.61 | -12.37 | 35.98 | 1.29 | 15.60 | 15.60 | -7.36 | 581 | 107 | 4 | 68.56 | 265 | 17 | 17 | 170 |
| *B. sylvaticum* | Bsyl_197 | 41.50 | 36.00 | 13.26 | 7.41 | 31.05 | 627.31 | 26.08 | 2.20 | 23.88 | 10.98 | 21.20 | 21.20 | 6.09 | 716 | 87 | 32 | 28.98 | 254 | 121 | 121 | 178 |
| *B. sylvaticum* | Bsyl_198 | 36.70 | 24.50 | 17.96 | 5.68 | 28.11 | 557.21 | 29.03 | 8.82 | 20.21 | 13.15 | 24.96 | 24.96 | 11.58 | 428 | 76 | 1 | 82.97 | 219 | 7 | 7 | 189 |
| *B. sylvaticum* | Bsyl_199 | 36.50 | 22.40 | 17.44 | 7.33 | 32.48 | 588.62 | 29.87 | 7.30 | 22.57 | 12.34 | 24.87 | 24.87 | 10.84 | 612 | 115 | 3 | 81.90 | 320 | 13 | 13 | 295 |
| *B. sylvaticum* | Bsyl_200 | 39.20 | 26.30 | 16.16 | 8.72 | 32.97 | 669.20 | 30.74 | 4.29 | 26.45 | 9.98 | 24.64 | 24.64 | 8.47 | 593 | 119 | 4 | 80.17 | 306 | 19 | 19 | 305 |
| *B. sylvaticum* | Bsyl_201 | 35.40 | 23.90 | 15.79 | 7.24 | 32.73 | 571.11 | 27.96 | 5.83 | 22.13 | 9.17 | 22.92 | 22.92 | 9.17 | 695 | 139 | 3 | 83.87 | 362 | 11 | 11 | 362 |
| *B. sylvaticum* | Bsyl_202 | 37.60 | 26.00 | 16.07 | 6.65 | 29.70 | 604.74 | 28.40 | 6.00 | 22.40 | 9.18 | 23.72 | 23.72 | 9.18 | 577 | 114 | 2 | 86.02 | 313 | 9 | 9 | 313 |
| *B. sylvaticum* | Bsyl_203 | 37.20 | 24.50 | 18.03 | 4.84 | 24.02 | 578.35 | 29.07 | 8.91 | 20.16 | 13.10 | 25.38 | 25.38 | 11.44 | 416 | 73 | 2 | 79.59 | 206 | 10 | 10 | 180 |
| *B. sylvaticum* | Bsyl_206 | 37.60 | 25.00 | 17.63 | 4.52 | 23.16 | 570.40 | 27.90 | 8.40 | 19.50 | 11.18 | 24.88 | 24.88 | 11.13 | 440 | 77 | 3 | 78.73 | 217 | 12 | 12 | 196 |
| *B. sylvaticum* | Bsyl_207 | 37.70 | 26.60 | 17.33 | 8.10 | 32.08 | 652.18 | 31.38 | 6.13 | 25.24 | 9.96 | 25.69 | 25.69 | 9.96 | 649 | 141 | 2 | 91.34 | 367 | 9 | 9 | 367 |
| *B. sylvaticum* | Bsyl_208 | 37.10 | 25.50 | 15.42 | 5.88 | 28.03 | 579.87 | 26.96 | 5.99 | 20.97 | 8.86 | 22.77 | 22.77 | 8.86 | 598 | 108 | 3 | 80.37 | 305 | 14 | 14 | 305 |
| *B. sylvaticum* | Bsyl_209 | 41.40 | 26.20 | 12.73 | 9.59 | 32.78 | 738.07 | 28.69 | -0.57 | 29.26 | 9.05 | 21.35 | 21.89 | 3.77 | 594 | 76 | 23 | 33.38 | 206 | 81 | 92 | 187 |
| *B. sylvaticum* | Bsyl_210 | 38.00 | 21.90 | 10.27 | 9.73 | 36.19 | 649.30 | 25.82 | -1.08 | 26.90 | 4.31 | 18.46 | 18.46 | 2.78 | 907 | 138 | 23 | 56.13 | 385 | 74 | 74 | 367 |
| *B. sylvaticum* | Bsyl_211 | 39.40 | 20.80 | 12.42 | 10.23 | 37.04 | 646.05 | 28.22 | 0.61 | 27.61 | 6.24 | 20.58 | 20.58 | 4.94 | 991 | 148 | 27 | 51.25 | 405 | 89 | 89 | 367 |
| *B. sylvaticum* | Bsyl_212 | 35.20 | 26.10 | 18.82 | 6.79 | 32.10 | 547.10 | 30.30 | 9.16 | 21.14 | 12.77 | 25.68 | 25.73 | 12.66 | 552 | 117 | 1 | 86.59 | 301 | 6 | 19 | 268 |
| *B. sylvaticum* | Bsyl_214 | 56.40 | 15.40 | 6.74 | 7.78 | 30.58 | 658.06 | 21.29 | -4.16 | 25.45 | 7.22 | 1.46 | 15.23 | -0.82 | 714 | 76 | 39 | 19.64 | 216 | 135 | 187 | 175 |
| *B. sylvaticum* | Bsyl_215 | 38.00 | 24.70 | 17.67 | 4.90 | 24.48 | 576.84 | 28.35 | 8.35 | 20.00 | 11.21 | 25.01 | 25.01 | 11.13 | 413 | 72 | 4 | 75.11 | 198 | 15 | 15 | 178 |
| *B. sylvaticum* | Bsyl_216 | 50.00 | 14.60 | 8.54 | 8.44 | 30.79 | 713.14 | 23.76 | -3.64 | 27.40 | 15.75 | -0.20 | 17.29 | -0.20 | 509 | 74 | 21 | 46.42 | 210 | 71 | 209 | 71 |
| *B. sylvaticum* | Bsyl_217 | 53.20 | 6.60 | 8.98 | 7.61 | 34.13 | 554.90 | 21.59 | -0.70 | 22.29 | 6.16 | 4.78 | 15.87 | 2.36 | 799 | 84 | 45 | 18.29 | 234 | 152 | 214 | 192 |
| *B. sylvaticum* | Bsyl_218 | 47.50 | 10.10 | 4.93 | 7.93 | 32.80 | 615.87 | 17.80 | -6.38 | 24.19 | 12.50 | -1.70 | 12.50 | -2.32 | 1568 | 190 | 100 | 24.46 | 543 | 312 | 543 | 318 |
| *B. sylvaticum* | Bsyl_219 | 48.00 | 10.20 | 8.00 | 8.92 | 32.96 | 689.04 | 22.86 | -4.21 | 27.06 | 16.61 | 0.90 | 16.61 | -0.31 | 1038 | 140 | 48 | 34.59 | 391 | 164 | 391 | 165 |
| *B. sylvaticum* | Bsyl_220 | 49.90 | 10.00 | 8.87 | 8.48 | 32.75 | 666.35 | 23.36 | -2.53 | 25.90 | 15.65 | 4.59 | 17.17 | 0.80 | 649 | 69 | 44 | 14.49 | 191 | 137 | 186 | 157 |
| *B. sylvaticum* | Bsyl_221 | 48.60 | 10.40 | 8.39 | 8.52 | 32.01 | 693.61 | 23.14 | -3.48 | 26.63 | 15.47 | 1.18 | 16.98 | -0.09 | 769 | 93 | 45 | 25.92 | 264 | 137 | 259 | 150 |
| *B. sylvaticum* | Bsyl_222 | 49.60 | 11.20 | 8.40 | 9.36 | 34.52 | 686.35 | 23.51 | -3.60 | 27.11 | 15.48 | 3.97 | 16.96 | 0.11 | 780 | 86 | 50 | 16.25 | 235 | 162 | 232 | 183 |
| *B. sylvaticum* | Bsyl_223 | 47.40 | 11.20 | 5.46 | 7.91 | 32.92 | 611.78 | 18.50 | -5.53 | 24.03 | 13.00 | -1.77 | 13.00 | -1.77 | 1277 | 175 | 61 | 37.97 | 503 | 199 | 503 | 199 |
| *B. sylvaticum* | Bsyl_224 | 47.20 | 4.40 | 9.58 | 8.49 | 33.82 | 619.16 | 23.98 | -1.11 | 25.09 | 6.07 | 5.41 | 17.37 | 2.17 | 854 | 86 | 56 | 12.56 | 230 | 181 | 213 | 209 |
| *B. sylvaticum* | Bsyl_225 | 45.90 | 1.90 | 9.73 | 7.93 | 35.03 | 556.66 | 22.55 | -0.08 | 22.63 | 10.16 | 5.84 | 16.76 | 3.20 | 994 | 98 | 67 | 11.55 | 273 | 224 | 228 | 249 |
| *B. sylvaticum* | Bsyl_227 | 45.40 | 2.40 | 10.02 | 9.61 | 38.66 | 570.27 | 23.88 | -0.99 | 24.87 | 12.25 | 4.18 | 17.19 | 3.23 | 779 | 85 | 47 | 17.06 | 224 | 157 | 201 | 170 |
| *B. sylvaticum* | Bsyl_228 | 45.00 | 3.60 | 8.09 | 9.28 | 36.65 | 584.82 | 22.66 | -2.66 | 25.32 | 12.93 | 2.09 | 15.53 | 1.41 | 799 | 91 | 48 | 21.02 | 238 | 154 | 192 | 166 |
| *B. sylvaticum* | Bsyl_229 | 43.40 | 2.30 | 10.42 | 8.43 | 35.11 | 568.13 | 23.93 | -0.07 | 24.00 | 7.51 | 17.70 | 17.70 | 4.02 | 922 | 95 | 41 | 20.44 | 273 | 169 | 169 | 252 |
| *B. sylvaticum* | Bsyl_230 | 56.80 | -3.90 | 5.99 | 7.01 | 37.55 | 440.07 | 16.59 | -2.08 | 18.68 | 1.80 | 7.62 | 11.68 | 1.02 | 1421 | 183 | 73 | 29.97 | 483 | 237 | 256 | 451 |
| *B. sylvaticum* | Bsyl_231 | 57.30 | -3.50 | 5.83 | 7.11 | 38.45 | 422.53 | 16.31 | -2.19 | 18.50 | 1.93 | 7.23 | 11.36 | 1.17 | 1177 | 140 | 70 | 24.48 | 376 | 215 | 229 | 349 |
| *B. sylvaticum* | Bsyl_233 | 45.00 | 0.70 | 12.55 | 9.50 | 40.08 | 540.17 | 25.96 | 2.26 | 23.70 | 9.49 | 19.34 | 19.34 | 6.33 | 997 | 106 | 56 | 19.15 | 305 | 185 | 185 | 285 |
| *B. sylvaticum* | Bsyl_234 | 49.80 | 1.30 | 10.20 | 6.80 | 34.42 | 483.75 | 21.32 | 1.57 | 19.75 | 7.89 | 16.13 | 16.19 | 4.45 | 763 | 82 | 51 | 17.72 | 238 | 162 | 174 | 206 |
| *B. sylvaticum* | Bsyl_235 | 48.40 | 1.90 | 10.49 | 8.59 | 35.21 | 570.14 | 24.99 | 0.60 | 24.39 | 7.27 | 6.55 | 17.66 | 3.77 | 623 | 60 | 42 | 10.27 | 167 | 143 | 147 | 155 |
| *B. sylvaticum* | Bsyl_236 | 45.80 | 2.80 | 8.53 | 9.24 | 37.90 | 567.83 | 22.25 | -2.12 | 24.37 | 13.81 | 2.69 | 15.70 | 1.89 | 807 | 93 | 46 | 21.54 | 240 | 151 | 228 | 164 |
| *B. sylvaticum* | Bsyl_237 | 46.30 | 9.60 | -0.08 | 7.94 | 33.20 | 584.66 | 12.37 | -11.55 | 23.92 | 7.25 | -6.71 | 7.25 | -6.71 | 1066 | 114 | 50 | 24.43 | 334 | 183 | 334 | 183 |
| *B. sylvaticum* | Bsyl_238 | 41.50 | 0.60 | 14.88 | 12.17 | 38.58 | 703.75 | 32.76 | 1.22 | 31.54 | 15.46 | 23.80 | 23.80 | 6.58 | 402 | 52 | 13 | 34.82 | 132 | 73 | 73 | 78 |
| *B. sylvaticum* | Bsyl_239 | 41.90 | 0.90 | 13.13 | 10.69 | 37.12 | 654.72 | 29.27 | 0.48 | 28.79 | 14.76 | 6.60 | 21.53 | 5.52 | 552 | 65 | 26 | 25.88 | 169 | 104 | 126 | 107 |
| *B. sylvaticum* | Bsyl_240 | 42.30 | 3.20 | 14.48 | 8.62 | 35.67 | 561.27 | 27.73 | 3.56 | 24.16 | 15.31 | 21.72 | 21.72 | 8.24 | 571 | 81 | 17 | 31.88 | 196 | 95 | 95 | 138 |
| *B. sylvaticum* | Bsyl_241 | 41.50 | 1.20 | 12.62 | 11.09 | 38.61 | 629.64 | 28.73 | 0.02 | 28.71 | 13.62 | 6.33 | 20.79 | 5.45 | 613 | 70 | 23 | 26.95 | 195 | 123 | 132 | 124 |
| *B. sylvaticum* | Bsyl_242 | 41.70 | 2.90 | 15.05 | 9.12 | 38.41 | 536.00 | 27.63 | 3.90 | 23.73 | 16.11 | 21.83 | 22.01 | 9.06 | 663 | 85 | 25 | 26.69 | 218 | 121 | 147 | 157 |
| *B. sylvaticum* | Bsyl_243 | 40.70 | 0.30 | 13.58 | 11.59 | 41.50 | 594.52 | 29.28 | 1.36 | 27.92 | 14.40 | 21.21 | 21.25 | 6.87 | 547 | 66 | 21 | 31.06 | 176 | 97 | 110 | 120 |
| *B. sylvaticum* | Bsyl_244 | 42.60 | 0.60 | 2.39 | 9.99 | 35.45 | 652.57 | 19.22 | -8.96 | 28.18 | -0.53 | 10.98 | 10.98 | -4.71 | 1392 | 150 | 72 | 20.86 | 420 | 253 | 253 | 379 |
| *B. sylvaticum* | Bsyl_245 | 42.00 | 2.40 | 12.27 | 9.34 | 36.25 | 587.15 | 26.29 | 0.52 | 25.77 | 17.27 | 6.34 | 19.81 | 5.60 | 755 | 84 | 42 | 21.27 | 224 | 148 | 188 | 159 |
| *B. sylvaticum* | Bsyl_246 | 42.00 | 1.60 | 12.29 | 10.22 | 37.58 | 607.37 | 27.29 | 0.10 | 27.19 | 13.66 | 5.40 | 20.16 | 5.40 | 748 | 94 | 36 | 28.85 | 239 | 122 | 204 | 122 |
| *B. sylvaticum* | Bsyl_247 | 42.50 | 1.40 | 4.55 | 9.97 | 36.52 | 628.03 | 20.50 | -6.80 | 27.30 | 1.72 | 12.78 | 12.78 | -2.33 | 1258 | 133 | 69 | 18.32 | 372 | 247 | 247 | 318 |
| *B. sylvaticum* | Bsyl_248 | 41.40 | 2.20 | 15.93 | 8.76 | 37.11 | 547.58 | 28.68 | 5.08 | 23.60 | 20.90 | 10.37 | 23.11 | 9.84 | 622 | 85 | 25 | 30.88 | 218 | 122 | 158 | 132 |
| *B. sylvaticum* | Bsyl_249 | 51.70 | -5.20 | 10.49 | 4.74 | 33.00 | 354.07 | 18.44 | 4.07 | 14.37 | 9.43 | 13.31 | 15.06 | 6.52 | 891 | 108 | 47 | 28.03 | 306 | 154 | 198 | 231 |
| *B. sylvaticum* | Bsyl_250 | 51.60 | -4.20 | 10.45 | 5.95 | 35.27 | 411.15 | 19.73 | 2.87 | 16.86 | 6.90 | 11.45 | 15.65 | 5.89 | 1087 | 128 | 59 | 26.26 | 366 | 192 | 224 | 319 |
| *B. sylvaticum* | Bsyl_251 | 47.40 | 1.50 | 11.43 | 10.40 | 40.26 | 592.95 | 26.38 | 0.56 | 25.82 | 7.81 | 18.91 | 18.91 | 4.31 | 686 | 70 | 48 | 11.54 | 187 | 149 | 149 | 176 |
| *B. sylvaticum* | Bsyl_252 | 46.90 | 2.90 | 10.78 | 10.18 | 39.51 | 594.34 | 25.51 | -0.24 | 25.76 | 16.55 | 6.77 | 18.30 | 3.69 | 746 | 74 | 49 | 12.22 | 204 | 154 | 197 | 180 |
| *B. sylvaticum* | Bsyl_253 | 47.90 | 4.30 | 9.73 | 8.74 | 34.80 | 596.86 | 24.20 | -0.91 | 25.11 | 3.53 | 5.65 | 17.28 | 2.66 | 856 | 85 | 60 | 10.52 | 243 | 197 | 199 | 234 |
| *B. sylvaticum* | Bsyl_254 | 47.70 | 0.80 | 10.79 | 8.61 | 36.75 | 552.68 | 24.74 | 1.30 | 23.44 | 5.13 | 17.77 | 17.77 | 4.35 | 699 | 73 | 48 | 13.81 | 207 | 146 | 146 | 195 |
| *B. sylvaticum* | Bsyl_255 | 49.70 | 4.60 | 9.23 | 8.32 | 35.57 | 568.74 | 22.50 | -0.88 | 23.38 | 3.39 | 8.49 | 16.38 | 2.49 | 887 | 89 | 60 | 12.51 | 257 | 185 | 226 | 248 |
| *B. sylvaticum* | Bsyl_256 | 46.20 | 4.10 | 10.76 | 9.44 | 36.48 | 615.51 | 25.49 | -0.40 | 25.88 | 18.56 | 4.56 | 18.56 | 3.50 | 753 | 86 | 44 | 22.32 | 233 | 142 | 233 | 148 |
| *B. sylvaticum* | Bsyl_257 | 55.90 | 9.90 | 7.68 | 6.61 | 25.96 | 646.08 | 19.20 | -6.26 | 25.46 | 8.61 | 2.30 | 15.65 | -0.07 | 636 | 69 | 34 | 22.91 | 201 | 109 | 176 | 123 |
| *B. sylvaticum* | Bsyl_258 | 55.40 | 9.90 | 8.14 | 6.61 | 28.60 | 603.76 | 20.25 | -2.87 | 23.12 | 9.07 | 3.07 | 15.66 | 1.12 | 643 | 68 | 35 | 20.56 | 195 | 114 | 176 | 147 |
| *B. sylvaticum* | Bsyl_259 | 54.90 | 9.90 | 8.50 | 5.12 | 24.73 | 581.69 | 19.84 | -0.88 | 20.72 | 15.50 | 10.48 | 15.74 | 1.82 | 779 | 86 | 42 | 18.89 | 235 | 156 | 227 | 180 |
| *B. sylvaticum* | Bsyl_261 | 55.30 | 11.80 | 8.62 | 4.51 | 20.77 | 656.65 | 20.50 | -1.21 | 21.71 | 16.21 | 3.19 | 16.96 | 0.94 | 596 | 63 | 32 | 19.69 | 178 | 109 | 171 | 136 |
| *B. sylvaticum* | Bsyl_262 | 55.90 | 12.20 | 7.90 | 5.78 | 25.63 | 648.85 | 20.38 | -2.17 | 22.55 | 15.29 | 3.01 | 16.02 | 0.13 | 587 | 65 | 28 | 23.17 | 187 | 102 | 179 | 125 |
| *B. sylvaticum* | Bsyl_263 | 35.10 | -5.10 | 11.83 | 10.60 | 37.71 | 607.83 | 28.03 | -0.08 | 28.12 | 5.41 | 19.66 | 19.66 | 4.80 | 1021 | 171 | 2 | 77.12 | 501 | 20 | 20 | 441 |
| *B. sylvaticum* | Bsyl_264 | 40.50 | -1.60 | 7.94 | 12.87 | 40.92 | 655.29 | 26.65 | -4.79 | 31.44 | 9.52 | 16.43 | 16.61 | 0.79 | 636 | 84 | 31 | 27.86 | 209 | 114 | 128 | 152 |
| *B. sylvaticum* | Bsyl_265 | 59.90 | 8.60 | 1.61 | 7.82 | 28.35 | 726.46 | 17.36 | -10.24 | 27.60 | 6.37 | -3.67 | 11.13 | -6.63 | 869 | 95 | 42 | 25.57 | 276 | 147 | 265 | 177 |
| *B. sylvaticum* | Bsyl_266 | 53.00 | -9.00 | 10.36 | 6.40 | 39.53 | 377.40 | 19.28 | 3.08 | 16.20 | 8.46 | 11.55 | 15.21 | 6.12 | 1184 | 127 | 67 | 21.84 | 375 | 219 | 260 | 337 |
| *B. sylvaticum* | Bsyl_268 | 51.80 | -10.10 | 9.83 | 5.31 | 39.09 | 320.00 | 17.38 | 3.78 | 13.60 | 7.09 | 12.53 | 13.98 | 6.36 | 1753 | 202 | 96 | 27.68 | 589 | 298 | 373 | 504 |
| *B. sylvaticum* | Bsyl_269 | 53.60 | -9.40 | 9.94 | 5.81 | 38.65 | 352.62 | 18.23 | 3.19 | 15.04 | 8.46 | 10.81 | 14.43 | 6.08 | 1153 | 129 | 64 | 25.95 | 381 | 206 | 228 | 333 |
| *B. sylvaticum* | Bsyl_270 | 52.40 | -6.40 | 10.23 | 5.61 | 37.03 | 358.28 | 18.65 | 3.50 | 15.14 | 7.06 | 13.22 | 14.85 | 6.28 | 962 | 109 | 54 | 24.59 | 312 | 172 | 215 | 300 |
| *B. sylvaticum* | Bsyl_271 | 49.00 | 4.60 | 10.26 | 9.30 | 36.74 | 593.29 | 24.93 | -0.39 | 25.32 | 4.12 | 6.18 | 17.76 | 3.26 | 714 | 69 | 48 | 10.23 | 193 | 154 | 190 | 175 |
| *B. sylvaticum* | Bsyl_272 | 47.60 | 2.80 | 10.59 | 9.36 | 37.49 | 590.70 | 24.87 | -0.09 | 24.96 | 7.21 | 6.58 | 18.04 | 3.51 | 706 | 69 | 49 | 10.69 | 192 | 157 | 170 | 179 |
| *B. sylvaticum* | Bsyl_273 | 47.90 | 3.60 | 10.62 | 8.95 | 35.89 | 597.96 | 24.88 | -0.05 | 24.93 | 10.96 | 6.58 | 18.18 | 3.44 | 671 | 65 | 44 | 11.52 | 184 | 144 | 168 | 164 |
| *B. sylvaticum* | Bsyl_274 | 59.40 | 24.30 | 5.96 | 6.21 | 21.62 | 798.60 | 21.49 | -7.25 | 28.74 | 11.73 | -0.59 | 16.04 | -3.41 | 641 | 74 | 32 | 32.67 | 217 | 100 | 192 | 135 |
| *B. sylvaticum* | Bsyl_275 | 58.40 | 26.80 | 5.25 | 8.13 | 25.63 | 854.63 | 22.51 | -9.20 | 31.71 | 14.49 | -0.92 | 15.96 | -5.05 | 617 | 79 | 28 | 34.90 | 224 | 91 | 223 | 105 |
| *B. sylvaticum* | Bsyl_276 | 58.50 | 21.90 | 6.69 | 4.80 | 19.75 | 708.67 | 19.67 | -4.65 | 24.32 | 7.99 | 0.36 | 15.74 | -1.45 | 584 | 73 | 31 | 31.70 | 205 | 94 | 153 | 106 |
| *B. sylvaticum* | Bsyl_277 | 58.90 | 25.40 | 5.42 | 8.20 | 25.86 | 836.36 | 22.86 | -8.86 | 31.72 | 14.74 | -0.80 | 15.97 | -4.56 | 695 | 86 | 34 | 32.22 | 238 | 106 | 229 | 136 |
| *B. sylvaticum* | Bsyl_278 | 58.00 | 22.10 | 7.22 | 4.99 | 19.23 | 746.34 | 21.06 | -4.90 | 25.96 | 8.68 | 0.52 | 16.61 | -1.48 | 561 | 66 | 27 | 33.19 | 190 | 83 | 167 | 94 |
| *B. sylvaticum* | Bsyl_279 | 34.40 | 36.20 | 10.52 | 12.00 | 40.63 | 665.26 | 27.06 | -2.48 | 29.54 | 2.48 | 18.21 | 18.36 | 2.42 | 800 | 165 | 0 | 90.91 | 444 | 5 | 5 | 416 |
| *B. sylvaticum* | Bsyl_280 | 57.80 | 14.30 | 5.53 | 8.36 | 30.85 | 674.58 | 20.70 | -6.39 | 27.09 | 13.10 | 0.21 | 14.18 | -2.33 | 703 | 78 | 36 | 22.37 | 220 | 125 | 209 | 152 |
| *B. sylvaticum* | Bsyl_282 | 56.80 | 14.00 | 6.23 | 8.10 | 30.73 | 679.59 | 21.13 | -5.23 | 26.36 | 13.79 | 1.08 | 14.89 | -1.78 | 755 | 82 | 43 | 20.92 | 229 | 135 | 214 | 175 |
| *B. sylvaticum* | Bsyl_283 | 48.40 | 12.80 | 8.38 | 8.51 | 31.01 | 718.83 | 23.25 | -4.20 | 27.45 | 17.21 | 0.92 | 17.21 | -0.42 | 897 | 117 | 49 | 30.70 | 330 | 164 | 330 | 166 |
| *B. sylvaticum* | Bsyl_284 | 50.50 | 10.30 | 7.71 | 8.20 | 31.72 | 673.37 | 22.06 | -3.78 | 25.85 | 0.67 | 3.29 | 16.07 | -0.46 | 760 | 81 | 50 | 14.51 | 213 | 162 | 203 | 199 |
| *B. sylvaticum* | Bsyl_285 | 57.00 | 21.40 | 6.63 | 6.16 | 23.84 | 729.75 | 20.40 | -5.43 | 25.82 | 7.68 | 0.81 | 15.75 | -2.00 | 749 | 92 | 32 | 32.02 | 253 | 116 | 218 | 154 |
| *B. sylvaticum* | Bsyl_286 | 50.70 | 34.50 | 6.79 | 8.30 | 25.00 | 955.73 | 24.22 | -8.98 | 33.20 | 18.23 | -3.76 | 18.23 | -5.03 | 592 | 79 | 33 | 27.25 | 208 | 108 | 208 | 118 |
| *B. sylvaticum* | Bsyl_287 | 43.60 | 4.10 | 14.82 | 8.91 | 34.48 | 609.30 | 28.88 | 3.04 | 25.84 | 11.36 | 22.59 | 22.59 | 7.68 | 665 | 105 | 19 | 37.82 | 230 | 93 | 93 | 189 |
| *B. sylvaticum* | Bsyl_288 | 41.40 | 9.20 | 16.59 | 6.67 | 31.39 | 533.32 | 28.40 | 7.16 | 21.24 | 14.71 | 23.23 | 23.50 | 10.68 | 487 | 70 | 8 | 50.95 | 205 | 40 | 60 | 162 |
| *B. sylvaticum* | Bsyl_289 | 44.10 | 4.60 | 14.00 | 10.06 | 35.52 | 658.72 | 29.80 | 1.50 | 28.31 | 14.36 | 22.41 | 22.41 | 6.21 | 818 | 133 | 32 | 35.83 | 289 | 135 | 135 | 194 |
| *B. sylvaticum* | Bsyl_290 | 43.40 | -0.10 | 12.56 | 9.23 | 40.65 | 502.82 | 24.67 | 1.95 | 22.72 | 7.49 | 18.82 | 18.95 | 6.83 | 1346 | 133 | 77 | 16.39 | 384 | 260 | 260 | 370 |
| *B. sylvaticum* | Bsyl_291 | 64.10 | 11.60 | 5.60 | 5.65 | 23.57 | 677.11 | 19.04 | -4.93 | 23.97 | 5.89 | 4.41 | 14.29 | -2.22 | 1035 | 117 | 55 | 24.14 | 321 | 180 | 245 | 289 |
| *B. sylvaticum* | Bsyl_292 | 57.50 | 34.60 | 4.21 | 8.15 | 24.02 | 954.05 | 22.29 | -11.65 | 33.94 | 15.92 | -1.78 | 15.92 | -7.60 | 633 | 88 | 30 | 34.68 | 234 | 96 | 234 | 113 |
| *B. sylvaticum* | Bsyl_293 | 52.30 | 40.00 | 5.49 | 8.77 | 22.86 | 1101.55 | 25.00 | -13.37 | 38.37 | 18.52 | -1.61 | 18.52 | -8.07 | 545 | 70 | 27 | 27.67 | 183 | 89 | 183 | 108 |
| *B. sylvaticum* | Bsyl_294 | 44.50 | 33.70 | 9.77 | 7.52 | 26.37 | 780.01 | 24.94 | -3.59 | 28.53 | 2.25 | 8.69 | 19.53 | 0.38 | 537 | 70 | 35 | 21.80 | 174 | 109 | 131 | 168 |
| *B. sylvaticum* | Bsyl_295 | 56.70 | 41.40 | 3.23 | 8.83 | 22.19 | 1109.11 | 22.98 | -16.81 | 39.79 | 16.23 | -4.72 | 16.23 | -10.68 | 598 | 76 | 27 | 31.08 | 209 | 93 | 209 | 110 |
| *B. sylvaticum* | Bsyl_296 | 54.80 | 42.40 | 4.17 | 8.15 | 21.97 | 1069.75 | 23.27 | -13.80 | 37.07 | 16.95 | -2.52 | 16.95 | -9.17 | 547 | 77 | 27 | 32.32 | 196 | 86 | 196 | 103 |
| *B. sylvaticum* | Bsyl_297 | 51.40 | 42.10 | 6.32 | 9.23 | 24.35 | 1103.28 | 26.25 | -11.67 | 37.92 | 19.47 | -0.79 | 19.47 | -7.27 | 539 | 58 | 31 | 20.84 | 158 | 98 | 158 | 131 |
| *B. sylvaticum* | Bsyl_298 | 53.80 | 57.20 | 1.98 | 10.55 | 25.59 | 1153.53 | 23.00 | -18.24 | 41.24 | 15.81 | -4.75 | 15.81 | -12.31 | 649 | 83 | 31 | 31.39 | 225 | 101 | 225 | 121 |
| *B. sylvaticum* | Bsyl_299 | 45.00 | 35.10 | 9.33 | 7.08 | 25.42 | 765.42 | 23.96 | -3.90 | 27.87 | 2.08 | 10.23 | 18.95 | 0.33 | 560 | 63 | 38 | 14.18 | 158 | 127 | 146 | 154 |
| *B. sylvaticum* | Bsyl_300 | 54.80 | 37.40 | 4.80 | 8.19 | 23.73 | 989.27 | 22.96 | -11.56 | 34.52 | 16.73 | -1.38 | 16.73 | -7.50 | 597 | 84 | 28 | 35.33 | 224 | 91 | 224 | 107 |
| *B. sylvaticum* | Bsyl_301 | 47.10 | 2.20 | 11.26 | 9.43 | 37.69 | 590.13 | 25.72 | 0.71 | 25.01 | 7.79 | 18.70 | 18.70 | 4.28 | 699 | 73 | 49 | 11.04 | 188 | 158 | 158 | 174 |
| *B. sylvaticum* | Bsyl_302 | 47.40 | 5.00 | 10.02 | 8.29 | 32.45 | 643.45 | 24.50 | -1.05 | 25.54 | 16.19 | 5.91 | 18.11 | 2.36 | 793 | 85 | 51 | 13.49 | 219 | 166 | 196 | 191 |
| *B. sylvaticum* | Bsyl_303 | 48.60 | 3.80 | 10.65 | 9.68 | 38.18 | 588.24 | 25.19 | -0.15 | 25.34 | 17.57 | 6.66 | 18.07 | 3.61 | 604 | 61 | 34 | 15.19 | 177 | 118 | 171 | 147 |
| *B. sylvaticum* | Bsyl_304 | 51.80 | -2.00 | 9.12 | 7.23 | 36.59 | 463.55 | 20.47 | 0.70 | 19.77 | 4.78 | 13.45 | 15.08 | 3.86 | 894 | 96 | 57 | 15.45 | 270 | 193 | 203 | 250 |
| *B. sylvaticum* | Bsyl_305 | 48.80 | -3.60 | 11.59 | 4.90 | 33.33 | 358.23 | 19.80 | 5.10 | 14.70 | 8.63 | 15.92 | 16.28 | 7.65 | 926 | 116 | 40 | 35.03 | 341 | 138 | 160 | 284 |
| *B. sylvaticum* | Bsyl_306 | 54.50 | 83.30 | 1.18 | 9.76 | 21.35 | 1335.37 | 24.72 | -21.00 | 45.72 | 17.20 | -13.70 | 17.20 | -15.58 | 448 | 64 | 15 | 41.46 | 179 | 58 | 179 | 75 |
| *B. sylvaticum* | Bsyl_307 | 53.60 | 88.50 | 0.03 | 11.59 | 25.13 | 1255.03 | 23.54 | -22.57 | 46.11 | 15.39 | -13.22 | 15.39 | -15.78 | 748 | 96 | 23 | 41.84 | 276 | 81 | 276 | 106 |
| *B. sylvaticum* | Bsyl_308 | 53.00 | 91.20 | -0.14 | 10.43 | 24.22 | 1221.58 | 21.74 | -21.34 | 43.08 | 14.57 | -13.15 | 14.57 | -15.47 | 520 | 95 | 12 | 64.74 | 253 | 43 | 253 | 51 |
| *B. sylvaticum* | Bsyl_309 | 54.80 | 94.90 | -1.50 | 12.70 | 28.49 | 1163.76 | 21.68 | -22.89 | 44.57 | 12.98 | -14.52 | 12.98 | -15.57 | 553 | 103 | 12 | 64.47 | 265 | 43 | 265 | 52 |
| *B. sylvaticum* | Bsyl_310 | 53.00 | 88.80 | -0.68 | 12.70 | 26.81 | 1268.03 | 23.31 | -24.05 | 47.36 | 14.75 | -13.91 | 14.75 | -16.74 | 778 | 103 | 24 | 42.99 | 289 | 81 | 289 | 107 |
| *B. sylvaticum* | Bsyl_311 | 52.30 | 88.70 | -0.42 | 12.00 | 26.53 | 1232.89 | 22.44 | -22.78 | 45.22 | 14.48 | -13.30 | 14.48 | -16.04 | 675 | 101 | 18 | 51.69 | 278 | 60 | 278 | 78 |
| *B. sylvaticum* | Bsyl_312 | 56.00 | 13.10 | 7.53 | 6.73 | 28.97 | 633.31 | 20.50 | -2.72 | 23.22 | 8.23 | 2.44 | 15.52 | 0.12 | 754 | 78 | 41 | 20.83 | 229 | 141 | 202 | 177 |
| *B. sylvaticum* | Bsyl_313 | 38.00 | -2.80 | 11.30 | 12.39 | 37.56 | 730.67 | 30.94 | -2.04 | 32.98 | 4.13 | 20.76 | 20.96 | 3.21 | 532 | 65 | 8 | 44.60 | 186 | 44 | 49 | 177 |
| *B. sylvaticum* | Bsyl_314 | 51.00 | 4.90 | 10.53 | 7.33 | 32.31 | 574.11 | 23.24 | 0.56 | 22.69 | 7.33 | 6.41 | 17.67 | 3.67 | 763 | 72 | 48 | 12.38 | 210 | 160 | 209 | 185 |
| *B. sylvaticum* | Bsyl_315 | 51.70 | 36.20 | 6.15 | 8.05 | 23.66 | 1003.10 | 23.92 | -10.10 | 34.02 | 18.14 | -5.07 | 18.14 | -6.23 | 625 | 79 | 35 | 25.14 | 207 | 113 | 207 | 123 |
| *B. sylvaticum* | Bsyl_316 | 58.60 | -3.50 | 7.71 | 5.25 | 36.83 | 334.52 | 15.65 | 1.38 | 14.27 | 6.44 | 8.35 | 11.97 | 3.97 | 1025 | 120 | 52 | 27.47 | 341 | 170 | 237 | 291 |
| *B. sylvaticum* | Bsyl_317 | 57.70 | -2.90 | 8.17 | 6.70 | 38.93 | 392.92 | 17.65 | 0.43 | 17.22 | 8.77 | 9.20 | 13.23 | 3.79 | 698 | 70 | 44 | 15.54 | 208 | 144 | 175 | 170 |
| *B. sylvaticum* | Bsyl_318 | 46.60 | 1.40 | 11.55 | 8.64 | 37.09 | 558.40 | 24.78 | 1.50 | 23.28 | 8.32 | 18.57 | 18.57 | 4.97 | 775 | 77 | 52 | 12.61 | 215 | 160 | 160 | 203 |
| *B. sylvaticum* | Bsyl_319 | 47.00 | 7.80 | 7.09 | 8.19 | 33.82 | 609.35 | 20.45 | -3.77 | 24.22 | 14.68 | 0.75 | 14.68 | -0.14 | 1357 | 145 | 95 | 14.64 | 413 | 293 | 413 | 309 |
| *B. sylvaticum* | Bsyl_320 | 45.10 | 0.00 | 12.63 | 9.64 | 40.54 | 536.31 | 25.90 | 2.13 | 23.77 | 6.99 | 19.32 | 19.32 | 6.37 | 904 | 100 | 49 | 21.25 | 287 | 163 | 163 | 265 |
| *B. sylvaticum* | Bsyl_321 | 44.70 | -0.50 | 12.80 | 10.10 | 41.55 | 534.94 | 26.07 | 1.76 | 24.30 | 7.14 | 19.46 | 19.46 | 6.50 | 891 | 99 | 46 | 21.21 | 282 | 163 | 163 | 261 |
| *B. sylvaticum* | Bsyl_322 | 45.10 | -1.00 | 12.72 | 9.74 | 42.65 | 501.58 | 25.19 | 2.36 | 22.84 | 10.04 | 18.86 | 18.86 | 6.76 | 926 | 108 | 45 | 25.10 | 302 | 154 | 154 | 275 |
| *B. sylvaticum* | Bsyl_323 | 45.70 | -0.50 | 12.79 | 9.36 | 40.16 | 533.45 | 25.93 | 2.62 | 23.30 | 7.20 | 19.44 | 19.44 | 6.53 | 849 | 100 | 43 | 24.85 | 278 | 143 | 143 | 251 |
| *B. sylvaticum* | Bsyl_324 | 56.30 | -2.70 | 8.17 | 7.47 | 40.19 | 425.49 | 18.55 | -0.04 | 18.59 | 8.73 | 5.03 | 13.61 | 3.26 | 655 | 66 | 40 | 15.29 | 188 | 133 | 158 | 172 |
| *B. sylvaticum* | Bsyl_325 | 58.30 | 16.90 | 6.73 | 6.65 | 25.39 | 701.52 | 21.15 | -5.03 | 26.18 | 15.05 | 4.78 | 15.86 | -1.29 | 579 | 62 | 32 | 22.09 | 183 | 109 | 172 | 131 |
| *B. sylvaticum* | Bsyl_326 | 53.10 | -1.90 | 7.90 | 6.50 | 35.02 | 446.49 | 18.47 | -0.09 | 18.56 | 5.80 | 12.07 | 13.65 | 2.89 | 1049 | 108 | 72 | 15.76 | 318 | 225 | 240 | 282 |
| *B. sylvaticum* | Bsyl_327 | 48.10 | -0.70 | 11.42 | 8.58 | 38.79 | 512.19 | 24.30 | 2.19 | 22.11 | 6.30 | 17.91 | 17.91 | 5.53 | 751 | 80 | 44 | 19.99 | 234 | 139 | 139 | 224 |
| *B. sylvaticum* | Bsyl_328 | 47.00 | 1.00 | 11.25 | 8.93 | 37.85 | 561.10 | 24.99 | 1.40 | 23.60 | 8.02 | 18.34 | 18.34 | 4.67 | 713 | 70 | 46 | 13.60 | 202 | 147 | 147 | 193 |
| *B. sylvaticum* | Bsyl_329 | 44.60 | 4.10 | 7.96 | 7.82 | 33.36 | 577.66 | 21.52 | -1.93 | 23.45 | 8.66 | 15.39 | 15.39 | 1.57 | 1132 | 151 | 54 | 26.53 | 364 | 203 | 203 | 282 |
| *B. sylvaticum* | Bsyl_330 | 51.20 | 2.90 | 10.02 | 6.82 | 33.12 | 499.86 | 21.41 | 0.84 | 20.58 | 7.74 | 6.04 | 16.23 | 4.14 | 718 | 80 | 42 | 19.84 | 226 | 136 | 188 | 170 |
| *B. sylvaticum* | Bsyl_331 | 51.10 | 4.00 | 10.48 | 7.29 | 33.24 | 550.05 | 22.74 | 0.80 | 21.94 | 7.51 | 6.47 | 17.32 | 3.91 | 786 | 78 | 49 | 14.91 | 227 | 159 | 204 | 186 |
| *B. sylvaticum* | Bsyl_332 | 59.40 | 6.10 | 6.83 | 6.72 | 31.65 | 545.29 | 18.52 | -2.73 | 21.24 | 4.00 | 9.27 | 13.78 | 0.58 | 2274 | 288 | 94 | 35.24 | 811 | 312 | 436 | 699 |
| *B. sylvaticum* | Bsyl_333 | 49.00 | 3.00 | 10.49 | 8.32 | 35.79 | 558.58 | 23.84 | 0.59 | 23.24 | 15.84 | 6.60 | 17.51 | 3.85 | 617 | 58 | 44 | 9.33 | 170 | 134 | 160 | 149 |
| *B. sylvaticum* | Bsyl_334 | 48.30 | 1.10 | 10.08 | 8.27 | 35.79 | 543.99 | 23.78 | 0.67 | 23.11 | 7.09 | 6.21 | 16.93 | 3.71 | 662 | 63 | 44 | 10.07 | 186 | 147 | 157 | 174 |
| *B. sylvaticum* | Bsyl_335 | 47.90 | 2.10 | 10.95 | 9.15 | 37.58 | 583.20 | 24.96 | 0.60 | 24.36 | 7.54 | 18.29 | 18.29 | 4.00 | 646 | 67 | 40 | 12.39 | 176 | 144 | 144 | 164 |
| *B. sylvaticum* | Bsyl_336 | 46.80 | 5.20 | 11.12 | 8.57 | 33.59 | 649.16 | 25.38 | -0.12 | 25.50 | 11.27 | 4.67 | 19.18 | 3.23 | 930 | 96 | 58 | 14.21 | 264 | 199 | 221 | 218 |
| *B. sylvaticum* | Bsyl_337 | 44.30 | 6.80 | 1.35 | 6.83 | 28.33 | 630.27 | 15.34 | -8.78 | 24.12 | -1.12 | 9.33 | 9.61 | -5.38 | 1133 | 126 | 45 | 25.44 | 359 | 181 | 194 | 323 |
| *B. sylvaticum* | Bsyl_338 | 43.40 | 4.80 | 15.15 | 8.15 | 32.67 | 611.02 | 28.71 | 3.75 | 24.96 | 15.78 | 22.90 | 22.90 | 7.94 | 636 | 105 | 17 | 39.42 | 226 | 88 | 88 | 169 |
| *B. sylvaticum* | Bsyl_339 | 41.30 | -7.30 | 13.08 | 8.97 | 39.80 | 490.19 | 25.75 | 3.22 | 22.52 | 8.26 | 19.31 | 19.44 | 7.53 | 1199 | 172 | 17 | 54.57 | 497 | 88 | 99 | 494 |
| *B. sylvaticum* | Bsyl_340 | 42.50 | 8.80 | 14.53 | 7.98 | 34.40 | 560.81 | 27.64 | 4.44 | 23.20 | 12.23 | 21.66 | 21.70 | 8.28 | 817 | 118 | 13 | 47.03 | 324 | 75 | 111 | 249 |
| *B. sylvaticum* | Bsyl_341 | 43.90 | 0.10 | 12.47 | 10.07 | 41.39 | 535.78 | 25.65 | 1.31 | 24.34 | 6.86 | 19.17 | 19.17 | 6.22 | 1122 | 110 | 66 | 14.85 | 323 | 224 | 224 | 312 |
| *B. sylvaticum* | Bsyl_342 | 46.30 | -0.60 | 12.32 | 8.85 | 39.03 | 529.98 | 25.10 | 2.43 | 22.67 | 6.80 | 18.92 | 18.92 | 6.05 | 786 | 102 | 37 | 26.98 | 264 | 134 | 134 | 227 |
| *B. sylvaticum* | Bsyl_343 | 49.30 | -0.10 | 11.49 | 6.18 | 34.45 | 444.60 | 21.45 | 3.50 | 17.95 | 9.63 | 16.95 | 17.13 | 6.39 | 709 | 80 | 42 | 22.48 | 229 | 132 | 146 | 212 |
| *B. sylvaticum* | Bsyl_344 | 46.40 | 6.60 | 10.11 | 7.46 | 30.91 | 623.44 | 23.88 | -0.25 | 24.13 | 18.03 | 5.64 | 18.03 | 2.94 | 1173 | 123 | 79 | 11.79 | 324 | 252 | 324 | 275 |
| *B. sylvaticum* | Bsyl_345 | 47.40 | -2.00 | 11.99 | 7.70 | 36.78 | 494.80 | 24.03 | 3.10 | 20.92 | 7.09 | 18.21 | 18.21 | 6.25 | 809 | 93 | 44 | 26.17 | 270 | 134 | 134 | 251 |
| *B. sylvaticum* | Bsyl_346 | 49.80 | 2.10 | 9.92 | 7.54 | 34.62 | 521.05 | 22.45 | 0.66 | 21.79 | 7.19 | 6.08 | 16.38 | 3.75 | 716 | 72 | 49 | 13.04 | 208 | 152 | 171 | 180 |
| *B. sylvaticum* | Bsyl_347 | 44.90 | 2.60 | 9.00 | 9.90 | 38.77 | 574.67 | 23.32 | -2.21 | 25.53 | 11.05 | 3.03 | 16.27 | 2.21 | 818 | 87 | 52 | 15.84 | 227 | 178 | 191 | 194 |
| *B. sylvaticum* | Bsyl_348 | 48.60 | -2.90 | 11.23 | 6.59 | 37.80 | 403.76 | 21.05 | 3.61 | 17.44 | 7.48 | 16.17 | 16.36 | 6.56 | 925 | 111 | 45 | 29.86 | 325 | 148 | 170 | 299 |
| *B. sylvaticum* | Bsyl_349 | 48.80 | -0.50 | 10.34 | 7.74 | 37.58 | 484.39 | 22.06 | 1.46 | 20.61 | 5.60 | 16.41 | 16.41 | 4.72 | 827 | 90 | 48 | 19.96 | 260 | 154 | 154 | 245 |
| *B. sylvaticum* | Bsyl_350 | 44.60 | -1.20 | 12.87 | 9.52 | 42.39 | 479.21 | 24.99 | 2.52 | 22.46 | 7.96 | 18.73 | 18.84 | 7.24 | 960 | 115 | 43 | 27.54 | 326 | 159 | 189 | 297 |
| *B. sylvaticum* | Bsyl_351 | 58.60 | 12.10 | 5.95 | 7.32 | 28.31 | 677.77 | 20.30 | -5.57 | 25.87 | 6.46 | 4.76 | 14.59 | -2.02 | 902 | 103 | 50 | 23.58 | 297 | 163 | 218 | 224 |
| *B. sylvaticum* | Bsyl_352 | 60.30 | 18.50 | 5.62 | 7.09 | 26.03 | 706.23 | 20.51 | -6.72 | 27.23 | 13.99 | -0.40 | 14.85 | -2.37 | 580 | 70 | 29 | 28.83 | 196 | 99 | 187 | 120 |
| *B. sylvaticum* | Bsyl_353 | 63.90 | 9.90 | 5.78 | 4.99 | 27.29 | 506.48 | 15.85 | -2.43 | 18.28 | 6.41 | 7.65 | 12.19 | 0.01 | 1221 | 145 | 59 | 28.03 | 402 | 206 | 267 | 343 |
| *B. sylvaticum* | Bsyl_354 | 60.00 | 20.40 | 5.57 | 4.53 | 18.64 | 700.81 | 18.30 | -6.00 | 24.30 | 7.02 | -0.75 | 14.53 | -2.46 | 553 | 66 | 27 | 32.36 | 188 | 87 | 160 | 96 |
| *B. sylvaticum* | Bsyl_355 | 60.40 | 19.80 | 5.62 | 6.39 | 24.17 | 695.09 | 19.72 | -6.74 | 26.46 | 10.90 | 3.14 | 14.70 | -2.30 | 565 | 66 | 26 | 30.46 | 185 | 92 | 169 | 104 |
| *B. sylvaticum* | Bsyl_357 | 47.00 | 9.80 | 2.56 | 8.51 | 34.11 | 607.82 | 15.68 | -9.26 | 24.94 | 10.02 | -4.02 | 10.02 | -4.55 | 1384 | 167 | 86 | 24.62 | 478 | 271 | 478 | 278 |
| *B. sylvaticum* | Bsyl_358 | 41.90 | -2.50 | 9.64 | 11.97 | 40.36 | 626.77 | 27.04 | -2.63 | 29.66 | 11.12 | 17.79 | 17.79 | 2.58 | 496 | 61 | 26 | 25.37 | 159 | 86 | 86 | 133 |
| *B. sylvaticum* | Bsyl_359 | 57.50 | 12.00 | 7.34 | 6.17 | 26.53 | 651.47 | 19.94 | -3.34 | 23.28 | 8.12 | 2.02 | 15.60 | -0.29 | 761 | 81 | 43 | 21.95 | 234 | 137 | 199 | 190 |
| *B. sylvaticum* | Bsyl_360 | 53.90 | -2.20 | 8.39 | 6.46 | 35.59 | 440.79 | 18.71 | 0.56 | 18.15 | 4.26 | 9.82 | 14.03 | 3.37 | 1275 | 140 | 73 | 23.02 | 412 | 236 | 274 | 363 |
| *B. sylvaticum* | Bsyl_361 | 50.20 | 2.90 | 9.86 | 7.73 | 34.68 | 535.90 | 22.80 | 0.52 | 22.27 | 6.98 | 5.99 | 16.49 | 3.49 | 694 | 67 | 45 | 11.73 | 195 | 149 | 174 | 166 |
| *B. sylvaticum* | Bsyl_362 | 59.00 | 23.70 | 6.08 | 7.34 | 24.42 | 808.21 | 22.56 | -7.52 | 30.07 | 11.69 | -0.46 | 16.11 | -3.47 | 631 | 77 | 30 | 36.17 | 224 | 93 | 191 | 124 |
| *B. sylvaticum* | Bsyl_363 | 52.60 | -1.40 | 9.18 | 8.00 | 39.29 | 459.60 | 20.84 | 0.48 | 20.36 | 4.79 | 5.62 | 15.07 | 3.94 | 665 | 64 | 45 | 11.94 | 187 | 143 | 165 | 170 |
| *B. sylvaticum* | Bsyl_364 | 46.70 | 6.00 | 7.38 | 8.07 | 32.81 | 625.43 | 21.09 | -3.50 | 24.59 | 3.92 | 3.02 | 15.14 | -0.13 | 1342 | 127 | 96 | 9.60 | 370 | 308 | 318 | 350 |
| *B. sylvaticum* | Bsyl_365 | 45.80 | 3.70 | 8.11 | 8.42 | 35.22 | 577.36 | 21.71 | -2.20 | 23.91 | 13.44 | 2.19 | 15.42 | 1.46 | 855 | 95 | 50 | 20.39 | 253 | 161 | 245 | 170 |
| *B. sylvaticum* | Bsyl_366 | 59.20 | 18.50 | 6.50 | 6.74 | 26.13 | 681.48 | 20.90 | -4.90 | 25.80 | 14.70 | 0.60 | 15.43 | -1.07 | 576 | 64 | 30 | 25.37 | 179 | 100 | 171 | 129 |
| *B. sylvaticum* | Bsyl_368 | 55.40 | -3.80 | 6.65 | 6.43 | 37.42 | 411.52 | 16.57 | -0.62 | 17.19 | 2.70 | 8.12 | 11.96 | 2.03 | 1765 | 206 | 93 | 30.39 | 602 | 286 | 313 | 551 |
| *B. sylvaticum* | Bsyl_369 | 57.70 | -5.50 | 8.60 | 6.12 | 39.60 | 369.05 | 17.03 | 1.57 | 15.46 | 6.79 | 9.75 | 13.24 | 4.33 | 1956 | 234 | 89 | 33.15 | 675 | 294 | 348 | 587 |
| *B. sylvaticum* | Bsyl_370 | 56.90 | -7.50 | 8.89 | 4.62 | 37.23 | 296.91 | 15.60 | 3.20 | 12.40 | 8.02 | 9.22 | 12.75 | 5.58 | 1311 | 152 | 60 | 29.90 | 439 | 199 | 312 | 361 |
| *B. sylvaticum* | Bsyl_371 | 57.40 | -1.90 | 7.44 | 7.03 | 36.20 | 453.99 | 17.90 | -1.52 | 19.42 | 7.95 | 4.59 | 13.05 | 1.89 | 758 | 79 | 51 | 15.88 | 227 | 160 | 178 | 193 |
| *B. sylvaticum* | Bsyl_372 | 56.30 | -4.50 | 5.70 | 5.23 | 34.23 | 394.31 | 14.35 | -0.93 | 15.28 | 2.03 | 6.94 | 10.68 | 1.35 | 2387 | 305 | 118 | 32.75 | 832 | 373 | 408 | 770 |
| *B. sylvaticum* | Bsyl_373 | 52.10 | 5.20 | 9.70 | 8.10 | 35.85 | 549.92 | 22.40 | -0.19 | 22.59 | 6.79 | 5.61 | 16.57 | 3.16 | 806 | 83 | 47 | 16.71 | 240 | 163 | 202 | 196 |
| *B. sylvaticum* | Bsyl_374 | 51.90 | 4.10 | 9.88 | 6.20 | 30.97 | 518.59 | 20.76 | 0.74 | 20.01 | 10.99 | 5.84 | 16.14 | 3.58 | 810 | 94 | 42 | 24.66 | 271 | 152 | 214 | 192 |
| *B. sylvaticum* | Bsyl_375 | 46.40 | -1.60 | 12.73 | 7.57 | 36.74 | 489.46 | 24.26 | 3.64 | 20.62 | 7.82 | 18.73 | 18.73 | 6.89 | 746 | 90 | 34 | 32.23 | 260 | 108 | 108 | 241 |
| *B. sylvaticum* | Bsyl_376 | 49.60 | -1.80 | 11.02 | 5.54 | 33.71 | 399.76 | 20.16 | 3.72 | 16.44 | 7.60 | 15.82 | 16.20 | 6.52 | 925 | 119 | 45 | 33.14 | 334 | 147 | 174 | 291 |
| *B. sylvaticum* | Bsyl_377 | 52.90 | -2.80 | 9.17 | 8.01 | 40.18 | 441.63 | 20.48 | 0.54 | 19.94 | 6.95 | 5.70 | 14.88 | 4.14 | 707 | 72 | 48 | 13.81 | 208 | 152 | 168 | 180 |
| *B. sylvaticum* | Bsyl_378 | 48.40 | 5.80 | 9.27 | 8.30 | 33.71 | 621.33 | 23.09 | -1.54 | 24.64 | 15.33 | 5.18 | 17.05 | 1.85 | 819 | 81 | 52 | 12.07 | 224 | 174 | 220 | 215 |
| *B. sylvaticum* | Bsyl_379 | 49.10 | 5.40 | 9.87 | 8.35 | 34.73 | 601.28 | 23.59 | -0.44 | 24.04 | 3.58 | 5.81 | 17.45 | 2.72 | 777 | 77 | 49 | 11.85 | 212 | 166 | 209 | 206 |
| *B. sylvaticum* | Bsyl_380 | 49.10 | 6.10 | 10.25 | 8.11 | 33.11 | 627.31 | 24.03 | -0.45 | 24.48 | 6.39 | 6.18 | 18.15 | 2.76 | 752 | 77 | 51 | 10.83 | 209 | 171 | 187 | 200 |
| *B. sylvaticum* | Bsyl_381 | 49.10 | 6.90 | 9.36 | 8.34 | 33.40 | 633.56 | 23.50 | -1.48 | 24.98 | 5.49 | 5.25 | 17.33 | 1.77 | 874 | 95 | 58 | 13.92 | 259 | 193 | 206 | 237 |
| *B. sylvaticum* | Bsyl_382 | 47.00 | 6.50 | 6.24 | 7.81 | 32.25 | 619.82 | 19.76 | -4.45 | 24.21 | -0.04 | 1.74 | 13.93 | -1.11 | 1371 | 131 | 97 | 8.47 | 371 | 313 | 344 | 360 |
| *B. sylvaticum* | Bsyl_383 | 44.50 | 3.30 | 9.36 | 9.20 | 36.08 | 594.28 | 23.85 | -1.63 | 25.48 | 10.03 | 16.95 | 16.95 | 2.56 | 787 | 101 | 38 | 23.00 | 240 | 152 | 152 | 201 |
| *B. sylvaticum* | Bsyl_384 | 39.30 | 8.80 | 17.04 | 9.76 | 37.00 | 612.37 | 32.14 | 5.76 | 26.38 | 14.46 | 24.83 | 24.91 | 10.29 | 540 | 86 | 5 | 53.99 | 217 | 33 | 53 | 179 |
| *B. sylvaticum* | Bsyl_385 | 40.20 | 9.50 | 13.57 | 6.08 | 28.35 | 577.99 | 25.94 | 4.50 | 21.44 | 11.24 | 20.93 | 20.98 | 7.21 | 621 | 91 | 10 | 47.41 | 237 | 52 | 70 | 222 |
| *B. sylvaticum* | Bsyl_386 | 40.30 | 8.80 | 12.95 | 8.05 | 33.06 | 596.92 | 27.48 | 3.13 | 24.35 | 7.54 | 20.47 | 20.62 | 6.33 | 960 | 145 | 8 | 58.00 | 401 | 63 | 73 | 374 |
| *B. sylvaticum* | Bsyl_387 | 40.80 | 9.30 | 14.92 | 7.07 | 31.40 | 578.42 | 27.93 | 5.41 | 22.52 | 12.58 | 22.30 | 22.37 | 8.56 | 612 | 96 | 8 | 52.31 | 253 | 50 | 61 | 223 |
| *B. sylvaticum* | Bsyl_388 | 39.80 | 8.70 | 16.74 | 8.31 | 34.92 | 579.05 | 30.36 | 6.55 | 23.81 | 14.45 | 23.99 | 24.20 | 10.34 | 556 | 90 | 6 | 55.31 | 231 | 33 | 49 | 185 |
| *B. sylvaticum* | Bsyl_389 | 41.10 | 8.30 | 15.32 | 7.57 | 35.75 | 504.10 | 27.32 | 6.13 | 21.19 | 13.65 | 21.57 | 21.87 | 9.83 | 513 | 85 | 3 | 59.77 | 225 | 29 | 51 | 162 |
| *B. sylvaticum* | Bsyl_390 | 52.10 | 17.40 | 8.70 | 8.48 | 30.58 | 737.38 | 24.10 | -3.64 | 27.74 | 17.80 | 0.71 | 17.80 | -0.33 | 500 | 74 | 21 | 37.28 | 194 | 76 | 194 | 85 |
| *B. sylvaticum* | Bsyl_391 | 50.40 | 30.50 | 8.09 | 7.73 | 24.79 | 903.91 | 24.30 | -6.88 | 31.18 | 18.96 | -1.95 | 18.96 | -3.03 | 610 | 86 | 33 | 32.72 | 225 | 108 | 225 | 120 |
| *B. sylvaticum* | Bsyl_392 | 50.90 | 14.90 | 8.71 | 8.21 | 31.04 | 697.03 | 23.48 | -2.97 | 26.46 | 17.27 | 1.24 | 17.27 | 0.19 | 705 | 96 | 34 | 37.27 | 268 | 118 | 268 | 121 |
| *B. sylvaticum* | Bsyl_393 | 52.20 | 10.80 | 7.98 | 7.39 | 31.26 | 615.36 | 21.18 | -2.46 | 23.64 | 15.64 | 3.47 | 15.64 | 0.62 | 731 | 74 | 47 | 12.91 | 209 | 157 | 209 | 177 |
| *B. sylvaticum* | Bsyl_394 | 53.70 | 12.50 | 8.42 | 7.56 | 30.39 | 664.07 | 22.30 | -2.56 | 24.86 | 16.62 | 3.65 | 16.62 | 0.38 | 555 | 62 | 33 | 20.64 | 180 | 106 | 180 | 120 |
| *B. sylvaticum* | Bsyl_395 | 51.90 | 8.20 | 9.50 | 8.25 | 34.68 | 588.72 | 22.79 | -1.00 | 23.79 | 16.88 | 5.30 | 16.88 | 2.47 | 757 | 78 | 42 | 16.37 | 225 | 149 | 225 | 182 |
| *B. sylvaticum* | Bsyl_396 | 49.40 | 8.60 | 10.86 | 8.42 | 32.36 | 676.11 | 25.51 | -0.52 | 26.04 | 17.70 | 3.96 | 19.32 | 2.74 | 745 | 80 | 48 | 18.46 | 235 | 149 | 212 | 160 |
| *B. sylvaticum* | Bsyl_397 | 53.70 | 13.30 | 8.51 | 7.56 | 30.28 | 663.49 | 22.46 | -2.50 | 24.96 | 16.72 | 3.85 | 16.72 | 0.44 | 532 | 64 | 29 | 25.09 | 181 | 99 | 181 | 105 |
| *B. sylvaticum* | Bsyl_398 | 50.20 | 7.40 | 9.21 | 7.69 | 32.18 | 619.63 | 22.72 | -1.19 | 23.91 | 15.47 | 5.07 | 17.00 | 1.80 | 658 | 65 | 42 | 12.36 | 190 | 139 | 184 | 150 |
| *B. sylvaticum* | Bsyl_399 | 52.80 | 9.90 | 9.20 | 8.03 | 31.85 | 659.01 | 23.32 | -1.90 | 25.22 | 2.42 | 4.72 | 17.40 | 1.23 | 744 | 77 | 49 | 13.75 | 209 | 161 | 202 | 194 |
| *B. sylvaticum* | Bsyl_400 | 49.40 | 9.30 | 8.85 | 8.52 | 32.81 | 670.37 | 23.54 | -2.43 | 25.96 | 4.65 | 4.56 | 17.26 | 0.78 | 826 | 88 | 54 | 13.36 | 234 | 187 | 205 | 228 |
| *B. sylvaticum* | Bsyl_401 | 52.00 | 14.40 | 8.96 | 8.22 | 31.18 | 690.32 | 23.47 | -2.88 | 26.35 | 17.50 | 4.35 | 17.50 | 0.55 | 563 | 70 | 33 | 24.64 | 188 | 105 | 188 | 117 |
| *B. sylvaticum* | Bsyl_402 | 54.30 | 11.00 | 8.66 | 6.49 | 29.04 | 604.30 | 21.48 | -0.88 | 22.35 | 15.91 | 3.96 | 16.34 | 1.67 | 657 | 67 | 38 | 16.12 | 187 | 129 | 184 | 156 |
| *B. sylvaticum* | Bsyl_403 | 51.30 | 8.00 | 8.11 | 8.36 | 34.60 | 585.31 | 21.55 | -2.60 | 24.15 | 2.32 | 10.91 | 15.51 | 1.18 | 1102 | 117 | 75 | 14.78 | 331 | 247 | 270 | 306 |
| *B. sylvaticum* | Bsyl_404 | 53.20 | 8.70 | 8.90 | 7.31 | 32.20 | 589.50 | 21.58 | -1.13 | 22.71 | 16.25 | 4.60 | 16.25 | 1.86 | 705 | 74 | 38 | 17.20 | 209 | 135 | 209 | 157 |
| *B. sylvaticum* | Bsyl_405 | 51.80 | 12.80 | 9.17 | 8.75 | 32.20 | 701.65 | 24.26 | -2.91 | 27.17 | 17.93 | 4.61 | 17.93 | 0.63 | 553 | 60 | 33 | 19.26 | 174 | 108 | 174 | 119 |
| *B. sylvaticum* | Bsyl_406 | 51.30 | 12.20 | 9.34 | 8.19 | 30.98 | 677.61 | 24.38 | -2.06 | 26.45 | 17.75 | 2.13 | 17.75 | 1.15 | 505 | 65 | 28 | 27.88 | 179 | 92 | 179 | 92 |
| *B. sylvaticum* | Bsyl_407 | 50.10 | 8.20 | 9.08 | 7.38 | 30.57 | 639.35 | 22.61 | -1.54 | 24.16 | 15.58 | 4.92 | 17.12 | 1.42 | 682 | 70 | 46 | 13.35 | 199 | 146 | 185 | 163 |
| *B. sylvaticum* | Bsyl_408 | 49.60 | 7.50 | 9.02 | 8.58 | 33.82 | 631.18 | 23.50 | -1.88 | 25.38 | 5.19 | 4.78 | 16.99 | 1.48 | 697 | 70 | 45 | 11.85 | 197 | 149 | 179 | 175 |
| *B. sylvaticum* | Bsyl_409 | 48.90 | 8.70 | 9.30 | 9.14 | 35.02 | 644.77 | 24.14 | -1.96 | 26.10 | 15.88 | 13.72 | 17.46 | 1.62 | 944 | 96 | 68 | 12.36 | 269 | 213 | 247 | 233 |
| *B. sylvaticum* | Bsyl_410 | 53.90 | 14.00 | 8.34 | 6.79 | 28.18 | 653.83 | 21.68 | -2.40 | 24.08 | 16.56 | 3.51 | 16.56 | 0.58 | 572 | 62 | 31 | 21.54 | 184 | 105 | 184 | 119 |
| *B. sylvaticum* | Bsyl_411 | 52.30 | 7.10 | 9.35 | 7.85 | 34.57 | 571.90 | 22.20 | -0.51 | 22.70 | 3.59 | 5.21 | 16.48 | 2.53 | 781 | 76 | 46 | 15.37 | 217 | 157 | 216 | 195 |
| *B. sylvaticum* | Bsyl_412 | 51.70 | 9.20 | 8.64 | 7.72 | 31.98 | 615.72 | 22.22 | -1.92 | 24.14 | 2.36 | 4.33 | 16.29 | 1.19 | 824 | 81 | 53 | 12.18 | 226 | 175 | 224 | 211 |
| *B. sylvaticum* | Bsyl_413 | 50.30 | 6.40 | 7.09 | 7.00 | 30.87 | 592.90 | 19.86 | -2.81 | 22.66 | 1.10 | 9.72 | 14.51 | 0.06 | 1191 | 129 | 72 | 17.36 | 368 | 253 | 257 | 348 |
| *B. sylvaticum* | Bsyl_414 | 51.50 | 14.90 | 8.89 | 8.15 | 30.91 | 698.64 | 23.46 | -2.91 | 26.37 | 17.51 | 4.28 | 17.51 | 0.37 | 647 | 90 | 39 | 27.98 | 224 | 122 | 224 | 135 |
| *B. sylvaticum* | Bsyl_415 | 54.50 | 13.30 | 8.24 | 6.85 | 29.32 | 626.91 | 21.39 | -1.98 | 23.36 | 16.15 | 3.26 | 16.15 | 0.95 | 590 | 72 | 33 | 22.96 | 189 | 107 | 189 | 118 |
| *B. sylvaticum* | Bsyl_416 | 51.90 | 11.50 | 9.14 | 8.49 | 32.81 | 650.38 | 23.87 | -2.00 | 25.86 | 15.55 | 2.18 | 17.22 | 1.30 | 475 | 55 | 24 | 27.57 | 162 | 84 | 161 | 87 |
| *B. sylvaticum* | Bsyl_417 | 52.00 | 10.00 | 8.44 | 7.68 | 31.80 | 617.22 | 22.00 | -2.17 | 24.16 | 16.15 | 4.04 | 16.15 | 1.02 | 768 | 77 | 49 | 12.70 | 218 | 165 | 218 | 188 |
| *B. sylvaticum* | Bsyl_418 | 51.20 | 11.40 | 8.65 | 8.24 | 31.74 | 657.61 | 23.25 | -2.72 | 25.97 | 15.08 | 1.61 | 16.78 | 0.69 | 595 | 71 | 35 | 22.29 | 191 | 116 | 189 | 118 |
| *B. sylvaticum* | Bsyl_419 | 52.80 | 9.10 | 9.07 | 8.18 | 33.51 | 609.54 | 23.00 | -1.40 | 24.40 | 16.71 | 4.65 | 16.71 | 1.83 | 707 | 73 | 43 | 15.17 | 206 | 143 | 206 | 172 |
| *B. sylvaticum* | Bsyl_420 | 55.50 | -1.60 | 8.21 | 7.21 | 37.94 | 445.04 | 18.83 | -0.17 | 18.99 | 3.72 | 5.02 | 13.80 | 2.94 | 671 | 66 | 45 | 13.58 | 193 | 143 | 170 | 173 |
| *B. sylvaticum* | Bsyl_421 | 47.70 | 15.90 | 8.17 | 9.90 | 35.39 | 677.81 | 23.00 | -4.96 | 27.96 | 16.55 | -0.07 | 16.55 | -0.07 | 899 | 118 | 43 | 32.92 | 327 | 144 | 327 | 144 |
| *B. sylvaticum* | Bsyl_422 | 53.10 | 32.30 | 5.67 | 8.75 | 26.77 | 926.00 | 22.93 | -9.78 | 32.70 | 16.77 | -4.61 | 16.77 | -5.82 | 614 | 80 | 30 | 32.27 | 225 | 99 | 225 | 109 |
| *B. sylvaticum* | Bsyl_423 | 41.50 | -1.80 | 11.96 | 12.72 | 41.34 | 643.72 | 29.82 | -0.96 | 30.78 | 13.69 | 20.24 | 20.29 | 4.69 | 410 | 57 | 23 | 28.82 | 142 | 77 | 91 | 94 |
| *B. sylvaticum* | Bsyl_424 | 42.30 | 13.30 | 10.23 | 8.41 | 31.99 | 666.68 | 24.95 | -1.35 | 26.30 | 11.21 | 3.08 | 18.65 | 2.44 | 847 | 105 | 40 | 27.30 | 281 | 138 | 230 | 150 |
| *B. sylvaticum* | Bsyl_425 | 40.00 | -0.60 | 12.46 | 13.13 | 42.68 | 631.07 | 30.07 | -0.68 | 30.75 | 14.10 | 6.25 | 20.71 | 5.41 | 415 | 55 | 21 | 30.44 | 137 | 71 | 95 | 78 |
| *B. sylvaticum* | Bsyl_426 | 42.30 | -7.20 | 12.25 | 10.15 | 41.12 | 524.80 | 26.40 | 1.72 | 24.69 | 6.92 | 18.95 | 19.01 | 6.24 | 1461 | 203 | 34 | 48.08 | 575 | 130 | 158 | 549 |
| *B. sylvaticum* | Bsyl_427 | 41.20 | -1.30 | 11.75 | 13.50 | 42.51 | 652.07 | 30.16 | -1.60 | 31.76 | 13.51 | 5.38 | 20.17 | 4.37 | 418 | 64 | 20 | 37.55 | 158 | 67 | 112 | 74 |
| *B. sylvaticum* | Bsyl_428 | 38.00 | 22.50 | 12.03 | 10.61 | 37.90 | 661.85 | 28.22 | 0.21 | 28.00 | 5.89 | 20.45 | 20.45 | 4.43 | 769 | 119 | 20 | 55.99 | 325 | 65 | 65 | 313 |
| *B. sylvaticum* | Bsyl_429 | 40.80 | -0.90 | 10.37 | 14.15 | 43.02 | 663.86 | 29.61 | -3.28 | 32.88 | 12.24 | 3.89 | 19.04 | 2.91 | 471 | 73 | 22 | 37.52 | 179 | 74 | 130 | 79 |
| *B. sylvaticum* | Bsyl_430 | 41.30 | 32.70 | 9.43 | 10.16 | 35.24 | 711.01 | 25.11 | -3.73 | 28.84 | 2.08 | 17.56 | 17.84 | 0.44 | 763 | 96 | 33 | 30.33 | 251 | 114 | 140 | 242 |
| *B. sylvaticum* | Bsyl_431 | 40.10 | 0.00 | 14.99 | 11.35 | 41.92 | 578.25 | 29.76 | 2.70 | 27.07 | 15.89 | 22.38 | 22.47 | 8.47 | 464 | 66 | 17 | 34.56 | 162 | 77 | 102 | 106 |
| *B. sylvaticum* | Bsyl_432 | 39.70 | 9.40 | 14.94 | 8.16 | 33.54 | 594.65 | 29.02 | 4.68 | 24.34 | 12.48 | 22.53 | 22.63 | 8.42 | 516 | 81 | 10 | 47.91 | 202 | 48 | 69 | 187 |
| *B. sylvaticum* | Bsyl_433 | 46.70 | 3.70 | 10.88 | 9.84 | 37.87 | 609.61 | 25.74 | -0.23 | 25.97 | 18.59 | 6.83 | 18.59 | 3.60 | 757 | 79 | 48 | 16.68 | 214 | 149 | 214 | 171 |
| *B. sylvaticum* | Bsyl_434 | 45.20 | 8.10 | 12.37 | 9.02 | 31.01 | 739.89 | 27.56 | -1.53 | 29.09 | 15.94 | 3.19 | 21.46 | 3.19 | 729 | 99 | 36 | 30.28 | 244 | 121 | 177 | 121 |
| *B. sylvaticum* | Bsyl_435 | 53.40 | 24.50 | 6.34 | 8.60 | 28.57 | 810.55 | 22.31 | -7.80 | 30.11 | 16.18 | -2.81 | 16.18 | -3.81 | 603 | 81 | 29 | 30.59 | 214 | 98 | 214 | 106 |
| *B. sylvaticum* | Bsyl_436 | 39.80 | -1.20 | 12.23 | 12.45 | 40.46 | 650.70 | 30.07 | -0.71 | 30.78 | 13.97 | 5.85 | 20.77 | 4.97 | 399 | 53 | 21 | 31.00 | 135 | 69 | 89 | 77 |
| *B. sylvaticum* | Bsyl_437 | 39.30 | -1.10 | 14.04 | 11.89 | 39.10 | 661.37 | 31.30 | 0.89 | 30.41 | 15.81 | 22.63 | 22.63 | 6.55 | 379 | 47 | 14 | 32.41 | 123 | 67 | 67 | 79 |
| *B. sylvaticum* | Bsyl_438 | 50.30 | 4.50 | 9.44 | 7.92 | 34.36 | 559.95 | 22.66 | -0.41 | 23.06 | 14.77 | 5.45 | 16.45 | 2.79 | 823 | 80 | 56 | 10.11 | 222 | 185 | 214 | 200 |
| *B. sylvaticum* | Bsyl_439 | 50.20 | 5.30 | 9.12 | 7.49 | 32.73 | 579.06 | 21.93 | -0.95 | 22.88 | 14.74 | 5.03 | 16.36 | 2.20 | 891 | 84 | 62 | 9.47 | 242 | 198 | 241 | 212 |
| *B. sylvaticum* | Bsyl_440 | 49.70 | 5.50 | 8.70 | 7.74 | 33.69 | 582.87 | 21.58 | -1.39 | 22.97 | 2.71 | 15.51 | 16.04 | 1.77 | 1022 | 114 | 66 | 16.01 | 312 | 223 | 227 | 297 |
| *B. sylvaticum* | Bsyl_441 | 53.00 | 4.80 | 9.71 | 5.49 | 28.31 | 517.77 | 20.30 | 0.90 | 19.40 | 11.03 | 5.28 | 16.08 | 3.68 | 762 | 97 | 38 | 31.73 | 279 | 133 | 206 | 181 |
| *B. sylvaticum* | Bsyl_442 | 49.70 | 18.60 | 8.00 | 9.21 | 32.19 | 730.92 | 23.22 | -5.38 | 28.60 | 16.84 | 0.11 | 16.84 | -1.22 | 789 | 105 | 36 | 38.70 | 305 | 116 | 305 | 122 |
| *B. sylvaticum* | Bsyl_443 | 50.50 | 14.50 | 8.39 | 7.41 | 28.55 | 697.87 | 22.57 | -3.37 | 25.94 | 16.94 | -0.13 | 16.94 | -0.13 | 589 | 85 | 28 | 42.33 | 235 | 92 | 235 | 92 |
| *B. sylvaticum* | Bsyl_445 | 47.70 | 6.40 | 9.64 | 9.63 | 36.51 | 636.65 | 24.41 | -1.97 | 26.38 | 5.73 | 5.58 | 17.58 | 1.98 | 1070 | 105 | 75 | 11.48 | 302 | 237 | 257 | 280 |
| *B. sylvaticum* | Bsyl_446 | 47.30 | -3.20 | 12.37 | 5.51 | 33.80 | 388.06 | 21.52 | 5.21 | 16.30 | 9.15 | 17.10 | 17.37 | 8.03 | 908 | 117 | 40 | 34.31 | 335 | 135 | 158 | 266 |
| *B. sylvaticum* | Bsyl_447 | 51.10 | 10.60 | 8.62 | 8.75 | 33.57 | 655.33 | 23.38 | -2.68 | 26.06 | 16.76 | 4.19 | 16.76 | 0.70 | 606 | 73 | 35 | 20.37 | 192 | 122 | 192 | 128 |
| *B. sylvaticum* | Bsyl_448 | 50.70 | 12.30 | 8.07 | 7.91 | 31.01 | 671.36 | 22.22 | -3.30 | 25.52 | 16.33 | -0.12 | 16.33 | -0.12 | 730 | 94 | 39 | 27.24 | 248 | 132 | 248 | 132 |
| *B. sylvaticum* | Bsyl_449 | 53.20 | 14.10 | 8.72 | 8.18 | 30.99 | 688.19 | 23.59 | -2.80 | 26.40 | 17.28 | 4.02 | 17.28 | 0.43 | 486 | 60 | 26 | 28.92 | 173 | 87 | 173 | 93 |
| *B. sylvaticum* | Bsyl_450 | 51.20 | 13.10 | 9.06 | 7.98 | 29.49 | 715.81 | 23.57 | -3.48 | 27.04 | 17.09 | 4.33 | 17.82 | 0.16 | 642 | 83 | 38 | 25.95 | 213 | 120 | 211 | 139 |
| *B. sylvaticum* | Bsyl_451 | 53.60 | 7.90 | 8.87 | 7.18 | 32.28 | 565.55 | 21.25 | -1.00 | 22.24 | 9.63 | 4.52 | 15.89 | 2.15 | 808 | 85 | 44 | 18.21 | 238 | 155 | 216 | 186 |
| *B. sylvaticum* | Bsyl_452 | 52.70 | 14.10 | 8.88 | 8.14 | 30.64 | 697.84 | 23.68 | -2.88 | 26.56 | 17.56 | 4.24 | 17.56 | 0.45 | 552 | 64 | 32 | 23.99 | 185 | 102 | 185 | 116 |
| *B. sylvaticum* | Bsyl_453 | 53.70 | 8.80 | 8.90 | 6.38 | 29.54 | 584.30 | 20.78 | -0.82 | 21.60 | 9.60 | 4.43 | 16.16 | 1.97 | 764 | 78 | 39 | 20.89 | 227 | 138 | 217 | 169 |
| *B. sylvaticum* | Bsyl_454 | 53.40 | 11.40 | 8.78 | 7.94 | 31.86 | 644.38 | 22.70 | -2.24 | 24.93 | 16.79 | 4.37 | 16.79 | 0.99 | 615 | 70 | 36 | 18.14 | 193 | 123 | 193 | 139 |
| *B. sylvaticum* | Bsyl_455 | 54.30 | 12.60 | 8.42 | 6.69 | 28.61 | 636.60 | 21.45 | -1.93 | 23.38 | 16.35 | 3.42 | 16.35 | 0.91 | 631 | 68 | 36 | 20.23 | 197 | 116 | 197 | 134 |
| *B. sylvaticum* | Bsyl_456 | 53.00 | 13.30 | 8.71 | 7.81 | 30.30 | 687.51 | 22.97 | -2.80 | 25.77 | 17.26 | 3.93 | 17.26 | 0.40 | 560 | 64 | 30 | 22.85 | 182 | 102 | 182 | 121 |
| *B. sylvaticum* | Bsyl_457 | 46.40 | 3.20 | 10.35 | 10.07 | 39.35 | 586.51 | 25.00 | -0.60 | 25.60 | 15.91 | 6.30 | 17.78 | 3.45 | 759 | 85 | 46 | 20.68 | 221 | 146 | 221 | 163 |
| *B. sylvaticum* | Bsyl_458 | 47.40 | 5.90 | 9.96 | 8.05 | 32.87 | 623.62 | 23.51 | -1.00 | 24.50 | 6.15 | 5.99 | 17.69 | 2.43 | 1095 | 105 | 76 | 11.34 | 307 | 247 | 253 | 275 |
| *B. sylvaticum* | Bsyl_459 | 54.10 | -6.20 | 8.76 | 7.18 | 39.36 | 417.53 | 19.08 | 0.83 | 18.24 | 4.55 | 12.72 | 14.12 | 3.97 | 1518 | 171 | 88 | 23.90 | 489 | 272 | 302 | 464 |
| *B. sylvaticum* | Bsyl_460 | 52.30 | -7.70 | 8.62 | 6.99 | 40.51 | 393.91 | 18.48 | 1.23 | 17.25 | 4.75 | 12.38 | 13.74 | 4.16 | 1617 | 188 | 91 | 24.71 | 521 | 293 | 305 | 515 |
| *B. sylvaticum* | Bsyl_461 | 52.90 | -6.30 | 8.85 | 6.35 | 39.81 | 369.63 | 17.92 | 1.96 | 15.96 | 5.42 | 12.14 | 13.61 | 4.80 | 1156 | 126 | 69 | 20.88 | 361 | 222 | 229 | 346 |
| *B. sylvaticum* | Bsyl_462 | 51.90 | -9.10 | 9.61 | 6.00 | 40.08 | 341.39 | 17.96 | 3.00 | 14.96 | 6.57 | 12.60 | 13.98 | 5.86 | 1085 | 132 | 54 | 31.63 | 381 | 175 | 181 | 366 |
| *B. sylvaticum* | Bsyl_463 | 53.30 | -7.80 | 9.61 | 7.35 | 41.98 | 390.80 | 19.59 | 2.09 | 17.50 | 7.55 | 10.86 | 14.71 | 5.24 | 957 | 99 | 60 | 18.19 | 292 | 191 | 217 | 263 |
| *B. sylvaticum* | Bsyl_464 | 52.90 | -8.20 | 9.77 | 7.19 | 41.49 | 388.88 | 19.50 | 2.17 | 17.33 | 7.74 | 10.99 | 14.82 | 5.44 | 1010 | 110 | 61 | 22.10 | 324 | 193 | 226 | 287 |
| *B. sylvaticum* | Bsyl_465 | 52.90 | -7.30 | 9.32 | 7.37 | 41.98 | 385.90 | 19.43 | 1.87 | 17.56 | 5.64 | 10.47 | 14.38 | 5.08 | 1151 | 122 | 70 | 20.13 | 357 | 218 | 246 | 337 |
| *B. sylvaticum* | Bsyl_466 | 51.90 | -8.10 | 9.97 | 6.65 | 40.28 | 373.52 | 19.17 | 2.67 | 16.50 | 6.35 | 11.14 | 14.71 | 5.64 | 1208 | 140 | 66 | 28.98 | 404 | 202 | 218 | 393 |
| *B. sylvaticum* | Bsyl_467 | 53.40 | -6.10 | 9.85 | 7.04 | 39.69 | 402.55 | 19.84 | 2.10 | 17.74 | 5.93 | 11.10 | 15.01 | 5.28 | 798 | 82 | 49 | 18.00 | 239 | 155 | 191 | 228 |
| *B. sylvaticum* | Bsyl_468 | 51.50 | -0.10 | 10.83 | 8.20 | 38.15 | 493.62 | 23.34 | 1.86 | 21.48 | 6.13 | 7.02 | 17.24 | 5.27 | 628 | 63 | 39 | 15.51 | 185 | 132 | 145 | 162 |
| *B. sylvaticum* | Bsyl_469 | 38.50 | 16.30 | 12.22 | 6.34 | 30.29 | 551.91 | 23.65 | 2.73 | 20.92 | 7.42 | 18.96 | 19.08 | 5.86 | 897 | 130 | 21 | 52.96 | 363 | 71 | 106 | 343 |
| *B. sylvaticum* | Bsyl_470 | 43.30 | 19.40 | 7.92 | 10.00 | 33.86 | 718.79 | 23.45 | -6.09 | 29.54 | 8.65 | 0.10 | 16.64 | -1.12 | 1062 | 115 | 74 | 14.32 | 311 | 226 | 248 | 248 |
| *B. sylvaticum* | Bsyl_471 | 46.00 | 14.70 | 8.75 | 9.79 | 33.96 | 725.04 | 24.43 | -4.40 | 28.82 | 17.58 | 1.28 | 17.58 | -0.31 | 1374 | 155 | 68 | 23.75 | 416 | 231 | 416 | 241 |
| *B. sylvaticum* | Bsyl_472 | 46.80 | 23.50 | 7.99 | 9.64 | 31.96 | 787.11 | 23.53 | -6.64 | 30.17 | 15.93 | -0.39 | 17.28 | -2.11 | 606 | 96 | 26 | 45.53 | 253 | 83 | 243 | 92 |
| *B. sylvaticum* | Bsyl_473 | 46.10 | 12.40 | 7.87 | 8.06 | 29.91 | 703.32 | 22.13 | -4.81 | 26.94 | 14.64 | -0.67 | 16.58 | -0.67 | 1035 | 117 | 45 | 31.16 | 337 | 150 | 336 | 150 |
| *B. sylvaticum* | Bsyl_474 | 52.10 | 23.70 | 7.74 | 8.10 | 27.81 | 808.97 | 23.47 | -5.68 | 29.14 | 17.54 | -1.26 | 17.54 | -2.25 | 574 | 80 | 28 | 35.52 | 217 | 87 | 217 | 98 |
| *B. sylvaticum* | Bsyl_475 | 54.20 | -1.10 | 8.54 | 7.17 | 37.60 | 442.39 | 19.51 | 0.44 | 19.06 | 4.31 | 5.12 | 14.21 | 3.50 | 729 | 72 | 48 | 14.24 | 213 | 154 | 179 | 191 |
| *B. sylvaticum* | Bsyl_476 | 40.80 | 39.10 | 9.18 | 8.65 | 32.28 | 682.77 | 23.25 | -3.54 | 26.79 | 12.09 | 17.14 | 17.14 | 0.70 | 709 | 82 | 33 | 26.87 | 229 | 116 | 116 | 167 |
| *B. sylvaticum* | Bsyl_477 | 40.60 | 22.00 | 8.82 | 9.10 | 33.45 | 692.33 | 23.95 | -3.26 | 27.20 | 5.24 | 16.91 | 17.36 | 0.48 | 595 | 62 | 39 | 17.75 | 172 | 120 | 127 | 136 |
| *B. sylvaticum* | Bsyl_478 | 53.70 | 18.40 | 7.53 | 7.55 | 28.36 | 719.30 | 22.00 | -4.64 | 26.64 | 16.39 | -0.44 | 16.39 | -1.26 | 597 | 82 | 26 | 33.81 | 218 | 91 | 218 | 104 |
| *B. sylvaticum* | Bsyl_479 | 52.70 | 23.90 | 6.47 | 7.74 | 27.38 | 769.63 | 21.46 | -6.82 | 28.28 | 15.80 | -2.23 | 15.80 | -3.09 | 597 | 82 | 29 | 33.34 | 220 | 92 | 220 | 104 |
| *B. sylvaticum* | Bsyl_480 | 50.20 | 19.50 | 7.57 | 8.92 | 31.46 | 734.00 | 22.41 | -5.94 | 28.35 | 16.47 | -0.41 | 16.47 | -1.57 | 764 | 105 | 36 | 38.72 | 299 | 117 | 299 | 123 |
| *B. sylvaticum* | Bsyl_481 | 39.80 | -8.50 | 15.51 | 10.08 | 43.62 | 459.60 | 28.38 | 5.28 | 23.10 | 10.88 | 21.12 | 21.41 | 10.11 | 854 | 122 | 10 | 54.92 | 345 | 56 | 60 | 345 |
| *B. sylvaticum* | Bsyl_482 | 38.40 | -7.30 | 16.11 | 11.60 | 44.81 | 504.79 | 31.04 | 5.15 | 25.90 | 11.16 | 22.31 | 22.68 | 10.28 | 545 | 87 | 4 | 58.41 | 232 | 29 | 34 | 218 |
| *B. sylvaticum* | Bsyl_483 | 38.80 | -9.40 | 15.66 | 6.84 | 40.99 | 346.20 | 24.66 | 7.98 | 16.68 | 12.27 | 19.73 | 20.17 | 11.58 | 715 | 112 | 8 | 60.73 | 315 | 39 | 46 | 297 |
| *B. sylvaticum* | Bsyl_484 | 39.30 | -8.00 | 16.16 | 10.48 | 42.55 | 504.90 | 30.27 | 5.65 | 24.62 | 11.05 | 22.45 | 22.66 | 10.29 | 710 | 105 | 5 | 58.51 | 299 | 35 | 44 | 288 |
| *B. sylvaticum* | Bsyl_485 | 37.50 | -8.70 | 15.84 | 8.78 | 44.55 | 382.81 | 26.86 | 7.16 | 19.70 | 12.23 | 20.43 | 20.76 | 11.36 | 573 | 99 | 2 | 68.15 | 268 | 20 | 26 | 251 |
| *B. sylvaticum* | Bsyl_486 | 42.10 | -8.20 | 11.84 | 8.97 | 40.01 | 489.88 | 24.67 | 2.24 | 22.43 | 6.87 | 17.97 | 18.09 | 6.10 | 1508 | 210 | 27 | 48.20 | 577 | 132 | 148 | 571 |
| *B. sylvaticum* | Bsyl_487 | 39.40 | -7.30 | 15.08 | 9.18 | 36.35 | 579.91 | 30.02 | 4.76 | 25.26 | 9.43 | 22.56 | 22.72 | 8.59 | 700 | 104 | 6 | 56.77 | 294 | 39 | 47 | 276 |
| *B. sylvaticum* | Bsyl_488 | 40.30 | -8.40 | 15.49 | 10.81 | 46.84 | 434.28 | 28.30 | 5.24 | 23.07 | 10.33 | 20.73 | 21.04 | 10.33 | 1009 | 144 | 15 | 51.95 | 406 | 79 | 79 | 406 |
| *B. sylvaticum* | Bsyl_489 | 40.10 | -7.10 | 14.65 | 9.81 | 38.30 | 572.98 | 29.46 | 3.84 | 25.62 | 8.92 | 22.06 | 22.09 | 8.21 | 827 | 121 | 9 | 56.89 | 347 | 48 | 55 | 332 |
| *B. sylvaticum* | Bsyl_490 | 38.90 | -8.30 | 16.51 | 10.69 | 44.32 | 475.58 | 30.35 | 6.22 | 24.13 | 11.71 | 22.34 | 22.60 | 10.91 | 662 | 105 | 4 | 61.77 | 290 | 29 | 37 | 276 |
| *B. sylvaticum* | Bsyl_491 | 37.20 | -7.90 | 16.04 | 10.14 | 41.63 | 497.00 | 30.08 | 5.71 | 24.37 | 11.47 | 22.35 | 22.63 | 10.50 | 594 | 114 | 2 | 70.95 | 285 | 19 | 26 | 267 |
| *B. sylvaticum* | Bsyl_492 | 40.90 | -8.20 | 13.40 | 8.98 | 40.88 | 463.12 | 25.84 | 3.88 | 21.96 | 8.01 | 19.19 | 19.28 | 8.01 | 1425 | 211 | 19 | 56.35 | 607 | 99 | 104 | 607 |
| *B. sylvaticum* | Bsyl_493 | 41.30 | -8.50 | 14.20 | 9.22 | 45.27 | 405.15 | 25.16 | 4.79 | 20.37 | 9.45 | 19.20 | 19.31 | 9.45 | 1381 | 201 | 20 | 54.08 | 571 | 101 | 111 | 571 |
| *B. sylvaticum* | Bsyl_494 | 39.90 | -7.80 | 14.05 | 9.27 | 37.49 | 557.33 | 28.59 | 3.88 | 24.71 | 8.46 | 21.24 | 21.25 | 7.76 | 1070 | 159 | 10 | 58.12 | 448 | 60 | 69 | 442 |
| *B. sylvaticum* | Bsyl_495 | 39.40 | -8.90 | 15.68 | 7.96 | 41.06 | 403.62 | 26.39 | 7.00 | 19.39 | 11.73 | 20.55 | 20.90 | 10.95 | 728 | 108 | 6 | 59.92 | 312 | 37 | 44 | 298 |
| *B. sylvaticum* | Bsyl_496 | 41.80 | -7.60 | 10.96 | 9.18 | 38.14 | 536.28 | 25.07 | 1.00 | 24.07 | 5.68 | 17.86 | 17.92 | 4.91 | 1299 | 173 | 23 | 48.43 | 496 | 108 | 120 | 493 |
| *B. sylvaticum* | Bsyl_497 | 41.80 | -6.90 | 11.56 | 10.30 | 38.76 | 569.86 | 26.69 | 0.11 | 26.58 | 5.90 | 18.89 | 18.95 | 5.12 | 935 | 139 | 21 | 49.42 | 379 | 89 | 96 | 365 |
| *B. sylvaticum* | Bsyl_498 | 37.80 | 42.60 | 8.67 | 10.20 | 28.05 | 951.30 | 28.83 | -7.52 | 36.35 | 1.33 | 20.45 | 20.45 | -2.68 | 805 | 124 | 5 | 66.39 | 339 | 20 | 20 | 287 |
| *B. sylvaticum* | Bsyl_499 | 44.40 | 9.90 | 11.20 | 9.83 | 35.82 | 645.18 | 26.70 | -0.74 | 27.44 | 7.56 | 19.37 | 19.37 | 3.61 | 1365 | 181 | 44 | 33.67 | 488 | 209 | 209 | 362 |
| *B. sylvaticum* | Bsyl_500 | 42.90 | 11.10 | 14.68 | 10.08 | 37.63 | 620.68 | 29.85 | 3.05 | 26.80 | 15.81 | 22.54 | 22.54 | 7.59 | 632 | 87 | 20 | 36.71 | 239 | 87 | 87 | 160 |
| *B. sylvaticum* | Bsyl_501 | 44.20 | 10.50 | 6.94 | 6.55 | 28.72 | 608.64 | 19.62 | -3.18 | 22.80 | 3.97 | 14.78 | 14.78 | 0.05 | 1656 | 210 | 68 | 30.24 | 576 | 265 | 265 | 425 |
| *B. sylvaticum* | Bsyl_502 | 43.00 | 11.80 | 12.20 | 7.92 | 31.01 | 650.08 | 26.81 | 1.26 | 25.55 | 9.03 | 20.39 | 20.39 | 4.69 | 630 | 79 | 28 | 26.26 | 211 | 107 | 107 | 157 |
| *B. sylvaticum* | Bsyl_503 | 42.40 | 11.40 | 15.40 | 9.52 | 36.73 | 615.01 | 29.99 | 4.06 | 25.93 | 12.48 | 23.19 | 23.23 | 8.45 | 541 | 74 | 10 | 42.09 | 199 | 57 | 93 | 166 |
| *B. sylvaticum* | Bsyl_504 | 42.80 | 10.20 | 15.70 | 4.99 | 25.53 | 546.86 | 26.60 | 7.04 | 19.56 | 13.44 | 22.53 | 22.69 | 9.49 | 546 | 76 | 9 | 43.19 | 211 | 58 | 86 | 161 |
| *B. sylvaticum* | Bsyl_505 | 43.30 | 10.70 | 14.60 | 9.34 | 35.78 | 616.06 | 29.21 | 3.11 | 26.10 | 15.64 | 22.36 | 22.36 | 7.42 | 691 | 95 | 22 | 36.65 | 263 | 103 | 103 | 161 |
| *B. sylvaticum* | Bsyl_506 | 43.70 | 11.70 | 10.01 | 7.21 | 29.69 | 634.48 | 23.84 | -0.45 | 24.30 | 6.90 | 18.05 | 18.05 | 2.71 | 1517 | 204 | 60 | 31.93 | 538 | 230 | 230 | 430 |
| *B. sylvaticum* | Bsyl_507 | 44.10 | 11.30 | 10.93 | 8.70 | 31.94 | 674.58 | 26.46 | -0.78 | 27.24 | 7.30 | 19.43 | 19.43 | 3.03 | 1396 | 188 | 46 | 33.39 | 499 | 210 | 210 | 380 |
| *B. sylvaticum* | Bsyl_508 | 43.70 | 10.40 | 14.40 | 10.64 | 38.70 | 626.58 | 29.64 | 2.15 | 27.49 | 15.35 | 22.30 | 22.30 | 7.07 | 899 | 137 | 24 | 41.91 | 356 | 125 | 125 | 202 |
| *B. sylvaticum* | Bsyl_509 | 55.50 | 14.00 | 7.78 | 6.81 | 28.74 | 633.48 | 21.24 | -2.44 | 23.68 | 8.58 | 2.67 | 15.84 | 0.43 | 654 | 71 | 38 | 22.02 | 210 | 126 | 158 | 160 |
| *B. sylvaticum* | Bsyl_510 | 59.90 | 16.60 | 5.73 | 8.63 | 28.84 | 777.22 | 22.33 | -7.59 | 29.92 | 15.69 | 0.08 | 15.69 | -3.51 | 613 | 75 | 28 | 28.89 | 206 | 103 | 206 | 116 |
| *B. sylvaticum* | Bsyl_511 | 59.60 | 17.80 | 6.07 | 7.62 | 27.15 | 741.68 | 21.56 | -6.52 | 28.08 | 14.46 | 0.15 | 15.66 | -2.54 | 542 | 73 | 25 | 30.92 | 189 | 89 | 185 | 102 |
| *B. sylvaticum* | Bsyl_512 | 56.00 | 14.00 | 7.24 | 7.51 | 31.36 | 624.31 | 21.16 | -2.77 | 23.94 | 7.81 | 2.30 | 15.25 | 0.04 | 688 | 68 | 41 | 18.51 | 203 | 132 | 187 | 164 |
| *B. sylvaticum* | Bsyl_513 | 58.60 | 15.90 | 6.62 | 7.68 | 27.74 | 719.81 | 21.96 | -5.73 | 27.69 | 14.79 | 0.75 | 15.92 | -1.68 | 574 | 69 | 30 | 26.11 | 189 | 100 | 185 | 121 |
| *B. sylvaticum* | Bsyl_514 | 38.40 | -4.00 | 15.30 | 12.34 | 38.56 | 694.45 | 33.83 | 1.83 | 32.00 | 11.16 | 24.23 | 24.34 | 7.45 | 440 | 65 | 5 | 51.16 | 171 | 32 | 34 | 157 |
| *B. sylvaticum* | Bsyl_515 | 41.10 | -3.90 | 10.51 | 10.59 | 36.37 | 673.46 | 27.57 | -1.55 | 29.12 | 6.61 | 19.21 | 19.27 | 2.88 | 508 | 62 | 19 | 32.64 | 164 | 72 | 80 | 139 |
| *B. sylvaticum* | Bsyl_516 | 34.10 | -5.50 | 16.75 | 11.73 | 41.16 | 578.61 | 32.94 | 4.43 | 28.50 | 10.10 | 23.96 | 24.41 | 10.10 | 600 | 89 | 1 | 66.58 | 248 | 16 | 17 | 248 |
| *B. sylvaticum* | Bsyl_517 | 39.00 | -5.30 | 15.61 | 12.86 | 41.03 | 650.05 | 33.80 | 2.46 | 31.34 | 8.99 | 23.97 | 24.01 | 8.23 | 538 | 82 | 6 | 54.36 | 222 | 38 | 38 | 203 |
| *B. sylvaticum* | Bsyl_518 | 40.70 | -7.60 | 13.62 | 9.62 | 41.18 | 495.05 | 27.17 | 3.80 | 23.37 | 7.99 | 19.89 | 19.97 | 7.99 | 1512 | 233 | 20 | 59.07 | 665 | 100 | 107 | 665 |
| *B. sylvaticum* | Bsyl_519 | 37.40 | 22.10 | 14.82 | 11.44 | 40.33 | 635.39 | 30.78 | 2.42 | 28.36 | 8.86 | 22.90 | 22.90 | 7.54 | 765 | 131 | 12 | 68.53 | 363 | 41 | 41 | 331 |
| *B. sylvaticum* | Bsyl_520 | 42.90 | -5.90 | 8.65 | 9.55 | 36.90 | 592.65 | 23.68 | -2.22 | 25.90 | 2.85 | 16.27 | 16.32 | 1.89 | 1018 | 121 | 43 | 31.17 | 348 | 149 | 153 | 326 |
| *B. sylvaticum* | Bsyl_521 | 41.10 | -4.50 | 11.60 | 11.85 | 39.79 | 642.03 | 29.09 | -0.70 | 29.79 | 7.80 | 19.69 | 19.90 | 4.24 | 396 | 50 | 14 | 33.06 | 126 | 61 | 66 | 108 |
| *B. sylvaticum* | Bsyl_522 | 41.80 | -6.10 | 11.22 | 10.75 | 38.14 | 608.11 | 27.60 | -0.58 | 28.19 | 5.11 | 19.11 | 19.11 | 4.35 | 793 | 110 | 21 | 43.91 | 301 | 87 | 87 | 296 |
| *B. sylvaticum* | Bsyl_523 | 43.30 | -4.50 | 12.92 | 9.03 | 43.73 | 431.31 | 23.55 | 2.90 | 20.66 | 10.78 | 18.22 | 18.41 | 7.83 | 869 | 97 | 47 | 25.40 | 277 | 143 | 153 | 244 |
| *B. sylvaticum* | Bsyl_524 | 40.30 | -5.10 | 7.41 | 8.56 | 32.27 | 646.70 | 23.47 | -3.04 | 26.52 | 1.48 | 16.04 | 16.04 | 0.46 | 976 | 130 | 21 | 44.03 | 361 | 91 | 91 | 334 |
| *B. sylvaticum* | Bsyl_525 | 58.10 | 11.70 | 7.25 | 6.87 | 28.02 | 648.67 | 20.60 | -3.90 | 24.50 | 7.98 | 2.02 | 15.45 | -0.41 | 819 | 93 | 45 | 23.39 | 266 | 151 | 202 | 199 |
| *B. sylvaticum* | Bsyl_526 | 57.00 | 16.20 | 6.41 | 9.02 | 32.58 | 680.78 | 22.25 | -5.44 | 27.69 | 14.17 | 1.06 | 15.27 | -1.42 | 568 | 62 | 35 | 19.65 | 176 | 107 | 169 | 126 |
| *B. sylvaticum* | Bsyl_527 | 58.90 | 17.70 | 6.57 | 6.32 | 24.91 | 689.02 | 20.37 | -5.02 | 25.38 | 7.66 | 0.54 | 15.51 | -1.18 | 559 | 61 | 28 | 26.47 | 176 | 96 | 162 | 103 |
| *B. sylvaticum* | Bsyl_528 | 56.90 | 18.20 | 7.37 | 4.71 | 21.65 | 636.09 | 19.68 | -2.07 | 21.75 | 8.62 | 4.74 | 15.60 | 0.28 | 513 | 55 | 30 | 20.12 | 157 | 96 | 134 | 110 |
| *B. sylvaticum* | Bsyl_529 | 51.40 | 5.50 | 9.86 | 8.48 | 36.52 | 556.58 | 23.00 | -0.22 | 23.23 | 4.26 | 5.83 | 16.81 | 3.24 | 768 | 74 | 46 | 12.65 | 216 | 166 | 194 | 196 |
| *B. sylvaticum* | Bsyl_530 | 49.90 | 3.60 | 9.73 | 8.35 | 35.83 | 555.24 | 23.24 | -0.07 | 23.30 | 6.62 | 5.83 | 16.68 | 3.13 | 749 | 71 | 53 | 9.96 | 207 | 170 | 186 | 187 |
| *B. sylvaticum* | Bsyl_531 | 48.60 | -2.20 | 11.93 | 6.27 | 35.07 | 436.91 | 21.98 | 4.12 | 17.87 | 7.77 | 17.31 | 17.45 | 6.87 | 773 | 93 | 41 | 27.58 | 264 | 129 | 145 | 238 |
| *B. sylvaticum* | Bsyl_533 | 53.90 | 33.70 | 5.07 | 8.04 | 23.57 | 966.82 | 22.41 | -11.68 | 34.10 | 16.58 | -5.87 | 16.58 | -6.83 | 629 | 82 | 28 | 32.57 | 231 | 101 | 231 | 111 |
| *B. sylvaticum* | Bsyl_534 | 44.80 | 8.70 | 13.04 | 8.82 | 30.26 | 747.39 | 28.77 | -0.39 | 29.16 | 13.67 | 22.26 | 22.26 | 3.82 | 826 | 125 | 33 | 31.99 | 282 | 151 | 151 | 179 |
| *B. sylvaticum* | Bsyl_535 | 46.90 | 15.50 | 8.95 | 9.67 | 32.25 | 772.90 | 24.47 | -5.51 | 29.98 | 18.20 | -0.85 | 18.20 | -0.85 | 895 | 122 | 28 | 43.71 | 362 | 109 | 362 | 109 |
| *B. sylvaticum* | Bsyl_536 | 46.60 | 14.50 | 8.20 | 10.30 | 32.60 | 798.16 | 24.62 | -6.98 | 31.60 | 17.74 | -0.02 | 17.74 | -1.99 | 1041 | 128 | 40 | 33.59 | 363 | 146 | 363 | 146 |
| *B. sylvaticum* | Bsyl_537 | 47.70 | 14.70 | 7.28 | 9.34 | 32.59 | 724.05 | 22.29 | -6.37 | 28.66 | 16.07 | -1.70 | 16.07 | -1.70 | 884 | 127 | 43 | 37.65 | 344 | 139 | 344 | 139 |
| *B. sylvaticum* | Bsyl_538 | 45.20 | 13.60 | 14.22 | 8.46 | 31.63 | 675.84 | 28.53 | 1.80 | 26.73 | 15.11 | 6.81 | 22.78 | 6.08 | 819 | 94 | 47 | 21.75 | 267 | 165 | 190 | 179 |
| *B. sylvaticum* | Bsyl_539 | 54.20 | -7.40 | 9.41 | 6.70 | 39.45 | 393.43 | 18.99 | 2.00 | 16.99 | 7.35 | 10.73 | 14.48 | 4.96 | 1046 | 110 | 62 | 20.65 | 325 | 194 | 233 | 291 |
| *B. sylvaticum* | Bsyl_540 | 50.60 | -4.90 | 10.88 | 4.94 | 34.56 | 347.41 | 18.80 | 4.50 | 14.30 | 8.17 | 13.63 | 15.42 | 7.05 | 927 | 113 | 49 | 29.67 | 326 | 159 | 198 | 252 |
| *B. sylvaticum* | Bsyl_541 | 52.10 | -1.40 | 9.36 | 7.97 | 38.29 | 471.64 | 21.48 | 0.67 | 20.81 | 4.90 | 5.70 | 15.45 | 4.03 | 693 | 67 | 47 | 11.66 | 196 | 150 | 169 | 179 |
| *B. sylvaticum* | Bsyl_542 | 52.50 | -0.60 | 9.35 | 7.63 | 37.36 | 469.97 | 21.11 | 0.70 | 20.41 | 4.92 | 5.66 | 15.39 | 4.01 | 644 | 60 | 43 | 10.65 | 177 | 139 | 168 | 161 |
| *B. sylvaticum* | Bsyl_543 | 52.10 | -0.10 | 9.75 | 8.00 | 37.62 | 487.30 | 21.98 | 0.72 | 21.26 | 10.53 | 5.94 | 15.91 | 4.16 | 576 | 54 | 36 | 11.84 | 160 | 124 | 144 | 139 |
| *B. sylvaticum* | Bsyl_544 | 55.40 | 37.00 | 4.65 | 7.79 | 23.38 | 961.60 | 22.31 | -10.99 | 33.30 | 16.37 | -1.19 | 16.37 | -7.23 | 632 | 85 | 29 | 34.16 | 233 | 94 | 233 | 116 |
| *B. sylvaticum* | Bsyl_545 | 55.30 | -5.60 | 9.30 | 6.01 | 39.24 | 359.17 | 17.77 | 2.45 | 15.32 | 6.06 | 10.27 | 13.81 | 5.24 | 1109 | 126 | 53 | 27.42 | 368 | 179 | 264 | 332 |
| *B. sylvaticum* | Bsyl_546 | 58.40 | -6.20 | 7.94 | 4.67 | 35.54 | 323.88 | 15.11 | 1.96 | 13.15 | 6.52 | 8.63 | 12.11 | 4.34 | 1317 | 156 | 61 | 31.48 | 448 | 203 | 288 | 402 |
| *B. sylvaticum* | Bsyl_547 | 57.50 | -7.30 | 8.94 | 4.88 | 36.15 | 331.44 | 16.32 | 2.82 | 13.51 | 7.52 | 9.73 | 13.15 | 5.30 | 1401 | 163 | 64 | 30.39 | 475 | 214 | 267 | 414 |
| *B. sylvaticum* | Bsyl_548 | 44.90 | 6.60 | 5.25 | 8.93 | 33.91 | 643.59 | 20.20 | -6.13 | 26.33 | 2.03 | 13.38 | 13.38 | -2.07 | 1216 | 130 | 64 | 18.94 | 377 | 232 | 232 | 336 |
| *B. sylvaticum* | Bsyl_549 | 37.00 | -1.90 | 17.86 | 11.00 | 42.34 | 543.25 | 31.97 | 5.98 | 25.99 | 15.26 | 24.68 | 24.98 | 11.73 | 235 | 33 | 2 | 50.80 | 93 | 15 | 19 | 78 |
| *B. sylvaticum* | Bsyl_550 | 37.50 | -4.40 | 14.91 | 13.10 | 41.21 | 653.43 | 33.48 | 1.70 | 31.78 | 8.45 | 23.36 | 23.45 | 7.59 | 594 | 89 | 4 | 58.02 | 248 | 31 | 33 | 233 |
| *B. sylvaticum* | Bsyl_551 | 38.60 | -4.90 | 15.37 | 12.93 | 40.36 | 670.78 | 33.97 | 1.93 | 32.04 | 8.51 | 24.01 | 24.03 | 7.74 | 531 | 81 | 6 | 54.77 | 218 | 36 | 37 | 198 |
| *B. sylvaticum* | Bsyl_552 | 58.70 | 7.60 | 4.50 | 6.39 | 25.99 | 673.66 | 18.16 | -6.42 | 24.58 | 5.11 | 7.58 | 13.11 | -3.31 | 1559 | 195 | 70 | 31.89 | 549 | 252 | 323 | 422 |
| *B. sylvaticum* | Bsyl_553 | 58.70 | 8.80 | 6.14 | 7.07 | 27.12 | 699.78 | 20.56 | -5.49 | 26.05 | 6.43 | 9.64 | 15.07 | -2.15 | 1191 | 154 | 60 | 30.59 | 434 | 209 | 270 | 268 |
| *B. sylvaticum* | Bsyl_554 | 59.50 | 10.70 | 6.46 | 7.35 | 27.06 | 727.06 | 21.32 | -5.84 | 27.16 | 6.68 | 1.06 | 15.77 | -2.12 | 795 | 98 | 41 | 26.14 | 270 | 145 | 208 | 165 |
| *B. sylvaticum* | Bsyl_555 | 62.10 | 5.20 | 5.46 | 3.84 | 24.70 | 454.20 | 13.96 | -1.60 | 15.56 | 6.26 | 6.96 | 11.18 | 0.25 | 2548 | 303 | 116 | 32.42 | 888 | 386 | 497 | 719 |
| *B. sylvaticum* | Bsyl_556 | 62.90 | 7.00 | 6.50 | 4.91 | 25.98 | 541.53 | 16.83 | -2.07 | 18.90 | 6.95 | 8.81 | 13.39 | 0.13 | 1830 | 217 | 82 | 31.02 | 621 | 285 | 355 | 523 |
| *B. sylvaticum* | Bsyl_557 | 59.00 | 5.70 | 7.70 | 6.21 | 31.48 | 511.19 | 18.38 | -1.34 | 19.72 | 8.46 | 9.61 | 14.13 | 1.89 | 1297 | 162 | 58 | 32.79 | 461 | 195 | 287 | 340 |
| *B. sylvaticum* | Bsyl_558 | 59.40 | 8.40 | 4.52 | 8.04 | 29.21 | 702.44 | 20.00 | -7.53 | 27.53 | 4.57 | -0.66 | 13.61 | -3.64 | 950 | 115 | 47 | 26.28 | 317 | 164 | 257 | 201 |
| *B. sylvaticum* | Bsyl_559 | 42.20 | -6.40 | 8.25 | 9.27 | 35.44 | 601.81 | 23.65 | -2.50 | 26.15 | 2.49 | 16.13 | 16.13 | 1.60 | 1072 | 143 | 34 | 40.02 | 398 | 132 | 132 | 379 |
| *B. sylvaticum* | Bsyl_560 | 43.10 | -8.30 | 12.57 | 8.71 | 45.36 | 386.49 | 23.39 | 4.20 | 19.20 | 8.99 | 17.36 | 17.66 | 8.25 | 1570 | 224 | 38 | 45.31 | 609 | 159 | 194 | 607 |
| *B. sylvaticum* | Bsyl_561 | 38.60 | -2.60 | 13.95 | 14.15 | 40.11 | 756.18 | 34.82 | -0.46 | 35.28 | 9.43 | 23.73 | 23.96 | 5.60 | 427 | 56 | 7 | 44.91 | 154 | 37 | 42 | 141 |
| *B. sylvaticum* | Bsyl_562 | 38.50 | -2.00 | 14.25 | 13.74 | 40.68 | 719.60 | 34.06 | 0.28 | 33.78 | 10.04 | 23.56 | 23.74 | 6.31 | 387 | 47 | 9 | 38.33 | 130 | 43 | 46 | 110 |
| *B. sylvaticum* | Bsyl_563 | 40.50 | -0.40 | 7.47 | 13.76 | 42.45 | 656.25 | 26.62 | -5.80 | 32.42 | 9.22 | 0.96 | 16.11 | 0.17 | 625 | 88 | 38 | 27.72 | 217 | 122 | 145 | 131 |
| *B. sylvaticum* | Bsyl_564 | 41.80 | -4.00 | 11.45 | 11.91 | 39.71 | 649.58 | 29.12 | -0.88 | 30.00 | 7.64 | 19.77 | 19.81 | 4.01 | 402 | 47 | 14 | 31.33 | 126 | 56 | 70 | 114 |
| *B. sylvaticum* | Bsyl_565 | 40.50 | -2.60 | 13.18 | 12.33 | 38.97 | 688.28 | 31.56 | -0.07 | 31.63 | 9.11 | 22.20 | 22.20 | 5.50 | 449 | 55 | 12 | 39.31 | 159 | 54 | 54 | 134 |
| *B. sylvaticum* | Bsyl_566 | 43.00 | -6.60 | 8.41 | 8.77 | 37.82 | 531.62 | 21.90 | -1.29 | 23.19 | 3.30 | 15.23 | 15.34 | 2.47 | 1303 | 160 | 48 | 36.13 | 463 | 166 | 177 | 428 |
| *B. sylvaticum* | Bsyl_567 | 42.80 | -3.40 | 11.75 | 9.99 | 41.31 | 518.04 | 25.61 | 1.42 | 24.19 | 6.55 | 18.42 | 18.42 | 5.85 | 755 | 83 | 38 | 24.51 | 233 | 128 | 128 | 212 |
| *B. sylvaticum* | Bsyl_568 | 43.20 | -7.20 | 12.17 | 9.44 | 45.20 | 424.21 | 24.08 | 3.19 | 20.90 | 8.07 | 17.45 | 17.70 | 7.37 | 1276 | 166 | 38 | 41.09 | 469 | 136 | 165 | 441 |
| *B. sylvaticum* | Bsyl_569 | 40.10 | -5.70 | 14.11 | 10.66 | 38.81 | 597.52 | 29.59 | 2.11 | 27.48 | 8.03 | 21.86 | 21.86 | 7.33 | 533 | 71 | 8 | 46.83 | 201 | 45 | 45 | 195 |
| *B. sylvaticum* | Bsyl_570 | 41.10 | -2.70 | 11.40 | 12.61 | 40.27 | 665.68 | 29.62 | -1.68 | 31.31 | 7.52 | 20.02 | 20.08 | 3.94 | 454 | 53 | 17 | 32.42 | 149 | 64 | 71 | 124 |
| *B. sylvaticum* | Bsyl_571 | 42.50 | -5.50 | 11.93 | 10.74 | 39.30 | 599.51 | 27.33 | 0.00 | 27.33 | 5.56 | 19.39 | 19.62 | 4.91 | 538 | 66 | 22 | 31.35 | 182 | 81 | 84 | 170 |
| *B. sylvaticum* | Bsyl_572 | 41.90 | -8.80 | 14.24 | 7.64 | 43.53 | 375.08 | 23.80 | 6.25 | 17.55 | 10.46 | 18.77 | 18.91 | 9.79 | 1498 | 221 | 30 | 51.31 | 606 | 121 | 154 | 598 |
| *B. sylvaticum* | Bsyl_573 | 42.50 | -8.60 | 13.67 | 6.88 | 39.59 | 403.31 | 23.45 | 6.06 | 17.38 | 8.96 | 18.69 | 18.80 | 8.96 | 1690 | 229 | 36 | 47.67 | 659 | 150 | 181 | 659 |
| *B. sylvaticum* | Bsyl_574 | 42.90 | -4.90 | 7.83 | 8.98 | 35.83 | 578.98 | 22.48 | -2.59 | 25.08 | 2.18 | 15.34 | 15.34 | 1.28 | 891 | 107 | 37 | 30.72 | 301 | 134 | 134 | 283 |
| *B. sylvaticum* | Bsyl_575 | 41.20 | -6.60 | 12.32 | 10.17 | 38.92 | 563.33 | 27.36 | 1.23 | 26.12 | 6.02 | 19.58 | 19.71 | 6.02 | 1074 | 159 | 17 | 55.19 | 457 | 86 | 86 | 457 |
| *B. sylvaticum* | Bsyl_576 | 42.30 | -3.00 | 8.73 | 10.16 | 38.82 | 579.24 | 24.18 | -2.00 | 26.17 | 10.06 | 16.24 | 16.24 | 2.21 | 643 | 75 | 30 | 26.30 | 196 | 106 | 106 | 180 |
| *B. sylvaticum* | Bsyl_577 | 39.50 | -2.10 | 13.76 | 12.73 | 38.99 | 711.01 | 32.83 | 0.17 | 32.66 | 9.51 | 22.93 | 23.13 | 5.86 | 428 | 50 | 12 | 35.96 | 138 | 54 | 60 | 123 |
| *B. sylvaticum* | Bsyl_578 | 36.90 | -4.10 | 15.56 | 12.02 | 42.38 | 583.35 | 31.87 | 3.51 | 28.36 | 10.02 | 23.09 | 23.25 | 9.11 | 537 | 80 | 3 | 59.86 | 229 | 25 | 29 | 216 |
| *B. sylvaticum* | Bsyl_579 | 37.60 | -6.50 | 16.58 | 11.72 | 41.19 | 585.27 | 32.85 | 4.38 | 28.46 | 10.80 | 24.02 | 24.19 | 9.88 | 571 | 95 | 3 | 65.06 | 262 | 25 | 29 | 237 |
| *B. sylvaticum* | Bsyl_580 | 41.10 | -3.30 | 10.30 | 11.06 | 37.54 | 658.70 | 27.66 | -1.81 | 29.47 | 6.59 | 18.91 | 18.95 | 3.02 | 475 | 58 | 17 | 32.17 | 152 | 69 | 76 | 127 |
| *B. sylvaticum* | Bsyl_581 | 40.40 | -4.00 | 14.36 | 11.61 | 37.82 | 684.45 | 32.10 | 1.40 | 30.70 | 10.27 | 23.15 | 23.32 | 6.65 | 374 | 45 | 11 | 36.90 | 129 | 50 | 51 | 106 |
| *B. sylvaticum* | Bsyl_582 | 43.60 | -7.60 | 13.25 | 7.76 | 45.06 | 350.84 | 22.74 | 5.53 | 17.21 | 10.10 | 17.49 | 17.89 | 9.36 | 1152 | 147 | 37 | 37.70 | 409 | 131 | 164 | 397 |
| *B. sylvaticum* | Bsyl_583 | 42.80 | -9.10 | 14.28 | 5.98 | 39.23 | 338.97 | 22.65 | 7.39 | 15.25 | 11.18 | 18.35 | 18.64 | 10.31 | 1047 | 149 | 24 | 44.92 | 418 | 112 | 136 | 375 |
| *B. sylvaticum* | Bsyl_584 | 39.00 | 1.50 | 17.52 | 10.13 | 40.44 | 555.79 | 31.38 | 6.33 | 25.05 | 15.20 | 24.52 | 24.74 | 11.32 | 444 | 70 | 5 | 50.22 | 180 | 38 | 74 | 135 |
| *B. sylvaticum* | Bsyl_585 | 39.90 | -6.40 | 15.76 | 10.71 | 38.88 | 601.66 | 31.48 | 3.93 | 27.55 | 9.66 | 23.48 | 23.52 | 8.88 | 576 | 86 | 7 | 53.83 | 240 | 41 | 43 | 220 |
| *B. sylvaticum* | Bsyl_586 | 42.70 | -1.20 | 11.59 | 9.49 | 36.91 | 566.75 | 26.64 | 0.93 | 25.71 | 8.90 | 19.08 | 19.08 | 5.18 | 775 | 82 | 37 | 23.65 | 235 | 130 | 130 | 217 |
| *B. sylvaticum* | Bsyl_587 | 42.40 | -1.80 | 13.46 | 10.63 | 38.49 | 599.97 | 29.42 | 1.81 | 27.61 | 14.70 | 21.29 | 21.29 | 6.59 | 519 | 59 | 30 | 22.66 | 162 | 92 | 92 | 130 |
| *B. sylvaticum* | Bsyl_588 | 40.90 | -5.40 | 11.89 | 11.99 | 40.68 | 623.25 | 29.08 | -0.39 | 29.47 | 8.23 | 19.77 | 19.88 | 4.71 | 412 | 48 | 13 | 34.29 | 137 | 57 | 63 | 123 |
| *B. sylvaticum* | Bsyl_589 | 41.80 | -3.10 | 9.85 | 10.94 | 38.19 | 631.83 | 26.68 | -1.97 | 28.65 | 11.50 | 17.99 | 18.03 | 2.73 | 455 | 53 | 18 | 27.12 | 139 | 72 | 81 | 127 |
| *B. sylvaticum* | Bsyl_590 | 42.40 | 0.00 | 11.14 | 10.24 | 36.74 | 630.48 | 27.32 | -0.56 | 27.88 | 7.91 | 19.36 | 19.36 | 4.02 | 746 | 80 | 35 | 21.89 | 226 | 151 | 151 | 189 |
| *B. sylvaticum* | Bsyl_591 | 40.90 | -1.90 | 10.33 | 13.81 | 43.41 | 641.04 | 28.87 | -2.94 | 31.81 | 12.01 | 18.59 | 18.69 | 3.21 | 537 | 71 | 23 | 31.26 | 171 | 88 | 96 | 135 |
| *B. sylvaticum* | Bsyl_592 | 38.10 | -6.30 | 14.55 | 11.94 | 39.37 | 635.50 | 32.47 | 2.14 | 30.32 | 8.35 | 22.76 | 22.89 | 7.41 | 685 | 111 | 5 | 61.51 | 303 | 35 | 38 | 279 |
| *B. sylvaticum* | Bsyl_593 | 36.20 | -5.40 | 17.97 | 7.86 | 39.98 | 421.67 | 29.01 | 9.36 | 19.65 | 14.18 | 23.13 | 23.60 | 13.28 | 734 | 144 | 1 | 77.78 | 374 | 16 | 22 | 345 |
| *B. sylvaticum* | Bsyl_594 | 39.50 | -5.30 | 13.21 | 10.73 | 37.27 | 636.93 | 29.90 | 1.12 | 28.78 | 6.92 | 21.58 | 21.58 | 6.11 | 583 | 79 | 8 | 48.67 | 222 | 44 | 44 | 212 |
| *B. sylvaticum* | Bsyl_595 | 43.40 | -5.20 | 13.25 | 8.41 | 41.60 | 434.38 | 23.65 | 3.43 | 20.22 | 11.09 | 18.59 | 18.79 | 8.12 | 867 | 96 | 44 | 26.29 | 280 | 138 | 148 | 252 |
| *B. sylvaticum* | Bsyl_596 | 40.50 | -3.30 | 13.50 | 12.32 | 38.39 | 698.05 | 32.14 | 0.05 | 32.10 | 9.39 | 22.63 | 22.63 | 5.67 | 429 | 61 | 9 | 44.15 | 166 | 45 | 45 | 128 |
| *B. sylvaticum* | Bsyl_597 | 38.50 | -7.90 | 16.12 | 12.44 | 45.53 | 522.76 | 32.40 | 5.08 | 27.32 | 10.95 | 22.60 | 22.92 | 10.11 | 610 | 99 | 4 | 60.12 | 264 | 30 | 36 | 250 |
| *B. sylvaticum* | Bsyl_598 | 40.10 | -2.00 | 10.45 | 12.57 | 39.69 | 676.11 | 29.16 | -2.49 | 31.66 | 12.14 | 19.21 | 19.40 | 3.01 | 556 | 69 | 20 | 31.97 | 175 | 83 | 91 | 149 |
| *B. sylvaticum* | Bsyl_599 | 41.50 | -5.30 | 12.47 | 11.66 | 41.03 | 603.87 | 28.82 | 0.40 | 28.41 | 6.08 | 20.09 | 20.16 | 5.45 | 475 | 58 | 15 | 34.79 | 163 | 64 | 67 | 152 |
| *B. sylvaticum* | Bsyl_600 | 39.40 | -4.50 | 12.98 | 10.82 | 36.28 | 672.44 | 30.20 | 0.37 | 29.84 | 9.09 | 21.72 | 21.78 | 5.48 | 499 | 66 | 9 | 43.21 | 179 | 46 | 53 | 163 |
| *B. sylvaticum* | Bsyl_601 | 43.40 | -3.70 | 14.11 | 9.19 | 47.94 | 377.23 | 24.46 | 5.29 | 19.17 | 12.76 | 18.67 | 19.07 | 9.99 | 1173 | 142 | 55 | 26.25 | 381 | 196 | 215 | 321 |
| *B. sylvaticum* | Bsyl_602 | 36.80 | -5.50 | 15.07 | 10.43 | 39.94 | 551.30 | 30.17 | 4.04 | 26.13 | 9.93 | 22.07 | 22.40 | 8.89 | 749 | 121 | 1 | 69.98 | 343 | 20 | 25 | 328 |
| *B. sylvaticum* | Bsyl_603 | 40.60 | -6.10 | 11.00 | 9.83 | 37.50 | 585.68 | 26.39 | 0.19 | 26.20 | 5.41 | 18.77 | 18.77 | 4.54 | 998 | 148 | 16 | 52.62 | 404 | 77 | 77 | 403 |
| *B. sylvaticum* | Bsyl_604 | 43.30 | -1.70 | 13.07 | 7.68 | 39.11 | 442.92 | 23.72 | 4.09 | 19.64 | 8.89 | 18.35 | 18.86 | 8.09 | 1323 | 147 | 71 | 21.11 | 406 | 241 | 259 | 368 |
| *B. sylvaticum* | Bsyl_605 | 43.00 | 6.20 | 14.95 | 6.74 | 34.29 | 471.21 | 25.73 | 6.08 | 19.65 | 13.20 | 20.86 | 21.12 | 9.87 | 698 | 108 | 7 | 47.45 | 271 | 68 | 91 | 227 |
| *B. sylvaticum* | Bsyl_606 | 44.90 | 4.60 | 10.32 | 9.03 | 34.60 | 622.13 | 25.32 | -0.78 | 26.10 | 10.78 | 4.07 | 18.26 | 3.12 | 951 | 119 | 56 | 24.11 | 306 | 191 | 203 | 196 |
| *B. sylvaticum* | Bsyl_607 | 40.80 | 40.50 | 10.91 | 8.63 | 31.72 | 695.49 | 25.10 | -2.10 | 27.20 | 7.91 | 5.64 | 19.16 | 2.35 | 955 | 115 | 58 | 21.23 | 303 | 194 | 213 | 233 |
| *B. sylvaticum* | Bsyl_609 | 59.10 | -3.10 | 7.63 | 4.61 | 35.42 | 310.13 | 14.93 | 1.92 | 13.01 | 6.59 | 10.07 | 11.64 | 4.25 | 1148 | 137 | 56 | 30.16 | 390 | 180 | 255 | 318 |
| *B. sylvaticum* | Bsyl_610 | 56.70 | -6.50 | 9.48 | 4.18 | 31.86 | 336.79 | 16.75 | 3.64 | 13.11 | 6.48 | 10.30 | 13.75 | 5.73 | 1618 | 190 | 69 | 33.89 | 559 | 232 | 381 | 490 |
| *B. sylvaticum* | Bsyl_611 | 62.40 | 7.90 | -0.14 | 5.10 | 24.76 | 569.65 | 11.50 | -9.09 | 20.60 | -5.47 | 1.83 | 7.38 | -6.37 | 1377 | 166 | 50 | 31.03 | 461 | 201 | 284 | 434 |
| *B. sylvaticum* | Bsyl_612 | 51.50 | 4.50 | 10.13 | 8.01 | 35.24 | 552.99 | 22.85 | 0.14 | 22.72 | 10.61 | 6.03 | 16.99 | 3.55 | 802 | 79 | 48 | 16.43 | 235 | 162 | 206 | 193 |
| *B. sylvaticum* | Bsyl_613 | 45.50 | 0.50 | 12.23 | 9.22 | 39.38 | 539.37 | 25.42 | 2.02 | 23.40 | 6.58 | 18.99 | 18.99 | 5.96 | 899 | 95 | 52 | 18.98 | 277 | 167 | 167 | 259 |
| *B. sylvaticum* | Bsyl_614 | 50.50 | 16.40 | 6.33 | 7.84 | 30.07 | 692.44 | 20.50 | -5.55 | 26.06 | 14.73 | -1.22 | 14.73 | -2.26 | 617 | 93 | 27 | 46.14 | 262 | 87 | 262 | 91 |
| *B. sylvaticum* | Bsyl_615 | 57.90 | -6.80 | 8.03 | 4.86 | 34.74 | 350.98 | 15.65 | 1.67 | 13.98 | 6.48 | 8.91 | 12.52 | 4.20 | 1532 | 177 | 71 | 30.68 | 521 | 236 | 282 | 453 |
| *B. sylvaticum* | Bsyl_616 | 50.50 | 3.50 | 10.24 | 7.88 | 35.18 | 542.94 | 23.09 | 0.70 | 22.39 | 7.21 | 6.37 | 17.05 | 3.80 | 734 | 73 | 47 | 13.07 | 208 | 159 | 188 | 173 |
| *B. sylvaticum* | Bsyl_617 | 47.70 | -2.70 | 12.08 | 5.82 | 34.52 | 410.87 | 21.54 | 4.69 | 16.85 | 8.38 | 17.11 | 17.31 | 7.40 | 882 | 106 | 45 | 28.33 | 305 | 143 | 163 | 279 |
| *B. sylvaticum* | Bsyl_618 | 43.50 | -6.70 | 12.85 | 8.79 | 46.47 | 381.99 | 23.32 | 4.42 | 18.91 | 11.22 | 17.54 | 17.83 | 8.56 | 1098 | 132 | 41 | 35.65 | 390 | 139 | 157 | 344 |
| *B. sylvaticum* | Bsyl_620 | 34.10 | -4.20 | 12.57 | 10.44 | 37.03 | 620.44 | 28.73 | 0.53 | 28.20 | 6.42 | 20.46 | 20.54 | 5.34 | 435 | 55 | 6 | 49.08 | 154 | 32 | 34 | 147 |
| *B. sylvaticum* | Bsyl_621 | 41.70 | -0.90 | 15.12 | 10.73 | 37.83 | 645.12 | 31.02 | 2.66 | 28.36 | 17.30 | 8.86 | 23.32 | 7.60 | 357 | 49 | 19 | 28.32 | 123 | 67 | 78 | 72 |
| *B. sylvaticum* | Bsyl_622 | 59.60 | 5.20 | 7.72 | 4.25 | 26.18 | 452.10 | 16.54 | 0.29 | 16.25 | 6.12 | 8.82 | 13.35 | 2.61 | 1944 | 235 | 87 | 34.99 | 699 | 277 | 379 | 473 |
| *B. sylvaticum* | Bsyl_623 | 42.70 | 27.70 | 12.61 | 9.46 | 32.18 | 757.17 | 28.53 | -0.86 | 29.40 | 8.97 | 21.63 | 22.06 | 3.67 | 537 | 59 | 33 | 15.54 | 158 | 113 | 119 | 130 |
| *B. sylvaticum* | Bsyl_624 | 34.10 | 50.70 | 14.25 | 13.66 | 35.27 | 920.29 | 35.16 | -3.57 | 38.73 | 4.14 | 25.18 | 25.50 | 2.76 | 184 | 34 | 0 | 78.16 | 94 | 2 | 4 | 84 |
| *B. sylvaticum* | Bsyl_625 | 47.30 | 26.60 | 8.56 | 9.67 | 31.21 | 825.57 | 25.11 | -5.87 | 30.98 | 17.18 | -0.27 | 18.61 | -1.54 | 557 | 91 | 21 | 52.82 | 245 | 69 | 239 | 70 |
| *B. sylvaticum* | Bsyl_626 | 41.30 | 23.40 | 11.49 | 10.00 | 33.11 | 757.58 | 28.02 | -2.18 | 30.19 | 7.12 | 20.25 | 20.90 | 2.26 | 496 | 54 | 28 | 18.77 | 147 | 95 | 114 | 126 |
| *B. sylvaticum* | Bsyl_627 | 41.00 | 39.80 | 13.99 | 6.82 | 29.95 | 599.76 | 25.85 | 3.07 | 22.77 | 11.98 | 19.06 | 21.44 | 7.01 | 970 | 127 | 52 | 26.89 | 333 | 188 | 211 | 226 |
| *B. sylvaticum* | Bsyl_628 | 49.40 | 20.40 | 6.94 | 9.56 | 32.53 | 745.04 | 22.22 | -7.16 | 29.38 | 14.49 | 2.33 | 15.88 | -2.49 | 958 | 140 | 43 | 39.79 | 375 | 151 | 371 | 189 |
| *B. sylvaticum* | Bsyl_629 | 34.80 | -4.60 | 13.79 | 11.68 | 39.74 | 612.05 | 30.73 | 1.33 | 29.40 | 7.45 | 21.69 | 21.69 | 6.67 | 731 | 110 | 2 | 69.25 | 316 | 19 | 19 | 299 |
| *B. sylvaticum* | Bsyl_630 | 37.00 | 22.50 | 16.69 | 9.79 | 36.83 | 634.84 | 31.68 | 5.09 | 26.59 | 10.92 | 24.81 | 24.81 | 9.52 | 632 | 114 | 7 | 75.04 | 314 | 25 | 25 | 288 |
| *B. sylvaticum* | Bsyl_631 | 42.00 | 28.00 | 13.41 | 7.30 | 28.73 | 689.94 | 27.08 | 1.68 | 25.40 | 7.34 | 22.08 | 22.08 | 5.55 | 584 | 73 | 27 | 30.30 | 205 | 95 | 95 | 181 |
| *B. sylvaticum* | Bsyl_632 | 45.10 | 28.70 | 10.76 | 9.87 | 30.43 | 865.03 | 28.16 | -4.26 | 32.42 | 19.61 | 1.28 | 21.44 | 0.17 | 443 | 57 | 25 | 23.89 | 145 | 85 | 137 | 85 |
| *B. sylvaticum* | Bsyl_633 | 44.50 | 23.50 | 11.30 | 10.72 | 32.12 | 860.98 | 28.97 | -4.39 | 33.36 | 19.90 | 2.33 | 21.57 | 0.44 | 619 | 76 | 36 | 24.78 | 212 | 114 | 188 | 122 |
| *B. sylvaticum* | Bsyl_634 | 41.00 | 22.40 | 11.81 | 10.34 | 33.84 | 756.85 | 28.40 | -2.15 | 30.55 | 15.45 | 3.81 | 21.20 | 2.56 | 477 | 53 | 29 | 19.61 | 142 | 99 | 111 | 100 |
| *B. sylvaticum* | Bsyl_635 | 41.50 | 24.30 | 6.95 | 9.07 | 33.59 | 676.46 | 21.63 | -5.37 | 27.00 | 10.06 | 12.21 | 15.16 | -1.28 | 681 | 75 | 38 | 20.29 | 215 | 126 | 173 | 172 |
| *B. sylvaticum* | Bsyl_636 | 40.90 | 24.20 | 12.90 | 8.41 | 30.57 | 723.51 | 27.83 | 0.32 | 27.51 | 5.74 | 21.37 | 21.99 | 4.26 | 573 | 70 | 19 | 32.93 | 203 | 79 | 99 | 177 |
| *B. sylvaticum* | Bsyl_637 | 39.50 | 21.60 | 13.90 | 10.85 | 35.23 | 728.28 | 30.80 | 0.00 | 30.80 | 10.47 | 22.90 | 22.90 | 5.07 | 779 | 108 | 22 | 47.55 | 302 | 75 | 75 | 275 |
| *B. sylvaticum* | Bsyl_638 | 54.70 | 20.50 | 7.95 | 7.16 | 26.89 | 741.97 | 22.42 | -4.18 | 26.61 | 13.10 | 2.64 | 17.18 | -1.00 | 793 | 84 | 41 | 25.88 | 251 | 128 | 232 | 176 |
| *B. sylvaticum* | Bsyl_639 | 38.70 | 21.40 | 15.11 | 11.34 | 39.36 | 648.93 | 31.46 | 2.66 | 28.80 | 9.00 | 23.30 | 23.30 | 7.63 | 849 | 144 | 16 | 62.14 | 370 | 54 | 54 | 334 |
| *B. sylvaticum* | Bsyl_640 | 42.40 | -3.70 | 11.16 | 11.71 | 40.87 | 609.95 | 28.04 | -0.61 | 28.65 | 4.90 | 19.08 | 19.08 | 4.25 | 528 | 63 | 25 | 28.76 | 163 | 86 | 86 | 148 |
| *B. sylvaticum* | Bsyl_641 | 39.10 | 22.20 | 13.75 | 10.77 | 36.01 | 714.13 | 30.47 | 0.55 | 29.92 | 10.20 | 22.76 | 22.76 | 5.32 | 653 | 91 | 20 | 47.07 | 252 | 65 | 65 | 232 |
| *B. sylvaticum* | Bsyl_642 | 41.20 | 44.80 | 10.94 | 10.99 | 33.53 | 786.75 | 28.04 | -4.72 | 32.77 | 14.39 | 1.38 | 20.46 | 1.38 | 513 | 83 | 17 | 49.73 | 223 | 62 | 167 | 62 |
| *B. sylvaticum* | Bsyl_643 | 36.90 | 14.50 | 17.52 | 8.37 | 37.41 | 526.56 | 29.86 | 7.48 | 22.38 | 16.00 | 23.85 | 24.25 | 11.64 | 421 | 75 | 1 | 74.26 | 208 | 12 | 39 | 135 |
| *B. sylvaticum* | Bsyl_644 | 39.60 | 27.00 | 15.35 | 9.90 | 34.72 | 705.92 | 31.15 | 2.62 | 28.52 | 8.68 | 24.26 | 24.26 | 7.14 | 689 | 138 | 8 | 73.22 | 345 | 35 | 35 | 330 |
| *B. sylvaticum* | Bsyl_645 | 36.80 | 28.80 | 18.70 | 10.31 | 38.24 | 644.56 | 33.70 | 6.74 | 26.96 | 11.25 | 26.79 | 26.79 | 11.25 | 1023 | 230 | 4 | 93.79 | 595 | 19 | 19 | 595 |
| *B. sylvaticum* | Bsyl_647 | 54.80 | -1.30 | 8.78 | 7.02 | 39.72 | 396.54 | 18.98 | 1.32 | 17.67 | 7.10 | 9.49 | 13.93 | 4.35 | 714 | 71 | 44 | 16.55 | 209 | 145 | 188 | 183 |
| *B. sylvaticum* | Bsyl_648 | 53.30 | -4.50 | 10.04 | 6.01 | 37.06 | 384.64 | 18.99 | 2.78 | 16.21 | 8.52 | 13.36 | 14.90 | 5.68 | 879 | 108 | 46 | 26.62 | 299 | 153 | 204 | 246 |
| *B. sylvaticum* | Bsyl_649 | 51.90 | -3.20 | 9.11 | 6.64 | 35.76 | 443.56 | 19.56 | 0.99 | 18.57 | 5.10 | 13.17 | 14.74 | 4.06 | 1153 | 131 | 61 | 25.05 | 386 | 205 | 222 | 355 |
| *B. sylvaticum* | Bsyl_650 | 52.80 | -4.00 | 7.58 | 6.25 | 37.72 | 393.79 | 16.99 | 0.42 | 16.57 | 4.08 | 8.64 | 12.62 | 3.25 | 1758 | 199 | 98 | 24.97 | 580 | 316 | 348 | 516 |
| *B. sylvaticum* | Bsyl_651 | 52.40 | -3.30 | 7.87 | 7.07 | 37.43 | 429.90 | 18.66 | -0.22 | 18.88 | 3.97 | 11.94 | 13.41 | 3.01 | 1426 | 171 | 77 | 28.31 | 496 | 243 | 265 | 457 |
| *B. sylvaticum* | Bsyl_652 | 46.50 | 10.30 | 2.89 | 9.72 | 35.36 | 650.06 | 17.20 | -10.27 | 27.47 | 10.87 | -4.10 | 10.87 | -4.95 | 521 | 69 | 21 | 40.06 | 199 | 73 | 199 | 74 |
| *B. sylvaticum* | Bsyl_653 | 49.20 | 19.20 | 6.29 | 10.15 | 34.09 | 735.90 | 22.00 | -7.77 | 29.77 | 15.14 | 1.67 | 15.14 | -2.96 | 923 | 108 | 56 | 23.99 | 311 | 176 | 311 | 188 |
| *B. sylvaticum* | Bsyl_654 | 50.90 | -1.90 | 9.95 | 8.05 | 40.58 | 443.01 | 21.26 | 1.42 | 19.84 | 5.89 | 14.00 | 15.63 | 4.98 | 793 | 95 | 40 | 26.12 | 268 | 145 | 149 | 245 |
| *B. sylvaticum* | Bsyl_655 | 50.80 | -2.80 | 9.92 | 7.08 | 38.74 | 417.89 | 20.32 | 2.05 | 18.27 | 6.22 | 10.91 | 15.27 | 5.27 | 949 | 106 | 62 | 21.09 | 307 | 186 | 195 | 290 |
| *B. sylvaticum* | Bsyl_656 | 45.80 | -1.20 | 13.02 | 7.44 | 36.26 | 498.04 | 24.43 | 3.90 | 20.53 | 10.54 | 19.11 | 19.20 | 7.13 | 804 | 98 | 36 | 30.83 | 278 | 120 | 146 | 252 |
| *B. sylvaticum* | Bsyl_657 | 60.50 | 17.50 | 5.36 | 7.95 | 27.89 | 735.29 | 21.19 | -7.31 | 28.50 | 13.81 | -0.37 | 14.92 | -3.18 | 617 | 77 | 34 | 27.27 | 212 | 109 | 198 | 128 |
| *B. sylvaticum* | Bsyl_658 | 39.30 | 16.30 | 13.93 | 7.07 | 30.96 | 591.45 | 26.93 | 4.10 | 22.83 | 11.47 | 21.34 | 21.41 | 7.21 | 941 | 136 | 17 | 55.23 | 393 | 70 | 99 | 353 |
| *B. sylvaticum* | Bsyl_659 | 43.00 | 12.90 | 10.39 | 7.98 | 30.59 | 671.76 | 25.20 | -0.89 | 26.09 | 11.27 | 3.24 | 18.95 | 2.65 | 862 | 94 | 49 | 19.53 | 273 | 167 | 202 | 176 |
| *B. sylvaticum* | Bsyl_660 | 43.50 | 13.50 | 13.84 | 9.87 | 35.64 | 665.63 | 29.10 | 1.41 | 27.69 | 14.80 | 6.73 | 22.22 | 5.98 | 703 | 78 | 44 | 19.03 | 222 | 145 | 168 | 150 |
| *B. sylvaticum* | Bsyl_661 | 39.90 | 16.40 | 11.89 | 6.36 | 28.06 | 612.85 | 24.80 | 2.12 | 22.67 | 9.20 | 19.72 | 19.72 | 5.06 | 867 | 122 | 20 | 45.94 | 346 | 92 | 92 | 302 |
| *B. sylvaticum* | Bsyl_662 | 56.50 | 9.80 | 7.48 | 7.26 | 30.48 | 606.01 | 19.84 | -3.96 | 23.81 | 12.01 | 2.49 | 15.07 | 0.43 | 681 | 72 | 37 | 23.17 | 213 | 120 | 188 | 161 |
| *B. sylvaticum* | Bsyl_663 | 55.40 | 9.10 | 7.89 | 7.16 | 31.87 | 576.58 | 20.60 | -1.88 | 22.48 | 8.76 | 3.21 | 15.06 | 1.13 | 882 | 101 | 48 | 24.79 | 289 | 156 | 226 | 205 |
| *B. sylvaticum* | Bsyl_665 | 48.00 | -3.40 | 11.32 | 6.99 | 38.40 | 423.97 | 21.57 | 3.36 | 18.21 | 7.29 | 16.61 | 16.63 | 6.44 | 995 | 123 | 46 | 31.48 | 356 | 154 | 177 | 326 |
| *B. sylvaticum* | Bsyl_666 | 53.50 | -2.80 | 9.48 | 6.99 | 37.80 | 441.62 | 19.72 | 1.24 | 18.49 | 7.31 | 10.97 | 15.08 | 4.41 | 861 | 95 | 53 | 21.38 | 277 | 171 | 199 | 218 |
| *B. sylvaticum* | Bsyl_667 | 51.00 | 0.50 | 10.12 | 7.02 | 36.15 | 463.39 | 21.03 | 1.60 | 19.43 | 8.03 | 14.24 | 15.94 | 4.83 | 766 | 88 | 46 | 25.87 | 255 | 150 | 154 | 214 |
| *B. sylvaticum* | Bsyl_668 | 50.80 | -0.50 | 10.12 | 8.09 | 40.08 | 457.46 | 21.54 | 1.36 | 20.19 | 7.91 | 14.35 | 15.91 | 4.86 | 733 | 83 | 42 | 27.02 | 246 | 133 | 140 | 215 |
| *B. sylvaticum* | Bsyl_669 | 54.00 | 43.90 | 4.60 | 8.48 | 21.93 | 1129.53 | 24.60 | -14.04 | 38.65 | 18.04 | -8.21 | 18.04 | -9.39 | 501 | 66 | 24 | 31.08 | 177 | 79 | 177 | 90 |
| *B. sylvaticum* | Bsyl_670 | 51.30 | 87.70 | -0.18 | 11.24 | 27.51 | 1133.69 | 21.08 | -19.77 | 40.85 | 13.46 | -12.26 | 13.46 | -14.36 | 566 | 98 | 12 | 67.55 | 273 | 38 | 273 | 45 |
| *B. sylvaticum* | Bsyl_671 | 54.00 | 39.00 | 4.89 | 8.55 | 23.67 | 1052.25 | 23.82 | -12.28 | 36.10 | 17.58 | -7.06 | 17.58 | -7.93 | 548 | 78 | 26 | 32.73 | 200 | 87 | 200 | 98 |
| *B. sylvaticum* | Bsyl_672 | 53.30 | 45.70 | 4.27 | 9.30 | 23.16 | 1157.83 | 25.14 | -15.03 | 40.17 | 17.85 | -3.81 | 17.85 | -9.92 | 549 | 66 | 28 | 25.15 | 181 | 92 | 181 | 112 |
| *B. sylvaticum* | Bsyl_673 | 54.60 | 40.70 | 5.10 | 8.61 | 22.73 | 1065.74 | 24.62 | -13.24 | 37.86 | 17.79 | -1.75 | 17.79 | -8.32 | 551 | 78 | 29 | 31.08 | 194 | 90 | 194 | 105 |
| *B. sylvaticum* | Bsyl_674 | 58.60 | 6.50 | 5.02 | 5.12 | 25.24 | 577.71 | 15.94 | -4.35 | 20.29 | 2.49 | 7.33 | 12.24 | -1.61 | 2411 | 303 | 103 | 37.59 | 895 | 331 | 430 | 741 |
| *B. sylvaticum* | Bsyl_675 | 55.50 | 10.70 | 8.70 | 5.55 | 25.80 | 603.26 | 20.91 | -0.60 | 21.50 | 9.79 | 3.62 | 16.25 | 1.71 | 539 | 54 | 29 | 19.42 | 160 | 97 | 154 | 118 |
| *B. sylvaticum* | Bsyl_676 | 49.30 | 3.70 | 10.14 | 8.74 | 36.34 | 570.68 | 24.01 | -0.04 | 24.05 | 15.65 | 6.19 | 17.34 | 3.37 | 640 | 59 | 45 | 8.68 | 174 | 142 | 166 | 154 |
| *B. sylvaticum* | Bsyl_677 | 48.30 | 14.40 | 8.38 | 8.99 | 31.82 | 734.05 | 23.41 | -4.84 | 28.25 | 17.26 | 0.77 | 17.26 | -0.77 | 782 | 107 | 46 | 30.14 | 287 | 148 | 287 | 151 |
| *B. sylvaticum* | Bsyl_678 | 56.70 | -5.20 | 7.66 | 5.75 | 37.05 | 384.00 | 16.26 | 0.74 | 15.52 | 4.05 | 8.97 | 12.51 | 3.32 | 2115 | 256 | 93 | 34.31 | 748 | 303 | 371 | 674 |
| *B. sylvaticum* | Bsyl_679 | 55.20 | -2.90 | 6.85 | 7.31 | 39.28 | 426.91 | 17.56 | -1.04 | 18.61 | 2.72 | 8.31 | 12.37 | 2.02 | 1478 | 164 | 87 | 23.81 | 478 | 263 | 297 | 445 |
| *B. sylvaticum* | Bsyl_680 | 52.90 | 1.10 | 9.49 | 7.00 | 36.05 | 461.92 | 20.50 | 1.09 | 19.40 | 7.30 | 5.80 | 15.33 | 4.21 | 683 | 70 | 43 | 14.82 | 201 | 140 | 171 | 174 |
| *B. sylvaticum* | Bsyl_681 | 48.90 | 8.00 | 10.04 | 8.83 | 34.48 | 649.10 | 24.49 | -1.12 | 25.60 | 16.64 | 3.35 | 18.16 | 2.18 | 824 | 92 | 55 | 17.55 | 251 | 167 | 225 | 183 |
| *B. sylvaticum* | Bsyl_682 | 48.50 | 7.10 | 7.12 | 6.81 | 29.68 | 609.95 | 19.86 | -3.09 | 22.96 | 3.80 | 2.72 | 14.79 | -0.07 | 1200 | 124 | 83 | 12.05 | 340 | 270 | 289 | 321 |
| *B. sylvaticum* | Bsyl_683 | 47.90 | 7.10 | 6.09 | 6.76 | 30.40 | 585.38 | 18.53 | -3.71 | 22.24 | 0.40 | 13.26 | 13.45 | -0.64 | 1460 | 150 | 104 | 10.98 | 414 | 330 | 351 | 399 |
| *B. sylvaticum* | Bsyl_684 | 63.60 | 10.90 | 5.44 | 6.35 | 28.45 | 589.69 | 17.60 | -4.71 | 22.31 | 9.60 | 4.32 | 13.07 | -1.23 | 1050 | 121 | 56 | 23.05 | 327 | 184 | 266 | 278 |
| *B. sylvaticum* | Bsyl_685 | 61.90 | 6.80 | 3.67 | 4.92 | 25.96 | 533.22 | 14.40 | -4.53 | 18.93 | 0.65 | 6.06 | 10.69 | -2.28 | 1756 | 220 | 64 | 36.27 | 610 | 239 | 308 | 572 |
| *B. sylvaticum* | Bsyl_686 | 43.20 | -1.00 | 13.11 | 8.97 | 42.07 | 453.21 | 25.00 | 3.68 | 21.32 | 8.82 | 18.66 | 19.11 | 8.13 | 1145 | 129 | 60 | 22.26 | 356 | 207 | 212 | 325 |
| *B. sylvaticum* | Bsyl_687 | 48.70 | 0.50 | 9.67 | 7.64 | 36.03 | 507.96 | 21.90 | 0.68 | 21.22 | 4.60 | 16.04 | 16.04 | 3.71 | 757 | 78 | 50 | 14.93 | 225 | 160 | 160 | 219 |
| *B. sylvaticum* | Bsyl_688 | 42.00 | -1.50 | 14.40 | 10.78 | 39.46 | 603.67 | 29.66 | 2.35 | 27.32 | 16.19 | 22.03 | 22.03 | 7.37 | 391 | 48 | 22 | 24.14 | 128 | 72 | 72 | 91 |
| *B. sylvaticum* | Bsyl_689 | 57.90 | 16.50 | 6.90 | 7.81 | 28.65 | 703.16 | 22.05 | -5.19 | 27.25 | 15.07 | 5.30 | 16.06 | -1.17 | 581 | 71 | 32 | 25.09 | 188 | 106 | 178 | 133 |
| *B. sylvaticum* | Bsyl_690 | 49.20 | 0.60 | 10.28 | 7.06 | 34.91 | 482.86 | 21.88 | 1.66 | 20.22 | 7.98 | 16.24 | 16.32 | 4.60 | 745 | 79 | 49 | 16.96 | 230 | 156 | 161 | 213 |
| *B. sylvaticum* | Bsyl_692 | 45.40 | 6.20 | 3.98 | 7.42 | 30.76 | 613.12 | 17.61 | -6.51 | 24.12 | 1.23 | 11.73 | 11.73 | -2.87 | 1509 | 161 | 88 | 16.01 | 448 | 302 | 302 | 416 |
| *B. sylvaticum* | Bsyl_693 | 44.50 | 2.30 | 10.93 | 10.13 | 38.57 | 591.41 | 25.72 | -0.54 | 26.27 | 13.11 | 18.10 | 18.44 | 3.97 | 761 | 80 | 47 | 14.14 | 214 | 171 | 172 | 189 |
| *B. sylvaticum* | Bsyl_694 | 46.10 | 0.20 | 11.71 | 9.08 | 38.86 | 543.68 | 25.20 | 1.83 | 23.37 | 6.06 | 18.54 | 18.54 | 5.39 | 860 | 96 | 48 | 22.21 | 277 | 152 | 152 | 262 |
| *B. sylvaticum* | Bsyl_695 | 43.00 | 1.30 | 10.89 | 9.62 | 39.49 | 546.29 | 24.15 | -0.21 | 24.36 | 12.34 | 17.85 | 17.85 | 4.69 | 975 | 107 | 63 | 14.20 | 294 | 216 | 216 | 224 |
| *B. sylvaticum* | Bsyl_696 | 44.00 | 3.60 | 12.51 | 8.68 | 34.99 | 589.96 | 26.35 | 1.56 | 24.80 | 9.21 | 20.06 | 20.06 | 5.74 | 705 | 99 | 28 | 29.16 | 226 | 120 | 120 | 189 |
| *B. sylvaticum* | Bsyl_697 | 44.50 | 1.30 | 12.61 | 9.91 | 39.12 | 570.82 | 26.88 | 1.55 | 25.33 | 9.28 | 19.79 | 19.79 | 5.92 | 847 | 85 | 55 | 12.59 | 230 | 178 | 178 | 229 |
| *B. sylvaticum* | Bsyl_698 | 44.00 | 2.80 | 11.98 | 9.34 | 36.08 | 603.19 | 26.45 | 0.55 | 25.90 | 8.64 | 19.66 | 19.66 | 4.98 | 688 | 74 | 35 | 17.56 | 196 | 140 | 140 | 179 |
| *B. sylvaticum* | Bsyl_699 | 46.60 | 0.40 | 11.47 | 9.54 | 39.76 | 551.90 | 25.50 | 1.50 | 24.00 | 8.30 | 18.47 | 18.47 | 5.08 | 683 | 70 | 41 | 17.41 | 204 | 136 | 136 | 189 |
| *B. sylvaticum* | Bsyl_700 | 41.80 | 8.90 | 14.12 | 8.05 | 35.29 | 545.89 | 26.75 | 3.95 | 22.80 | 12.00 | 20.93 | 21.08 | 7.97 | 669 | 98 | 11 | 48.56 | 269 | 55 | 83 | 210 |
| *B. sylvaticum* | Bsyl_701 | 46.80 | -0.20 | 11.39 | 9.22 | 38.90 | 545.00 | 25.27 | 1.58 | 23.69 | 5.82 | 18.28 | 18.28 | 5.05 | 773 | 88 | 45 | 21.48 | 248 | 142 | 142 | 227 |
| *B. sylvaticum* | Bsyl_702 | 45.40 | 4.00 | 8.14 | 8.92 | 35.76 | 589.32 | 22.52 | -2.43 | 24.95 | 12.89 | 2.12 | 15.62 | 1.35 | 844 | 94 | 47 | 20.69 | 249 | 155 | 230 | 163 |
| *B. sylvaticum* | Bsyl_703 | 42.70 | -2.50 | 10.35 | 9.79 | 40.17 | 530.81 | 24.40 | 0.02 | 24.38 | 4.97 | 17.17 | 17.17 | 4.21 | 707 | 79 | 38 | 24.41 | 210 | 119 | 119 | 192 |
| *B. sylvaticum* | Bsyl_705 | 37.50 | -3.10 | 14.57 | 14.12 | 41.23 | 708.57 | 34.60 | 0.34 | 34.26 | 7.44 | 23.69 | 23.85 | 6.61 | 414 | 54 | 5 | 48.99 | 155 | 30 | 31 | 145 |
| *B. sylvaticum* | Bsyl_706 | 36.40 | -6.00 | 17.53 | 8.80 | 39.26 | 484.20 | 29.95 | 7.54 | 22.41 | 13.05 | 23.49 | 23.93 | 12.04 | 665 | 112 | 0 | 72.05 | 317 | 17 | 22 | 292 |
| *B. sylvaticum* | Bsyl_707 | 37.90 | -4.80 | 17.55 | 12.92 | 41.54 | 645.13 | 35.22 | 4.12 | 31.11 | 10.89 | 25.72 | 25.87 | 10.13 | 540 | 87 | 3 | 62.33 | 241 | 26 | 31 | 210 |
| *B. sylvaticum* | Bsyl_708 | 36.90 | -2.80 | 8.69 | 12.99 | 38.54 | 726.02 | 28.96 | -4.76 | 33.72 | 1.76 | 18.10 | 18.34 | 0.66 | 692 | 87 | 10 | 48.81 | 254 | 52 | 55 | 249 |
| *B. sylvaticum* | Bsyl_709 | 38.00 | -6.90 | 15.44 | 11.44 | 41.17 | 573.28 | 31.60 | 3.80 | 27.80 | 9.85 | 22.73 | 22.95 | 8.94 | 612 | 100 | 4 | 62.04 | 270 | 29 | 33 | 250 |
| *B. sylvaticum* | Bsyl_710 | 36.60 | -4.80 | 15.17 | 10.15 | 39.86 | 544.39 | 29.81 | 4.34 | 25.46 | 10.10 | 22.16 | 22.37 | 9.11 | 683 | 113 | 1 | 71.39 | 320 | 18 | 24 | 306 |
| *B. sylvaticum* | Bsyl_711 | 37.10 | -4.80 | 16.19 | 12.82 | 43.00 | 601.82 | 33.38 | 3.57 | 29.81 | 10.32 | 23.84 | 24.12 | 9.42 | 616 | 104 | 2 | 66.76 | 286 | 22 | 27 | 264 |
| *B. sylvaticum* | Bsyl_712 | 37.90 | -5.60 | 14.74 | 11.99 | 40.00 | 625.39 | 32.24 | 2.26 | 29.98 | 8.60 | 22.78 | 22.94 | 7.68 | 670 | 108 | 4 | 62.37 | 296 | 31 | 34 | 271 |
| *B. sylvaticum* | Bsyl_713 | 36.90 | -3.40 | 16.19 | 11.85 | 41.58 | 591.73 | 32.34 | 3.84 | 28.50 | 10.41 | 23.81 | 23.91 | 9.54 | 392 | 57 | 2 | 56.40 | 161 | 21 | 22 | 151 |
| *B. sylvaticum* | Bsyl_714 | 37.70 | -3.80 | 15.12 | 13.26 | 40.97 | 672.57 | 33.99 | 1.62 | 32.37 | 8.34 | 23.86 | 23.86 | 7.55 | 466 | 66 | 4 | 52.68 | 182 | 30 | 30 | 169 |
| *B. sylvaticum* | Bsyl_715 | 56.40 | 38.60 | 4.48 | 8.03 | 21.93 | 1052.02 | 23.88 | -12.76 | 36.64 | 17.35 | -3.57 | 17.35 | -8.13 | 647 | 88 | 29 | 33.66 | 235 | 97 | 235 | 120 |
| *B. sylvaticum* | Bsyl_716 | 56.60 | 39.80 | 3.81 | 8.47 | 23.30 | 1045.17 | 23.03 | -13.33 | 36.36 | 16.52 | -8.02 | 16.52 | -8.97 | 599 | 87 | 26 | 34.63 | 218 | 91 | 218 | 107 |
| *B. sylvaticum* | Bsyl_717 | 56.20 | 42.40 | 3.64 | 8.16 | 21.56 | 1086.99 | 22.99 | -14.87 | 37.86 | 16.59 | -3.30 | 16.59 | -9.96 | 565 | 75 | 26 | 32.11 | 199 | 86 | 199 | 103 |
| *B. sylvaticum* | Bsyl_718 | 55.80 | 42.00 | 4.01 | 8.23 | 21.92 | 1063.65 | 23.15 | -14.40 | 37.55 | 16.71 | -2.68 | 16.71 | -9.30 | 565 | 78 | 25 | 34.49 | 205 | 83 | 205 | 101 |
| *B. sylvaticum* | Bsyl_719 | 55.30 | 41.30 | 4.28 | 8.76 | 23.00 | 1063.22 | 23.90 | -14.21 | 38.11 | 17.06 | -2.35 | 17.06 | -9.03 | 597 | 82 | 28 | 31.75 | 210 | 92 | 210 | 114 |
| *B. sylvaticum* | Bsyl_720 | 56.00 | 39.40 | 4.58 | 8.25 | 22.40 | 1056.78 | 24.06 | -12.76 | 36.82 | 17.50 | -7.87 | 17.50 | -8.04 | 617 | 89 | 28 | 34.62 | 225 | 95 | 225 | 111 |
| *B. sylvaticum* | Bsyl_721 | 56.10 | 40.40 | 4.23 | 8.35 | 22.91 | 1045.34 | 23.16 | -13.27 | 36.43 | 16.83 | -7.36 | 16.83 | -8.83 | 600 | 82 | 26 | 32.10 | 211 | 92 | 211 | 110 |
| *B. sylvaticum* | Bsyl_722 | 51.50 | 105.30 | -2.34 | 10.29 | 24.07 | 1205.09 | 18.95 | -23.80 | 42.75 | 12.58 | -15.57 | 12.58 | -17.23 | 581 | 131 | 9 | 83.41 | 325 | 35 | 325 | 37 |
| *B. sylvaticum* | Bsyl_723 | 53.30 | 93.50 | 0.71 | 12.20 | 26.03 | 1288.23 | 24.26 | -22.60 | 46.86 | 16.08 | -12.96 | 16.08 | -15.82 | 540 | 97 | 13 | 61.52 | 254 | 47 | 254 | 60 |
| *B. sylvaticum* | Bsyl_724 | 52.20 | 87.40 | 2.05 | 12.84 | 27.94 | 1224.60 | 25.30 | -20.66 | 45.96 | 16.74 | -10.92 | 16.74 | -13.41 | 825 | 123 | 19 | 57.73 | 351 | 61 | 351 | 73 |
| *B. sylvaticum* | Bsyl_725 | 53.30 | 87.50 | 1.33 | 11.31 | 25.25 | 1226.37 | 24.25 | -20.54 | 44.80 | 16.18 | -11.74 | 16.18 | -14.07 | 716 | 97 | 23 | 43.25 | 274 | 78 | 274 | 100 |
| *B. sylvaticum* | Bsyl_726 | 53.40 | 85.70 | 2.06 | 10.16 | 22.40 | 1304.70 | 25.43 | -19.90 | 45.34 | 17.63 | -12.63 | 17.63 | -14.30 | 563 | 71 | 20 | 37.33 | 211 | 74 | 211 | 94 |
| *B. sylvaticum* | Bsyl_727 | 54.00 | 87.10 | 1.74 | 10.36 | 23.28 | 1277.03 | 25.28 | -19.20 | 44.48 | 17.28 | -12.13 | 17.28 | -14.07 | 600 | 81 | 20 | 41.52 | 232 | 73 | 232 | 93 |
| *B. sylvaticum* | Bsyl_728 | 51.90 | 88.30 | -3.08 | 11.22 | 26.69 | 1165.81 | 18.79 | -23.24 | 42.04 | 11.27 | -15.40 | 11.27 | -17.41 | 666 | 100 | 18 | 52.86 | 280 | 60 | 280 | 74 |
| *B. sylvaticum* | Bsyl_729 | 44.90 | 44.10 | 10.51 | 10.33 | 27.20 | 1042.26 | 31.52 | -6.46 | 37.98 | 21.11 | -1.08 | 23.35 | -2.31 | 409 | 60 | 20 | 35.32 | 153 | 65 | 149 | 69 |
| *B. sylvaticum* | Bsyl_730 | 44.80 | 37.50 | 11.45 | 7.90 | 27.62 | 760.56 | 26.59 | -2.02 | 28.61 | 4.06 | 20.52 | 21.05 | 2.43 | 658 | 93 | 41 | 26.68 | 229 | 133 | 141 | 214 |
| *B. sylvaticum* | Bsyl_731 | 44.60 | 38.40 | 11.10 | 7.54 | 27.58 | 727.91 | 25.36 | -1.98 | 27.33 | 3.95 | 19.79 | 20.26 | 2.47 | 946 | 133 | 60 | 27.50 | 336 | 188 | 200 | 312 |
| *B. sylvaticum* | Bsyl_732 | 44.90 | 39.00 | 11.84 | 9.90 | 30.60 | 846.48 | 28.96 | -3.38 | 32.34 | 3.03 | 21.36 | 22.23 | 1.41 | 801 | 92 | 51 | 18.67 | 254 | 168 | 193 | 231 |
| *B. sylvaticum* | Bsyl_733 | 44.30 | 39.00 | 11.59 | 7.98 | 29.37 | 718.29 | 26.02 | -1.16 | 27.18 | 4.58 | 18.33 | 20.53 | 2.96 | 1342 | 191 | 79 | 30.84 | 498 | 255 | 265 | 455 |
| *B. sylvaticum* | Bsyl_734 | 44.60 | 39.50 | 11.65 | 9.16 | 30.23 | 797.42 | 27.58 | -2.71 | 30.28 | 3.39 | 20.74 | 21.37 | 1.74 | 1132 | 143 | 76 | 23.93 | 391 | 235 | 244 | 351 |
| *B. sylvaticum* | Bsyl_735 | 44.60 | 40.20 | 10.49 | 9.38 | 30.20 | 816.86 | 26.77 | -4.30 | 31.07 | 2.03 | 5.32 | 20.27 | 0.21 | 1025 | 108 | 66 | 15.60 | 304 | 208 | 259 | 265 |
| *B. sylvaticum* | Bsyl_736 | 43.70 | 40.00 | 9.91 | 9.34 | 33.52 | 696.63 | 24.72 | -3.14 | 27.86 | 6.44 | 18.39 | 18.39 | 1.32 | 1619 | 192 | 105 | 22.67 | 537 | 333 | 333 | 460 |
| *B. sylvaticum* | Bsyl_737 | 43.90 | 40.60 | 3.18 | 9.57 | 32.99 | 715.82 | 18.67 | -10.34 | 29.01 | 6.59 | -5.16 | 11.80 | -5.56 | 1264 | 133 | 67 | 20.27 | 385 | 222 | 348 | 257 |
| *B. sylvaticum* | Bsyl_738 | 44.20 | 43.10 | 9.16 | 10.07 | 28.48 | 939.74 | 28.00 | -7.35 | 35.35 | 18.53 | -2.54 | 20.57 | -2.54 | 520 | 81 | 19 | 45.62 | 213 | 66 | 196 | 66 |
| *B. sylvaticum* | Bsyl_739 | 43.70 | 42.80 | 4.56 | 8.42 | 31.07 | 700.73 | 18.22 | -8.89 | 27.11 | 11.22 | -4.05 | 13.00 | -4.05 | 734 | 118 | 24 | 55.22 | 319 | 77 | 312 | 77 |
| *B. sylvaticum* | Bsyl_740 | 43.50 | 43.40 | 6.80 | 7.84 | 27.72 | 764.15 | 21.34 | -6.95 | 28.29 | 14.23 | -2.57 | 16.06 | -2.57 | 795 | 117 | 31 | 45.09 | 319 | 103 | 306 | 103 |
| *B. sylvaticum* | Bsyl_741 | 43.50 | 45.50 | 10.79 | 10.08 | 27.54 | 991.37 | 30.46 | -6.13 | 36.59 | 20.51 | -1.64 | 22.80 | -1.64 | 452 | 66 | 22 | 38.43 | 178 | 70 | 167 | 70 |
| *B. sylvaticum* | Bsyl_742 | 42.10 | 47.90 | 10.82 | 7.36 | 24.92 | 833.99 | 26.20 | -3.34 | 29.54 | 12.03 | 1.19 | 21.13 | 0.82 | 446 | 57 | 21 | 27.72 | 143 | 76 | 116 | 76 |
| *B. sylvaticum* | Bsyl_743 | 41.70 | 48.40 | 11.48 | 8.08 | 25.75 | 857.59 | 28.07 | -3.30 | 31.37 | 12.58 | 1.64 | 22.18 | 1.30 | 403 | 49 | 24 | 24.05 | 134 | 81 | 81 | 85 |
| *B. sylvaticum* | Bsyl_744 | 44.10 | 47.10 | 12.21 | 8.26 | 24.06 | 993.41 | 30.78 | -3.57 | 34.35 | 17.63 | 0.98 | 24.57 | 0.24 | 264 | 31 | 15 | 20.22 | 81 | 51 | 69 | 51 |
| *B. sylvaticum* | Bsyl_745 | 53.60 | 92.20 | 1.02 | 11.99 | 25.89 | 1256.34 | 24.18 | -22.15 | 46.33 | 16.19 | -12.65 | 16.19 | -14.89 | 465 | 88 | 9 | 71.22 | 238 | 32 | 238 | 41 |
| *B. sylvaticum* | Bsyl_746 | 52.90 | 92.80 | -1.15 | 10.43 | 24.42 | 1212.61 | 20.80 | -21.91 | 42.71 | 13.51 | -13.67 | 13.51 | -16.38 | 790 | 131 | 21 | 53.70 | 344 | 76 | 344 | 96 |
| *B. sylvaticum* | Bsyl_747 | 51.80 | 85.90 | 1.72 | 12.44 | 29.67 | 1118.62 | 23.17 | -18.74 | 41.91 | 15.15 | -10.66 | 15.15 | -12.01 | 673 | 103 | 20 | 53.64 | 286 | 60 | 286 | 72 |
| *B. sylvaticum* | Bsyl_748 | 51.90 | 86.70 | 2.05 | 11.91 | 28.25 | 1131.48 | 23.70 | -18.48 | 42.18 | 15.69 | -10.21 | 15.69 | -12.03 | 749 | 116 | 18 | 59.00 | 328 | 56 | 328 | 66 |
| *B. sylvaticum* | Bsyl_749 | 50.50 | 86.90 | -2.12 | 9.97 | 27.07 | 1033.30 | 17.05 | -19.80 | 36.85 | 10.39 | -13.25 | 10.39 | -14.91 | 464 | 82 | 9 | 68.13 | 227 | 30 | 227 | 36 |
| *B. sylvaticum* | Bsyl_750 | 37.00 | 9.90 | 18.04 | 11.43 | 41.32 | 610.01 | 33.71 | 6.04 | 27.67 | 12.52 | 25.63 | 25.80 | 11.19 | 542 | 81 | 4 | 59.53 | 226 | 24 | 50 | 223 |
| *B. sylvaticum* | Bsyl_751 | 36.60 | 10.60 | 18.90 | 8.92 | 36.86 | 566.66 | 32.16 | 7.96 | 24.20 | 13.95 | 25.80 | 26.21 | 12.47 | 480 | 69 | 3 | 58.81 | 196 | 19 | 50 | 193 |
| *B. sylvaticum* | Bsyl_752 | 37.90 | 14.40 | 11.85 | 6.87 | 29.78 | 600.65 | 25.14 | 2.06 | 23.08 | 6.72 | 19.40 | 19.51 | 5.26 | 603 | 91 | 8 | 55.32 | 246 | 34 | 74 | 229 |
| *B. sylvaticum* | Bsyl_753 | 35.90 | 14.50 | 19.17 | 7.26 | 34.26 | 520.07 | 30.99 | 9.79 | 21.20 | 15.04 | 25.50 | 25.86 | 13.35 | 517 | 101 | 0 | 83.55 | 270 | 6 | 48 | 180 |
| *B. sylvaticum* | Bsyl_754 | 38.70 | 23.70 | 12.86 | 7.47 | 30.33 | 656.75 | 26.67 | 2.03 | 24.64 | 5.26 | 21.17 | 21.17 | 5.26 | 614 | 94 | 14 | 54.22 | 249 | 51 | 51 | 249 |
| *B. sylvaticum* | Bsyl_755 | 40.50 | 25.60 | 11.98 | 7.36 | 28.73 | 687.78 | 26.15 | 0.55 | 25.60 | 5.60 | 20.12 | 20.61 | 3.95 | 659 | 103 | 17 | 48.83 | 270 | 70 | 73 | 258 |
| *B. sylvaticum* | Bsyl_756 | 40.50 | 23.70 | 11.96 | 8.46 | 31.76 | 694.88 | 26.41 | -0.24 | 26.65 | 8.36 | 20.13 | 20.68 | 3.66 | 535 | 62 | 23 | 27.52 | 176 | 80 | 94 | 160 |
| *B. sylvaticum* | Bsyl_757 | 41.00 | 29.60 | 12.75 | 7.94 | 32.61 | 629.19 | 25.62 | 1.29 | 24.34 | 7.22 | 20.47 | 20.47 | 5.28 | 790 | 118 | 31 | 41.04 | 307 | 111 | 111 | 236 |
| *B. sylvaticum* | Bsyl_758 | 37.70 | 27.40 | 17.60 | 11.18 | 39.89 | 645.69 | 32.98 | 4.95 | 28.04 | 9.92 | 25.70 | 25.70 | 9.92 | 696 | 145 | 3 | 87.14 | 388 | 17 | 17 | 388 |
| *B. sylvaticum* | Bsyl_759 | 37.10 | 27.80 | 16.48 | 9.60 | 35.05 | 679.16 | 32.01 | 4.62 | 27.39 | 8.71 | 24.82 | 25.10 | 8.71 | 841 | 173 | 6 | 86.29 | 471 | 26 | 28 | 471 |
| *B. sylvaticum* | Bsyl_760 | 35.20 | 33.60 | 19.71 | 12.87 | 41.63 | 695.02 | 36.57 | 5.65 | 30.92 | 11.64 | 28.03 | 28.28 | 11.59 | 363 | 83 | 1 | 87.21 | 200 | 8 | 9 | 160 |
| *B. sylvaticum* | Bsyl_761 | 37.10 | 36.70 | 17.23 | 11.72 | 34.50 | 849.18 | 35.41 | 1.44 | 33.97 | 6.73 | 27.29 | 27.48 | 6.73 | 753 | 141 | 2 | 79.89 | 389 | 18 | 18 | 389 |
| *B. sylvaticum* | Bsyl_762 | 41.30 | 34.10 | 7.75 | 10.13 | 34.96 | 720.05 | 23.92 | -5.05 | 28.98 | 11.35 | 15.97 | 16.39 | -1.25 | 562 | 76 | 29 | 30.57 | 199 | 91 | 121 | 147 |
| *B. sylvaticum* | Bsyl_763 | 38.30 | 36.80 | 7.92 | 13.79 | 37.35 | 813.51 | 28.43 | -8.49 | 36.92 | -0.33 | 17.57 | 17.62 | -2.32 | 509 | 69 | 8 | 50.34 | 180 | 31 | 46 | 175 |
| *B. sylvaticum* | Bsyl_764 | 40.00 | 41.20 | 5.71 | 12.75 | 30.30 | 1037.45 | 27.84 | -14.22 | 42.07 | 10.38 | 17.74 | 17.74 | -7.73 | 422 | 66 | 16 | 41.23 | 165 | 60 | 60 | 82 |
| *B. sylvaticum* | Bsyl_765 | 39.40 | 42.80 | 5.93 | 12.04 | 29.36 | 1022.91 | 28.13 | -12.87 | 41.00 | 4.08 | 18.16 | 18.16 | -6.85 | 526 | 75 | 16 | 43.37 | 198 | 53 | 53 | 124 |
| *B. sylvaticum* | Bsyl_766 | 39.50 | 49.00 | 14.84 | 8.55 | 27.56 | 828.35 | 31.72 | 0.68 | 31.04 | 10.82 | 25.07 | 25.07 | 5.02 | 364 | 57 | 5 | 46.38 | 133 | 33 | 33 | 96 |
| *B. sylvaticum* | Bsyl_767 | 37.20 | 49.70 | 15.53 | 8.63 | 31.70 | 706.92 | 29.43 | 2.21 | 27.22 | 12.78 | 21.99 | 24.06 | 7.04 | 1198 | 196 | 32 | 56.64 | 530 | 111 | 125 | 385 |
| *B. sylvaticum* | Bsyl_768 | 37.60 | 45.40 | 11.34 | 8.53 | 25.94 | 893.40 | 28.34 | -4.52 | 32.86 | 10.01 | 21.81 | 22.00 | 0.10 | 291 | 52 | 2 | 65.56 | 137 | 11 | 20 | 81 |
| *B. sylvaticum* | Bsyl_769 | 36.00 | 52.60 | 3.80 | 10.76 | 28.80 | 968.53 | 23.40 | -13.95 | 37.35 | 2.78 | 14.95 | 15.56 | -8.27 | 222 | 44 | 2 | 72.41 | 119 | 15 | 23 | 57 |
| *B. sylvaticum* | Bsyl_770 | 36.40 | 53.90 | 9.67 | 10.98 | 29.91 | 961.98 | 28.50 | -8.21 | 36.71 | 9.15 | 20.54 | 21.15 | -2.57 | 205 | 37 | 4 | 60.37 | 96 | 15 | 20 | 61 |
| *B. sylvaticum* | Bsyl_771 | 34.90 | 50.50 | 17.17 | 13.77 | 35.00 | 944.74 | 38.15 | -1.18 | 39.34 | 6.83 | 27.97 | 28.63 | 5.22 | 255 | 41 | 1 | 73.43 | 118 | 4 | 5 | 116 |
| *B. sylvaticum* | Bsyl_772 | 33.70 | 48.80 | 14.08 | 14.96 | 38.51 | 874.65 | 35.46 | -3.38 | 38.84 | 4.34 | 24.59 | 25.00 | 3.41 | 345 | 63 | 0 | 83.29 | 169 | 0 | 6 | 160 |
| *B. sylvaticum* | Bsyl_773 | 33.70 | 47.20 | 17.43 | 15.06 | 38.10 | 909.91 | 39.34 | -0.19 | 39.53 | 7.43 | 28.20 | 28.70 | 6.37 | 443 | 85 | 0 | 86.52 | 222 | 2 | 3 | 211 |
| *B. sylvaticum* | Bsyl_774 | 44.00 | 45.20 | 10.86 | 10.60 | 28.17 | 1030.34 | 31.54 | -6.10 | 37.64 | 21.32 | -0.56 | 23.52 | -1.96 | 402 | 59 | 20 | 36.78 | 155 | 65 | 148 | 66 |
| *B. sylvaticum* | Bsyl_775 | 47.00 | 32.20 | 9.81 | 10.16 | 29.40 | 942.06 | 28.66 | -5.89 | 34.56 | 19.49 | -0.59 | 21.38 | -1.67 | 436 | 52 | 26 | 22.91 | 144 | 83 | 141 | 92 |
| *B. sylvaticum* | Bsyl_776 | 47.70 | 34.70 | 9.49 | 9.41 | 27.41 | 958.69 | 27.95 | -6.39 | 34.34 | 19.44 | 3.29 | 21.12 | -2.30 | 468 | 54 | 26 | 21.00 | 149 | 97 | 142 | 121 |
| *B. sylvaticum* | Bsyl_777 | 49.20 | 34.60 | 8.08 | 8.78 | 26.04 | 978.34 | 25.89 | -7.82 | 33.71 | 18.39 | 2.02 | 19.76 | -4.04 | 528 | 59 | 31 | 18.23 | 164 | 105 | 161 | 125 |
| *B. sylvaticum* | Bsyl_778 | 54.90 | 56.10 | 3.39 | 10.11 | 23.55 | 1221.63 | 25.28 | -17.64 | 42.92 | 17.95 | -3.94 | 17.95 | -11.80 | 573 | 65 | 25 | 27.98 | 186 | 90 | 186 | 124 |
| *B. sylvaticum* | Bsyl_779 | 55.80 | 55.30 | 3.02 | 9.20 | 21.74 | 1229.76 | 24.90 | -17.45 | 42.34 | 17.73 | -4.57 | 17.73 | -12.23 | 518 | 64 | 20 | 32.81 | 181 | 74 | 181 | 100 |
| *B. sylvaticum* | Bsyl_780 | 55.20 | 53.30 | 3.15 | 9.18 | 22.54 | 1178.49 | 24.30 | -16.44 | 40.75 | 17.25 | -4.18 | 17.25 | -11.36 | 492 | 65 | 21 | 34.67 | 179 | 75 | 179 | 86 |
| *B. sylvaticum* | Bsyl_781 | 54.70 | 51.70 | 3.79 | 9.41 | 22.15 | 1275.76 | 25.55 | -16.93 | 42.48 | 18.57 | -5.18 | 18.57 | -11.90 | 531 | 64 | 27 | 28.63 | 182 | 87 | 182 | 104 |
| *B. sylvaticum* | Bsyl_782 | 56.30 | 50.10 | 2.99 | 8.42 | 20.77 | 1200.77 | 24.28 | -16.25 | 40.54 | 17.42 | -4.37 | 17.42 | -11.97 | 559 | 71 | 28 | 32.64 | 201 | 89 | 201 | 99 |
| *B. sylvaticum* | Bsyl_783 | 60.80 | 50.70 | 0.86 | 8.72 | 21.53 | 1107.39 | 22.16 | -18.35 | 40.51 | 14.64 | -5.40 | 14.64 | -12.94 | 581 | 78 | 23 | 36.29 | 214 | 80 | 214 | 102 |
| *B. sylvaticum* | Bsyl_784 | 59.80 | 47.40 | 1.79 | 8.40 | 21.12 | 1091.13 | 22.19 | -17.57 | 39.76 | 15.23 | -4.28 | 15.23 | -11.88 | 594 | 77 | 24 | 35.25 | 216 | 82 | 216 | 104 |
| *B. sylvaticum* | Bsyl_785 | 58.20 | 45.80 | 2.86 | 8.93 | 22.65 | 1091.70 | 23.34 | -16.06 | 39.40 | 16.23 | -3.56 | 16.23 | -10.72 | 617 | 80 | 26 | 32.93 | 219 | 90 | 219 | 113 |
| *B. sylvaticum* | Bsyl_786 | 57.10 | 44.40 | 3.77 | 7.70 | 21.11 | 1051.98 | 22.66 | -13.82 | 36.48 | 16.51 | -2.37 | 16.51 | -9.42 | 611 | 80 | 27 | 31.69 | 211 | 91 | 211 | 113 |
| *B. sylvaticum* | Bsyl_787 | 57.40 | 22.20 | 6.50 | 6.36 | 23.61 | 747.50 | 20.99 | -5.92 | 26.92 | 15.33 | 0.36 | 15.93 | -2.26 | 646 | 74 | 27 | 33.15 | 218 | 95 | 205 | 123 |
| *B. sylvaticum* | Bsyl_788 | 56.70 | 23.40 | 6.81 | 7.27 | 24.48 | 819.92 | 23.03 | -6.67 | 29.70 | 15.96 | -2.22 | 17.01 | -3.32 | 585 | 76 | 25 | 33.11 | 209 | 86 | 206 | 99 |
| *B. sylvaticum* | Bsyl_789 | 56.90 | 24.40 | 6.10 | 7.12 | 24.90 | 811.17 | 21.50 | -7.10 | 28.60 | 15.01 | -3.06 | 16.14 | -3.78 | 647 | 78 | 28 | 30.87 | 226 | 98 | 213 | 114 |
| *B. sylvaticum* | Bsyl_790 | 58.40 | 24.60 | 5.81 | 6.62 | 22.58 | 823.74 | 21.37 | -7.96 | 29.33 | 6.34 | -0.70 | 16.21 | -3.91 | 694 | 81 | 32 | 31.10 | 228 | 106 | 208 | 144 |
| *B. sylvaticum* | Bsyl_791 | 58.90 | 22.60 | 6.26 | 5.48 | 21.46 | 730.13 | 20.23 | -5.29 | 25.52 | 7.35 | -0.10 | 15.60 | -2.07 | 638 | 78 | 32 | 33.20 | 227 | 100 | 170 | 111 |
| *B. sylvaticum* | Bsyl_792 | 63.20 | 9.70 | 4.21 | 6.62 | 29.63 | 605.24 | 16.88 | -5.47 | 22.34 | 0.54 | 7.04 | 11.90 | -2.81 | 1115 | 130 | 55 | 24.21 | 345 | 191 | 252 | 329 |
| *B. sylvaticum* | Bsyl_793 | 59.00 | 13.20 | 6.20 | 7.70 | 28.03 | 723.48 | 21.39 | -6.08 | 27.48 | 10.93 | 0.77 | 15.40 | -2.39 | 635 | 70 | 35 | 25.26 | 206 | 110 | 188 | 132 |
| *B. sylvaticum* | Bsyl_794 | 56.00 | 9.00 | 8.01 | 7.08 | 31.57 | 590.02 | 20.59 | -1.84 | 22.43 | 8.78 | 3.34 | 15.34 | 1.05 | 834 | 97 | 45 | 25.49 | 280 | 148 | 202 | 198 |
| *B. sylvaticum* | Bsyl_795 | 56.80 | 8.70 | 8.02 | 6.22 | 28.74 | 581.50 | 20.10 | -1.55 | 21.66 | 9.01 | 3.20 | 15.24 | 1.26 | 785 | 94 | 41 | 28.35 | 273 | 136 | 187 | 183 |
| *B. sylvaticum* | Bsyl_796 | 54.20 | -4.50 | 8.91 | 6.22 | 38.60 | 375.35 | 17.97 | 1.87 | 16.10 | 7.26 | 12.30 | 13.67 | 4.67 | 978 | 110 | 58 | 23.77 | 321 | 177 | 199 | 273 |
| *B. sylvaticum* | Bsyl_797 | 51.30 | 80.90 | 2.89 | 12.08 | 24.96 | 1391.45 | 27.88 | -20.50 | 48.38 | 17.69 | -13.10 | 19.37 | -14.34 | 347 | 59 | 15 | 39.23 | 131 | 56 | 131 | 63 |
| *B. sylvaticum* | Bsyl_798 | 51.10 | 85.70 | -3.00 | 10.54 | 28.93 | 985.75 | 16.43 | -19.99 | 36.42 | 9.30 | -13.77 | 9.30 | -14.71 | 666 | 101 | 17 | 56.48 | 287 | 55 | 287 | 63 |
| *B. sylvaticum* | Bsyl_799 | 53.40 | 84.80 | 1.66 | 10.66 | 23.24 | 1323.70 | 25.07 | -20.80 | 45.87 | 17.42 | -13.09 | 17.42 | -14.91 | 516 | 66 | 19 | 33.75 | 188 | 72 | 188 | 94 |
| *B. sylvaticum* | Bsyl_800 | 52.60 | 85.70 | 2.50 | 11.91 | 25.44 | 1291.48 | 26.08 | -20.72 | 46.80 | 17.90 | -12.07 | 17.90 | -13.63 | 616 | 82 | 21 | 39.99 | 234 | 75 | 234 | 94 |
| *B. sylvaticum* | Bsyl_801 | 53.30 | 86.70 | 2.04 | 10.35 | 23.50 | 1236.64 | 24.79 | -19.26 | 44.05 | 17.07 | -11.34 | 17.07 | -13.26 | 654 | 87 | 22 | 41.30 | 249 | 77 | 249 | 97 |
| *B. sylvaticum* | Bsyl_802 | 54.70 | 88.00 | 0.19 | 9.54 | 23.06 | 1166.23 | 22.45 | -18.92 | 41.38 | 14.92 | -12.26 | 14.92 | -13.90 | 621 | 86 | 19 | 46.54 | 252 | 65 | 252 | 80 |
| *B. sylvaticum* | Bsyl_803 | 52.70 | 103.70 | 0.37 | 10.91 | 22.22 | 1400.76 | 24.41 | -24.68 | 49.10 | 16.96 | -14.33 | 16.96 | -17.73 | 432 | 102 | 7 | 88.66 | 254 | 26 | 254 | 32 |
| *B. sylvaticum* | Bsyl_804 | 51.20 | 109.80 | -3.69 | 15.37 | 29.56 | 1344.79 | 21.24 | -30.74 | 51.99 | 12.60 | -18.05 | 12.60 | -20.66 | 385 | 103 | 3 | 108.14 | 259 | 11 | 259 | 15 |
| *B. sylvaticum* | Bsyl_805 | 52.30 | 116.30 | -3.24 | 14.17 | 23.52 | 1735.73 | 25.40 | -34.86 | 60.26 | 16.71 | -21.94 | 16.71 | -26.01 | 357 | 101 | 3 | 113.96 | 250 | 10 | 250 | 12 |
| *B. sylvaticum* | Bsyl_806 | 40.70 | 45.10 | 8.09 | 11.05 | 33.18 | 765.02 | 24.73 | -8.57 | 33.30 | 11.13 | -1.13 | 17.37 | -1.13 | 508 | 84 | 18 | 51.01 | 221 | 63 | 120 | 63 |
| *B. sylvaticum* | Bsyl_807 | 39.50 | 46.30 | 8.25 | 10.75 | 30.85 | 865.65 | 26.46 | -8.37 | 34.84 | 11.50 | -2.25 | 18.85 | -2.25 | 592 | 88 | 27 | 38.62 | 233 | 91 | 121 | 91 |
| *B. sylvaticum* | Bsyl_808 | 41.40 | 42.20 | 6.05 | 10.50 | 32.15 | 799.14 | 23.14 | -9.51 | 32.65 | 2.49 | 0.29 | 15.41 | -4.02 | 1000 | 102 | 62 | 16.24 | 295 | 208 | 222 | 250 |
| *B. sylvaticum* | Bsyl_809 | 40.40 | 43.20 | 2.86 | 11.80 | 31.10 | 931.82 | 23.04 | -14.89 | 37.92 | 6.49 | -8.77 | 14.01 | -8.77 | 560 | 92 | 26 | 41.91 | 227 | 89 | 122 | 89 |
| *B. sylvaticum* | Bsyl_810 | 43.20 | 40.40 | 13.02 | 8.28 | 32.19 | 654.51 | 26.29 | 0.57 | 25.72 | 9.82 | 19.12 | 21.17 | 5.11 | 1609 | 192 | 102 | 23.19 | 536 | 320 | 330 | 462 |
| *B. sylvaticum* | Bsyl_811 | 43.00 | 42.80 | 3.06 | 7.90 | 28.95 | 708.48 | 17.37 | -9.93 | 27.30 | 11.62 | -5.25 | 11.62 | -5.55 | 1167 | 123 | 68 | 16.94 | 346 | 222 | 346 | 246 |
| *B. sylvaticum* | Bsyl_812 | 42.40 | 41.90 | 14.12 | 9.35 | 36.47 | 635.49 | 26.84 | 1.19 | 25.65 | 21.87 | 13.09 | 21.87 | 6.28 | 1819 | 176 | 105 | 15.05 | 514 | 361 | 514 | 456 |
| *B. sylvaticum* | Bsyl_813 | 42.30 | 42.60 | 14.41 | 9.99 | 36.51 | 675.95 | 28.48 | 1.11 | 27.37 | 7.70 | 13.58 | 22.44 | 5.92 | 1543 | 180 | 91 | 18.73 | 477 | 309 | 361 | 461 |
| *B. sylvaticum* | Bsyl_814 | 41.90 | 41.90 | 13.96 | 8.82 | 36.08 | 629.49 | 26.46 | 2.02 | 24.43 | 11.27 | 12.33 | 21.64 | 6.37 | 2067 | 244 | 83 | 29.36 | 687 | 322 | 482 | 599 |
| *B. sylvaticum* | Bsyl_815 | 41.50 | 42.90 | 3.31 | 10.34 | 32.56 | 789.69 | 19.90 | -11.85 | 31.75 | 10.36 | -5.96 | 12.58 | -6.52 | 849 | 110 | 53 | 26.25 | 295 | 163 | 218 | 172 |
| *B. sylvaticum* | Bsyl_816 | 41.90 | 43.50 | 8.36 | 8.44 | 29.21 | 752.94 | 24.09 | -4.80 | 28.89 | 11.29 | 17.47 | 17.47 | -0.77 | 896 | 101 | 57 | 17.62 | 271 | 190 | 190 | 232 |
| *B. sylvaticum* | Bsyl_817 | 42.40 | 43.60 | 8.55 | 8.88 | 30.49 | 747.60 | 24.24 | -4.88 | 29.11 | 11.62 | -0.34 | 17.45 | -0.51 | 1109 | 112 | 75 | 12.58 | 315 | 251 | 252 | 282 |
| *B. sylvaticum* | Bsyl_818 | 42.30 | 44.40 | 5.62 | 9.18 | 32.00 | 748.68 | 20.76 | -7.94 | 28.70 | 8.70 | -3.56 | 14.49 | -3.56 | 1047 | 140 | 58 | 32.42 | 381 | 179 | 277 | 179 |
| *B. sylvaticum* | Bsyl_819 | 41.50 | 44.20 | 7.78 | 11.10 | 35.77 | 738.19 | 24.00 | -7.03 | 31.02 | 11.02 | -1.22 | 16.68 | -1.22 | 676 | 113 | 21 | 52.48 | 291 | 77 | 239 | 77 |
| *B. sylvaticum* | Bsyl_820 | 41.90 | 45.50 | 11.48 | 10.28 | 31.89 | 818.55 | 28.37 | -3.86 | 32.23 | 15.50 | 1.40 | 21.56 | 1.40 | 728 | 110 | 27 | 44.31 | 292 | 92 | 247 | 92 |
| *B. sylvaticum* | Bsyl_821 | 41.60 | 46.80 | 9.46 | 9.18 | 30.72 | 764.18 | 24.87 | -5.02 | 29.89 | 12.70 | 0.36 | 18.85 | 0.36 | 711 | 105 | 23 | 44.38 | 278 | 80 | 239 | 80 |
| *B. sylvaticum* | Bsyl_822 | 40.60 | 46.30 | 12.27 | 9.90 | 30.08 | 837.43 | 30.11 | -2.79 | 32.90 | 15.95 | 2.15 | 22.60 | 2.15 | 430 | 75 | 15 | 51.80 | 187 | 53 | 143 | 53 |
| *B. sylvaticum* | Bsyl_823 | 40.30 | 46.80 | 12.96 | 9.39 | 29.24 | 821.66 | 30.62 | -1.48 | 32.09 | 16.52 | 3.18 | 23.29 | 3.18 | 430 | 68 | 15 | 47.71 | 180 | 52 | 124 | 52 |
| *B. sylvaticum* | Bsyl_824 | 41.00 | 47.40 | 13.09 | 10.15 | 29.74 | 879.88 | 31.24 | -2.91 | 34.15 | 17.01 | 2.46 | 24.00 | 2.46 | 531 | 66 | 29 | 30.38 | 186 | 94 | 127 | 94 |
| *B. sylvaticum* | Bsyl_825 | 40.80 | 48.00 | 11.61 | 9.89 | 29.70 | 869.07 | 29.18 | -4.10 | 33.29 | 15.82 | 1.02 | 22.35 | 1.02 | 607 | 76 | 35 | 29.14 | 202 | 112 | 135 | 112 |
| *B. sylvaticum* | Bsyl_826 | 40.30 | 48.40 | 14.67 | 9.73 | 28.74 | 892.73 | 33.08 | -0.78 | 33.86 | 13.12 | 25.83 | 25.83 | 4.04 | 336 | 43 | 10 | 32.35 | 106 | 49 | 49 | 82 |
| *B. sylvaticum* | Bsyl_827 | 41.30 | 49.00 | 12.58 | 7.94 | 25.97 | 847.13 | 29.30 | -1.25 | 30.56 | 8.57 | 23.20 | 23.20 | 2.77 | 293 | 37 | 11 | 29.22 | 98 | 42 | 42 | 77 |
| *B. sylvaticum* | Bsyl_828 | 40.80 | 49.20 | 10.45 | 7.90 | 25.43 | 827.69 | 27.69 | -3.36 | 31.05 | 8.68 | 20.96 | 20.96 | 0.81 | 413 | 51 | 18 | 29.67 | 138 | 70 | 70 | 88 |
| *B. sylvaticum* | Bsyl_829 | 39.10 | 46.60 | 13.05 | 10.68 | 31.87 | 840.51 | 30.65 | -2.87 | 33.52 | 12.14 | 22.90 | 23.29 | 2.63 | 436 | 71 | 16 | 46.07 | 172 | 60 | 79 | 74 |
| *B. sylvaticum* | Bsyl_830 | 39.00 | 48.60 | 14.05 | 8.93 | 28.45 | 799.22 | 31.63 | 0.24 | 31.39 | 15.36 | 23.96 | 23.96 | 4.52 | 686 | 122 | 11 | 51.04 | 287 | 69 | 69 | 179 |
| *B. sylvaticum* | Bsyl_831 | 38.50 | 48.60 | 9.41 | 9.77 | 32.46 | 690.58 | 26.03 | -4.05 | 30.08 | 11.02 | 17.61 | 17.61 | 0.81 | 521 | 77 | 15 | 38.76 | 176 | 71 | 71 | 106 |
| *B. sylvaticum* | Bsyl_832 | 40.00 | 23.40 | 15.70 | 8.06 | 30.14 | 709.52 | 30.28 | 3.54 | 26.74 | 12.07 | 24.05 | 24.72 | 7.33 | 437 | 59 | 16 | 38.21 | 163 | 57 | 58 | 140 |
| *B. sylvaticum* | Bsyl_833 | 40.20 | 24.30 | 12.78 | 6.41 | 27.13 | 656.51 | 25.58 | 1.94 | 23.64 | 6.68 | 20.59 | 21.09 | 5.14 | 561 | 77 | 17 | 42.76 | 217 | 65 | 73 | 201 |
| *B. sylvaticum* | Bsyl_834 | 35.60 | 27.10 | 17.19 | 5.71 | 28.83 | 540.89 | 27.89 | 8.10 | 19.79 | 11.09 | 23.92 | 23.92 | 11.01 | 722 | 150 | 2 | 86.84 | 397 | 10 | 10 | 345 |
| *B. sylvaticum* | Bsyl_835 | 35.20 | 24.80 | 10.00 | 7.83 | 32.82 | 615.54 | 23.29 | -0.56 | 23.86 | 2.93 | 17.68 | 17.68 | 2.93 | 1046 | 208 | 6 | 79.46 | 542 | 33 | 33 | 500 |
| *B. sylvaticum* | Bsyl_836 | 38.50 | 22.60 | 11.52 | 10.55 | 37.06 | 681.31 | 27.71 | -0.76 | 28.47 | 5.17 | 20.15 | 20.15 | 3.63 | 763 | 109 | 22 | 48.24 | 296 | 75 | 75 | 288 |
| *B. sylvaticum* | Bsyl_837 | 40.10 | 21.00 | 7.18 | 8.71 | 32.76 | 654.82 | 22.19 | -4.40 | 26.58 | 4.46 | 15.34 | 15.34 | -0.48 | 919 | 111 | 45 | 30.04 | 312 | 137 | 137 | 254 |
| *B. sylvaticum* | Bsyl_838 | 40.80 | 21.40 | 10.85 | 10.90 | 34.84 | 751.64 | 27.75 | -3.54 | 31.28 | 6.67 | 20.06 | 20.06 | 1.59 | 734 | 93 | 38 | 26.47 | 251 | 130 | 130 | 193 |
| *B. sylvaticum* | Bsyl_839 | 41.20 | 24.90 | 12.50 | 9.08 | 31.56 | 736.78 | 28.14 | -0.62 | 28.76 | 5.21 | 21.15 | 21.73 | 3.72 | 580 | 76 | 22 | 31.80 | 204 | 82 | 104 | 184 |
| *B. sylvaticum* | Bsyl_840 | 39.20 | 23.50 | 17.79 | 5.13 | 22.72 | 664.33 | 29.86 | 7.27 | 22.59 | 11.65 | 26.37 | 26.37 | 10.16 | 450 | 70 | 10 | 54.73 | 186 | 37 | 37 | 178 |
| *B. sylvaticum* | Bsyl_841 | 41.40 | 19.90 | 11.78 | 9.81 | 35.98 | 638.08 | 26.83 | -0.44 | 27.28 | 5.70 | 19.82 | 19.82 | 4.36 | 1150 | 153 | 44 | 34.81 | 410 | 152 | 152 | 365 |
| *B. sylvaticum* | Bsyl_842 | 43.30 | 17.80 | 15.34 | 10.27 | 33.43 | 749.77 | 32.43 | 1.72 | 30.71 | 10.96 | 24.86 | 24.86 | 6.32 | 1110 | 135 | 57 | 24.85 | 371 | 191 | 191 | 310 |
| *B. sylvaticum* | Bsyl_843 | 45.20 | 14.60 | 14.13 | 7.92 | 31.17 | 656.13 | 28.26 | 2.86 | 25.40 | 14.53 | 22.54 | 22.54 | 6.45 | 1232 | 163 | 51 | 34.20 | 462 | 218 | 218 | 302 |
| *B. sylvaticum* | Bsyl_844 | 44.90 | 22.40 | 9.40 | 9.40 | 31.04 | 785.05 | 25.64 | -4.65 | 30.29 | 17.21 | 0.97 | 18.92 | -0.42 | 710 | 88 | 42 | 24.18 | 238 | 129 | 218 | 139 |
| *B. sylvaticum* | Bsyl_845 | 45.50 | 25.30 | 5.28 | 10.10 | 33.98 | 757.75 | 20.44 | -9.30 | 29.73 | 12.88 | -4.30 | 14.32 | -4.30 | 665 | 100 | 30 | 47.49 | 279 | 93 | 275 | 93 |
| *B. sylvaticum* | Bsyl_846 | 47.80 | 27.20 | 9.35 | 9.84 | 30.45 | 860.40 | 26.20 | -6.11 | 32.31 | 18.35 | -0.20 | 19.63 | -1.37 | 554 | 87 | 25 | 45.95 | 233 | 79 | 225 | 83 |
| *B. sylvaticum* | Bsyl_847 | 40.90 | 14.70 | 11.80 | 7.12 | 30.33 | 614.70 | 24.76 | 1.29 | 23.48 | 8.99 | 19.54 | 19.54 | 4.72 | 817 | 112 | 29 | 37.98 | 297 | 96 | 96 | 256 |
| *B. sylvaticum* | Bsyl_848 | 45.40 | 9.90 | 12.52 | 9.40 | 31.23 | 780.66 | 28.29 | -1.80 | 30.09 | 18.20 | 2.74 | 21.98 | 2.74 | 890 | 109 | 44 | 22.18 | 279 | 173 | 225 | 173 |
| *B. sylvaticum* | Bsyl_849 | 38.10 | 13.40 | 18.37 | 6.94 | 33.01 | 529.93 | 30.01 | 8.99 | 21.01 | 16.77 | 24.84 | 25.25 | 12.50 | 463 | 69 | 3 | 58.64 | 197 | 24 | 58 | 146 |
| *B. sylvaticum* | Bsyl_850 | 46.50 | 11.40 | 9.37 | 9.34 | 33.08 | 717.38 | 24.63 | -3.60 | 28.23 | 18.30 | 0.74 | 18.30 | 0.74 | 718 | 88 | 28 | 37.43 | 259 | 96 | 259 | 96 |
| *B. sylvaticum* | Bsyl_851 | 46.70 | 9.00 | -1.30 | 5.96 | 28.56 | 548.06 | 10.04 | -10.83 | 20.86 | 5.52 | -7.25 | 5.71 | -7.25 | 1643 | 168 | 102 | 13.30 | 482 | 346 | 474 | 346 |
| *B. sylvaticum* | Bsyl_852 | 47.10 | 10.60 | 4.96 | 8.39 | 33.01 | 639.76 | 18.58 | -6.84 | 25.42 | 12.80 | -1.80 | 12.80 | -2.72 | 909 | 120 | 52 | 33.00 | 345 | 162 | 345 | 166 |
| *B. sylvaticum* | Bsyl_853 | 46.10 | 18.20 | 9.85 | 7.96 | 28.37 | 764.04 | 24.82 | -3.24 | 28.05 | 19.15 | 1.86 | 19.15 | 0.37 | 667 | 85 | 33 | 26.88 | 219 | 106 | 219 | 121 |
| *B. sylvaticum* | Bsyl_854 | 47.40 | 21.70 | 9.99 | 10.19 | 32.48 | 819.11 | 26.52 | -4.86 | 31.38 | 18.37 | 1.35 | 19.73 | -0.43 | 557 | 76 | 29 | 30.33 | 201 | 95 | 188 | 107 |
| *B. sylvaticum* | Bsyl_855 | 47.20 | 19.50 | 10.19 | 9.13 | 30.26 | 806.59 | 26.17 | -4.00 | 30.17 | 18.27 | 1.75 | 19.81 | 0.00 | 517 | 59 | 27 | 24.29 | 167 | 86 | 158 | 100 |
| *B. sylvaticum* | Bsyl_856 | 46.80 | 17.80 | 10.72 | 8.07 | 27.83 | 799.12 | 26.10 | -2.90 | 29.00 | 20.41 | 2.32 | 20.41 | 0.74 | 574 | 67 | 26 | 27.86 | 185 | 86 | 185 | 99 |
| *B. sylvaticum* | Bsyl_857 | 56.40 | 10.80 | 7.96 | 7.34 | 31.91 | 600.17 | 21.06 | -1.93 | 22.98 | 8.85 | 3.01 | 15.53 | 1.02 | 569 | 60 | 28 | 22.92 | 176 | 98 | 164 | 116 |
| *B. sylvaticum* | Bsyl_858 | 57.00 | 10.00 | 7.57 | 7.22 | 31.32 | 600.95 | 20.36 | -2.70 | 23.06 | 12.13 | 2.64 | 15.16 | 0.60 | 622 | 70 | 32 | 24.20 | 200 | 105 | 176 | 133 |
| *B. sylvaticum* | Bsyl_859 | 53.60 | 19.40 | 7.51 | 7.87 | 28.33 | 749.39 | 22.51 | -5.27 | 27.78 | 16.61 | -0.81 | 16.61 | -1.81 | 620 | 80 | 27 | 33.78 | 228 | 95 | 228 | 109 |
| *B. sylvaticum* | Bsyl_860 | 54.10 | 22.20 | 6.81 | 7.47 | 26.34 | 800.66 | 22.12 | -6.24 | 28.36 | 16.47 | -2.20 | 16.47 | -3.13 | 576 | 76 | 26 | 32.86 | 213 | 92 | 213 | 101 |
| *B. sylvaticum* | Bsyl_861 | 52.30 | 19.30 | 8.09 | 8.40 | 28.50 | 788.94 | 23.92 | -5.56 | 29.48 | 17.48 | -0.50 | 17.48 | -2.05 | 529 | 75 | 22 | 36.61 | 203 | 82 | 203 | 89 |
| *B. sylvaticum* | Bsyl_862 | 49.40 | 22.30 | 7.24 | 8.51 | 30.76 | 726.26 | 21.86 | -5.82 | 27.68 | 15.94 | -0.70 | 15.94 | -1.95 | 805 | 111 | 39 | 36.96 | 306 | 122 | 306 | 135 |
| *B. sylvaticum* | Bsyl_863 | 49.80 | 19.90 | 7.56 | 9.51 | 33.14 | 724.94 | 22.62 | -6.07 | 28.69 | 16.37 | -0.30 | 16.37 | -1.43 | 871 | 123 | 41 | 40.37 | 346 | 135 | 346 | 142 |
| *B. sylvaticum* | Bsyl_864 | 59.70 | 18.70 | 5.88 | 7.01 | 25.73 | 725.44 | 21.01 | -6.22 | 27.24 | 14.33 | 0.00 | 15.29 | -2.52 | 593 | 73 | 27 | 29.55 | 199 | 97 | 190 | 121 |
| *B. sylvaticum* | Bsyl_865 | 57.40 | 13.00 | 5.94 | 7.98 | 31.11 | 656.03 | 20.36 | -5.30 | 25.66 | 6.44 | 4.70 | 14.34 | -1.72 | 1058 | 112 | 59 | 22.67 | 332 | 191 | 260 | 275 |
| *B. sylvaticum* | Bsyl_866 | 58.20 | 12.80 | 6.57 | 7.40 | 28.35 | 682.70 | 21.10 | -5.00 | 26.10 | 6.98 | 1.21 | 15.25 | -1.46 | 718 | 80 | 39 | 23.26 | 228 | 125 | 201 | 157 |
| *B. sylvaticum* | Bsyl_867 | 57.80 | 13.40 | 5.79 | 7.72 | 29.51 | 672.78 | 20.45 | -5.73 | 26.18 | 6.14 | 0.51 | 14.41 | -2.07 | 807 | 88 | 45 | 21.40 | 246 | 144 | 223 | 188 |
| *B. sylvaticum* | Bsyl_868 | 58.50 | 13.50 | 6.26 | 7.69 | 28.57 | 707.72 | 21.19 | -5.75 | 26.94 | 14.08 | 0.84 | 15.26 | -2.10 | 617 | 70 | 32 | 23.67 | 194 | 107 | 185 | 127 |
| *B. sylvaticum* | Bsyl_869 | 58.60 | 14.60 | 6.81 | 6.67 | 27.13 | 654.65 | 20.68 | -3.90 | 24.58 | 15.36 | 1.25 | 15.36 | -0.49 | 580 | 67 | 30 | 26.72 | 193 | 96 | 193 | 98 |
| *B. sylvaticum* | Bsyl_870 | 58.20 | 15.20 | 5.98 | 7.80 | 28.96 | 701.16 | 21.24 | -5.71 | 26.95 | 13.85 | 0.54 | 14.99 | -2.19 | 563 | 72 | 27 | 28.19 | 191 | 96 | 188 | 112 |
| *B. sylvaticum* | Bsyl_871 | 57.30 | 14.50 | 5.73 | 8.12 | 30.19 | 691.86 | 20.98 | -5.93 | 26.90 | 13.43 | 0.48 | 14.58 | -2.38 | 733 | 81 | 41 | 20.47 | 219 | 132 | 209 | 166 |
| *B. sylvaticum* | Bsyl_872 | 56.90 | 15.00 | 5.98 | 7.94 | 30.45 | 674.93 | 20.81 | -5.26 | 26.07 | 13.56 | 0.74 | 14.65 | -1.84 | 680 | 77 | 38 | 20.83 | 207 | 122 | 202 | 151 |
| *B. sylvaticum* | Bsyl_873 | 56.50 | 13.00 | 7.59 | 7.46 | 30.66 | 649.32 | 21.10 | -3.24 | 24.34 | 14.92 | 2.50 | 15.76 | -0.06 | 805 | 92 | 44 | 23.35 | 253 | 147 | 243 | 181 |
| *B. sylvaticum* | Bsyl_874 | 55.50 | 13.10 | 8.02 | 6.15 | 26.68 | 639.35 | 20.99 | -2.08 | 23.07 | 8.88 | 2.81 | 16.08 | 0.58 | 589 | 62 | 33 | 18.82 | 176 | 113 | 153 | 141 |
| *B. sylvaticum* | Bsyl_875 | 56.40 | 14.40 | 6.44 | 7.70 | 30.47 | 659.01 | 20.88 | -4.38 | 25.26 | 6.77 | 1.35 | 14.87 | -1.23 | 749 | 76 | 45 | 18.87 | 222 | 142 | 203 | 181 |
| *B. sylvaticum* | Bsyl_876 | 60.00 | 10.90 | 4.83 | 7.44 | 26.37 | 759.38 | 20.55 | -7.67 | 28.22 | 4.82 | -0.65 | 14.57 | -4.13 | 929 | 108 | 46 | 26.89 | 310 | 162 | 264 | 183 |
| *B. sylvaticum* | Bsyl_877 | 59.80 | 9.60 | 3.93 | 8.66 | 28.62 | 800.11 | 20.77 | -9.48 | 30.25 | 8.91 | -1.32 | 14.08 | -5.70 | 843 | 102 | 38 | 29.10 | 279 | 141 | 254 | 148 |
| *B. sylvaticum* | Bsyl_878 | 58.30 | 8.30 | 6.57 | 7.11 | 28.70 | 647.00 | 20.20 | -4.56 | 24.76 | 7.21 | 9.45 | 14.82 | -1.03 | 1252 | 157 | 62 | 31.14 | 447 | 206 | 258 | 321 |
| *B. sylvaticum* | Bsyl_879 | 58.10 | 7.10 | 6.92 | 4.30 | 22.67 | 555.87 | 17.03 | -1.94 | 18.97 | 4.59 | 8.88 | 13.88 | 0.53 | 1813 | 225 | 81 | 35.88 | 649 | 260 | 327 | 529 |
| *B. sylvaticum* | Bsyl_880 | 59.10 | 6.80 | 1.77 | 5.26 | 23.69 | 618.61 | 14.38 | -7.82 | 22.20 | -1.01 | 3.75 | 9.86 | -4.96 | 1969 | 239 | 76 | 35.23 | 703 | 282 | 376 | 524 |
| *B. sylvaticum* | Bsyl_881 | 59.90 | 6.60 | 2.45 | 5.96 | 26.09 | 630.28 | 15.20 | -7.64 | 22.84 | -0.66 | 5.10 | 10.62 | -4.62 | 2111 | 264 | 83 | 35.99 | 741 | 282 | 393 | 674 |
| *B. sylvaticum* | Bsyl_882 | 60.40 | 7.20 | 0.36 | 5.97 | 25.53 | 655.05 | 13.06 | -10.32 | 23.38 | 0.92 | 3.15 | 8.90 | -6.86 | 1303 | 153 | 57 | 27.95 | 425 | 201 | 293 | 334 |
| *B. sylvaticum* | Bsyl_883 | 60.50 | 6.30 | 0.95 | 4.30 | 21.85 | 561.73 | 12.07 | -7.60 | 19.66 | -1.65 | 2.67 | 8.30 | -5.11 | 2598 | 318 | 101 | 35.73 | 901 | 349 | 487 | 817 |
| *B. sylvaticum* | Bsyl_884 | 60.40 | 5.40 | 5.24 | 4.75 | 27.01 | 487.55 | 15.06 | -2.52 | 17.58 | 2.84 | 7.12 | 11.51 | -0.18 | 3114 | 376 | 135 | 32.79 | 1062 | 451 | 591 | 940 |
| *B. sylvaticum* | Bsyl_885 | 61.40 | 7.60 | -1.58 | 5.73 | 25.44 | 625.22 | 10.98 | -11.54 | 22.52 | -7.37 | 0.90 | 6.73 | -8.45 | 1354 | 157 | 48 | 30.47 | 442 | 197 | 298 | 415 |
| *B. sylvaticum* | Bsyl_886 | 61.00 | 5.60 | 3.35 | 3.59 | 19.88 | 538.77 | 13.83 | -4.21 | 18.04 | 0.71 | 5.20 | 10.43 | -2.52 | 3608 | 444 | 155 | 35.42 | 1241 | 501 | 643 | 1142 |
| *B. sylvaticum* | Bsyl_887 | 61.60 | 5.70 | 5.22 | 4.18 | 22.42 | 544.05 | 15.94 | -2.70 | 18.64 | 2.19 | 7.48 | 12.32 | -0.85 | 2924 | 359 | 117 | 38.07 | 1027 | 393 | 483 | 947 |
| *B. sylvaticum* | Bsyl_888 | 62.60 | 8.80 | 0.79 | 6.02 | 27.57 | 589.45 | 13.27 | -8.58 | 21.85 | -2.46 | 3.08 | 8.58 | -5.58 | 1076 | 120 | 47 | 25.42 | 331 | 176 | 262 | 310 |
| *B. sylvaticum* | Bsyl_889 | 63.10 | 8.50 | 5.77 | 5.25 | 28.19 | 511.61 | 16.18 | -2.45 | 18.63 | 3.01 | 7.78 | 12.38 | 0.06 | 1306 | 156 | 57 | 28.80 | 431 | 208 | 265 | 389 |
| *B. sylvaticum* | Bsyl_890 | 63.60 | 9.00 | 6.23 | 5.14 | 27.90 | 509.68 | 16.57 | -1.87 | 18.44 | 3.48 | 8.22 | 12.68 | 0.38 | 1194 | 139 | 54 | 29.17 | 392 | 187 | 253 | 352 |
| *B. sylvaticum* | Bsyl_891 | 54.70 | 39.80 | 5.00 | 8.76 | 23.70 | 1063.16 | 24.20 | -12.76 | 36.96 | 17.67 | -7.05 | 17.67 | -8.26 | 568 | 80 | 26 | 31.65 | 202 | 91 | 202 | 107 |
| *B. sylvaticum* | Bsyl_892 | 35.00 | 32.80 | 13.57 | 10.96 | 37.70 | 684.19 | 29.45 | 0.38 | 29.07 | 5.38 | 21.87 | 21.87 | 5.38 | 750 | 154 | 7 | 85.94 | 418 | 24 | 24 | 418 |
| *B. sylvaticum* | Bsyl_893 | 37.00 | 30.00 | 9.69 | 12.76 | 38.27 | 771.35 | 28.28 | -5.06 | 33.34 | 0.37 | 19.08 | 19.19 | 0.37 | 558 | 90 | 13 | 55.74 | 249 | 43 | 60 | 249 |
| *B. sylvaticum* | Bsyl_894 | 41.30 | 31.40 | 12.95 | 7.43 | 32.95 | 574.30 | 24.85 | 2.30 | 22.55 | 8.04 | 14.80 | 20.07 | 6.10 | 1067 | 151 | 53 | 33.70 | 387 | 167 | 205 | 283 |
| *B. sylvaticum* | Bsyl_895 | 41.40 | 41.40 | 14.58 | 7.78 | 34.58 | 574.48 | 26.42 | 3.92 | 22.50 | 12.61 | 12.36 | 21.74 | 7.96 | 1990 | 254 | 86 | 33.95 | 715 | 304 | 501 | 457 |
| *B. sylvaticum* | Bsyl_896 | 36.10 | 9.60 | 17.57 | 11.49 | 36.95 | 706.35 | 35.55 | 4.44 | 31.11 | 9.49 | 26.73 | 26.73 | 9.49 | 470 | 59 | 7 | 43.03 | 172 | 42 | 42 | 172 |
| *B. sylvaticum* | Bsyl_897 | 36.90 | 4.00 | 17.04 | 9.35 | 38.81 | 537.47 | 30.26 | 6.17 | 24.09 | 12.09 | 23.54 | 24.08 | 10.91 | 952 | 177 | 1 | 72.32 | 475 | 28 | 56 | 432 |
| *B. sylvaticum* | Bsyl_898 | 36.30 | 8.00 | 15.40 | 11.41 | 38.84 | 657.72 | 32.23 | 2.84 | 29.38 | 7.85 | 23.67 | 23.82 | 7.85 | 778 | 120 | 5 | 61.27 | 343 | 36 | 54 | 343 |
| *B. sylvaticum* | Bsyl_899 | 35.80 | -5.90 | 17.55 | 6.80 | 37.18 | 406.90 | 26.98 | 8.70 | 18.28 | 14.02 | 22.35 | 22.82 | 12.79 | 780 | 132 | 1 | 75.02 | 372 | 15 | 23 | 338 |
| *B. sylvaticum* | Bsyl_900 | 31.20 | -8.00 | 12.58 | 11.43 | 39.62 | 605.02 | 28.23 | -0.61 | 28.84 | 8.64 | 20.18 | 20.29 | 5.39 | 512 | 71 | 3 | 57.52 | 196 | 23 | 28 | 171 |
| *B. sylvaticum* | Bsyl_901 | 35.50 | -5.40 | 16.60 | 9.77 | 39.02 | 531.08 | 30.41 | 5.36 | 25.04 | 10.37 | 23.32 | 23.48 | 10.37 | 814 | 149 | 0 | 85.93 | 424 | 9 | 16 | 424 |
| *B. sylvaticum* | Bsyl_902 | 28.50 | -13.90 | 20.54 | 8.06 | 52.28 | 264.96 | 28.73 | 13.32 | 15.41 | 18.42 | 22.02 | 24.05 | 17.51 | 122 | 29 | 0 | 85.80 | 67 | 2 | 6 | 66 |
| *B. sylvaticum* | Bsyl_903 | 36.10 | 51.80 | 6.28 | 11.02 | 29.10 | 982.05 | 26.22 | -11.66 | 37.88 | 5.27 | 17.59 | 18.15 | -6.05 | 230 | 45 | 2 | 68.44 | 116 | 14 | 21 | 65 |
| *B. sylvaticum* | Bsyl_904 | 36.40 | 44.40 | 17.60 | 12.16 | 30.45 | 1015.68 | 39.13 | -0.80 | 39.93 | 6.55 | 30.13 | 30.13 | 5.18 | 852 | 187 | 0 | 98.11 | 489 | 0 | 0 | 461 |
| *B. sylvaticum* | Bsyl_905 | 58.40 | 22.90 | 6.49 | 6.34 | 23.27 | 763.45 | 21.40 | -5.84 | 27.24 | 7.47 | 0.06 | 16.19 | -2.26 | 599 | 70 | 31 | 32.50 | 208 | 95 | 172 | 120 |
| *B. sylvaticum* | Bsyl_906 | 59.20 | 27.60 | 4.75 | 7.70 | 23.66 | 882.84 | 22.12 | -10.43 | 32.55 | 14.43 | -5.41 | 15.66 | -6.13 | 627 | 80 | 27 | 36.76 | 232 | 90 | 216 | 105 |
| *B. sylvaticum* | Bsyl_907 | 56.30 | 21.10 | 6.92 | 6.32 | 25.00 | 725.95 | 20.36 | -4.91 | 25.28 | 7.87 | 1.16 | 15.91 | -1.73 | 721 | 86 | 29 | 34.39 | 245 | 104 | 218 | 143 |
| *B. sylvaticum* | Bsyl_908 | 56.80 | 22.60 | 6.05 | 7.22 | 26.38 | 749.05 | 21.14 | -6.24 | 27.38 | 14.41 | 0.34 | 15.49 | -2.83 | 666 | 81 | 30 | 29.65 | 225 | 109 | 218 | 126 |
| *B. sylvaticum* | Bsyl_909 | 57.20 | 23.10 | 6.38 | 7.42 | 25.85 | 749.09 | 22.10 | -6.60 | 28.70 | 14.95 | -2.04 | 15.93 | -2.43 | 635 | 75 | 29 | 30.73 | 218 | 98 | 210 | 114 |
| *B. sylvaticum* | Bsyl_910 | 57.40 | 24.70 | 5.34 | 6.82 | 23.55 | 839.30 | 20.96 | -8.00 | 28.96 | 14.55 | -4.16 | 15.69 | -4.84 | 671 | 80 | 26 | 33.17 | 236 | 97 | 219 | 116 |
| *B. sylvaticum* | Bsyl_911 | 56.90 | 26.10 | 4.93 | 7.83 | 25.69 | 841.27 | 21.40 | -9.08 | 30.48 | 13.91 | -1.15 | 15.29 | -5.42 | 770 | 93 | 37 | 25.20 | 252 | 134 | 242 | 154 |
| *B. sylvaticum* | Bsyl_912 | 53.60 | 25.80 | 5.33 | 6.85 | 23.45 | 833.44 | 20.51 | -8.70 | 29.21 | 15.44 | -4.16 | 15.44 | -5.13 | 715 | 86 | 38 | 25.00 | 239 | 127 | 239 | 136 |
| *B. sylvaticum* | Bsyl_913 | 54.60 | 41.50 | 4.76 | 8.32 | 22.57 | 1057.92 | 23.90 | -12.94 | 36.85 | 17.44 | -1.74 | 17.44 | -8.41 | 575 | 80 | 30 | 29.92 | 200 | 94 | 200 | 118 |
| *B. sylvaticum* | Bsyl_914 | 54.50 | 36.20 | 4.93 | 8.21 | 24.39 | 966.11 | 22.75 | -10.92 | 33.66 | 16.65 | -1.12 | 16.65 | -6.99 | 625 | 83 | 31 | 31.73 | 227 | 101 | 227 | 117 |
| *B. sylvaticum* | Bsyl_915 | 54.30 | 35.30 | 4.86 | 8.02 | 23.75 | 977.60 | 22.50 | -11.29 | 33.79 | 16.63 | -6.02 | 16.63 | -7.23 | 621 | 83 | 30 | 33.67 | 230 | 97 | 230 | 109 |
| *B. sylvaticum* | Bsyl_916 | 54.30 | 42.20 | 4.56 | 8.38 | 22.33 | 1082.64 | 23.87 | -13.65 | 37.52 | 17.46 | -2.13 | 17.46 | -8.98 | 550 | 77 | 29 | 30.79 | 194 | 89 | 194 | 110 |
| *B. sylvaticum* | Bsyl_917 | 54.50 | 39.10 | 4.84 | 8.51 | 23.79 | 1046.30 | 23.71 | -12.04 | 35.75 | 17.44 | -6.87 | 17.44 | -8.01 | 558 | 80 | 25 | 33.50 | 205 | 87 | 205 | 102 |
| *B. sylvaticum* | Bsyl_918 | 54.00 | 40.60 | 5.37 | 8.93 | 22.54 | 1110.61 | 25.80 | -13.82 | 39.62 | 18.52 | -2.25 | 18.52 | -8.52 | 544 | 78 | 28 | 31.29 | 191 | 86 | 191 | 106 |
| *B. sylvaticum* | Bsyl_919 | 56.70 | 37.30 | 4.25 | 8.02 | 22.37 | 988.31 | 22.35 | -13.50 | 35.86 | 16.13 | -1.81 | 16.13 | -8.22 | 642 | 88 | 28 | 34.84 | 236 | 94 | 236 | 114 |
| *B. sylvaticum* | Bsyl_920 | 56.20 | 36.90 | 4.11 | 7.95 | 22.93 | 978.27 | 22.10 | -12.57 | 34.67 | 16.02 | -6.73 | 16.02 | -8.01 | 646 | 90 | 27 | 36.62 | 244 | 94 | 244 | 112 |
| *B. sylvaticum* | Bsyl_921 | 55.20 | 39.00 | 4.86 | 8.20 | 23.08 | 1022.35 | 23.61 | -11.94 | 35.55 | 17.36 | -2.33 | 17.36 | -7.51 | 589 | 85 | 28 | 34.27 | 215 | 91 | 215 | 108 |
| *B. sylvaticum* | Bsyl_922 | 56.00 | 35.70 | 3.90 | 8.00 | 22.56 | 987.49 | 22.08 | -13.36 | 35.45 | 15.88 | -7.37 | 15.88 | -8.21 | 636 | 84 | 28 | 35.36 | 236 | 92 | 236 | 107 |
| *B. sylvaticum* | Bsyl_923 | 55.70 | 36.40 | 4.27 | 7.93 | 23.11 | 971.61 | 22.22 | -12.11 | 34.33 | 16.13 | -6.49 | 16.13 | -7.69 | 643 | 85 | 29 | 33.69 | 236 | 97 | 236 | 114 |
| *B. sylvaticum* | Bsyl_924 | 55.90 | 38.50 | 5.29 | 8.22 | 21.19 | 1106.89 | 25.67 | -13.14 | 38.81 | 18.70 | -4.92 | 18.70 | -8.62 | 653 | 91 | 30 | 34.29 | 238 | 98 | 238 | 103 |
| *B. sylvaticum* | Bsyl_925 | 52.60 | 38.40 | 5.44 | 8.67 | 24.03 | 1048.11 | 24.22 | -11.87 | 36.08 | 17.95 | -6.62 | 17.95 | -7.36 | 572 | 78 | 29 | 30.93 | 208 | 97 | 208 | 111 |
| *B. sylvaticum* | Bsyl_926 | 50.70 | 37.00 | 6.38 | 7.70 | 21.84 | 1020.36 | 24.40 | -10.84 | 35.24 | 18.52 | -0.12 | 18.52 | -6.40 | 579 | 70 | 34 | 21.66 | 189 | 110 | 189 | 122 |
| *B. sylvaticum* | Bsyl_927 | 54.90 | 44.70 | 4.14 | 8.22 | 21.56 | 1110.97 | 23.78 | -14.34 | 38.12 | 17.44 | -8.27 | 17.44 | -9.64 | 552 | 73 | 25 | 30.32 | 190 | 87 | 190 | 105 |
| *B. sylvaticum* | Bsyl_928 | 55.10 | 43.90 | 3.54 | 8.34 | 21.70 | 1103.57 | 23.42 | -15.02 | 38.44 | 16.78 | -9.21 | 16.78 | -10.24 | 516 | 76 | 23 | 36.42 | 189 | 76 | 189 | 88 |
| *B. sylvaticum* | Bsyl_929 | 54.20 | 45.20 | 4.51 | 8.43 | 21.09 | 1145.51 | 24.50 | -15.46 | 39.96 | 17.99 | -8.74 | 17.99 | -9.77 | 515 | 70 | 24 | 30.91 | 181 | 83 | 181 | 93 |
| *B. sylvaticum* | Bsyl_930 | 55.60 | 58.40 | 2.22 | 9.54 | 22.91 | 1195.76 | 23.37 | -18.28 | 41.65 | 16.43 | -10.94 | 16.43 | -12.83 | 550 | 83 | 20 | 43.89 | 220 | 71 | 220 | 83 |
| *B. sylvaticum* | Bsyl_931 | 45.10 | 42.00 | 9.74 | 9.28 | 27.23 | 930.84 | 27.88 | -6.22 | 34.10 | 19.04 | -0.79 | 21.07 | -1.76 | 535 | 80 | 24 | 35.33 | 197 | 77 | 185 | 86 |
| *B. sylvaticum* | Bsyl_932 | 44.00 | 42.00 | 6.82 | 11.49 | 34.98 | 795.56 | 23.47 | -9.37 | 32.84 | 14.52 | -3.23 | 16.32 | -3.23 | 661 | 106 | 21 | 52.39 | 279 | 69 | 275 | 69 |
| *B. sylvaticum* | Bsyl_933 | 43.50 | 41.80 | 2.45 | 9.06 | 32.44 | 677.82 | 17.54 | -10.40 | 27.94 | 5.43 | -5.46 | 10.73 | -5.65 | 1262 | 125 | 65 | 19.98 | 364 | 214 | 355 | 245 |
| *B. sylvaticum* | Bsyl_934 | 43.20 | 44.30 | 10.15 | 9.51 | 28.59 | 895.20 | 27.27 | -5.98 | 33.25 | 18.98 | -1.07 | 20.96 | -1.07 | 669 | 99 | 30 | 41.02 | 266 | 96 | 236 | 96 |
| *B. sylvaticum* | Bsyl_935 | 43.00 | 45.70 | 9.70 | 9.06 | 27.94 | 856.73 | 26.50 | -5.94 | 32.44 | 18.04 | -1.03 | 20.08 | -1.03 | 561 | 89 | 21 | 52.33 | 244 | 65 | 231 | 65 |
| *B. sylvaticum* | Bsyl_936 | 42.80 | 47.20 | 10.31 | 9.27 | 27.76 | 894.22 | 28.05 | -5.36 | 33.41 | 19.20 | -0.62 | 21.36 | -0.62 | 443 | 58 | 17 | 40.97 | 165 | 59 | 155 | 59 |
| *B. sylvaticum* | Bsyl_937 | 59.50 | 39.10 | 2.67 | 7.84 | 21.78 | 1005.14 | 21.64 | -14.38 | 36.02 | 15.06 | -8.23 | 15.06 | -9.76 | 553 | 77 | 22 | 38.99 | 216 | 77 | 216 | 88 |
| *B. sylvaticum* | Bsyl_938 | 49.10 | 23.50 | 6.69 | 8.85 | 30.71 | 746.78 | 21.32 | -7.48 | 28.81 | 13.81 | -1.93 | 15.59 | -2.76 | 902 | 130 | 40 | 41.38 | 359 | 136 | 358 | 144 |
| *B. sylvaticum* | Bsyl_939 | 52.30 | 12.40 | 9.33 | 8.36 | 31.87 | 685.15 | 24.14 | -2.10 | 26.23 | 16.35 | 4.81 | 17.85 | 1.07 | 558 | 61 | 34 | 20.00 | 176 | 111 | 176 | 122 |
