## Appendix 2 for "A Palearctic divide, niche conservatism and host-fungal endophyte interactions shaped the phylogeography of the grass *Brachypodium sylvaticum*"

### **2. Expanded Materials and methods**

#### ***2.1 Sampling and genome skimming sequencing***

A total of 94 samples belonging to the *B. sylvaticum* complex were selected for the genomic study. Sampling included 85 individuals of *B. sylvaticum* *s. s.*, four *B. miserum,* two *B. breviglume,* and one *B. kurilense, B. spryginii* and *B. glaucovirens*. The individuals were strategically collected across the respective native ranges, aiming to sample a comprehensive representation of the species' genetic, geographical, and ecological diversities (Figure 1, Appendix 1). Six samples of other diploid *Brachypodium* species were selected as appropriate outgroups (two *B. stacei,* two *B. distachyon* and two *B. arbuscula*) and three samples of the closely related *B. pinnatum* were selected as close ingroups, to adequately reconstruct the evolutionary framework of the *B. sylvaticum* complex samples under study (Catalán et al., 2023). Total genomic DNA was extracted from silica gel dried leaf tissue or herbarium specimens using the CTAB method (Doyle & Doyle, 1987), with modifications to improve yield and purity (Decena et al., 2024).

##### ***2.******2 Plastome Assembly and annotation***

Whole plastome sequences of the studied *B. sylvaticum* complex samples were assembled from the filtered genome skimming reads using Novoplasty 4.3.3 (Dierckxsens et al., 2017) and *B. sylvaticum* Ain-1 reference plastome (https://phytozome-next.jgi.doe.gov/) as a seed to initiate the extension and assembly of the plastome samples under study. *de novo* annotation of plastome genes was performed using GeSeq (Tillich et al., 2017) implemented in Chlorobox (https://chlorobox.mpimp-golm.mpg.de/)

*2.3 Brachypodium sylvaticum complex* and *Epichloë sylvatica read-filtering from Illumina genome skimming raw reads*

To analyze genomic variation and gene sequences in the *B. sylvaticum* complex samples and their associated *E. sylvatica* fungal endophytes, we employed a customized bioinformatic pipeline (SplitReads.sh). This workflow facilitated the mapping reads to both grass and endophyte reference genomes, the extraction of gene sequences, and the filtering of genomic variants (SNPs). Initially, raw reads were mapped to a concatenation of two reference genomes, those of *Brachypodium sylvaticum* Ain1 (https://phytozome-next.jgi.doe.gov/info/B*sylvaticum*Ain_1_v2_1, Lei et al., 2024) and *Epichloë sylvatica* ASM100826v1 (https://www.ncbi.nlm.nih.gov/datasets/genome/GCA_001008265.1/), using Minimap2 ( Li, 2018). The output alignment files were sorted and indexed and filtered into host ant fungal alingments using Samtools (Danecek et al., 2021).

##### ***2.4 Gene sequence extraction and SNP detection in host Brachypodium sylvaticum complex samples***

Mapped reads for host *B. sylvaticum* complex taxa were aligned to a concatenated CDSs reference from *B. sylvaticum* Ain1 to extract gene sequences for our *Brachypodium* samples. Consensus sequences were generated using Samtools, ensuring a minimum depth threshold of 10 reads per position to account for sequencing errors, obtaining relatively high-confidence host gene coding sequences for subsequent analyses. To identify genetic variants, a SNP-calling workflow was implemented using BCFtools (Danecek et al., 2021). Reads were processed with bcftools mpileup, using the reference genome to generate per-base sequencing depth and variant likelihoods. SNPs and small indels were called using bcftools and individual VCF files were merged using bcftools merge to create a consensus dataset of variants. We then used VCFtools (Danecek et al., 2011) to retain only biallelic SNPs and a maximum missing data threshold of 10%. The combined VCF file was subsequently filtered for linkage disequilibrium (LD) using plink2 (Chang et al., 2015), with an LD threshold of 0.2, minimum allelic frequency (MAF) of 0.05, and a sliding window of 10,000 bp.

##### ***2.5 ITS sequence extraction for Epichloë sylvatica endophytes from the holobiont samples***

The pre-filtered *Epichloë sylvatica* reads were further mapped against a concatenated CDSs reference from *E. festucae* NCBI ID:35717 to obtain gene annotations for comparative analysis. However, given the higher recovery of ribosomal genes over single-copy genes, an additional round of mapping was performed using only the complete rDNA 35S cistron (18S + ITS1 + 5.8S + ITS2 + 28S) to refine the detection and characterization of the *E.* *sylvatica* 35S sequences. To extract high-confidence fungal sequences, consensus sequences were generated using Samtools. Furthermore, the obtained data were subjected to BLAST analysis to verify that the sequences belonged to *Epichloë* rather than representing potential contaminations, other endophytes, or chimeric sequences.

#### ***2.6 Population genomics and phylogeographic analyses of host Brachypodium sylvaticum complex plants***

Phylogenomic trees of host plants were constructed using maximum likelihood (ML) methods to infer the evolutionary relationships among the studied *B. sylvaticum* complex taxa and populations. From an initial variant matrix comprising all high quality mapped sites (8,372,248), we pruned for maximum missing data per site ≤ 10%, linkage disequilibrium pruning (r² < 0.2) and minor allele frequency ≥ 0.05, yielding a final panel of 160,124 SNPs used to build an ML tree using IQ-TREE2 (Nguyen et al., 2015) with the best substitution model selected by the program, and 1000 ultrafast bootstrap replicates to assess branch support. Similarly, a ML phylogenomic tree was constructed for the whole plastome sequence dataset using IQ-TREE2. To investigate potential cytonuclear topological discordances between the nuclear and plastome genomes, we tested for potential cophylogenetic trees using the R packages ape and phytools. To quantify the extent of discordance, we employed the generalized Robinson–Foulds (gRF) distance, as implemented in the TreeDist R package (Smith, 2020).

A divergence dating analysis of the nuclear *B. sylvaticum* complex tree was performed using TreePL (Smith & O’Meara, 2012), a penalized likelihood method for estimating divergence times. The input topology was the maximum-likelihood phylogeny constructed from the concatenated nuclear SNP dataset (160,124 SNPs). Secondary calibrations were imposed on key nodes, following Catalán et al. (2023) and Sancho et al. (2018) estimations for the *Brachypodium* crown ancestor split (11.6 Ma) and the core-perennial ancestor split (2.71 Ma). The optimal smoothing parameter was determined using cross-validation. Additionally, we generated 1,000 bootstrap trees from the SNP dataset and dated each with TreePL to assess temporal uncertainty.

To infer the phylogeographic patterns of the *B. sylvaticum* complex population groups, we applied the BioGeoBEARS R package (Matzke, 2013), which implements likelihood-based models to reconstruct ancestral areas and infer biogeographic events such as dispersal, extinction, and vicariance. We performed the analysis on the dated phylogeny allowing null ranges and setting the maximum range size to nine. Geographic areas were manually defined based on the current distribution of the sampled populations, and their main evolutionary lineages (see Results), covering an East-to-West gradient: East Palearctic group (Kuril Islands, Japan, Himalayas), and West Palearctic group (East Europe, Middle East, Eastern Mediterranean, British Islands, Pyrenees + Alps, and Morocco + Southern Spain). To evaluate the fit of alternative biogeographic models, we performed model selection among the standard DEC, DIVALIKE, and BAYAREALIKE models, as well as their respective +J variants that incorporate founder-event speciation (jump dispersal). Model comparisons were based on log-likelihood and Akaike Information Criterion (AIC) scores, which were extracted and compared across models.

To investigate population structure and genomic ancestry, we employed ADMIXTURE v1.3.0 (Alexander et al., 2009) using the 160,124 nuclear SNPs*.* This dataset was reduced to include only *B. sylvaticum* complex samples. This approach estimates individual ancestry by iteratively updating allele frequencies and ancestry fractions. We tested hypothetical genomic groups ranging from K=1 to K=10 and selected the best K using StructureSelector (Li & Liu, 2018), The selection of the best K was based on Puechmaille’s method (Puechmaille, 2016) .

#### ***2.7 Phylogenetic analysis of Epichloë sylvatica endophytes and potential*** ***plant-endophyte co-evolution***

To investigate the potential co-evolution between the *B. sylvaticum* host plant and their
*E. sylvatica* fungal endophytes, we also constructed an *Epichloë* ML phylogenetic tree using the *E. sylvatica* ITS region (ITS1 + 5.8S + ITS2) alignment. We employed the Procrustes Application to Cophylogenetic (PACo; Balbuena et al., 2013) analysis to evaluate host-endophyte phylogenetic congruence using the PACo package and a custom PACo.R script. For each 1,000 ultrafast‑bootstrap trees of hosts and endophytes we computed patristic distance matrices, then applied PACo with 1,000 permutation replicates per tree to extract global goodness‑of‑fit statistics (m² and p‑values) and squared residuals for each host–endophyte link. Residuals were normalized within each replicate, and for each link we examined the distribution of its 1,000 normalized residuals by binning into quartiles. Discordance for a given link was quantified as the combined proportion of replicates falling in the third and fourth quartiles (i.e., > 50th and > 75th percentiles). We also performed a statistical ParaFit analysis of coevolution (Legendre et al., 2002), which test globally and individually for specific host-endophyte interactions (links). Parafit assumes the null hypothesis of independent evolutionary pathways versus the alternative hypothesis of coevolution; the analysis is based on patristic distances of both host and endophyte trees, and a host-endophyte interaction matrix over 1000 replicates is conducted using the ape package and the Parafit script (ParaFit.R) in R.

#### ***2.8 Genomic diversity of Brachypodium sylvaticum complex taxa and genomic groups and impact of spatial isolation and environment isolation***

Genomic diversity metrics, including observed heterozygosity (Ho), expected heterozygosity (He), and the inbreeding coefficient (Fis), were estimated for the *B. sylvaticum* complex taxa and genomic groups retrieved from Admixture using the 160,124 SNP dataset with plink2 (Chang et al., 2015). The selfing rate (s) was further calculated as s = 2Fis/(1 + Fis) following Ritland (1990). To explore patterns of isolation by distance (IBD) and isolation by environment (IBE) in the
*B. sylvaticum* complex populations, we first classified individuals into their corresponding populations (Table S1) using the dartR v2.9.7 library (Mijangos et al., 2022). We then computed pairwise geographic distances between individuals using the Haversine method with the geosphere v1.5-20 package, and pairwise bioclimatic distances based on a set of 19 bioclimatic variables retrieved from worldclim (current climate) using stats package in R. Pairwise genetic distances were estimated using Manhattan distances between individuals using the vegan R package.

To assess the relative contributions of geographic and environmental factors to the genomic differentiation of *B. sylvaticum* complex populations, we applied distance-based redundancy analyses (dbRDA) using the capscale function from the vegan R package. Genetic, geographic, and bioclimatic distance matrices were computed and transformed via Principal Coordinates Analysis (PCoA); axes of each variable were used as predictors. We fitted three models: (i) a full model including both environmental and geographic predictors, (ii) an environment-only model (IBE), and (iii) a geography-only model (IBD). The significance of each model and of individual predictors was assessed using permutation tests (n = 10,000). To disentangle the individual effects of IBE and IBD, we compared nested models using partial dbRDA, evaluating the contribution of environmental distances while controlling geography, and vice versa. This approach allowed us to test whether bioclimatic differences significantly explained genetic structure after accounting for spatial autocorrelation (indicative of IBE), and whether geographic distances explained structure independently of environmental effects (indicative of IBD). Variance partitioning and ANOVA comparisons further quantified the unique and shared contributions of both types of distances to population genomic differentiation.

#### ***2.9 Environmental niche modeling analysis of the Brachypodium sylvaticum complex***

We performed environmental niche modeling (ENM) with occurrence data from the 94
*B. sylvaticum* complex samples used in the genomic study. To further increase the coverage of the species distributions, we also incorporated occurrence records downloaded from GBIF of taxonomically studied vouchers: 168 for *B. breviglume*, three for *B. glaucovirens*, 28 for *B. kurilense*, 921 for *B. sylvaticum s.s.*, and 48 for *B. miserum* (Figure 1; Table S1; Appendix 1). To minimize sampling bias, occurrences with pair-distance <50 km were excluded using the 'thin' function from the R package spThin (Aiello‐Lammens et al., 2015). The final dataset included 1,262 confirmed occurrence sites for *B. sylvaticum* complex samples classified into their respective taxa. Niche models were constructed for the whole *B. sylvaticum* complex dataset and separately for the Eastern Palearctic and Western Palearctic groups.

A set of 19 bioclimatic variables was retrieved from the WorldClim database (http://www.worldclim.org) at a spatial resolution of 2.5 arc minutes (~4.5 km² at the equator) for two different geoclimatic scenarios: present and Last Glacial Maximum (LGM, ~21 kya). The ENM analysis was conducted using the Maxent 3.4.4 algorithm (Steven J Phillips et al., 2017) implemented in R through the ENMTools (Warren et al., 2021) and Maxnet (Phillips et al., 2021) packages. The most suitable climatic variables were selected by removing highly correlated variables based on the Variance Inflation Factor (VIF). To further mitigate sampling bias, we overlapped 100 independent Maxnet models with climate data through bootstrap analysis. In each model iteration, 75% of the occurrence records were used for training, while 25% were used for testing. Model performance was assessed using the area under the curve (AUC) metric, which evaluates the discrimination ability of a model against a random one (Lawson et al., 2014). To eliminate areas with low environmental suitability and reduce overprediction, a presence threshold based on a 90% sensitivity criterion was applied, following model evaluation with presence–background data using the dismo package.

To evaluate the potential effect of the Quaternary glacial on the expansions or contractions of the *B. sylvaticum* complex niche, we constructed ENMs for Last Glacial Maximum (LGM, ~22,000 ya), using paleoclimatic data from the ‘Community Climate System Model’ at 2.5 min resolution (CCSM; Gent et al., 2011) and assuming that the species' climate-driven biological requirements were the same in the past as today.

***2.10 Comparing environmental niches and testing niche overlaps of the main
B. sylvaticum groups***

We compared the current environmental conditions of the *B. sylvaticum* complex*,* Eastern and Western groups, aiming to detect significant differences among them across 19 bioclimatic variables. In addition, we explored potential environmental variation within the *B. sylvaticum* complex using the same set of variables. Statistical analyses were performed using non-parametric Kruskal–Wallis tests followed by pairwise multiple comparisons to identify climatic differences among groups and Principal Component Analyses (PCA) to discriminate major environmental gradients, using the ecospat package in R (Broennimann et al., 2012).

We used a niche comparison framework based on niche overlaps and niche identity tests, aiming to evaluate niche conservatism *vs* divergence between the two main *B. sylvaticum* groups (see Results). Niche overlap between the Eastern and Western groups was quantified using Schoener’s D metric and the Hellinger-based I index, both ranging from 0 (no overlap) to 1 (complete overlap). Additionally, niche breadth was calculated using Levins' B2 standardized index. To assess the significance of niche differences, we performed niche equivalency and niche similarity tests, based on 1000 random permutations, which evaluate whether observed niche overlaps are greater or lower than expected by chance. Following Warren et al. (2021), we interpreted significantly low overlap in the similarity test (P < 0.05 or P < 0.01) as evidence of ecological niche differentiation, while non-significant results were associated with niche conservatism. We further compared niches from the same Last Glacial Maximum (LGM) group with present-day ones to assess changes in both niche breadth and overlap.
