## Supplementary material for "A Palearctic divide, niche conservatism and host-fungal endophyte interactions shaped the phylogeography of the grass *Brachypodium sylvaticum*": Table S4

**Table S4.** Results of Procrustean Approach to Cophylogenetics (PACo) analysis for individual *Brachypodium sylvaticum* complex *– Epichloë* *sylvatica* associations. The sum of squared normalized residual values represents the deviation of each host–endophyte link from the expected coevolutionary pattern, with higher residuals indicating greater phylogenetic incongruence. Associations are classified as Congruent (median residual ≤ 0.012) or Incongruent (median residual > 0.012), using 0.012 as the threshold defined by the combined Q3+Q4 quartiles of the bootstrap distributions.

| **Global M2:** 0.8310 | | **P-Value:** 0.004 | |
| --- | --- | --- | --- |
| ***Brachypodium sylvaticum s. l.*** | ***Epichloë sylvatica*** | **Sum of squared normalized residuals** | **Congruence** |
| Bkurilense | Esyl_Bkurilense | 0.0146 | Incongruent |
| Bspryginii | Esyl_Bspryginii | 0.0167 | Incongruent |
| Bsyl29H | Esyl_Bsyl29H | 0.0149 | Incongruent |
| Bsyl30H | Esyl_Bsyl30H | 0.0153 | Incongruent |
| Bsyl31H | Esyl_Bsyl31H | 0.0184 | Incongruent |
| Bsyl32H | Esyl_Bsyl32H | 0.0147 | Incongruent |
| Bsyl35H | Esyl_Bsyl35H | 0.0147 | Incongruent |
| Bsyl36H | Esyl_Bsyl36H | 0.0153 | Incongruent |
| **Bsyl37H** | **Esyl_Bsyl37H** | **0.0102** | **Congruent** |
| **Bsyl38H** | **Esyl_Bsyl38H** | **0.0058** | **Congruent** |
| Bsyl39H | Esyl_Bsyl39H | 0.0137 | Incongruent |
| Bsyl41H | Esyl_Bsyl41H | 0.0147 | Incongruent |
| **Bsyl466-13** | **Esyl_Bsyl466-13** | **0.0117** | **Congruent** |
| **Bsyl466-2** | **Esyl_Bsyl466-2** | **0.0102** | **Congruent** |
| **Bsyl466-7B** | **Esyl_Bsyl466-7B** | **0.0098** | **Congruent** |
| **Bsyl467-10** | **Esyl_Bsyl467-10** | **0.0107** | **Congruent** |
| **Bsyl467-2** | **Esyl_Bsyl467-2** | **0.0120** | **Congruent** |
| **Bsyl467-7** | **Esyl_Bsyl467-7** | **0.0117** | **Congruent** |
| **Bsyl467-9B** | **Esyl_Bsyl467-9B** | **0.0097** | **Congruent** |
| Bsyl470-4 | Esyl_Bsyl470-4 | 0.0192 | Incongruent |
| Bsyl470-8B | Esyl_Bsyl470-8B | 0.0190 | Incongruent |
| Bsyl470-9 | Esyl_Bsyl470-9 | 0.0168 | Incongruent |
| Bsyl476-10 | Esyl_Bsyl476-10 | 0.0137 | Incongruent |
| Bsyl476-11 | Esyl_Bsyl476-11 | 0.0144 | Incongruent |
| Bsyl476-14 | Esyl_Bsyl476-14 | 0.0137 | Incongruent |
| Bsyl476-9 | Esyl_Bsyl476-9 | 0.0161 | Incongruent |
| **Bsyl477-1** | **Esyl_Bsyl477-1** | **0.0105** | **Congruent** |
| **Bsyl477-10** | **Esyl_Bsyl477-10** | **0.0094** | **Congruent** |
| Bsyl477-11 | Esyl_Bsyl477-11 | 0.0126 | Incongruent |
| **Bsyl477-3** | **Esyl_Bsyl477-3** | **0.0104** | **Congruent** |
| **Bsyl500-2** | **Esyl_Bsyl500-2** | **0.0109** | **Congruent** |
| **Bsyl500-3** | **Esyl_Bsyl500-3** | **0.0111** | **Congruent** |
| **Bsyl500-6** | **Esyl_Bsyl500-6** | **0.0106** | **Congruent** |
| **Bsyl501-1** | **Esyl_Bsyl501-1** | **0.0117** | **Congruent** |
| **Bsyl501-5** | **Esyl_Bsyl501-5** | **0.0106** | **Congruent** |
| **Bsyl501-6** | **Esyl_Bsyl501-6** | **0.0108** | **Congruent** |
| **Bsyl501-7** | **Esyl_Bsyl501-7** | **0.0115** | **Congruent** |
| **Bsyl502-2** | **Esyl_Bsyl502-2** | **0.0114** | **Congruent** |
| **Bsyl502-3** | **Esyl_Bsyl502-3** | **0.0113** | **Congruent** |
| **Bsyl502-4** | **Esyl_Bsyl502-4** | **0.0115** | **Congruent** |
| **Bsyl502-5** | **Esyl_Bsyl502-5** | **0.0099** | **Congruent** |
| **Bsyl505-2** | **Esyl_Bsyl505-2** | **0.0105** | **Congruent** |
| **Bsyl505-4** | **Esyl_Bsyl505-4** | **0.0108** | **Congruent** |
| **Bsyl505-6** | **Esyl_Bsyl505-6** | **0.0106** | **Congruent** |
| **Bsyl506-2** | **Esyl_Bsyl506-2** | **0.0107** | **Congruent** |
| **Bsyl506-3** | **Esyl_Bsyl506-3** | **0.0101** | **Congruent** |
| **Bsyl506-6** | **Esyl_Bsyl506-6** | **0.0107** | **Congruent** |
| **Bsyl508-2** | **Esyl_Bsyl508-2** | **0.0104** | **Congruent** |
| **Bsyl508-3** | **Esyl_Bsyl508-3** | **0.0103** | **Congruent** |
| **Bsyl508-6** | **Esyl_Bsyl508-6** | **0.0109** | **Congruent** |
| Bsyl54-3 | Esyl_Bsyl54-3 | 0.0131 | Incongruent |
| Bsyl54-9 | Esyl_Bsyl54-9 | 0.0134 | Incongruent |
| **Bsyl550-10** | **Esyl_Bsyl550-10** | **0.0102** | **Congruent** |
| **Bsyl550-1B** | **Esyl_Bsyl550-1B** | **0.0115** | **Congruent** |
| **Bsyl550-3B** | **Esyl_Bsyl550-3B** | **0.0109** | **Congruent** |
| **Bsyl550-8B** | **Esyl_Bsyl550-8B** | **0.0115** | **Congruent** |
| **Bsyl552-1** | **Esyl_Bsyl552-1** | **0.0101** | **Congruent** |
| **Bsyl552-2** | **Esyl_Bsyl552-2** | **0.0111** | **Congruent** |
| **Bsyl552-5** | **Esyl_Bsyl552-5** | **0.0112** | **Congruent** |
| **Bsyl553-2B** | **Esyl_Bsyl553-2B** | **0.0114** | **Congruent** |
| **Bsyl553-3B** | **Esyl_Bsyl553-3B** | **0.0103** | **Congruent** |
| Bsyl553-5B | Esyl_Bsyl553-5B | 0.0157 | Incongruent |
| **Bsyl554-1** | **Esyl_Bsyl554-1** | **0.0108** | **Congruent** |
| Bsyl554-6 | Esyl_Bsyl554-6 | 0.0122 | Incongruent |
| **Bsyl554-7** | **Esyl_Bsyl554-7** | **0.0108** | **Congruent** |
| **Bsyl555-1** | **Esyl_Bsyl555-1** | **0.0099** | **Congruent** |
| **Bsyl555-3B** | **Esyl_Bsyl555-3B** | **0.0100** | **Congruent** |
| **Bsyl555-5** | **Esyl_Bsyl555-5** | **0.0096** | **Congruent** |
| **Bsyl557-2** | **Esyl_Bsyl557-2** | **0.0102** | **Congruent** |
| **Bsyl557-7** | **Esyl_Bsyl557-7** | **0.0106** | **Congruent** |
| Bsyl59-1 | Esyl_Bsyl59-1 | 0.0122 | Incongruent |
| Bsyl59-4 | Esyl_Bsyl59-4 | 0.0125 | Incongruent |
| Bsyl59-5 | Esyl_Bsyl59-5 | 0.0123 | Incongruent |
| **Bsyl62-8** | **Esyl_Bsyl62-8** | **0.0102** | **Congruent** |
| Bsyl63-2A | Esyl_Bsyl63-2A | 0.0125 | Incongruent |
| **Bsyl63-2B** | **Esyl_Bsyl63-2B** | **0.0118** | **Congruent** |
| Bsyl63-4 | Esyl_Bsyl63-4 | 0.0129 | Incongruent |
| Bsyl63-5 | Esyl_Bsyl63-5 | 0.0126 | Incongruent |
| Bsyl72-3 | Esyl_Bsyl72-3 | 0.0123 | Incongruent |
| Bsyl72-4 | Esyl_Bsyl72-4 | 0.0125 | Incongruent |
| Bsyl73-1 | Esyl_Bsyl73-1 | 0.0123 | Incongruent |
| Bsyl73-3 | Esyl_Bsyl73-3 | 0.0123 | Incongruent |
| Bsyl73-4 | Esyl_Bsyl73-4 | 0.0125 | Incongruent |
