## Supplementary material for "A Palearctic divide, niche conservatism and host-fungal endophyte interactions shaped the phylogeography of the grass *Brachypodium sylvaticum*": Table S5

**Table S5.** Genetic diversity parameters at the Individual level for each *Brachypodium sylvaticum* complex sample calculated using plink2. For each accession, observed heterozygosity (Ho), expected heterozygosity (He) and the method-of-moments F coefficient are shown in the table.

| **Accession** | **Heterozigosity (Ho)** | **Heterozigosity (He)** | **F** |
| --- | --- | --- | --- |
| **Bbrev33H** | 0.074 | 0.166 | 0.556 |
| **Bbrev34H** | 0.083 | 0.128 | 0.349 |
| **Bglaucovirens** | 0.114 | 0.128 | 0.113 |
| **Bkurilense** | 0.077 | 0.128 | 0.394 |
| **Bmis66_3** | 0.069 | 0.145 | 0.524 |
| **Bmis66-4** | 0.08 | 0.138 | 0.424 |
| **Bmis67-1** | 0.067 | 0.149 | 0.553 |
| **Bmis67-2** | 0.059 | 0.158 | 0.628 |
| **Bspryginii** | 0.093 | 0.140 | 0.337 |
| **Bsyl29H** | 0.089 | 0.161 | 0.448 |
| **Bsyl30H** | 0.091 | 0.167 | 0.455 |
| **Bsyl31H** | 0.08 | 0.170 | 0.526 |
| **Bsyl32H** | 0.079 | 0.166 | 0.523 |
| **Bsyl35H** | 0.097 | 0.163 | 0.407 |
| **Bsyl36H** | 0.088 | 0.169 | 0.483 |
| **Bsyl37H** | 0.133 | 0.154 | 0.139 |
| **Bsyl38H** | 0.111 | 0.162 | 0.313 |
| **Bsyl39H** | 0.104 | 0.158 | 0.341 |
| **Bsyl41H** | 0.088 | 0.148 | 0.406 |
| **Bsyl466-13** | 0.073 | 0.146 | 0.502 |
| **Bsyl466-2** | 0.075 | 0.143 | 0.472 |
| **Bsyl466-7b** | 0.104 | 0.115 | 0.094 |
| **Bsyl467-10** | 0.092 | 0.103 | 0.113 |
| **Bsyl467-2** | 0.074 | 0.145 | 0.491 |
| **Bsyl467-7** | 0.075 | 0.139 | 0.461 |
| **Bsyl467-9b** | 0.086 | 0.148 | 0.421 |
| **Bsyl470-10** | 0.063 | 0.153 | 0.59 |
| **Bsyl470-4** | 0.075 | 0.153 | 0.507 |
| **Bsyl470-8b** | 0.084 | 0.149 | 0.439 |
| **Bsyl470-9** | 0.097 | 0.110 | 0.118 |
| **Bsyl476-10** | 0.089 | 0.102 | 0.127 |
| **Bsyl476-11** | 0.098 | 0.116 | 0.15 |
| **Bsyl476-14** | 0.092 | 0.104 | 0.114 |
| **Bsyl476-9** | 0.083 | 0.149 | 0.443 |
| **Bsyl477-10** | 0.087 | 0.140 | 0.383 |
| **Bsyl477-11** | 0.092 | 0.101 | 0.089 |
| **Bsyl477-1** | 0.096 | 0.131 | 0.266 |
| **Bsyl477-3** | 0.096 | 0.141 | 0.322 |
| **Bsyl500-2** | 0.082 | 0.141 | 0.422 |
| **Bsyl500-3** | 0.087 | 0.100 | 0.128 |
| **Bsyl500-6** | 0.094 | 0.108 | 0.13 |
| **Bsyl501-1** | 0.097 | 0.114 | 0.147 |
| **Bsyl501-5** | 0.093 | 0.103 | 0.094 |
| **Bsyl501-6** | 0.094 | 0.114 | 0.178 |
| **Bsyl501-7** | 0.09 | 0.104 | 0.132 |
| **Bsyl502-2** | 0.098 | 0.111 | 0.115 |
| **Bsyl502-3** | 0.096 | 0.108 | 0.111 |
| **Bsyl502-4** | 0.091 | 0.102 | 0.106 |
| **Bsyl502-5** | 0.098 | 0.146 | 0.328 |
| **Bsyl505-2** | 0.092 | 0.128 | 0.278 |
| **Bsyl505-4** | 0.086 | 0.099 | 0.123 |
| **Bsyl505-6** | 0.101 | 0.112 | 0.098 |
| **Bsyl506-2** | 0.099 | 0.121 | 0.182 |
| **Bsyl506-3** | 0.096 | 0.129 | 0.254 |
| **Bsyl506-6** | 0.094 | 0.104 | 0.101 |
| **Bsyl508-2** | 0.089 | 0.139 | 0.363 |
| **Bsyl508-3** | 0.094 | 0.146 | 0.36 |
| **Bsyl508-6** | 0.098 | 0.111 | 0.114 |
| **Bsyl54-3** | 0.092 | 0.108 | 0.148 |
| **Bsyl54-9** | 0.072 | 0.146 | 0.503 |
| **Bsyl550-10** | 0.094 | 0.103 | 0.085 |
| **Bsyl550-1B** | 0.094 | 0.122 | 0.23 |
| **Bsyl550-3B** | 0.093 | 0.103 | 0.1 |
| **Bsyl550-8b** | 0.095 | 0.137 | 0.305 |
| **Bsyl552-1** | 0.097 | 0.124 | 0.216 |
| **Bsyl552-2** | 0.079 | 0.140 | 0.438 |
| **Bsyl552-5** | 0.095 | 0.107 | 0.114 |
| **Bsyl553-2B** | 0.094 | 0.115 | 0.18 |
| **Bsyl553-3b** | 0.14 | 0.149 | 0.059 |
| **Bsyl553-5b** | 0.089 | 0.141 | 0.368 |
| **Bsyl554-1** | 0.079 | 0.139 | 0.434 |
| **Bsyl554-3B** | 0.077 | 0.138 | 0.443 |
| **Bsyl554-6** | 0.096 | 0.109 | 0.119 |
| **Bsyl554-7** | 0.096 | 0.127 | 0.243 |
| **Bsyl555-1** | 0.091 | 0.149 | 0.386 |
| **Bsyl555-3B** | 0.096 | 0.107 | 0.101 |
| **Bsyl555-5** | 0.098 | 0.110 | 0.105 |
| **Bsyl555-8** | 0.093 | 0.145 | 0.363 |
| **Bsyl557-2** | 0.105 | 0.122 | 0.142 |
| **Bsyl557-3** | 0.031 | 0.171 | 0.82 |
| **Bsyl557-7** | 0.079 | 0.145 | 0.457 |
| **Bsyl557-8** | 0.024 | 0.173 | 0.863 |
| **Bsyl59-1** | 0.077 | 0.140 | 0.447 |
| **Bsyl59-2** | 0.088 | 0.102 | 0.134 |
| **Bsyl59-4** | 0.083 | 0.152 | 0.456 |
| **Bsyl59-5** | 0.073 | 0.143 | 0.488 |
| **Bsyl62-8** | 0.097 | 0.144 | 0.329 |
| **Bsyl63-2A** | 0.079 | 0.139 | 0.431 |
| **Bsyl63-2B** | 0.092 | 0.103 | 0.111 |
| **Bsyl63-4** | 0.075 | 0.144 | 0.482 |
| **Bsyl63-5** | 0.084 | 0.132 | 0.365 |
| **Bsyl72-1** | 0.099 | 0.130 | 0.242 |
| **Bsyl72-3** | 0.103 | 0.124 | 0.167 |
| **Bsyl72-4** | 0.091 | 0.133 | 0.316 |
| **Bsyl73-1** | 0.09 | 0.135 | 0.333 |
| **Bsyl73-3** | 0.09 | 0.132 | 0.321 |
| **Bsyl73-4** | 0.093 | 0.129 | 0.279 |
