## Supplementary material for "A Palearctic divide, niche conservatism and host-fungal endophyte interactions shaped the phylogeography of the grass *Brachypodium sylvaticum*": Table S1

**Table S1.** List of the 94 *Brachypodium sylvaticum* complex samples (85 *B. sylvaticum s. s., 2 B. breviglume, 1 B. glaucovirens, 1 B. kurilense, 4 B. miserum, 1 B. spryginii*) included in the phylogeographic study of the complex. Information on taxon name, population code, sampling size (N), geographic location, georeferenced coordinates, and ENA accession code is provided in the table, and that on samples from seed germplasms [RIKEN, Japanese Center for Sustainable Resource Science] or herbarium specimens [B (Berlin), M (München), LD (Lund), VLA (Vladivostok)] under Location. The remaining samples were collected in the field. All studied *B. sylvaticum* complex samples are diploids. Taxonomic identities are based in the taxonomic works of Keng (1982), Probatova (1985), Schippmann (1991), Catalán et al. (2016), Tzvelev (2015), Tzvelev & Probatova (2019). Samples corresponding to the outgroups used for this study (*B. stacei, B. distachyon, B. arbuscula*, *B. pinnatum*) are listed at the bottom of the table. Newly studied accessions are indicated in bold.

| **Taxon** | **Code / Pop** | **N** | **Location** | **Latitude** | **Longitude** | **Accession** |
| --- | --- | --- | --- | --- | --- | --- |
| *B. breviglume* | Bbrev33H | 1 | Tibet: Gongbogyamda, LD135398 | 30.359 | 68.997 | **ERS16317778** |
| *B. breviglume* | Bbrev34H | 1 | Pakistan: Hazara, Paras-Shogran, M0175636 | 31.795 | 83.465 | **ERS16317779** |
| *B. glaucovirens* | Bglaucovirens | 1 | Greece: Crete, B3151 | 35.256 | 24.828 | **ERS16317783** |
| *B. kurilense* | Bkurilense | 1 | Russia: Kuril Islands, Iturup, VLA 1625 | 45.063 | 147.797 | **ERS16317785** |
| *B. miserum* | Bmis66 | 2 | Japan: Honshu, Kanagawa | 35.401 | 139.046 | ERS24700336, ERS24700337 |
| *B. miserum* | Bmis67 | 2 | Japan: Honshu, Sendai, RIKEN | 38.315 | 140.604 | **ERS16317790,** ERS24700339 |
| *B. spryginii* | Bspryginii | 1 | Russia: Krasnodarskii Krai, VLA 11652 | 45.08 | 38.912 | **ERS16318043** |
| *B. silvaticum* | Bsyl29H | 1 | Greece: Evrytania, NO Viniani. B100281411 | 38.988 | 21.687 | **ERS16317844** |
| *B. silvaticum* | Bsyl30H | 1 | Denmark: Zealand, Vrangeskov, north of Ringsted. B100566906 | 55.489 | 11.777 | **ERS16317845** |
| *B. silvaticum* | Bsyl31H | 1 | Germany: Bayern, Regierungsbezirk Oberpfalz, Landkreis Regensburg. M-0177011 | 48.215 | 9.41 | **ERS16317846** |
| *B. silvaticum* | Bsyl32H | 1 | Russia: Kaluga, distr. Kondrovo. Vallis fluv. Ugra. M-01769847 | 54.536 | 36.116 | ERS24700341 |
| *B. silvaticum* | Bsyl35H | 1 | Greece: N Ano Vlasia, 3.6.2010, Eichenwald. Mus. Bot. Berol. B100404658 | 38.018 | 21.937 | ERS24700342 |
| *B. silvaticum* | Bsyl36H | 1 | Iran: Luristan, Bisheh. LD-1810427 | 33.515 | 47.77 | **ERS16317847** |
| *B. silvaticum* | Bsyl37H | 1 | Iran: Shahpasand. Museum Botanicum Hauniense 5/2017 103 | 35.179 | 50.127 | ERS24700344 |
| *B. silvaticum* | Bsyl38H | 1 | Turkey: Hatay, Amanos Mts. Between Dörtyol and Daz Dagy. B100565016 | 36.834 | 36.203 | ERS24700346 |
| *B. silvaticum* | Bsyl39H | 1 | Turkey: Samsun, 16.6 Km East Bafra. LD-1800582 | 41.282 | 36.291 | ERS24700347 |
| *B. silvaticum* | Bsyl41H | 1 | Afganistan: Kabul, prov. Istalif gardens. MSB-173320 | 34.484 | 69.297 | ERS24700348 |
| *B. silvaticum* | Bsyl466 | 3 | Spain: Huesca, S. Guara | 42.245 | -0.242 | ERS24700372, ERS24700374, ERS24700376 |
| *B. silvaticum* | Bsyl467 | 4 | Spain: Huesca, Bespen | 42.061 | -0.118 | **ERS16317852,** ERS24700377, ERS24700378, ERS24700379 |
| *B. silvaticum* | Bsyl470 | 3 | Spain: San Sebastian, Cristina Enea | 43.316 | -1.974 | **ERS16317853,** ERS24700380, ERS24799074 |
| *B. silvaticum* | Bsyl476 | 4 | France: Hautes Pyrenees, Barbazan | 43.033 | -0.623 | **ERS16317854**, ERS24700382, ERS24700383, ERS24700385 |
| *B. silvaticum* | Bsyl477 | 4 | Spain: Lleida, Bausen | 42.832 | 0.718 | ERS24700386, ERS24700387 |
| *B. silvaticum* | Bsyl500 | 3 | France: Nans les Pins | 43.338 | 5.741 | ERS24700392, ERS24700393, ERS24700394 |
| *B. silvaticum* | Bsyl501 | 4 | Frances: Alpes Maritime | 43.664 | 7.036 | **ERS16317855,** ERS24700395, ERS24700397, ERS24700399 |
| *B. silvaticum* | Bsyl502 | 4 | France: Le Cannet des Maures | 43.368 | 6.369 | ERS24700401, ERS24700402, ERS24700403, ERS24700405 |
| *B. silvaticum* | Bsyl505 | 3 | France: Haerault, Causse de la Selle | 43.803 | 3.644 | ERS24700407, ERS24700408, ERS24700409 |
| *B. silvaticum* | Bsyl506 | 3 | France: Haerault, Saint Jean de Fos | 43.712 | 3.544 | **ERS16317856,** ERS24700410, ERS24700411 |
| *B. silvaticum* | Bsyl508 | 3 | France: Aude, Villeseque des Corbieres | 43.041 | 2.856 | ERS24700412, ERS24700413, ERS24700414 |
| *B. silvaticum* | Bsyl54 | 2 | Morocco: Rif Mts. Bab Barret | 35.019 | -5.01 | ERS24700350, ERS24700352 |
| *B. silvaticum* | Bsyl550 | 4 | Spain: Salamanca, S. de Francia | 40.504 | -6.176 | ERS24700415, ERS24700416, ERS24700418, ERS24700419 |
| *B. silvaticum* | Bsyl552 | 3 | Spain: Cadiz, Tarifa | 36.15 | -5.628 | **ERS16317858,** ERS24700420, ERS24700421 |
| *B. silvaticum* | Bsyl553 | 3 | Spain: Malaga, Cartama | 36.718 | -4.605 | **ERS24700424, ERS24700426, ERS24700428** |
| *B. silvaticum* | Bsyl554 | 4 | Spain: Malaga, Canillas del Aceituno | 36.891 | -4.064 | **ERS16317859,** ERS24700429, ERS24700430, ERS24700431 |
| *B. silvaticum* | Bsyl555 | 4 | Spain: Granada, S. Nevada | 37.13 | -3.448 | ERS24700432, ERS24700433, ERS24700434, ERS24700435 |
| *B. silvaticum* | Bsyl557 | 2 | Spain: Girona, Ampurdan, Maçanet de Cabrenys | 42.389 | 2.744 | ERS24700436, ERS24700438 |
| *B. silvaticum* | Bsyl59 | 4 | Morocco: Middle Atlas, Bab Bou Idir | 33.532 | -4.685 | ERS24700353, ERS24700355, ERS24700356, ERS24700357 |
| *B. silvaticum* | Bsyl62 | 1 | Morocco: Middle Atlas, Ifrane NP | 33.518 | -5.169 | ERS24700358 |
| *B. silvaticum* | Bsyl63 | 4 | Morocco: Mulay-Idriss | 34.057 | -5.525 | **ERS16317849** |
| *B. silvaticum* | Bsyl72 | 3 | UK: Cambridgeshire, Dray Dayton | 52.241 | 0.015 | ERS24700362, ERS24700363, ERS24700364 |
| *B. silvaticum* | Bsyl73 | 3 | UK: East Suffolk, Stowmarket | 52.192 | 1.134 | ERS24700366, ERS24700368, ERS24700370 |
| *B. stacei* | ABR114 | 1 | Spain: Majorca | 39.534 | 2.858 | **SRS844132** |
| *B. stacei* | Bsta_ECI | 1 | Israel: Evolution Canyon I | 32.710 | 34.970 | **SRR17303409** |
| *B. distachyon* | Bd21 | 1 | Iraq: near Salakudin | 33.223 | 43.679 | **SRS1615350** |
| *B. distachyon* | LPA3.2 | 1 | Italy: Sicily, Lago di Piana degli Albanesi | 37.985 | 13.287 | **SRS1670020** |
| *B. arbuscula* | Barb502 | 1 | Spain: Canary Islands, La Gomera | 28.169 | -17.249 | **ERS16317771** |
| *B. arbuscula* | Barb405 | 1 | Spain: Canary Islands, Tenerife | 28.343 | -16.878 | **ERS16317772** |
| *B. pinnatum* | Bpin502 | 1 | Iraq: USDA_PI 185135 | 33.223 | 43.679 | **ERS16317809** |
| *B. pinnatum* | Bpin505 | 1 | Norway: USDA_PI 345964 | 60.446 | 8.428 | **ERS16317810** |
| *B. pinnatum* | Bpin515 | 1 | Kazakhstan: USDA PI 440176 | 48.019 | 66.923 | **ERS16317812** |
