## Supplementary material for "A Palearctic divide, niche conservatism and host-fungal endophyte interactions shaped the phylogeography of the grass *Brachypodium sylvaticum*": Table S2

**Table S2.** Summary statistics of the genomic data for the *Brachypodium sylvaticum* complex samples under study and their *Epichloë* endophytes. Information is provided on taxon name, population code, total number of Illumina raw reads generated from genome skimming analysis (RawReads), number of *Brachypodium* reads that successfully mapped to the *B. sylvaticum* reference genome Ain1 (BsylReads), number of *B. sylvaticum* protein-coding genes with at least 90% of sequence positions covered at a minimum depth of 10 (Genes90), assembled plastome length (Plastome), number of *Epichloë* reads mapped against *Epichloë* reference genomes (EpichloëReads), percentage of fungal endophyte rDNA 35S cistron recovered (Cistron%), and BLAST result (Blast) indicating whether the recovered fungal cistron belongs to *Epichloë* *sylvatica* (Yes) or to other *Epichloë* endophyte (No).

| **Taxon** | **Code** | **RawReads** | ***Brachypodium sylvaticum*** | | | ***Epichloë sylvatica*** | | |
| --- | --- | --- | --- | --- | --- | --- | --- | --- |
|  |  |  | **BsylReads** | **Genes90** | **Plastome** | **EpichloëReads** | **Cistron%** | **Blast** |
| *B. breviglume* | Bbrev33H | 33191622 | 28619108 | 16396 | 136,523 | 30342 | 63.04% | No |
| *B. breviglume* | Bbrev34H | 16686598 | 15208453 | 10892 | 136,312 | 35138 | 73.97% | No |
| *B. glaucovirens* | Bglaucovirens | 25016658 | 22593832 | 15430 | 136251 | 9070 | 12.54% | No |
| *B. kurilense* | Bkurilense | 11563438 | 10246338 | 8020 | 136,369 | 18089 | 68.49% | Yes |
| *B. misserum* | Bmis66-3 | 26069552 | 23077113 | 14164 | 136,326 | 10145 | 54.66% | No |
| *B. misserum* | Bmis66-4 | 15534082 | 21853030 | 14399 | 136,355 | 6563 | 4.94% | No |
| *B. misserum* | Bmis67-1 | 36235534 | 30409573 | 14434 | 136,350 | 15545 | 3.96% | No |
| *B. misserum* | Bmis67-2 | 26432496 | 23392396 | 15010 | 136,352 | 10596 | 38.24% | No |
| *B. spryginii* | Bspryginii | 10904196 | 17861035 | 10777 | 136,073 | 56508 | 86.60% | Yes |
| *B. sylvaticum* | Bsyl29H | 34829662 | 31686322 | 18580 | 136332 | 198413 | 77.89% | Yes |
| *B. sylvaticum* | Bsyl30H | 43089680 | 40184481 | 21937 | 136540 | 146004 | 76.30% | Yes |
| *B. sylvaticum* | Bsyl31H | 40115538 | 37330839 | 19997 | 136561 | 331211 | 81.97% | Yes |
| *B. sylvaticum* | Bsyl32H | 41607486 | 31448835 | 17727 | 136526 | 265881 | 82.60% | Yes |
| *B. sylvaticum* | Bsyl35H | 38710164 | 34693724 | 19201 | 136332 | 374559 | 82.60% | Yes |
| *B. sylvaticum* | Bsyl36H | 48966188 | 45697368 | 24287 | 136322 | 159039 | 81.09% | Yes |
| *B. sylvaticum* | Bsyl37H | 34857760 | 24410216 | 16377 | 136490 | 148592 | 71.38% | Yes |
| *B. sylvaticum* | Bsyl38H | 37856010 | 33127059 | 18533 | 136332 | 151170 | 74.57% | Yes |
| *B. sylvaticum* | Bsyl39H | 32417792 | 29397988 | 18227 | 136331 | 102031 | 72.96% | Yes |
| *B. sylvaticum* | Bsyl41H | 33512792 | 26128113 | 11532 | 136311 | 138552 | 76.16% | Yes |
| *B. sylvaticum* | Bsyl466-13 | 41827070 | 36979700 | 16541 | 136332 | 564201 | 86.72% | Yes |
| *B. sylvaticum* | Bsyl466-2 | 38991340 | 34544848 | 16407 | 136332 | 193487 | 77.75% | Yes |
| *B. sylvaticum* | Bsyl466-7B | 21194800 | 19132740 | 15157 | 136339 | 87246 | 72.96% | Yes |
| *B. sylvaticum* | Bsyl467-10 | 14476330 | 13208274 | 12661 | 136333 | 40079 | 74.14% | Yes |
| *B. sylvaticum* | Bsyl467-2 | 39407178 | 35785772 | 16318 | 136343 | 195354 | 79.20% | Yes |
| *B. sylvaticum* | Bsyl467-7 | 37152968 | 32870468 | 16206 | 136320 | 178624 | 84.01% | Yes |
| *B. sylvaticum* | Bsyl467-9B | 35167754 | 31778564 | 16213 | 136337 | 115943 | 75.63% | Yes |
| *B. sylvaticum* | Bsyl470-4 | 41966418 | 35730235 | 16337 | 136332 | 130204 | 78.12% | Yes |
| *B. sylvaticum* | Bsyl470-8B | 36591956 | 33081221 | 16266 | 136334 | 90681 | 78.91% | Yes |
| *B. sylvaticum* | Bsyl470-9 | 16207000 | 14352274 | 13535 | 136333 | 54662 | 72.77% | Yes |
| *B. sylvaticum* | Bsyl476-10 | 12729210 | 11530781 | 10762 | 136332 | 63216 | 74.79% | Yes |
| *B. sylvaticum* | Bsyl476-11 | 19206152 | 17134405 | 14445 | 136337 | 86087 | 77.20% | Yes |
| *B. sylvaticum* | Bsyl476-14 | 13289450 | 12028115 | 11193 | 136350 | 81158 | 73.57% | Yes |
| *B. sylvaticum* | Bsyl476-9 | 37107122 | 33622113 | 16157 | 136337 | 129727 | 82.09% | Yes |
| *B. sylvaticum* | Bsyl477-1 | 28562278 | 25829858 | 15948 | 136333 | 85642 | 76.63% | Yes |
| *B. sylvaticum* | Bsyl477-10 | 29817608 | 26913810 | 15848 | 136338 | 115179 | 83.97% | Yes |
| *B. sylvaticum* | Bsyl477-11 | 14194404 | 12658369 | 12203 | 136350 | 65876 | 75.16% | Yes |
| *B. sylvaticum* | Bsyl477-3 | 35342516 | 31942448 | 16512 | 136332 | 104865 | 83.36% | Yes |
| *B. sylvaticum* | Bsyl500-2 | 34294702 | 31128141 | 16080 | 136334 | 103982 | 78.36% | Yes |
| *B. sylvaticum* | Bsyl500-3 | 13393326 | 12189524 | 11221 | 136333 | 68424 | 73.34% | Yes |
| *B. sylvaticum* | Bsyl500-6 | 17864626 | 16130482 | 14050 | 136310 | 56404 | 78.42% | Yes |
| *B. sylvaticum* | Bsyl501-1 | 18801458 | 17037987 | 14099 | 136332 | 104451 | 78.48% | Yes |
| *B. sylvaticum* | Bsyl501-5 | 14756232 | 13086715 | 12851 | 136332 | 150856 | 74.42% | Yes |
| *B. sylvaticum* | Bsyl501-6 | 21353818 | 19391621 | 14930 | 136351 | 64129 | 76.30% | Yes |
| *B. sylvaticum* | Bsyl501-7 | 15721834 | 14246150 | 13119 | 136332 | 63373 | 74.18% | Yes |
| *B. sylvaticum* | Bsyl502-2 | 18649728 | 16740846 | 14628 | 136332 | 83063 | 77.93% | Yes |
| *B. sylvaticum* | Bsyl502-3 | 17674700 | 15923072 | 14053 | 136333 | 77749 | 75.50% | Yes |
| *B. sylvaticum* | Bsyl502-4 | 14979082 | 13353934 | 12506 | 136332 | 120035 | 81.87% | Yes |
| *B. sylvaticum* | Bsyl502-5 | 41844720 | 38007281 | 16909 | 136333 | 110169 | 76.32% | Yes |
| *B. sylvaticum* | Bsyl505-2 | 26665078 | 24255751 | 15705 | 136342 | 83964 | 74.44% | Yes |
| *B. sylvaticum* | Bsyl505-4 | 12926000 | 11456470 | 10693 | 136342 | 59019 | 77.10% | Yes |
| *B. sylvaticum* | Bsyl505-6 | 19207084 | 17454765 | 14571 | 136333 | 67970 | 76.52% | Yes |
| *B. sylvaticum* | Bsyl506-2 | 22935592 | 20831179 | 15393 | 136332 | 61002 | 73.89% | Yes |
| *B. sylvaticum* | Bsyl506-3 | 30621490 | 27198453 | 15852 | 136333 | 74925 | 74.71% | Yes |
| *B. sylvaticum* | Bsyl506-6 | 15066976 | 13540913 | 13088 | 136351 | 54472 | 74.24% | Yes |
| *B. sylvaticum* | Bsyl508-2 | 33880076 | 30714754 | 16245 | 136333 | 103503 | 79.36% | Yes |
| *B. sylvaticum* | Bsyl508-3 | 41601618 | 36650626 | 16750 | 136334 | 208174 | 79.52% | Yes |
| *B. sylvaticum* | Bsyl508-6 | 18084480 | 16327635 | 14270 | 136333 | 68879 | 78.95% | Yes |
| *B. sylvaticum* | Bsyl54-9 | 39675328 | 34226860 | 16445 | 136315 | 121085 | 86.27% | Yes |
| *B. sylvaticum* | Bsyl54-3 | 15528142 | 13749954 | 12563 | 136452 | 83317 | 84.27% | Yes |
| *B. sylvaticum* | Bsyl550-10 | 15012626 | 13549327 | 13056 | 136348 | 68193 | 82.91% | Yes |
| *B. sylvaticum* | Bsyl550-1B | 24491776 | 21867035 | 15502 | 136349 | 69709 | 86.60% | Yes |
| *B. sylvaticum* | Bsyl550-3B | 14182836 | 12778277 | 12175 | 136349 | 41504 | 81.54% | Yes |
| *B. sylvaticum* | Bsyl550-8B | 32320354 | 29231275 | 16253 | 136348 | 82390 | 86.50% | Yes |
| *B. sylvaticum* | Bsyl552-1 | 24078706 | 21557572 | 15629 | 136332 | 138016 | 76.52% | Yes |
| *B. sylvaticum* | Bsyl552-2 | 33625706 | 30247060 | 16201 | 136335 | 127819 | 84.70% | Yes |
| *B. sylvaticum* | Bsyl552-5 | 16111992 | 14440880 | 13665 | 136335 | 95807 | 89.44% | Yes |
| *B. sylvaticum* | Bsyl553-2B | 18846652 | 16927723 | 14187 | 136334 | 94675 | 79.73% | Yes |
| *B. sylvaticum* | Bsyl553-3B | 43053436 | 38207967 | 16926 | 136336 | 175185 | 74.48% | Yes |
| *B. sylvaticum* | Bsyl553-5B | 30605374 | 27690781 | 15889 | 136348 | 139834 | 76.18% | Yes |
| *B. sylvaticum* | Bsyl554-1 | 38576144 | 33741904 | 16331 | 136333 | 126717 | 74.61% | Yes |
| *B. sylvaticum* | Bsyl554-3B | 36963004 | 32918856 | 16080 | 136337 | 15224 | 22.94% | No |
| *B. sylvaticum* | Bsyl554-6 | 17961536 | 16074859 | 14287 | 136321 | 94897 | 75.18% | Yes |
| *B. sylvaticum* | Bsyl554-7 | 26569440 | 23679390 | 15736 | 136336 | 110466 | 78.42% | Yes |
| *B. sylvaticum* | Bsyl555-1 | 42833128 | 38272906 | 16932 | 136331 | 188724 | 77.14% | Yes |
| *B. sylvaticum* | Bsyl555-3B | 16786190 | 15251540 | 14076 | 136339 | 165522 | 81.70% | Yes |
| *B. sylvaticum* | Bsyl555-5 | 18192534 | 16356801 | 14495 | 136339 | 67979 | 79.14% | Yes |
| *B. sylvaticum* | Bsyl555-8 | 39254442 | 35808426 | 16817 | 136328 | 121352 | 74.55% | Yes |
| *B. sylvaticum* | Bsyl557-2 | 22789514 | 20716506 | 15536 | 136327 | 89612 | 76.20% | Yes |
| *B. sylvaticum* | Bsyl557-7 | 38414974 | 34728940 | 16411 | 136332 | 220168 | 75.12% | Yes |
| *B. sylvaticum* | Bsyl59-1 | 33320104 | 29865241 | 16226 | 136336 | 193045 | 82.70% | Yes |
| *B. sylvaticum* | Bsyl59-2 | 13487444 | 12116012 | 11519 | 136336 | 41636 | 63.82% | Yes |
| *B. sylvaticum* | Bsyl59-4 | 47040888 | 43336588 | 17263 | 136336 | 148139 | 74.99% | Yes |
| *B. sylvaticum* | Bsyl59-5 | 37024556 | 33847526 | 16508 | 136335 | 136524 | 74.61% | Yes |
| *B. sylvaticum* | Bsyl62-8 | 38069342 | 34372005 | 16686 | 136332 | 171679 | 81.46% | Yes |
| *B. sylvaticum* | Bsyl63-2A | 34736564 | 30467933 | 16171 | 136330 | 209259 | 88.09% | Yes |
| *B. sylvaticum* | Bsyl63-2B | 14428404 | 12806555 | 12578 | 136332 | 45064 | 71.63% | Yes |
| *B. sylvaticum* | Bsyl63-4 | 36599890 | 32728482 | 16315 | 136329 | 112308 | 74.28% | Yes |
| *B. sylvaticum* | Bsyl63-5 | 31041492 | 26845294 | 15733 | 136335 | 155692 | 76.40% | Yes |
| *B. sylvaticum* | Bsyl72-1 | 27259616 | 24612443 | 15871 | 136328 | 92430 | 83.13% | Yes |
| *B. sylvaticum* | Bsyl72-3 | 24074628 | 21462065 | 15618 | 136333 | 101990 | 77.85% | Yes |
| *B. sylvaticum* | Bsyl72-4 | 28959836 | 25879378 | 15840 | 136333 | 159723 | 75.24% | Yes |
| *B. sylvaticum* | Bsyl73-1 | 29560274 | 26449176 | 16010 | 136337 | 71003 | 77.18% | Yes |
| *B. sylvaticum* | Bsyl73-3 | 27714626 | 24918680 | 15885 | 136337 | 92626 | 75.16% | Yes |
| *B. sylvaticum* | Bsyl73-4 | 26580862 | 23870981 | 15776 | 136332 | 141298 | 78.61% | Yes |
| Total |  | 2606923782 |  |  |  | 10684833 |  |  |
