## Supplementary material for "A Palearctic divide, niche conservatism and host-fungal endophyte interactions shaped the phylogeography of the grass *Brachypodium sylvaticum*": Table S3

**Table S3** Results of co-evolutionary ParaFit analysis for individual *Brachypodium sylvaticum* complex *– Epichloë* *sylvatica* associations detected over 1000 replicates. The F1 statistic quantifies the degree of phylogenetic congruence between each *Brachypodium* host plant and its associated *Epichloë* endophyte. The p-values indicate the statistical significance of each association, with p<0.05 values (highlighted in bold) supporting significant co-evolutionary relationships. Global ParaFit value and p-value are also indicated.

| **Global ParaFit Value:** 87,009,603 | | **P- Value:** 0.006 | |
| --- | --- | --- | --- |
| ***Brachypodium sylvaticum s. l.*** | ***Epichloë sylvatica*** | **F1_Stat** | **P-Value** |
| Bkurilense | Esyl_Bkurilense | 1642523 | 0.1631 |
| **Bspryginii** | **Esyl_Bspryginii** | **7546561** | **0.0029** |
| Bsyl29H | Esyl_Bsyl29H | 3824211 | 0.0544 |
| Bsyl30H | Esyl_Bsyl30H | -914326 | 0.7812 |
| **Bsyl31H** | **Esyl_Bsyl31H** | **9246550** | **0.0015** |
| **Bsyl32H** | **Esyl_Bsyl32H** | **9301931** | **0.0015** |
| Bsyl35H | Esyl_Bsyl35H | 3758048 | 0.0516 |
| Bsyl36H | Esyl_Bsyl36H | -919326 | 0.7755 |
| Bsyl37H | Esyl_Bsyl37H | 1638289 | 0.1637 |
| Bsyl38H | Esyl_Bsyl38H | 3813426 | 0.0548 |
| Bsyl39H | Esyl_Bsyl39H | 3631555 | 0.0553 |
| **Bsyl41H** | **Esyl_Bsyl41H** | **7867518** | **0.0027** |
| Bsyl466-13 | Esyl_Bsyl466-13 | -2673842 | 0.9525 |
| Bsyl466-2 | Esyl_Bsyl466-2 | 1668800 | 0.2540 |
| Bsyl466-7B | Esyl_Bsyl466-7B | 1674040 | 0.2535 |
| Bsyl467-10 | Esyl_Bsyl467-10 | 2105062 | 0.1578 |
| Bsyl467-2 | Esyl_Bsyl467-2 | -2269065 | 0.9350 |
| Bsyl467-7 | Esyl_Bsyl467-7 | -1679150 | 0.8884 |
| **Bsyl467-9B** | **Esyl_Bsyl467-9B** | **9308221** | **0.0014** |
| Bsyl470-4 | Esyl_Bsyl470-4 | -5357635 | 0.9998 |
| Bsyl470-8B | Esyl_Bsyl470-8B | -6685273 | 0.9993 |
| Bsyl470-9 | Esyl_Bsyl470-9 | -7663194 | 0.9998 |
| **Bsyl476-10** | **Esyl_Bsyl476-10** | **8074542** | **0.0024** |
| **Bsyl476-11** | **Esyl_Bsyl476-11** | **8791270** | **0.0017** |
| **Bsyl476-14** | **Esyl_Bsyl476-14** | **8769632** | **0.0017** |
| **Bsyl476-9** | **Esyl_Bsyl476-9** | **9069109** | **0.0015** |
| Bsyl477-1 | Esyl_Bsyl477-1 | 1212570 | 0.3397 |
| **Bsyl477-10** | **Esyl_Bsyl477-10** | **9189058** | **0.0015** |
| Bsyl477-11 | Esyl_Bsyl477-11 | 1785372 | 0.1915 |
| Bsyl477-3 | Esyl_Bsyl477-3 | 2919597 | 0.0852 |
| Bsyl500-2 | Esyl_Bsyl500-2 | 3792181 | 0.0560 |
| Bsyl500-3 | Esyl_Bsyl500-3 | 2094699 | 0.1879 |
| Bsyl500-6 | Esyl_Bsyl500-6 | 1246173 | 0.2955 |
| Bsyl501-1 | Esyl_Bsyl501-1 | -1310867 | 0.8436 |
| Bsyl501-5 | Esyl_Bsyl501-5 | 2170102 | 0.1631 |
| Bsyl501-6 | Esyl_Bsyl501-6 | 1485414 | 0.2288 |
| Bsyl501-7 | Esyl_Bsyl501-7 | -1306358 | 0.8430 |
| Bsyl502-2 | Esyl_Bsyl502-2 | -1517982 | 0.8638 |
| Bsyl502-3 | Esyl_Bsyl502-3 | -1634525 | 0.8740 |
| Bsyl502-4 | Esyl_Bsyl502-4 | -1546175 | 0.8668 |
| Bsyl502-5 | Esyl_Bsyl502-5 | 1333201 | 0.2362 |
| Bsyl505-2 | Esyl_Bsyl505-2 | 3457459 | 0.0678 |
| Bsyl505-4 | Esyl_Bsyl505-4 | 1911873 | 0.1665 |
| Bsyl505-6 | Esyl_Bsyl505-6 | 3789420 | 0.0563 |
| Bsyl506-2 | Esyl_Bsyl506-2 | 3325833 | 0.0666 |
| Bsyl506-3 | Esyl_Bsyl506-3 | 1475927 | 0.2502 |
| Bsyl506-6 | Esyl_Bsyl506-6 | 2069947 | 0.1928 |
| Bsyl508-2 | Esyl_Bsyl508-2 | 3157415 | 0.0788 |
| Bsyl508-3 | Esyl_Bsyl508-3 | 2655920 | 0.0862 |
| Bsyl508-6 | Esyl_Bsyl508-6 | 3715046 | 0.0531 |
| Bsyl54-3 | Esyl_Bsyl54-3 | -2569357 | 0.9467 |
| Bsyl54-9 | Esyl_Bsyl54-9 | -2302093 | 0.9404 |
| Bsyl550-10 | Esyl_Bsyl550-10 | 471109 | 0.4612 |
| Bsyl550-1B | Esyl_Bsyl550-1B | -2476390 | 0.9414 |
| Bsyl550-3B | Esyl_Bsyl550-3B | 469531 | 0.4631 |
| Bsyl550-8B | Esyl_Bsyl550-8B | -2039091 | 0.9202 |
| Bsyl552-1 | Esyl_Bsyl552-1 | 840185 | 0.4242 |
| Bsyl552-2 | Esyl_Bsyl552-2 | -1497010 | 0.8582 |
| Bsyl552-5 | Esyl_Bsyl552-5 | -1480194 | 0.8571 |
| Bsyl553-2B | Esyl_Bsyl553-2B | 2008111 | 0.2005 |
| Bsyl553-3B | Esyl_Bsyl553-3B | -1922511 | 0.9336 |
| **Bsyl553-5B** | **Esyl_Bsyl553-5B** | **9256623** | **0.0015** |
| Bsyl554-1 | Esyl_Bsyl554-1 | 1207299 | 0.3165 |
| Bsyl554-6 | Esyl_Bsyl554-6 | -1666379 | 0.8858 |
| Bsyl554-7 | Esyl_Bsyl554-7 | 1517541 | 0.2745 |
| Bsyl555-1 | Esyl_Bsyl555-1 | 1242509 | 0.3352 |
| Bsyl555-3B | Esyl_Bsyl555-3B | 2274458 | 0.1671 |
| Bsyl555-5 | Esyl_Bsyl555-5 | 2055848 | 0.1584 |
| Bsyl557-2 | Esyl_Bsyl557-2 | 1590467 | 0.2682 |
| Bsyl557-7 | Esyl_Bsyl557-7 | 3790019 | 0.0540 |
| Bsyl59-1 | Esyl_Bsyl59-1 | 2138099 | 0.1844 |
| Bsyl59-4 | Esyl_Bsyl59-4 | 1960416 | 0.1399 |
| Bsyl59-5 | Esyl_Bsyl59-5 | 1274013 | 0.2981 |
| Bsyl62-8 | Esyl_Bsyl62-8 | 2183850 | 0.1717 |
| Bsyl63-2A | Esyl_Bsyl63-2A | 301656 | 0.5551 |
| Bsyl63-2B | Esyl_Bsyl63-2B | 2239949 | 0.1137 |
| Bsyl63-4 | Esyl_Bsyl63-4 | 2253034 | 0.1039 |
| Bsyl63-5 | Esyl_Bsyl63-5 | 2185670 | 0.1783 |
| Bsyl72-3 | Esyl_Bsyl72-3 | -2229904 | 0.9365 |
| Bsyl72-4 | Esyl_Bsyl72-4 | -2222063 | 0.9358 |
| Bsyl73-1 | Esyl_Bsyl73-1 | -2086155 | 0.9352 |
| Bsyl73-3 | Esyl_Bsyl73-3 | -2086155 | 0.9357 |
| Bsyl73-4 | Esyl_Bsyl73-4 | -2570498 | 0.9519 |
